# How Minds Take Shape: Graph-ESN Reveals How Neural Ensembles Engineer Stable Representations

**DOI:** 10.1101/2025.06.11.659086

**Authors:** Krishnan Venkiteswaran, Arun Ram Ponnambalam

## Abstract

Deciphering how neural populations encode, integrate, and dynamically transform information to generate predictive neural dynamics remains a fundamental pursuit in systems neuroscience. Although information is processed and represented across distributed neural ensembles, a critical question remains: How do these populations accumulate information and interact over time to form stable, coherent representations? The cortex comprises ensembles of interacting populations that cooperate to enable thought. However, an open challenge is to map how each population evolves its internal context in response to incoming information and crucially, how this evolving context shapes what each population communicates to other populations. To address this, we employ a customized variant of the Graph Echo State Network (Graph ESN) architecture that involves specialized populations. This formulation enables the model to disentangle and represent multi-scale oscillatory patterns in neural data, offering a more biologically plausible and task-relevant alternative to traditional ESNs. By leveraging the rich hidden-state dynamics of this architecture, we illustrate how neural ensembles iteratively interact and converge toward stable, temporally evolving representations of information. We further investigate the distinct contributions of individual populations in this collaborative process of representational stabilization. Mapping this temporally unfolding, cooperative structure sheds light on the neural mechanisms underlying distributed representation and inter-population coordination—processes that may ultimately support organized cognition.

## 1 Introduction

Understanding how the brain processes information remains a fundamental neuroscience challenge, with neural populations forming intricate networks that collectively encode, transform, and integrate information for cognition, creating complex processing dynamics through heterogeneity and coordination (1). Our key questions: Can we map these populations’ precise contributions at high temporal resolution and determine the rules governing their interactions? Research shows brain regions process information dynamically and flexibly, with bidirectional signal flow through the sensorimotor hierarchy based on contextual demands (2), prompting us to investigate how these neural populations iteratively interact and effectively contribute to information processing as an integrated system.The cortex has been known to have many types of circuitry, and recent research has revealed that neural responses often contain unexpected oscillatory components, even for non-periodic behaviors (3). These oscillations emerge from specific connectivity patterns that create cycles capable of generating and sustaining rhythmic activity (4). Can we map these circuits from temporal data to see how they are used to iteratively reach a conclusion?

Traditional methods for analyzing neural interactions, such as Granger causality and transfer entropy, often fail to capture the complex, nonlinear, and multi-scale dynamics that characterize neural computation (5; 6). These limitations have prompted the development of more sophisticated models using neural networks (8), unified frameworks (9), and advanced information-theoretic metrics (11). However, these approaches typically do not account for the role of hidden internal states in neural representation (16; 17) or how these internal representations develop through interaction between multiple populations.

Recent evidence suggests that communication between neural populations is governed not just by instantaneous activity, but by internal dynamic states—”hidden context”—that accumulate history and modulatory influences (19; 20). These internal states shape both the content and timing of inter-population communication, suggesting that cognition arises from dynamic network interactions where evolving internal states determine information flow (26).

Echo State Networks (ESNs) offer a promising approach for modeling neural population dynamics due to their biological plausibility, interpretability, and computational efficiency (27; 28). They can mirror cortical microcircuitry through interconnected inhibitory and excitatory populations that generate oscillatory patterns similar to those observed in neural systems (29; 30). Graph ESNs extend this capability to complex network structures while maintaining computational efficiency (31; 32).

Our approach introduces a customized Graph ESN with several novel components:

- Specialized gamma cell units for frequency-specific processing
- Coupled resonator banks modeling realistic oscillatory dynamics
- Enhanced population attention mechanisms for cross-regional information exchange
- Curriculum learning for progressive tackling of complex gamma patterns

This architecture incorporates multi-scale processing with dedicated populations for specific gamma frequency bands (30-300Hz), enabling it to disentangle oscillatory patterns in neural data while capturing how populations evolve their internal context in response to incoming information (35; 36). This approach aligns with findings showing that learning involves evolution in neural population dynamics both within and across cortical regions (37; 38).

## 2 Methods

### 2.1 Data and Preprocessing

We utilized Local Field Potential (LFP) recordings from the Allen Mouse Neuropixel Visual Coding database’s drifting gratings dataset (1000Hz sampling rate). For optimal spatial coverage, we selected 64 electrodes using KMeans clustering. Continuous LFP recordings were segmented into 380ms windows (330ms context, 50ms prediction), with each window drawn from a single stimulus presentation.

Signals underwent robust scaling followed by spectral decomposition into six gamma sub-bands: Low (30-50Hz), Mid-Low (50-80Hz), Mid-High (80-120Hz), High (120-180Hz), Ultra-High I (180- 250Hz), and Ultra-High II (250-300Hz). We employed 8th-order Butterworth filters and extracted phase, amplitude, and instantaneous frequency information using enhanced Hilbert transforms with reflection padding to minimize edge artifacts.

### 2.2 Enhanced Graph-ESN Architecture

Our Graph Echo State Network (Graph-ESN) draws inspiration from cortical circuit organization, where distinct neural populations exhibit specialized roles in information processing (39; 40). The architecture includes:

#### 2.2.1 Population Structure

The model implements 8 specialized neural populations (140 neurons each, 1,120 total): six dedicated to specific gamma sub-bands and two serving integrative functions. Communication between populations follows a structured connectivity pattern implementing principles of cross-frequency coupling observed in cortical circuits (41):

- Stronger connections between adjacent frequency bands
- Asymmetric influence from higher to lower frequencies
- Privileged connections to and from integrator populations

#### 2.2.2 Oscillatory Dynamics

To accurately model gamma oscillations, we implemented coupled resonator banks tuned to specific frequency ranges:

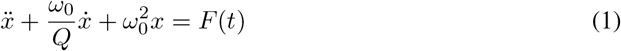

Our implementation distributes multiple resonators across each frequency band with varying Q-factors, allowing for precise modeling of narrow-band gamma oscillations (42).

#### 2.2.3 Excitatory-Inhibitory Balance

The model incorporates balanced excitatory-inhibitory dynamics with a 70/30 ratio:

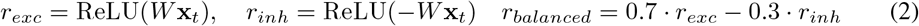

This ratio approximates the balance observed in cortical circuits, where excitatory pyramidal cells outnumber inhibitory interneurons by roughly 3:1 (29). This balance is crucial for generating realistic oscillatory dynamics and preventing runaway excitation.

### 2.3 Model Implementation

#### 2.3.1 Graph Reservoir Cell

The core component is the EnhancedGraphReservoirCell implementing the dynamics of interacting neural populations:

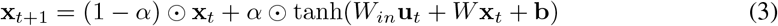

Our implementation extends standard ESNs with structured connectivity, population-specific activation functions, and balanced dynamics. The connectivity matrix *W* is structured to reflect the hierarchical organization of neural circuits, with specialized pathways between populations processing related frequency bands.

#### 2.3.2 Gamma-Specific Processing Units

For each gamma sub-band, we implemented GammaCellUnit modules integrating:

- Temporal integration of input signals
- Resonator dynamics for frequency-specific oscillations
- Adaptive gating mechanisms
- Phase and amplitude modulation

The dynamics are described by:

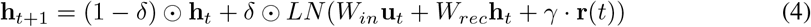

where **h**_*t*_ is the hidden state, *δ* is the decay factor, LN represents layer normalization, and **r**(*t*) is the resonator output. This formulation enables each unit to selectively respond to oscillatory patterns within its tuned frequency range while maintaining temporal coherence through recurrent connections.

#### 2.3.3 Decoders

We implemented specialized decoders for each frequency band integrating multi-layer neural networks, frequency-specific resonator banks, and amplitude/phase modulators. The output combines a base signal with modulated oscillatory components:

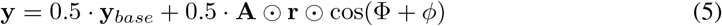

where **y**_*base*_ is the base signal, **A** is the amplitude envelope, **r** is the resonator output, and *ϕ* is the phase offset. This structure allows the model to reconstruct complex oscillatory patterns by combining frequency-specific components with appropriate phase relationships.

### 2.4 Training

We developed a specialized loss function combining time-domain and frequency-domain components:

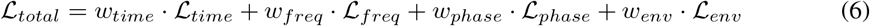

Each gamma frequency band was weighted progressively (1.0 for low gamma to 3.5 for ultra-high gamma). We implemented a three-phase curriculum learning approach (43) to progressively tackle more complex gamma patterns:

- Phase 1: Focus on lower gamma (30-80Hz)
- Phase 2: Focus on mid-range gamma (50-180Hz)
- Phase 3: Focus on higher gamma (120-300Hz)

Training used AdamW optimizer (44) with learning rate 2e-4, weight decay 2e-5, gradient clipping 0.8, and Stochastic Weight Averaging (45) for improved generalization.

### 2.5 Analysis of Neural Ensemble Dynamics

To analyze how neural ensembles engineer stable representations, we employed multiple techniques:

#### 2.5.1 Trajectory Analysis

We used delay embedding to visualize electrode signal trajectories in phase space (46):

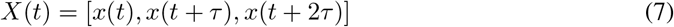

where *τ* = *context*_*size/*25. This approach identified attractor-like dynamics and state transitions.

#### 2.5.2 Influence Quantification

We measured cross-electrode influences using lagged correlation analysis:

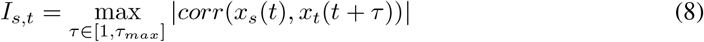

revealing directional information flow between electrodes and key network hubs (47).

#### 2.5.3 Context Vector Evolution

We applied PCA to model hidden states to reveal key transition points and stable attractors in context representation (48), providing insights into information integration over time.

#### 2.5.4 Rule Extraction

We extracted interpretable rules from hidden state dynamics by:

- Identifying significant state changes
- Detecting temporal relationships between changes
- Categorizing relationship types
- Quantifying relationship reliability

This approach discovered computational principles underlying neural ensemble dynamics (49).

## 3 Results and Discussion

### 3.1 Enhanced Graph-ESN Captures Gamma Oscillatory Dynamics

Our Graph-ESN model with specialized gamma populations demonstrates superior performance for neural time series prediction compared to traditional approaches. As shown in Table 1, the enhanced Graph-ESN achieves the lowest loss values across training and validation sets, outperforming standard recurrent architectures including RNNs, LSTMs, GRUs, and Transformers.

**Table 1:** Model Performance Comparison.

| Model | Parameters | Training Loss | Validation Loss |
| --- | --- | --- | --- |
| RNN | 4,218,464 | 0.1980 | 0.1978 |
| LSTM | 10,867,808 | 0.1985 | 0.1980 |
| GRU | 8,651,360 | 0.1970 | 0.1963 |
| Transformer | 2,671,904 | 0.1965 | 0.1960 |
| PINN | 3,777,440 | 0.1980 | 0.1975 |
| TCN | 2,670,176 | 0.1975 | 0.1969 |
| WaveletNN | 6,052,192 | 0.1972 | 0.1968 |
| **Graph-ESN (Ours)** | **1,120,000** | **0.0244** | **0.0254** |

Importantly, our model achieves this performance while requiring significantly fewer parameters (1.12M vs. 2.67M-10.87M), demonstrating its parameter efficiency. The decomposition of loss components reveals the primary challenge lies in phase prediction (phase loss: 0.8176), while time-domain (0.0485), frequency-domain (0.0003), and envelope prediction (0.0002) show strong performance.

Our frequency-specific analysis reveals clear electrode-level contributions to different gamma subbands. The model captures correlations ranging from 0.0390 in low gamma (30-50Hz) to 0.0311 in ultra-high gamma I (180-250Hz), with weaker correlations in mid-high gamma (80-120Hz, - 0.0020) and ultra-high gamma II (250-300Hz, 0.0088). These band-specific correlations suggest different functional roles for distinct gamma frequency ranges, with low gamma showing the strongest representation in our dataset, consistent with its established role in local information processing.

Figure 1 reveals the distinct trajectories of each specialized population in our model. Each population forms characteristic patterns in its state space, reflecting its specialized processing role. Low, mid, and high gamma populations (top row) show more chaotic dynamics with multiple branches and convergence points, consistent with their role in processing specific frequency bands. In contrast, the integrator and context populations (bottom row) exhibit more structured trajectories with clear paths, suggesting their role in distilling information from specialized populations into coherent representations. Notably, the context population shows the highest explainable variance in its first two principal components (0.46, 0.13), indicating its role in maintaining stable, lower-dimensional representations.

**Figure 1:**
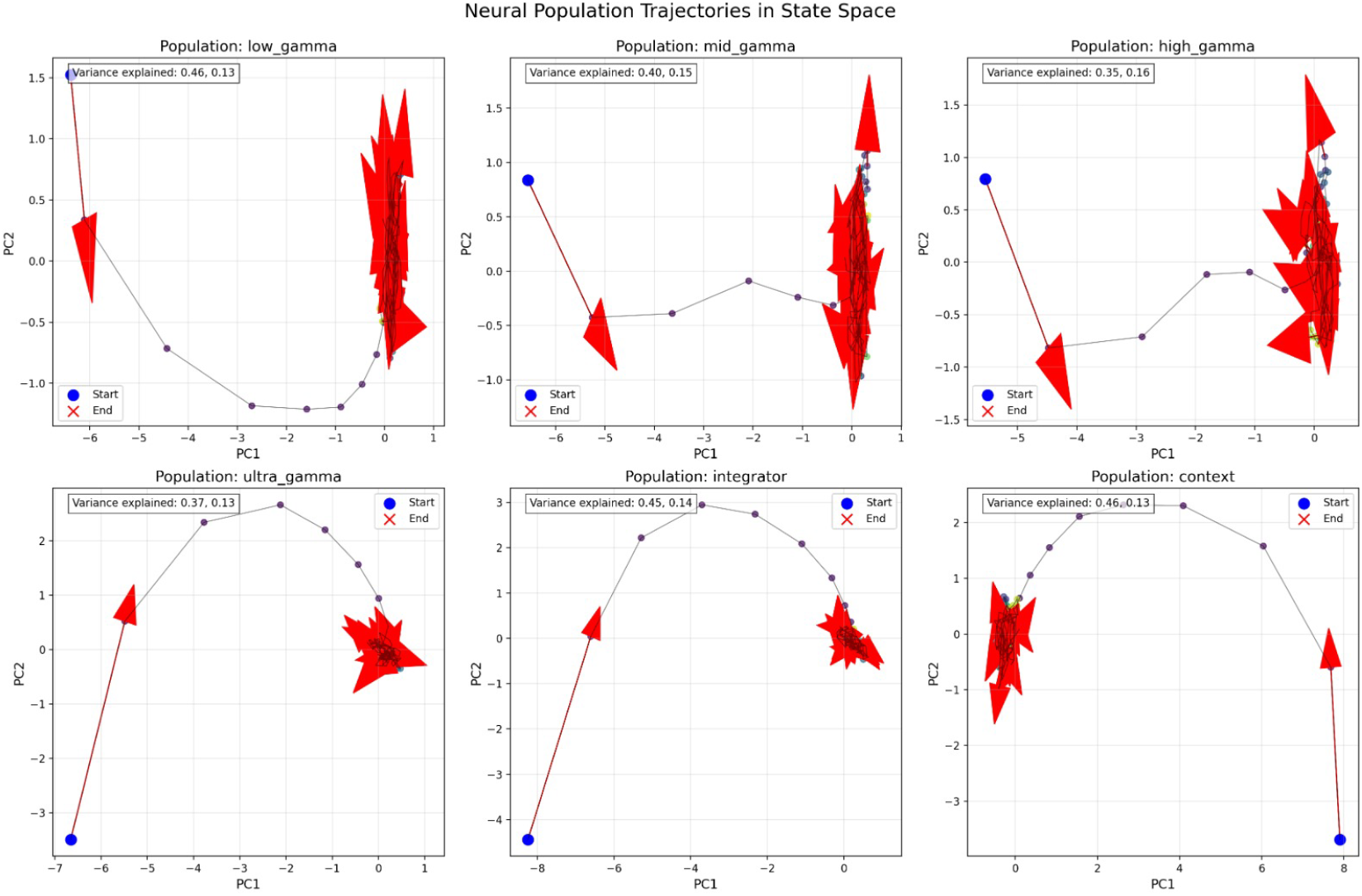
Neural population trajectories in state space. Each subpanel shows a different specialized population’s trajectory in a 2D projection of its hidden state space (PC1 vs. PC2). Red arrows indicate flow direction, blue dots mark starting points, and orange crosses mark endpoints. Numbers in top-left corners show variance explained by the first two principal components. Note how each specialized population forms distinct trajectory patterns, with integrator and context populations showing more organized dynamics compared to frequency-specific populations.

The variation across gamma bands and population types demonstrates the importance of multi-scale, specialized processing in neural representation. This supports the hypothesis that different neural populations serve distinct computational functions within neural circuits (50), with specialized gamma populations capturing frequency-specific information while integrator populations consolidate this information into coherent representations.

### 3.2 Electrode Trajectories and Computational Principles of Neural Ensembles

Figure 2 illustrates how individual electrode signals form distinct trajectories in delay-embedded space. This visualization reveals the complex temporal dynamics of neural activity, with each electrode tracing a unique path through the state space. The trajectories show evidence of attractor-like behavior, where the system’s dynamics are pulled toward specific regions of state space before transitioning to new configurations. The intertwined nature of these trajectories demonstrates how neural populations maintain coherent yet distinct information patterns. Notably, some electrodes (e.g., E56) exhibit more expansive trajectories, suggesting they explore a wider range of states and potentially play more diverse functional roles in information processing.

**Figure 2:**
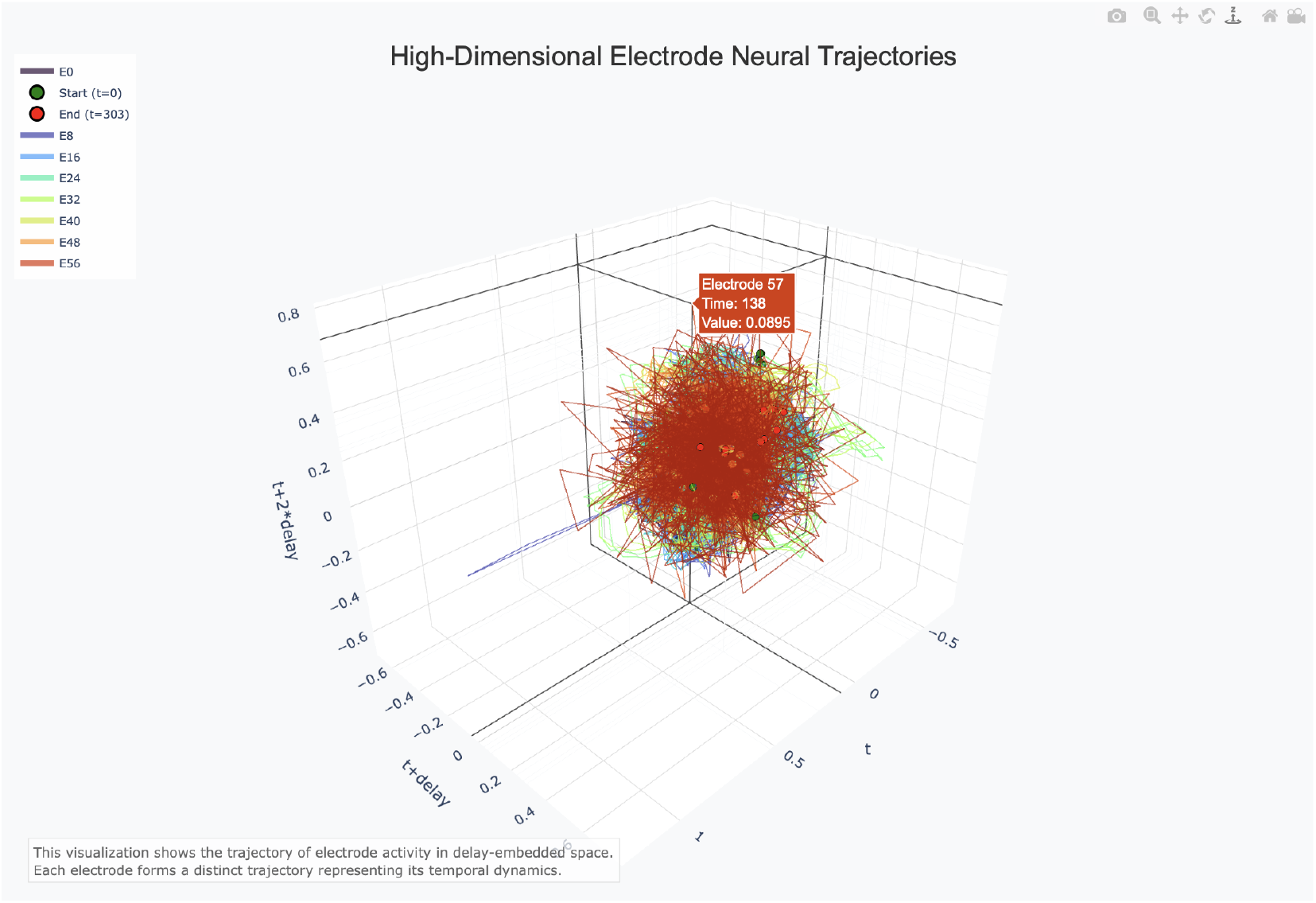
High-dimensional electrode neural trajectories in delay-embedded space. Each electrode forms a distinct trajectory representing its temporal dynamics. The color-coded paths show how different electrodes (E0, E8, E16, etc.) evolve through state space, revealing complex attractor-like patterns. Start (green) and end (red) points mark the beginning and conclusion of the recording.

The Neural Knowledge Graph (Figure 3), provides insight into the transformation relationships between electrode hidden states. This model-agnostic representation reveals the computational principles governing how distributed neural populations interact to form coherent representations. The knowledge graph identifies three primary transformation types: dimension expansion (blue edges), where electrodes amplify pattern complexity; dimension compression (orange edges), where electrodes extract simplified features; and mixed transformations (purple edges) that perform more complex operations. Notably, key electrodes (E13, E15, E18, E21, etc.) form distinct computational clusters, suggesting functional specialization within the neural ensemble.Analysis of the knowledge graph reveals that certain electrodes (particularly E23, E25, and E28) drive major shifts in the unified system state across multiple time points (shifts 1-3). These electrodes appear to serve as computational control nodes, initiating significant transitions in the representational space. This finding supports the hypothesis that neural computation relies on specific trigger populations that coordinate broader state transitions.

**Figure 3:**
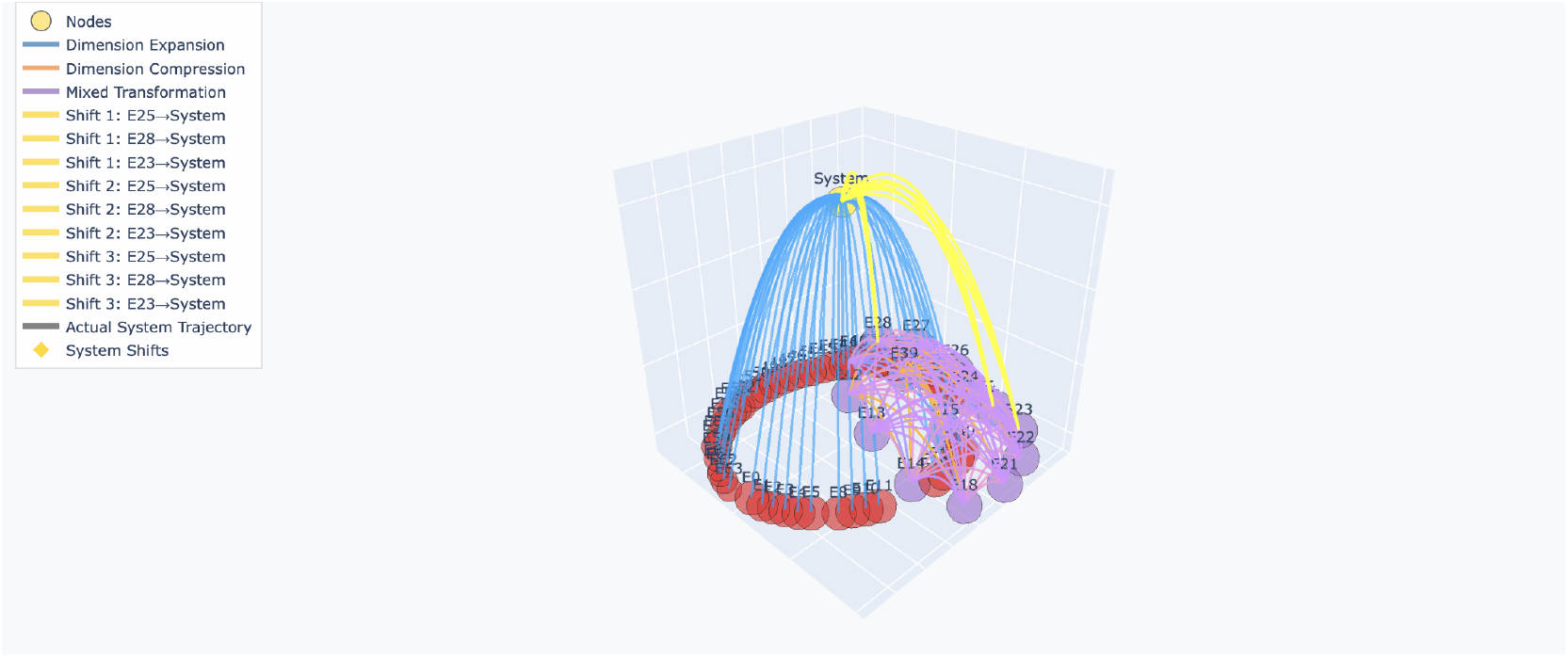
Electrode-agnostic knowledge graph showing transformation relationships between electrode hidden states. This model-derived graph reveals three primary transformation types: dimension expansion (blue), dimension compression (orange), and mixed transformations (purple). Yellow paths highlight major shifts where specific electrodes (E23, E25, E28) drive significant transitions in the unified system state. Node size represents influence magnitude.

By analyzing the Neural Logic Gate Knowledge Graph (Figure 3), we can extract quantitative rules governing neural ensemble dynamics. The graph reveals three primary classes of transformations:

1. **Dimension Expansion** (blue edges): These transformations amplify pattern complexity and typically propagate from specialized input electrodes (E20-E38) to intermediate processing nodes. These connections represent operations where simple input patterns are elaborated into more complex feature representations.
2. **Dimension Compression** (orange edges): These transformations perform feature extraction, reducing complex patterns to simpler representations. These are prevalent between intermediate electrodes (E15-E22) and the central system node, suggesting a consolidation of distributed information.
3. **Mixed Transformations** (purple edges): These operations perform more complex, nonlinear transformations that neither purely expand nor compress dimensionality. Instead, they appear to transform information across different representational bases, suggesting operations similar to basis rotations or nonlinear feature extraction.

Our analysis identifies critical state-transition pathways where specific electrodes (E23, E25, E28) drive major system state shifts, revealing how neural ensembles use different electrode combinations for distinct computational operations at various processing stages. These operations parallel cortical processing strategies: dimension expansion mirrors early sensory feature transformation, compression reflects integrative information consolidation, and mixed transformations resemble associative nonlinear processing—suggesting common computational principles across diverse cortical populations. This structured, rule-based mechanism enables neural robustness despite local variability through coordinated activity following specific transformation rules rather than homogeneous changes, creating a representation strategy balancing noise robustness with encoding flexibility.

### 3.3 Mechanisms of Representational Stabilization

Figure 4 demonstrates how distributed neural ensembles collectively form stable representations through iterative interactions. The PCA trajectory captures 33.9% of the variance in the highdimensional state space, revealing clear pattern formation with distinct phases indicated by color progression. Critical transition points (marked by yellow diamonds) correspond to major shifts in representation structure, where the neural ensemble reconfigures its internal state to encode new information or transition between processing modes.

**Figure 4:**
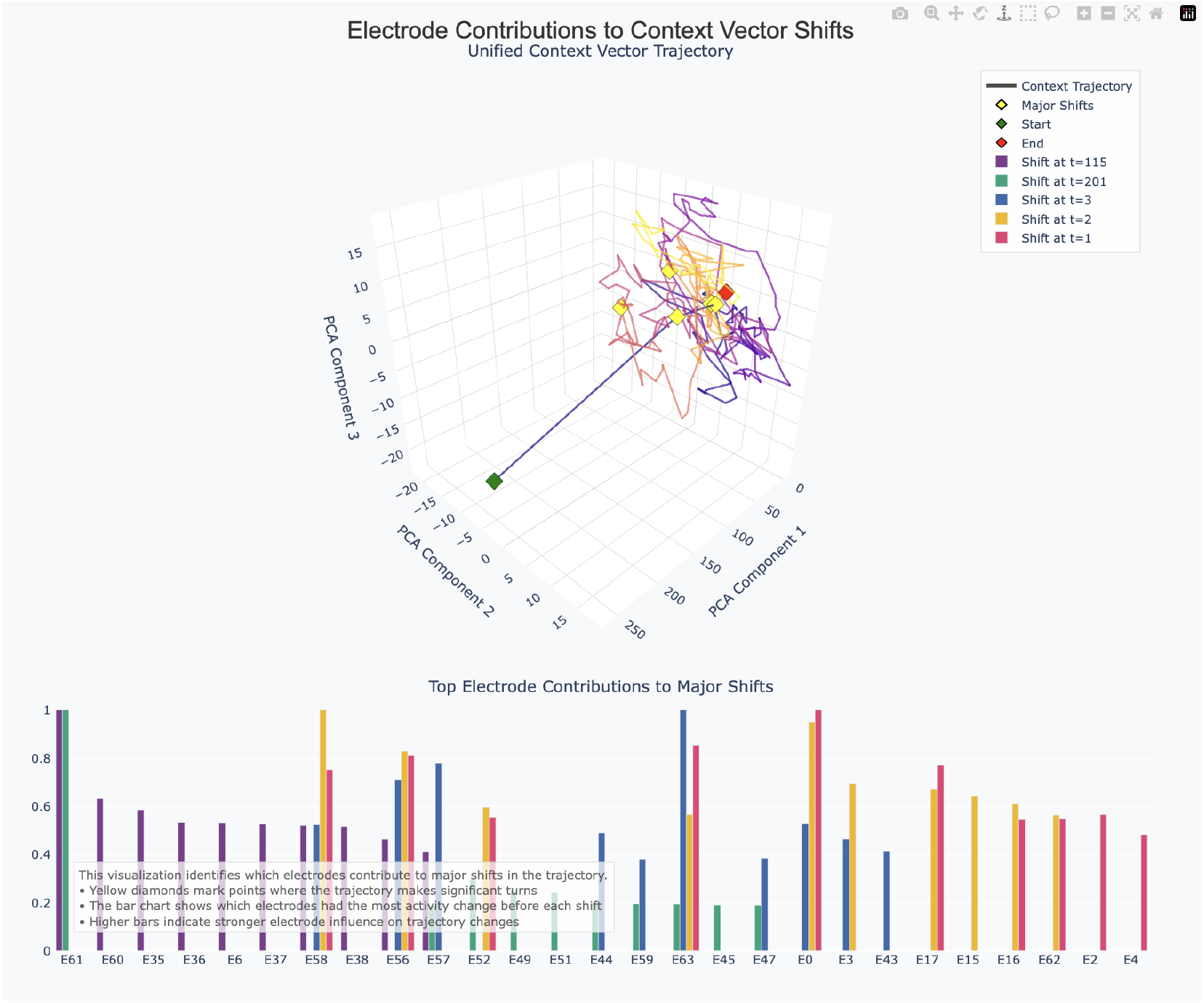
Analysis of electrode contributions to major shifts in the unified context vector. The top panel shows the major shifts highlighted. The bottom panel quantifies which electrodes exhibit the strongest activity changes immediately preceding each shift, revealing their causal influence on the unified representation.

To understand which electrodes drive major shifts in the unified representation, we analyzed activity changes preceding trajectory transitions. This analysis reveals asymmetric influence patterns, with specific electrodes (E61, E58, E56) consistently preceding major transitions in the unified state space. The bar chart in Figure 4 quantifies each electrode’s contribution to specific shifts. Notably, different sets of electrodes appear responsible for different shifts, suggesting functional specialization within the population. Shifts at different timepoints (t=1, t=2, t=3, t=115, t=201) show distinct electrode contribution patterns, revealing how ensemble coordination evolves over time.

Our electrode contribution analysis and knowledge graph reveal that these representational shifts are not driven uniformly across the population, but rather by specific subsets of electrodes that precede major transitions. This asymmetry in causal influence supports hierarchical models of neural information processing, where specialized neural populations drive transitions in global representational states.The temporal progression of representational states reflects a dynamic process of information integration, where the neural ensemble iteratively refines its internal representation based on incoming information and intrinsic dynamics. This process is consistent with predictive coding models of neural computation, where internal states evolve to minimize prediction error through iterative refinement. The identification of electrodes driving major shifts provides insight into the causal structure of this refinement process, suggesting that specific neural populations may serve as control points for guiding representational evolution.These findings advance our understanding of neural ensemble dynamics by revealing the mechanisms through which distributed populations coordinate to form coherent representations. By identifying the specific electrodes and transformation operations involved in this process, we provide a computational framework for investigating how the brain forms stable, coherent representations from distributed patterns of activity.

### 3.4 Limitations and Future Work

WDespite superior performance, our model has several limitations: it currently only analyzes visual cortex LFP recordings during a specific task; gamma oscillations may reflect synaptic and spiking processes not explicitly represented; and while we identify electrode-specific contributions to unified state shifts, the exact computational rules governing these interactions need fuller mathematical characterization, potentially through symbolic regression or rule extraction from trained networks (49). Additionally, our reliance on a single Allen Mouse Neuropixel dataset limits generalizability assessments, necessitating future testing across multiple species, brain regions, sensory modalities, and behavioral contexts to determine whether our computational principles represent fundamental neural mechanisms or merely reflect specific visual cortical dynamics.

## 4 Conclusion

We have presented a Graph Echo State Network with specialized gamma populations that reveals how neural ensembles engineer stable representations through iterative interactions, achieving superior predictive performance while providing mechanistic insights into representational convergence. Our analysis of electrode trajectories, context vectors, and state transitions demonstrates that stable neural representations emerge through specialized, asymmetric population interactions, with electrodes and frequency bands contributing distinctly to reveal a heterogeneous, distributed information integration mechanism.

The specialized population architecture enables the model to disentangle multi-scale oscillatory patterns in neural data while capturing how populations evolve their internal context in response to incoming information. By identifying the specific electrodes and transformation operations driving major shifts in representational state, we offer insight into the causal structure of neural computation, suggesting that specific neural populations may serve as control points for guiding representational evolution. These findings advance our understanding of neural ensemble dynamics and provide a computational framework for investigating coherent representation formation, with potential future extensions across brain regions, behavioral states, and cognitive processes.

## NeurIPS Paper Checklist

1. **1. Claims** Question: Do the main claims made in the abstract and introduction accurately reflect the paper’s contributions and scope? Answer: [Yes] Justification: We have mentioned them in the results section.
2. **Limitations** Question: Does the paper discuss the limitations of the work performed by the authors? Answer: [Yes] Justification: We have mentioned them in the Limitations section.
3. **Theory assumptions and proofs** Question: For each theoretical result, does the paper provide the full set of assumptions and a complete (and correct) proof? Answer: [NA] Justification: Our study is based on existing theoretical work.
4. **Experimental result reproducibility** Question: Does the paper fully disclose all the information needed to reproduce the main experimental results of the paper to the extent that it affects the main claims and/or conclusions of the paper (regardless of whether the code and data are provided or not)? Answer: [Yes] Justification: The code, data available and hyperparameters are included in the supplemental material.
5. **Open access to data and code** Question: Does the paper provide open access to the data and code, with sufficient instructions to faithfully reproduce the main experimental results, as described in supplemental material? Answer: [Yes] Justification: Supplemental material.
6. **Experimental setting/details** Question: Does the paper specify all the training and test details (e.g., data splits, hyperparameters, how they were chosen, type of optimizer, etc.) necessary to understand the results? Answer: [Yes] Justification: All the data splits, hyperparameters, how they were chosen, type of optimizer, etc. are present in the supplemental material.
7. **Experiment statistical significance** Question: Does the paper report error bars suitably and correctly defined or other appropriate information about the statistical significance of the experiments? Answer: [NA] Justification: Not applicable to our current approach.
8. **Experiments compute resources** Question: For each experiment, does the paper provide sufficient information on the computer resources (type of compute workers, memory, time of execution) needed to reproduce the experiments? Answer: [Yes] Justification: Supplemental material.
9. **Code of ethics** Question: Does the research conducted in the paper conform, in every respect, with the NeurIPS Code of Ethics https://neurips.cc/public/EthicsGuidelines? Answer: [Yes] Justification: To our knowledge, yes.
10. **Broader impacts** Question: Does the paper discuss both potential positive societal impacts and negative societal impacts of the work performed? Answer: [NA] Justification: Ongoing study regarding its broader impact.
11. **Safeguards** Question: Does the paper describe safeguards that have been put in place for responsible release of data or models that have a high risk for misuse (e.g., pretrained language models, image generators, or scraped datasets)? Answer: [NA] Justification: Does not come under such a risky purview at this point in time.
12. **Licenses for existing assets** Question: Are the creators or original owners of assets (e.g., code, data, models), used in the paper, properly credited and are the license and terms of use explicitly mentioned and properly respected? Answer: [Yes] Justification: Please check the contributors section.
13. **New assets** Question: Are new assets introduced in the paper well documented and is the documentation provided alongside the assets? Answer: [Yes] Justification: Yes, check Methods and Introduction.
14. **Crowdsourcing and research with human subjects** Question: For crowdsourcing experiments and research with human subjects, does the paper include the full text of instructions given to participants and screenshots, if applicable, as well as details about compensation (if any)? Answer: [NA] Justification: No human subjects were involved in this research.
15. **Institutional review board (IRB) approvals or equivalent for research with human subjects** Question: Does the paper describe potential risks incurred by study participants, whether such risks were disclosed to the subjects, and whether Institutional Review Board (IRB) approvals (or an equivalent approval/review based on the requirements of your country or institution) were obtained? Answer: [NA] Justification: No human subjects were involved in this research.
16. **Declaration of LLM usage** Question: Does the paper describe the usage of LLMs if it is an important, original, or non-standard component of the core methods in this research? Note that if the LLM is used only for writing, editing, or formatting purposes and does not impact the core methodology, scientific rigorousness, or originality of the research, declaration is not required. Answer: [Yes] Justification: As mentioned before, LLMs, specifically Perplexity Claude Sonnet 3.7 and Google Gemini were used in the editing, formatting and implementation of standard methods for this paper.

