## Supplementary material for "How Minds Take Shape: Graph-ESN Reveals How Neural Ensembles Engineer Stable Representations": Comparison.ipynb

```
# %% [code]
import os
import numpy as np
import pandas as pd
import matplotlib.pyplot as plt
import torch
import torch.nn as nn
import torch.nn.functional as F
from torch.utils.data import Dataset, DataLoader
from sklearn.model_selection import train_test_split
from sklearn.preprocessing import RobustScaler
from sklearn.metrics import mean_squared_error, r2_score
from scipy import signal
import warnings
from tqdm import tqdm
import time
import math
from torch.optim.lr_scheduler import ReduceLROnPlateau
import traceback

# Suppress warnings
warnings.filterwarnings('ignore')

# Check for GPU
device = torch.device('cuda' if torch.cuda.is_available() else 'cpu')
print(f"Using device: {device}")

# Set random seed for reproducibility
SEED = 42
np.random.seed(SEED)
torch.manual_seed(SEED)
if torch.cuda.is_available():
    torch.cuda.manual_seed_all(SEED)
    torch.backends.cudnn.deterministic = True
    torch.backends.cudnn.benchmark = False


class Config:
    def __init__(self):
        # Data parameters
        self.NUM_ELECTRODES = 64  # Choose a subset of the 95 available channels
        self.CONTEXT_SIZE = 330   # Size of input window
        self.PREDICT_SIZE = 50    # Size of prediction window
        self.WINDOW_SIZE = self.CONTEXT_SIZE + self.PREDICT_SIZE
        self.SAMPLING_RATE = 1000  # Based on the timestamp differences in the data
        self.MAX_SEQUENCES = 1000    # Number of sequences to extract
        self.BATCH_SIZE = 16
        self.STRIDE = 50

        # Enhanced gamma frequency bands - keep as is
        self.FREQ_BANDS = [
            (30, 50),       # Low Gamma
            (50, 80),       # Mid-Low Gamma
            (80, 120),      # Mid-High Gamma
            (120, 180),     # High Gamma
            (180, 250),     # Ultra High Gamma I
            (250, 300),     # Ultra High Gamma II
        ]

        # Improved learning parameters for gamma prediction
        self.LEARNING_RATE = 2e-4
        self.WEIGHT_DECAY = 2e-5
        self.EPOCHS = 10
        self.PATIENCE = 15
        self.CLIP_GRAD_NORM = 0.8
        self.LR_WARMUP_EPOCHS = 5
        self.GRADIENT_ACCUMULATION_STEPS = 2

        # Enhanced network architecture parameters
        self.HIDDEN_SIZE = 192           # Larger hidden representations
        self.READOUT_HIDDEN = 384        # Larger readout layer
        self.READOUT_LAYERS = 3          # More layers for better decoding
        self.DROPOUT = 0.25              # More regularization
        self.USE_ATTENTION = True
        self.ATTENTION_HEADS = 4         # More attention heads
        self.USE_SKIP_CONNECTIONS = True # Add skip connections
        self.USE_LAYER_NORM = True       # Use layer normalization
        self.USE_BATCH_NORM = True       # Use batch normalization for better training

        # Enhanced loss function parameters
        self.TIME_LOSS_WEIGHT = 0.25     # Less weight for time domain (overall shape)
        self.FREQ_LOSS_WEIGHT = 0.75     # More weight for frequency domain (oscillations)

        # Progressive gamma band weighting (heavily weight higher frequencies)
        self.GAMMA_BAND_WEIGHTS = {
            (30, 50): 1.0,      # Low gamma weight
            (50, 80): 1.5,      # Mid-Low gamma weight
            (80, 120): 2.0,     # Mid-High gamma weight
            (120, 180): 2.5,    # High gamma weight
            (180, 250): 3.0,    # Ultra high gamma I weight
            (250, 300): 3.5     # Ultra high gamma II weight
        }

        # Special weights for phase accuracy
        self.PHASE_LOSS_WEIGHT = 0.4     # Weight for phase prediction accuracy
        self.ENVELOPE_LOSS_WEIGHT = 0.2  # Weight for amplitude envelope
        self.USE_PHASE_CONSISTENCY_LOSS = True  # Use phase consistency loss

        # Directory setup
        self.OUTPUT_DIR = "model_comparison_results"
        self.create_directories()

    def create_directories(self):
        """Create necessary directories for outputs."""
        os.makedirs(self.OUTPUT_DIR, exist_ok=True)
        for subdir in ['models', 'visualizations', 'logs']:
            os.makedirs(os.path.join(self.OUTPUT_DIR, subdir), exist_ok=True)

# Initialize configuration
config = Config()


def find_data_file(path, filename):
    """Find the specified data file in the given path or its subdirectories."""
    data_file = os.path.join(path, filename)
    if os.path.exists(data_file):
        return data_file

    for root, dirs, files in os.walk(path):
        if filename in files:
            return os.path.join(root, filename)

    raise FileNotFoundError(f"Could not find {filename} in {path}")

def load_electrode_positions(path):
    """Load electrode position data from CSV file."""
    print("Loading electrode positions...")
    positions_file = find_data_file(path, 'limbic_insular_probe_channels.csv')

    # Updated column mapping for the new dataset
    positions = pd.read_csv(positions_file)

    # Rename columns to match the expected format
    positions = positions.rename(columns={
        'channel_index': 'channel_num',
        'probe_horizontal_position': 'x_position',
        'probe_vertical_position': 'y_position'
    })

    # IMPORTANT: Filter positions to only include channels 0-94 which are in the data file
    max_channel = 94
    positions = positions[positions['channel_num'] <= max_channel].reset_index(drop=True)

    print(f"Loaded {len(positions)} electrodes (filtered to channels 0-{max_channel})")
    return positions

def select_electrodes(positions, num_electrodes=None):
    """Select electrodes using KMeans clustering for optimal spatial coverage."""
    if num_electrodes is None:
        num_electrodes = config.NUM_ELECTRODES

    print(f"Selecting {num_electrodes} electrodes for analysis...")

    # Normalize positions for clustering
    x_norm = (positions['x_position'] - positions['x_position'].min()) / (positions['x_position'].max() - positions['x_position'].min())
    y_norm = (positions['y_position'] - positions['y_position'].min()) / (positions['y_position'].max() - positions['y_position'].min())
    coords = np.column_stack((x_norm.values, y_norm.values))

    # Use KMeans to select representative electrodes
    from sklearn.cluster import KMeans
    kmeans = KMeans(n_clusters=num_electrodes, random_state=SEED, n_init=10)
    kmeans.fit(coords)

    # Select the electrode closest to each cluster center
    selected_indices = []
    for cluster_idx in range(num_electrodes):
        cluster_points = np.where(kmeans.labels_ == cluster_idx)[0]
        center = kmeans.cluster_centers_[cluster_idx]
        distances = np.sqrt(np.sum((coords[cluster_points] - center)**2, axis=1))
        closest_idx = cluster_points[np.argmin(distances)]
        selected_indices.append(closest_idx)

    # Get channel numbers for selected electrodes
    channel_nums = positions.iloc[selected_indices]['channel_num'].astype(int).values

    print(f"Selected {len(channel_nums)} electrodes with good spatial coverage")
    return channel_nums, selected_indices

def load_lfp_data(path, channel_nums, max_rows=500000):
    """Load LFP data from CSV file."""
    print(f"Loading LFP data for {len(channel_nums)} channels...")
    data_file = find_data_file(path, 'drifting_gratings_full (1).csv')

    # Include presentation_id in the columns to load
    cols_to_use = ['timestamp', 'presentation_id'] + [f'channel_{ch}' for ch in channel_nums]

    print(f"Columns to load (first few): {cols_to_use[:min(5, len(cols_to_use))]}... (and {len(cols_to_use)-5} more)")

    # Load data in chunks to handle large files
    data = pd.read_csv(data_file, usecols=cols_to_use, nrows=max_rows)
    print(f"Loaded {len(data)} rows of LFP data")

    return data

def extract_windows(data, context_size=None, predict_size=None, stride=None, max_windows=None):
    """
    Extract time windows from continuous data for training and prediction,
    ensuring each window comes from a single presentation_id if available.
    """
    if context_size is None:
        context_size = config.CONTEXT_SIZE
    if predict_size is None:
        predict_size = config.PREDICT_SIZE
    if stride is None:
        stride = config.STRIDE
    if max_windows is None:
        max_windows = config.MAX_SEQUENCES

    print(f"Extracting windows with context={context_size}, predict={predict_size}, stride={stride}")

    # Check if presentation_id is available
    use_presentation_id = 'presentation_id' in data.columns
    if use_presentation_id:
        print(f"Using presentation_id to ensure windows come from same presentation")
        print(f"Found {data['presentation_id'].nunique()} unique presentation_ids")
    else:
        print(f"Warning: presentation_id not found in data, windows may cross presentation boundaries")

    # Get channel columns (all columns except timestamp and presentation_id)
    data_cols = [col for col in data.columns if col.startswith('channel_')]

    # Calculate window size
    window_size = context_size + predict_size

    # Extract windows
    windows = []

    if use_presentation_id:
        # Group data by presentation_id
        presentation_groups = data.groupby('presentation_id')

        for presentation_id, group in presentation_groups:
            # Skip presentations that are too short
            if len(group) < window_size:
                continue

            # Get data values for this presentation
            presentation_data = group[data_cols].values

            # Extract windows from this presentation
            for start in range(0, len(presentation_data) - window_size + 1, stride):
                if len(windows) >= max_windows:
                    break

                window = presentation_data[start:start + window_size]
                if np.isnan(window).any():
                    continue

                windows.append(window)

                if len(windows) % 100 == 0:
                    print(f"Extracted {len(windows)} windows...")

            if len(windows) >= max_windows:
                break
    else:
        # Original window extraction logic (not grouping by presentation_id)
        data_values = data[data_cols].values

        for start in range(0, len(data_values) - window_size, stride):
            if len(windows) >= max_windows:
                break

            window = data_values[start:start + window_size]
            if np.isnan(window).any():
                continue

            windows.append(window)

            if len(windows) % 100 == 0:
                print(f"Extracted {len(windows)} windows...")

    if not windows:
        print("No valid windows extracted!")
        return np.array([])

    windows = np.array(windows)

    print(f"Extracted {len(windows)} windows with shape: {windows.shape}")

    return windows

def create_enhanced_filter_bank(fs, freq_bands, order=8):
    """Create enhanced filter bank with higher order filters."""
    filter_bank = []
    nyquist = fs / 2.0

    for band_idx, (low_freq, high_freq) in enumerate(freq_bands):
        # Normalize frequencies
        low = low_freq / nyquist
        high = high_freq / nyquist

        # Create higher order Butterworth filter for sharper cutoffs
        # Using sos (second-order sections) format for better numerical stability
        sos = signal.butter(order, [low, high], btype='bandpass', output='sos')
        filter_bank.append((sos, band_idx, (low_freq, high_freq)))

    return filter_bank

def extract_enhanced_frequency_bands(windows, fs=None, freq_bands=None):
    """Extract frequency bands using enhanced filters with parallel processing."""
    if fs is None:
        fs = config.SAMPLING_RATE
    if freq_bands is None:
        freq_bands = config.FREQ_BANDS

    print("Extracting enhanced frequency bands...")
    n_samples, window_size, n_channels = windows.shape
    n_bands = len(freq_bands)

    # Create filter bank
    filter_bank = create_enhanced_filter_bank(fs, freq_bands, order=8)

    # Initialize output array for filtered data
    filtered_data = np.zeros((n_samples, n_bands, window_size, n_channels))

    # Process each band with enhanced filters
    for (sos, band_idx, (low_freq, high_freq)) in filter_bank:
        print(f"Filtering band {band_idx+1}/{n_bands}: {low_freq}-{high_freq} Hz")

        # Process each sample and channel
        for i in tqdm(range(n_samples), desc=f"Band {band_idx+1}"):
            for j in range(n_channels):
                signal_i = windows[i, :, j]

                # Apply zero-phase filtering with second-order sections (more stable)
                filtered = signal.sosfiltfilt(sos, signal_i)

                # Store filtered signal
                filtered_data[i, band_idx, :, j] = filtered

    print(f"Filtered data shape: {filtered_data.shape}")
    return filtered_data

def enhanced_hilbert_transform(signal_data, pad_factor=2):
    """Apply enhanced Hilbert transform with better padding to reduce edge effects."""
    # Determine padding size (minimum 200 points or half signal length)
    n = len(signal_data)
    pad_size = max(int(n * pad_factor), 200)

    # Apply reflection padding (better than zero padding for reducing edge effects)
    padded_signal = np.pad(signal_data, (pad_size, pad_size), mode='reflect')

    # Apply Hilbert transform to get analytic signal
    analytic_signal = signal.hilbert(padded_signal)

    # Extract phase and amplitude from the central (non-padded) portion
    phase = np.angle(analytic_signal[pad_size:pad_size+n])
    amplitude = np.abs(analytic_signal[pad_size:pad_size+n])

    # Get instantaneous frequency (derivative of phase)
    unwrapped_phase = np.unwrap(phase)
    inst_freq = np.diff(unwrapped_phase) / (2.0 * np.pi) * config.SAMPLING_RATE
    # Pad the instantaneous frequency to match original length
    inst_freq = np.append(inst_freq, inst_freq[-1])

    return phase, amplitude, inst_freq

def extract_phase_amplitude(filtered_data):
    """Extract phase, amplitude and instantaneous frequency using enhanced Hilbert transform."""
    print("Extracting phase, amplitude and instantaneous frequency information...")
    n_samples, n_bands, n_timepoints, n_channels = filtered_data.shape

    # Initialize arrays for phase, amplitude and frequency
    phase = np.zeros_like(filtered_data)
    amplitude = np.zeros_like(filtered_data)
    inst_freq = np.zeros_like(filtered_data)

    for i in tqdm(range(n_samples), desc="Processing samples"):
        for b in range(n_bands):
            for c in range(n_channels):
                try:
                    # Apply enhanced Hilbert transform
                    p, a, f = enhanced_hilbert_transform(filtered_data[i, b, :, c])

                    # Store results
                    phase[i, b, :, c] = p
                    amplitude[i, b, :, c] = a
                    inst_freq[i, b, :, c] = f
                except Exception as e:
                    print(f"Error in Hilbert transform: {e}")
                    # Fill with zeros on error
                    phase[i, b, :, c] = 0
                    amplitude[i, b, :, c] = 0
                    inst_freq[i, b, :, c] = 0

    # Normalize amplitude (per band) for better stability using 99th percentile
    for b in range(n_bands):
        max_amp = np.percentile(amplitude[:, b, :, :], 99)
        if max_amp > 0:
            amplitude[:, b, :, :] /= max_amp

    return phase, amplitude, inst_freq

def compute_spectral_features(data, fs=None, nperseg=64, noverlap=32):
    """Compute time-frequency features using spectrogram instead of wavelets."""
    if fs is None:
        fs = config.SAMPLING_RATE

    n_samples, n_timepoints, n_channels = data.shape

    # Compute spectrogram parameters
    f, _, _ = signal.spectrogram(np.zeros(nperseg), fs=fs, nperseg=nperseg, noverlap=noverlap)
    n_freq_bins = len(f)
    n_time_bins = 1 + (n_timepoints - nperseg) // (nperseg - noverlap)

    # Initialize output arrays
    spec_power = np.zeros((n_samples, n_freq_bins, n_time_bins, n_channels))

    # Compute spectrogram for each sample and channel
    print("Computing spectral features...")
    for i in tqdm(range(n_samples), desc="Spectral analysis"):
        for c in range(n_channels):
            signal_i = data[i, :, c]

            # Compute spectrogram
            f, t, Sxx = signal.spectrogram(
                signal_i,
                fs=fs,
                nperseg=nperseg,
                noverlap=noverlap,
                scaling='density'
            )

            # Store power
            spec_power[i, :, :Sxx.shape[1], c] = Sxx

    # Focus on gamma frequencies (>30 Hz)
    gamma_mask = f >= 30
    spec_power_gamma = spec_power[:, gamma_mask, :, :]
    freq_bins_gamma = f[gamma_mask]

    # Normalize spectrogram power
    for f_idx in range(len(freq_bins_gamma)): # Iterate using index
        max_power = np.percentile(spec_power_gamma[:, f_idx, :, :], 99)
        if max_power > 0:
            spec_power_gamma[:, f_idx, :, :] /= max_power

    return spec_power_gamma, freq_bins_gamma

def compute_power_features(amplitude_data, window_size=25):
    """Compute enhanced power-related features from amplitude data."""
    # Get dimensions
    n_samples, n_bands, n_timepoints, n_channels = amplitude_data.shape

    # Compute power (square of amplitude)
    power_data = amplitude_data ** 2

    # Use a more appropriate window size based on signal length
    window_size = min(window_size, n_timepoints // 5)
    stride = max(1, window_size // 2)  # 50% overlap

    # Number of windows
    n_windows = (n_timepoints - window_size) // stride + 1

    # Initialize power features array
    power_features = np.zeros((n_samples, n_bands, n_windows, n_channels))

    # Compute power in each window
    for i in range(n_samples):
        for b in range(n_bands):
            for c in range(n_channels):
                for w in range(n_windows):
                    start_idx = w * stride
                    end_idx = start_idx + window_size
                    power_features[i, b, w, c] = np.mean(power_data[i, b, start_idx:end_idx, c])

    return power_features

def preprocess_data(windows, context_size=None, predict_size=None):
    """Enhanced preprocessing with more sophisticated spectral features for gamma focus."""
    if context_size is None:
        context_size = config.CONTEXT_SIZE
    if predict_size is None:
        predict_size = config.PREDICT_SIZE

    print("Preprocessing data with enhanced gamma focus...")
    num_windows, window_size, num_electrodes = windows.shape
    assert window_size == context_size + predict_size

    # Split windows into context and target parts
    context_windows = windows[:, :context_size, :]
    target_windows = windows[:, context_size:, :]

   # Adjust the scaler to handle the very small amplitude ranges
    scaler = RobustScaler(quantile_range=(1, 99))

    # For very small amplitudes, multiply by 1e6 to make them more numerically stable
    context_flat = context_windows.reshape(-1, num_electrodes) * 1e6
    target_flat = target_windows.reshape(-1, num_electrodes) * 1e6

    # Scale the data
    scaler.fit(context_flat)
    context_scaled = scaler.transform(context_flat).reshape(context_windows.shape)
    target_scaled = scaler.transform(target_flat).reshape(target_windows.shape)

    # Similarly for full_flat
    full_flat = windows.reshape(-1, num_electrodes) * 1e6
    full_scaled = scaler.transform(full_flat).reshape(windows.shape)

    # Extract frequency band information with enhanced filters
    filtered_windows = extract_enhanced_frequency_bands(full_scaled, config.SAMPLING_RATE, config.FREQ_BANDS)

    # Extract phase, amplitude, and instantaneous frequency information
    phase_data, amplitude_data, inst_freq_data = extract_phase_amplitude(filtered_windows)

    # Compute spectral features
    spectral_power, freq_bins = compute_spectral_features(full_scaled, config.SAMPLING_RATE)

    # Compute power features for enhanced spectral representation
    power_features = compute_power_features(amplitude_data)

    # Split features into context and target parts
    phase_context = phase_data[:, :, :context_size, :]
    amplitude_context = amplitude_data[:, :, :context_size, :]
    freq_context = inst_freq_data[:, :, :context_size, :]
    filtered_context = filtered_windows[:, :, :context_size, :]

    # Calculate number of time bins in spectral data
    nperseg = 64
    noverlap = 32
    n_time_bins_context = 1 + (context_size - nperseg) // (nperseg - noverlap)
    spectral_context = spectral_power[:, :, :n_time_bins_context, :]

    # Calculate number of windows for downsampled features (power_features)
    power_feature_window_size = 25 # from compute_power_features default
    power_feature_stride = max(1, power_feature_window_size // 2)
    n_windows_context_power = (context_size - power_feature_window_size) // power_feature_stride + 1

    # Create dataset with all features
    X_data = {
        'raw': context_scaled,
        'phase': phase_context,
        'amplitude': amplitude_context,
        'inst_freq': freq_context,
        'filtered': filtered_context,
        'power': power_features[:, :, :n_windows_context_power, :],
        'spectral': spectral_context
    }

    y_data = {
        'target': target_scaled,
        'phase_target': phase_data[:, :, context_size:, :],
        'amplitude_target': amplitude_data[:, :, context_size:, :],
        'freq_target': inst_freq_data[:, :, context_size:, :]
    }

    # Split into training, validation, and test sets (70/15/15 split)
    indices = np.arange(len(context_scaled))
    train_idx, temp_idx = train_test_split(indices, test_size=0.3, random_state=SEED)
    val_idx, test_idx = train_test_split(temp_idx, test_size=0.5, random_state=SEED)

    # Create PyTorch datasets
    train_dataset = EnhancedLFPDataset(
        {k: v[train_idx] for k, v in X_data.items()},
        {k: v[train_idx] for k, v in y_data.items()}
    )

    val_dataset = EnhancedLFPDataset(
        {k: v[val_idx] for k, v in X_data.items()},
        {k: v[val_idx] for k, v in y_data.items()}
    )

    test_dataset = EnhancedLFPDataset(
        {k: v[test_idx] for k, v in X_data.items()},
        {k: v[test_idx] for k, v in y_data.items()}
    )

    # Create data loaders
    train_loader = DataLoader(train_dataset, batch_size=config.BATCH_SIZE, shuffle=True)
    val_loader = DataLoader(val_dataset, batch_size=config.BATCH_SIZE)
    test_loader = DataLoader(test_dataset, batch_size=config.BATCH_SIZE)

    print(f"Created data loaders with {len(train_loader)} training batches, "
          f"{len(val_loader)} validation batches, and {len(test_loader)} test batches")

    return train_loader, val_loader, test_loader, scaler

class EnhancedLFPDataset(Dataset):
    """Enhanced dataset for LFP data with gamma-specific spectral decomposition."""

    def __init__(self, X, y):
        """Initialize dataset with extended input features and target."""
        self.X_raw = torch.FloatTensor(X['raw'])
        self.X_phase = torch.FloatTensor(X['phase'])
        self.X_amplitude = torch.FloatTensor(X['amplitude'])
        self.X_inst_freq = torch.FloatTensor(X['inst_freq'])
        self.X_filtered = torch.FloatTensor(X['filtered'])
        self.X_power = torch.FloatTensor(X['power'])
        self.X_spectral = torch.FloatTensor(X['spectral'])

        self.y_target = torch.FloatTensor(y['target'])
        self.y_phase = torch.FloatTensor(y['phase_target'])
        self.y_amplitude = torch.FloatTensor(y['amplitude_target'])
        self.y_freq = torch.FloatTensor(y['freq_target'])

    def __len__(self):
        return len(self.y_target)

    def __getitem__(self, idx):
        return {
            'raw': self.X_raw[idx],
            'phase': self.X_phase[idx],
            'amplitude': self.X_amplitude[idx],
            'inst_freq': self.X_inst_freq[idx],
            'filtered': self.X_filtered[idx],
            'power': self.X_power[idx],
            'spectral': self.X_spectral[idx]
        }, {
            'target': self.y_target[idx],
            'phase': self.y_phase[idx],
            'amplitude': self.y_amplitude[idx],
            'freq': self.y_freq[idx]
        }


def compute_fft_torch(signal, fs=1000):
    """Compute FFT using PyTorch for better GPU compatibility."""
    n = signal.shape[-1] # Operate on last dimension (time)
    rfft_output = torch.fft.rfft(signal, n=n, dim=-1) # Use dim=-1 for [B,C,T] or [B,T]
    magnitudes = torch.abs(rfft_output) / n
    phases = torch.angle(rfft_output)
    frequencies = torch.fft.rfftfreq(n, d=1.0/fs, device=signal.device) # Correct d for sampling interval
    return frequencies, magnitudes, phases

class PhaseConsistencyLoss(nn.Module):
    """Loss function that measures consistency of phase relationships."""
    def __init__(self):
        super().__init__()

    def forward(self, pred_phases, true_phases):
        """
        Compute phase consistency loss.
        Args:
            pred_phases: Predicted phases [..., frequencies, (time_stft_bins)] or [..., frequencies] for FFT
            true_phases: True phases   [..., frequencies, (time_stft_bins)] or [..., frequencies] for FFT
        """
        phase_diff = pred_phases - true_phases
        # Wrap phase difference to [-pi, pi] for stability before cos/sin
        phase_diff_wrapped = torch.remainder(phase_diff + np.pi, 2 * np.pi) - np.pi

        real_part = torch.cos(phase_diff_wrapped)
        imag_part = torch.sin(phase_diff_wrapped)
        complex_diff = torch.complex(real_part, imag_part)

        # Mean over frequencies (or time_stft_bins then frequencies)
        # If FFT output [..., freqs], mean over freqs
        mean_vector = torch.mean(complex_diff, dim=-1) # Mean over last dim (freqs)
        if mean_vector.ndim > complex_diff.ndim - 1 and mean_vector.shape[-1] != 1: # if STFT output
             mean_vector = torch.mean(mean_vector, dim=-1) # then mean over time bins

        consistency = torch.abs(mean_vector)
        loss = 1.0 - consistency
        return torch.mean(loss) # Mean over batch and other dims

class EnhancedGammaLoss(nn.Module):
    """Enhanced loss function optimized for gamma oscillation prediction."""
    def __init__(self, config):
        super().__init__()
        self.config = config
        self.time_weight = config.TIME_LOSS_WEIGHT
        self.freq_weight = config.FREQ_LOSS_WEIGHT
        self.phase_weight = config.PHASE_LOSS_WEIGHT
        self.envelope_weight = config.ENVELOPE_LOSS_WEIGHT
        self.gamma_band_weights = config.GAMMA_BAND_WEIGHTS

        self.mse_loss = nn.MSELoss()
        self.phase_consistency_loss = PhaseConsistencyLoss()
        self.sampling_rate = float(config.SAMPLING_RATE) # Ensure float

        self.freq_band_map = {i: band for i, band in enumerate(config.FREQ_BANDS)}

    def gamma_frequency_loss(self, pred, target):
        batch_size, seq_len, num_channels = pred.shape

        # pred and target are [batch, time, channels]
        # For FFT, want [batch*channels, time]
        pred_reshaped = pred.permute(0, 2, 1).reshape(-1, seq_len)
        target_reshaped = target.permute(0, 2, 1).reshape(-1, seq_len)

        pred_freqs, pred_mags, pred_phases = compute_fft_torch(pred_reshaped, self.sampling_rate)
        target_freqs, target_mags, target_phases = compute_fft_torch(target_reshaped, self.sampling_rate)
        # freqs shape [n_fft_bins], mags/phases shape [batch*channels, n_fft_bins]

        # Initialize band weights tensor on the correct device
        weights = torch.ones_like(pred_freqs, device=pred.device) * 0.1
        for band_idx, (low, high) in self.freq_band_map.items():
            band_mask = (pred_freqs >= low) & (pred_freqs <= high)
            band_weight_val = self.gamma_band_weights.get((low, high), 1.0)
            weights = torch.where(band_mask, torch.tensor(band_weight_val, device=weights.device), weights)

        # Weighted MSE for magnitude (spectrum)
        # pred_mags, target_mags are [B*C, n_fft_bins]. weights is [n_fft_bins]
        mag_diff_sq = (pred_mags - target_mags)**2
        # Multiply by weights (broadcasts over B*C dim): [B*C, n_fft_bins] * [1, n_fft_bins]
        weighted_mag_loss = torch.mean(mag_diff_sq * weights.unsqueeze(0))
        freq_loss = weighted_mag_loss

        # Phase consistency loss for gamma frequencies
        phase_loss_val = torch.tensor(0.0, device=pred.device)
        if self.config.USE_PHASE_CONSISTENCY_LOSS:
            gamma_mask = pred_freqs >= 30
            if gamma_mask.any(): # Check if any gamma frequencies exist
                # pred_phases/target_phases are [B*C, n_fft_bins]
                # Select gamma frequencies: [B*C, n_gamma_bins]
                pred_gamma_phases = pred_phases[:, gamma_mask]
                target_gamma_phases = target_phases[:, gamma_mask]
                if pred_gamma_phases.shape[-1] > 0: # Ensure there are bins
                     phase_loss_val = self.phase_consistency_loss(pred_gamma_phases, target_gamma_phases)

        # Envelope loss using magnitude of gamma frequencies
        envelope_loss_val = torch.tensor(0.0, device=pred.device)
        gamma_mask_env = pred_freqs >= 30 # Re-evaluate or use gamma_mask
        if gamma_mask_env.any():
            pred_gamma_mags = pred_mags[:, gamma_mask_env]
            target_gamma_mags = target_mags[:, gamma_mask_env]
            if pred_gamma_mags.shape[-1] > 0:
                 envelope_loss_val = F.mse_loss(pred_gamma_mags, target_gamma_mags)

        return freq_loss, phase_loss_val, envelope_loss_val


    def forward(self, pred, target):
        time_loss = self.mse_loss(pred, target)
        freq_loss, phase_loss, envelope_loss = self.gamma_frequency_loss(pred, target)

        total_loss = (
            self.time_weight * time_loss +
            self.freq_weight * freq_loss +
            self.phase_weight * phase_loss +
            self.envelope_weight * envelope_loss
        )
        return total_loss, time_loss, freq_loss, phase_loss, envelope_loss

#----------------------------------------------------------------------
# Training and Evaluation Functions
#----------------------------------------------------------------------

def train_epoch(model, train_loader, optimizer, criterion, scheduler, device, config, epoch, clip_grad=None):
    """Enhanced training for one epoch with gradient accumulation and curriculum learning."""
    model.train()
    total_loss_val, time_losses_val, freq_losses_val, phase_losses_val, envelope_losses_val = 0.0, 0.0, 0.0, 0.0, 0.0

    if epoch < config.LR_WARMUP_EPOCHS and config.LR_WARMUP_EPOCHS > 0: # Avoid division by zero
        warmup_factor = (epoch + 1) / config.LR_WARMUP_EPOCHS
        for param_group in optimizer.param_groups:
            param_group['lr'] = config.LEARNING_RATE * warmup_factor

    progress_bar = tqdm(train_loader, desc=f"Training Epoch {epoch+1}")
    optimizer.zero_grad() # Initialize gradients

    for batch_idx, batch_data in enumerate(progress_bar):
        try:
            inputs, targets_dict = batch_data
            inputs = {k: v.to(device) for k, v in inputs.items()}
            target_signal = targets_dict['target'].to(device)

            outputs = model(inputs)
            loss, time_loss, freq_loss, phase_loss, envelope_loss = criterion(outputs, target_signal)

            if torch.isnan(loss):
                print(f"Warning: NaN loss encountered at epoch {epoch}, batch {batch_idx}. Skipping batch.")
                # If an optimizer step was planned, ensure gradients are zeroed
                if (batch_idx + 1) % config.GRADIENT_ACCUMULATION_STEPS == 0 or \
                   (batch_idx + 1) == len(train_loader):
                    optimizer.zero_grad() # Ensure grads are zeroed before next accumulation cycle
                continue

            scaled_loss = loss / config.GRADIENT_ACCUMULATION_STEPS
            scaled_loss.backward()

            if (batch_idx + 1) % config.GRADIENT_ACCUMULATION_STEPS == 0 or \
               (batch_idx + 1) == len(train_loader):
                if clip_grad is not None:
                    torch.nn.utils.clip_grad_norm_(model.parameters(), clip_grad)
                optimizer.step()
                optimizer.zero_grad() # Zero gradients after step

            total_loss_val += loss.item()
            time_losses_val += time_loss.item() if torch.is_tensor(time_loss) else time_loss
            freq_losses_val += freq_loss.item() if torch.is_tensor(freq_loss) else freq_loss
            phase_losses_val += phase_loss.item() if torch.is_tensor(phase_loss) else phase_loss
            envelope_losses_val += envelope_loss.item() if torch.is_tensor(envelope_loss) else envelope_loss

            progress_bar.set_postfix({
                'loss': loss.item(),
                'time_l': time_loss.item() if torch.is_tensor(time_loss) else time_loss,
                'freq_l': freq_loss.item() if torch.is_tensor(freq_loss) else freq_loss
            })

        except Exception as e:
            print(f"Error in training batch (epoch {epoch}, batch {batch_idx}): {type(e).__name__} - {e}")
            traceback.print_exc()
            # Ensure gradients are zeroed if an optimizer step was planned
            if (batch_idx + 1) % config.GRADIENT_ACCUMULATION_STEPS == 0 or \
                (batch_idx + 1) == len(train_loader):
                optimizer.zero_grad()
            continue

    num_batches = len(train_loader)
    if num_batches == 0: return {'loss': 0, 'time_loss': 0, 'freq_loss': 0, 'phase_loss': 0, 'envelope_loss': 0}

    return {
        'loss': total_loss_val / num_batches,
        'time_loss': time_losses_val / num_batches,
        'freq_loss': freq_losses_val / num_batches,
        'phase_loss': phase_losses_val / num_batches,
        'envelope_loss': envelope_losses_val / num_batches
    }

def evaluate(model, val_loader, criterion, device, compute_metrics=True):
    """Enhanced evaluation with detailed gamma-specific metrics."""
    model.eval()
    total_loss = 0
    time_losses = 0
    freq_losses = 0
    phase_losses = 0
    envelope_losses = 0

    # Metrics storage with separate gamma band metrics
    gamma_metrics = {band: {'mse': [], 'r2': [], 'corr': []} for band in config.FREQ_BANDS}

    all_metrics = {
        'mse': [],
        'r2': [],
        'corr': [],
        'gamma_metrics': gamma_metrics,
        'spectrum_corr': []  # Correlation in frequency domain
    }

    # Store sample states for visualization
    sample_states = None
    sample_inputs = None
    sample_targets = None
    sample_outputs = None

    with torch.no_grad():
        for batch_idx, batch_data in enumerate(tqdm(val_loader, desc="Evaluating")):
            try:
                # Unpack data
                inputs, targets = batch_data

                # Move to device
                inputs = {k: v.to(device) for k, v in inputs.items()}
                target_signal = targets['target'].to(device)

                # Forward pass with state tracking
                if hasattr(model, 'return_states') and callable(getattr(model, 'return_states')):
                    outputs, states = model(inputs, return_states=True)
                else:
                    outputs = model(inputs)
                    states = None

                # Store one set of samples for visualization
                if sample_states is None:
                    sample_states = states
                    sample_inputs = inputs
                    sample_targets = target_signal.cpu().numpy()
                    sample_outputs = outputs.cpu().numpy()

                # Compute loss
                loss, time_loss, freq_loss, phase_loss, envelope_loss = criterion(outputs, target_signal)

                # Track losses
                total_loss += loss.item()
                time_losses += time_loss.item()
                freq_losses += freq_loss.item()
                phase_losses += phase_loss.item()
                envelope_losses += envelope_loss.item()

                # Compute metrics if requested
                if compute_metrics:
                    # Move to CPU for numpy processing
                    outputs_np = outputs.cpu().numpy()
                    targets_np = target_signal.cpu().numpy()

                    # Get frequency band data for filtering
                    phase_target = targets['phase'].cpu().numpy()
                    amp_target = targets['amplitude'].cpu().numpy()

                    for i in range(outputs_np.shape[0]):
                        # Full signal metrics for limited electrodes
                        for e in range(min(5, outputs_np.shape[2])):
                            # Extract electrode signals
                            pred_signal = outputs_np[i, :, e]
                            true_signal = targets_np[i, :, e]

                            # Calculate full signal metrics
                            try:
                                mse = np.mean((pred_signal - true_signal)**2)
                                all_metrics['mse'].append(mse)

                                r2 = r2_score(true_signal, pred_signal)
                                if not np.isnan(r2):
                                    all_metrics['r2'].append(r2)

                                corr = np.corrcoef(pred_signal, true_signal)[0, 1]
                                if not np.isnan(corr):
                                    all_metrics['corr'].append(corr)

                                # Compute spectrum correlation
                                fs = config.SAMPLING_RATE
                                pred_fft = np.abs(np.fft.rfft(pred_signal))
                                true_fft = np.abs(np.fft.rfft(true_signal))
                                # Focus on gamma range frequencies (30-300 Hz)
                                freq_bins = np.fft.rfftfreq(len(true_signal), 1/fs)
                                gamma_mask = (freq_bins >= 30) & (freq_bins <= 300)
                                pred_fft_gamma = pred_fft[gamma_mask]
                                true_fft_gamma = true_fft[gamma_mask]
                                spec_corr = np.corrcoef(pred_fft_gamma, true_fft_gamma)[0, 1]
                                if not np.isnan(spec_corr):
                                    all_metrics['spectrum_corr'].append(spec_corr)

                            except Exception as e:
                                print(f"Error computing metrics: {e}")

                            # Compute band-specific metrics
                            for band_idx, (low_freq, high_freq) in enumerate(config.FREQ_BANDS):
                                try:
                                    # Extract band-specific components using target amplitude/phase
                                    # Use the ground truth filtered signal as reference
                                    if 'filtered' in inputs:
                                        band_true = inputs['filtered'][i, band_idx, :, e].cpu().numpy()
                                    else:
                                        band_true = true_signal  # Fallback if filtered not available

                                    # Skip filtering for short signals
                                    if len(pred_signal) < 20:  # Skip if signal is too short
                                        continue

                                    # Apply same band-pass filter to predicted signal
                                    nyquist = config.SAMPLING_RATE / 2
                                    low = low_freq / nyquist
                                    high = high_freq / nyquist

                                    # Use a much smaller filter order for short signals
                                    filter_order = min(2, len(pred_signal) // 10)  # Use only order 1 or 2 for short signals
                                    if filter_order < 1:
                                        continue  # Skip if we can't even use order 1

                                    # Design filter with reduced order
                                    sos = signal.butter(filter_order, [low, high], btype='bandpass', output='sos')

                                    # Forward filtering only (no zero-phase) - much less demanding on signal length
                                    band_pred = signal.sosfilt(sos, pred_signal)
                                    band_true_filtered = signal.sosfilt(sos, true_signal)

                                    # Calculate band-specific metrics
                                    band_mse = np.mean((band_pred - band_true_filtered)**2)
                                    gamma_metrics[(low_freq, high_freq)]['mse'].append(band_mse)

                                    band_r2 = r2_score(band_true_filtered, band_pred)
                                    if not np.isnan(band_r2):
                                        gamma_metrics[(low_freq, high_freq)]['r2'].append(band_r2)

                                    band_corr = np.corrcoef(band_pred, band_true_filtered)[0, 1]
                                    if not np.isnan(band_corr):
                                        gamma_metrics[(low_freq, high_freq)]['corr'].append(band_corr)

                                except Exception as exc:
                                    # Use exc instead of e to avoid variable name conflicts
                                    print(f"Error in band-specific metrics: {exc}")

            except Exception as e:
                print(f"Error in evaluation batch: {e}")
                continue

    # Calculate average losses
    num_batches = max(1, len(val_loader))
    avg_loss = total_loss / num_batches
    avg_time_loss = time_losses / num_batches
    avg_freq_loss = freq_losses / num_batches
    avg_phase_loss = phase_losses / num_batches
    avg_envelope_loss = envelope_losses / num_batches

    # Calculate average metrics
    avg_metrics = {
        'loss': avg_loss,
        'time_loss': avg_time_loss,
        'freq_loss': avg_freq_loss,
        'phase_loss': avg_phase_loss,
        'envelope_loss': avg_envelope_loss
    }

    for key in ['mse', 'r2', 'corr', 'spectrum_corr']:
        values = all_metrics.get(key, [])
        if values:
            avg_metrics[key] = np.mean(values)
        else:
            avg_metrics[key] = float('nan')

    # Calculate average band-specific metrics
    band_avg_metrics = {}
    for band, metrics in gamma_metrics.items():
        band_avg_metrics[band] = {}
        for metric_name, values in metrics.items():
            if values:
                band_avg_metrics[band][metric_name] = np.mean(values)
            else:
                band_avg_metrics[band][metric_name] = float('nan')

    avg_metrics['gamma_metrics'] = band_avg_metrics

    # Return also sample data for visualization
    return avg_metrics, sample_states, sample_inputs, sample_targets, sample_outputs


class EnhancedRNNModel(nn.Module):
    """RNN model optimized for gamma oscillation prediction."""
    def __init__(self, config):
        super().__init__()
        self.config = config

        # Input dimensions
        self.num_electrodes = config.NUM_ELECTRODES
        self.num_freq_bands = len(config.FREQ_BANDS)
        self.predict_size = config.PREDICT_SIZE

        # Enhanced feature projections
        self.raw_projection = nn.Linear(self.num_electrodes, config.HIDDEN_SIZE // 4)
        self.phase_projection = nn.Linear(self.num_freq_bands * self.num_electrodes, config.HIDDEN_SIZE // 4)
        self.amplitude_projection = nn.Linear(self.num_freq_bands * self.num_electrodes, config.HIDDEN_SIZE // 4)
        self.freq_projection = nn.Linear(self.num_freq_bands * self.num_electrodes, config.HIDDEN_SIZE // 4)

        # Feature fusion
        self.feature_fusion = nn.Sequential(
            nn.Linear(config.HIDDEN_SIZE, config.HIDDEN_SIZE),
            nn.LayerNorm(config.HIDDEN_SIZE),
            nn.GELU(),
            nn.Dropout(config.DROPOUT),
            nn.Linear(config.HIDDEN_SIZE, config.HIDDEN_SIZE),
            nn.LayerNorm(config.HIDDEN_SIZE),
            nn.GELU()
        )

        # RNN layers
        self.rnn = nn.RNN(
            input_size=config.HIDDEN_SIZE,
            hidden_size=config.READOUT_HIDDEN,
            num_layers=3,
            batch_first=True,
            dropout=config.DROPOUT,
            bidirectional=True
        )

        # Bidirectional projection
        self.bidirectional_proj = nn.Linear(config.READOUT_HIDDEN * 2, config.READOUT_HIDDEN)

        # Common readout layers
        readout_layers = []
        current_dim = config.READOUT_HIDDEN
        for i in range(config.READOUT_LAYERS - 1):
            next_dim = current_dim // 2
            readout_layers.extend([
                nn.Linear(current_dim, next_dim),
                nn.LayerNorm(next_dim),
                nn.GELU(),
                nn.Dropout(0.1)
            ])
            current_dim = next_dim

        self.readout_common = nn.Sequential(*readout_layers)
        self.final_readout_dim = current_dim

        # Main decoder
        self.main_decoder = nn.Linear(self.final_readout_dim, self.num_electrodes)

        # Gamma band-specific decoders
        self.gamma_decoders = nn.ModuleList([
            nn.Linear(self.final_readout_dim, self.num_electrodes)
            for _ in config.FREQ_BANDS
        ])

        # Time-specific prediction heads
        self.timestep_heads = nn.ModuleList([
            nn.Linear(self.num_electrodes * (len(config.FREQ_BANDS) + 1), self.num_electrodes)
            for _ in range(config.PREDICT_SIZE)
        ])

    def forward(self, x_dict, return_states=False):
        x_raw = x_dict['raw']
        x_phase = x_dict['phase']
        x_amplitude = x_dict['amplitude']
        x_inst_freq = x_dict['inst_freq']

        batch_size = x_raw.shape[0]
        context_size = x_raw.shape[1]
        device = x_raw.device

        # Process each timestep
        sequence = []
        for t in range(context_size):
            # Raw features
            raw_t = x_raw[:, t, :]
            raw_features = self.raw_projection(raw_t)

            # Phase features
            phase_t = x_phase[:, :, t, :].reshape(batch_size, -1)
            phase_features = self.phase_projection(phase_t)

            # Amplitude features
            amplitude_t = x_amplitude[:, :, t, :].reshape(batch_size, -1)
            amplitude_features = self.amplitude_projection(amplitude_t)

            # Frequency features
            freq_t = x_inst_freq[:, :, t, :].reshape(batch_size, -1)
            freq_features = self.freq_projection(freq_t)

            # Combine features
            combined_features = torch.cat([
                raw_features, phase_features, amplitude_features, freq_features
            ], dim=1)

            # Apply feature fusion
            fused_features = self.feature_fusion(combined_features)
            sequence.append(fused_features)

        # Stack into sequence
        sequence_tensor = torch.stack(sequence, dim=1)  # [batch, time, features]

        # Pass through RNN
        rnn_out, h_n = self.rnn(sequence_tensor)

        # Use final hidden state from both directions
        h_forward = h_n[-2, :, :]
        h_backward = h_n[-1, :, :]
        bidirectional_hidden = torch.cat([h_forward, h_backward], dim=1)
        final_hidden = self.bidirectional_proj(bidirectional_hidden)

        # Apply common readout
        common_features = self.readout_common(final_hidden)

        # Decoder outputs
        main_output = self.main_decoder(common_features)
        gamma_outputs = [decoder(common_features) for decoder in self.gamma_decoders]

        # Combine all outputs
        all_outputs_stacked = torch.stack([main_output] + gamma_outputs, dim=1)
        timestep_input_reshaped = all_outputs_stacked.reshape(batch_size, -1)

        # Predict each timestep
        predictions = torch.zeros(batch_size, self.predict_size, self.num_electrodes, device=device)
        for t_pred in range(self.predict_size):
            predictions[:, t_pred, :] = self.timestep_heads[t_pred](timestep_input_reshaped)

        # Optionally return states for visualization
        states = sequence_tensor if return_states else None
        return (predictions, states) if return_states else predictions

class EnhancedLSTMModel(nn.Module):
    """LSTM model optimized for gamma oscillation prediction."""
    def __init__(self, config):
        super().__init__()
        self.config = config

        # Input dimensions
        self.num_electrodes = config.NUM_ELECTRODES
        self.num_freq_bands = len(config.FREQ_BANDS)
        self.predict_size = config.PREDICT_SIZE

        # Enhanced feature projections
        self.raw_projection = nn.Linear(self.num_electrodes, config.HIDDEN_SIZE // 4)
        self.phase_projection = nn.Linear(self.num_freq_bands * self.num_electrodes, config.HIDDEN_SIZE // 4)
        self.amplitude_projection = nn.Linear(self.num_freq_bands * self.num_electrodes, config.HIDDEN_SIZE // 4)
        self.freq_projection = nn.Linear(self.num_freq_bands * self.num_electrodes, config.HIDDEN_SIZE // 4)

        # Feature fusion
        self.feature_fusion = nn.Sequential(
            nn.Linear(config.HIDDEN_SIZE, config.HIDDEN_SIZE),
            nn.LayerNorm(config.HIDDEN_SIZE),
            nn.GELU(),
            nn.Dropout(config.DROPOUT),
            nn.Linear(config.HIDDEN_SIZE, config.HIDDEN_SIZE),
            nn.LayerNorm(config.HIDDEN_SIZE),
            nn.GELU()
        )

        # LSTM layers
        self.lstm = nn.LSTM(
            input_size=config.HIDDEN_SIZE,
            hidden_size=config.READOUT_HIDDEN,
            num_layers=3,
            batch_first=True,
            dropout=config.DROPOUT,
            bidirectional=True
        )

        # Bidirectional projection
        self.bidirectional_proj = nn.Linear(config.READOUT_HIDDEN * 2, config.READOUT_HIDDEN)

        # Common readout layers
        readout_layers = []
        current_dim = config.READOUT_HIDDEN
        for i in range(config.READOUT_LAYERS - 1):
            next_dim = current_dim // 2
            readout_layers.extend([
                nn.Linear(current_dim, next_dim),
                nn.LayerNorm(next_dim),
                nn.GELU(),
                nn.Dropout(0.1)
            ])
            current_dim = next_dim

        self.readout_common = nn.Sequential(*readout_layers)
        self.final_readout_dim = current_dim

        # Main decoder
        self.main_decoder = nn.Linear(self.final_readout_dim, self.num_electrodes)

        # Gamma band-specific decoders
        self.gamma_decoders = nn.ModuleList([
            nn.Linear(self.final_readout_dim, self.num_electrodes)
            for _ in config.FREQ_BANDS
        ])

        # Time-specific prediction heads
        self.timestep_heads = nn.ModuleList([
            nn.Linear(self.num_electrodes * (len(config.FREQ_BANDS) + 1), self.num_electrodes)
            for _ in range(config.PREDICT_SIZE)
        ])

    def forward(self, x_dict, return_states=False):
        x_raw = x_dict['raw']
        x_phase = x_dict['phase']
        x_amplitude = x_dict['amplitude']
        x_inst_freq = x_dict['inst_freq']

        batch_size = x_raw.shape[0]
        context_size = x_raw.shape[1]
        device = x_raw.device

        # Process each timestep
        sequence = []
        for t in range(context_size):
            # Raw features
            raw_t = x_raw[:, t, :]
            raw_features = self.raw_projection(raw_t)

            # Phase features
            phase_t = x_phase[:, :, t, :].reshape(batch_size, -1)
            phase_features = self.phase_projection(phase_t)

            # Amplitude features
            amplitude_t = x_amplitude[:, :, t, :].reshape(batch_size, -1)
            amplitude_features = self.amplitude_projection(amplitude_t)

            # Frequency features
            freq_t = x_inst_freq[:, :, t, :].reshape(batch_size, -1)
            freq_features = self.freq_projection(freq_t)

            # Combine features
            combined_features = torch.cat([
                raw_features, phase_features, amplitude_features, freq_features
            ], dim=1)

            # Apply feature fusion
            fused_features = self.feature_fusion(combined_features)
            sequence.append(fused_features)

        # Stack into sequence
        sequence_tensor = torch.stack(sequence, dim=1)  # [batch, time, features]

        # Pass through LSTM
        lstm_out, (h_n, c_n) = self.lstm(sequence_tensor)

        # Use final hidden state from both directions
        h_forward = h_n[-2, :, :]
        h_backward = h_n[-1, :, :]
        bidirectional_hidden = torch.cat([h_forward, h_backward], dim=1)
        final_hidden = self.bidirectional_proj(bidirectional_hidden)

        # Apply common readout
        common_features = self.readout_common(final_hidden)

        # Decoder outputs
        main_output = self.main_decoder(common_features)
        gamma_outputs = [decoder(common_features) for decoder in self.gamma_decoders]

        # Combine all outputs
        all_outputs_stacked = torch.stack([main_output] + gamma_outputs, dim=1)
        timestep_input_reshaped = all_outputs_stacked.reshape(batch_size, -1)

        # Predict each timestep
        predictions = torch.zeros(batch_size, self.predict_size, self.num_electrodes, device=device)
        for t_pred in range(self.predict_size):
            predictions[:, t_pred, :] = self.timestep_heads[t_pred](timestep_input_reshaped)

        # Optionally return states for visualization
        states = lstm_out if return_states else None
        return (predictions, states) if return_states else predictions


class EnhancedGRUModel(nn.Module):
    """GRU model optimized for gamma oscillation prediction."""
    def __init__(self, config):
        super().__init__()
        self.config = config

        # Input dimensions
        self.num_electrodes = config.NUM_ELECTRODES
        self.num_freq_bands = len(config.FREQ_BANDS)
        self.predict_size = config.PREDICT_SIZE

        # Enhanced feature projections
        self.raw_projection = nn.Linear(self.num_electrodes, config.HIDDEN_SIZE // 4)
        self.phase_projection = nn.Linear(self.num_freq_bands * self.num_electrodes, config.HIDDEN_SIZE // 4)
        self.amplitude_projection = nn.Linear(self.num_freq_bands * self.num_electrodes, config.HIDDEN_SIZE // 4)
        self.freq_projection = nn.Linear(self.num_freq_bands * self.num_electrodes, config.HIDDEN_SIZE // 4)

        # Feature fusion
        self.feature_fusion = nn.Sequential(
            nn.Linear(config.HIDDEN_SIZE, config.HIDDEN_SIZE),
            nn.LayerNorm(config.HIDDEN_SIZE),
            nn.GELU(),
            nn.Dropout(config.DROPOUT),
            nn.Linear(config.HIDDEN_SIZE, config.HIDDEN_SIZE),
            nn.LayerNorm(config.HIDDEN_SIZE),
            nn.GELU()
        )

        # GRU layers
        self.gru = nn.GRU(
            input_size=config.HIDDEN_SIZE,
            hidden_size=config.READOUT_HIDDEN,
            num_layers=3,
            batch_first=True,
            dropout=config.DROPOUT,
            bidirectional=True
        )

        # Bidirectional projection
        self.bidirectional_proj = nn.Linear(config.READOUT_HIDDEN * 2, config.READOUT_HIDDEN)

        # Common readout layers
        readout_layers = []
        current_dim = config.READOUT_HIDDEN
        for i in range(config.READOUT_LAYERS - 1):
            next_dim = current_dim // 2
            readout_layers.extend([
                nn.Linear(current_dim, next_dim),
                nn.LayerNorm(next_dim),
                nn.GELU(),
                nn.Dropout(0.1)
            ])
            current_dim = next_dim

        self.readout_common = nn.Sequential(*readout_layers)
        self.final_readout_dim = current_dim

        # Main decoder
        self.main_decoder = nn.Linear(self.final_readout_dim, self.num_electrodes)

        # Gamma band-specific decoders
        self.gamma_decoders = nn.ModuleList([
            nn.Linear(self.final_readout_dim, self.num_electrodes)
            for _ in config.FREQ_BANDS
        ])

        # Time-specific prediction heads
        self.timestep_heads = nn.ModuleList([
            nn.Linear(self.num_electrodes * (len(config.FREQ_BANDS) + 1), self.num_electrodes)
            for _ in range(config.PREDICT_SIZE)
        ])

    def forward(self, x_dict, return_states=False):
        x_raw = x_dict['raw']
        x_phase = x_dict['phase']
        x_amplitude = x_dict['amplitude']
        x_inst_freq = x_dict['inst_freq']

        batch_size = x_raw.shape[0]
        context_size = x_raw.shape[1]
        device = x_raw.device

        # Process each timestep
        sequence = []
        for t in range(context_size):
            # Raw features
            raw_t = x_raw[:, t, :]
            raw_features = self.raw_projection(raw_t)

            # Phase features
            phase_t = x_phase[:, :, t, :].reshape(batch_size, -1)
            phase_features = self.phase_projection(phase_t)

            # Amplitude features
            amplitude_t = x_amplitude[:, :, t, :].reshape(batch_size, -1)
            amplitude_features = self.amplitude_projection(amplitude_t)

            # Frequency features
            freq_t = x_inst_freq[:, :, t, :].reshape(batch_size, -1)
            freq_features = self.freq_projection(freq_t)

            # Combine features
            combined_features = torch.cat([
                raw_features, phase_features, amplitude_features, freq_features
            ], dim=1)

            # Apply feature fusion
            fused_features = self.feature_fusion(combined_features)
            sequence.append(fused_features)

        # Stack into sequence
        sequence_tensor = torch.stack(sequence, dim=1)  # [batch, time, features]

        # Pass through GRU
        gru_out, h_n = self.gru(sequence_tensor)

        # Use final hidden state from both directions
        h_forward = h_n[-2, :, :]
        h_backward = h_n[-1, :, :]
        bidirectional_hidden = torch.cat([h_forward, h_backward], dim=1)
        final_hidden = self.bidirectional_proj(bidirectional_hidden)

        # Apply common readout
        common_features = self.readout_common(final_hidden)

        # Decoder outputs
        main_output = self.main_decoder(common_features)
        gamma_outputs = [decoder(common_features) for decoder in self.gamma_decoders]

        # Combine all outputs
        all_outputs_stacked = torch.stack([main_output] + gamma_outputs, dim=1)
        timestep_input_reshaped = all_outputs_stacked.reshape(batch_size, -1)

        # Predict each timestep
        predictions = torch.zeros(batch_size, self.predict_size, self.num_electrodes, device=device)
        for t_pred in range(self.predict_size):
            predictions[:, t_pred, :] = self.timestep_heads[t_pred](timestep_input_reshaped)

        # Optionally return states for visualization
        states = gru_out if return_states else None
        return (predictions, states) if return_states else predictions


class PositionalEncoding(nn.Module):
    def __init__(self, d_model, max_len=5000):
        super().__init__()
        pe = torch.zeros(max_len, d_model)
        position = torch.arange(0, max_len, dtype=torch.float).unsqueeze(1)
        div_term = torch.exp(torch.arange(0, d_model, 2).float() * (-math.log(10000.0) / d_model))
        pe[:, 0::2] = torch.sin(position * div_term)
        pe[:, 1::2] = torch.cos(position * div_term)
        pe = pe.unsqueeze(0)
        self.register_buffer('pe', pe)

    def forward(self, x):
        return x + self.pe[:, :x.size(1)]

class EnhancedTransformerModel(nn.Module):
    """Transformer model optimized for gamma oscillation prediction."""
    def __init__(self, config):
        super().__init__()
        self.config = config

        # Input dimensions
        self.num_electrodes = config.NUM_ELECTRODES
        self.num_freq_bands = len(config.FREQ_BANDS)
        self.predict_size = config.PREDICT_SIZE

        # Enhanced feature projections
        self.raw_projection = nn.Linear(self.num_electrodes, config.HIDDEN_SIZE // 4)
        self.phase_projection = nn.Linear(self.num_freq_bands * self.num_electrodes, config.HIDDEN_SIZE // 4)
        self.amplitude_projection = nn.Linear(self.num_freq_bands * self.num_electrodes, config.HIDDEN_SIZE // 4)
        self.freq_projection = nn.Linear(self.num_freq_bands * self.num_electrodes, config.HIDDEN_SIZE // 4)

        # Feature fusion
        self.feature_fusion = nn.Sequential(
            nn.Linear(config.HIDDEN_SIZE, config.HIDDEN_SIZE),
            nn.LayerNorm(config.HIDDEN_SIZE),
            nn.GELU(),
            nn.Dropout(config.DROPOUT),
            nn.Linear(config.HIDDEN_SIZE, config.HIDDEN_SIZE),
            nn.LayerNorm(config.HIDDEN_SIZE),
            nn.GELU()
        )

        # Positional encoding
        self.pos_encoder = PositionalEncoding(config.HIDDEN_SIZE)

        # Transformer encoder
        encoder_layers = nn.TransformerEncoderLayer(
            d_model=config.HIDDEN_SIZE,
            nhead=config.ATTENTION_HEADS,
            dim_feedforward=config.READOUT_HIDDEN,
            dropout=config.DROPOUT,
            batch_first=True
        )
        self.transformer_encoder = nn.TransformerEncoder(encoder_layers, num_layers=3)

        # Projection
        self.encoder_proj = nn.Linear(config.HIDDEN_SIZE, config.READOUT_HIDDEN)

        # Common readout layers
        readout_layers = []
        current_dim = config.READOUT_HIDDEN
        for i in range(config.READOUT_LAYERS - 1):
            next_dim = current_dim // 2
            readout_layers.extend([
                nn.Linear(current_dim, next_dim),
                nn.LayerNorm(next_dim),
                nn.GELU(),
                nn.Dropout(0.1)
            ])
            current_dim = next_dim

        self.readout_common = nn.Sequential(*readout_layers)
        self.final_readout_dim = current_dim

        # Main decoder
        self.main_decoder = nn.Linear(self.final_readout_dim, self.num_electrodes)

        # Gamma band-specific decoders
        self.gamma_decoders = nn.ModuleList([
            nn.Linear(self.final_readout_dim, self.num_electrodes)
            for _ in config.FREQ_BANDS
        ])

        # Time-specific prediction heads
        self.timestep_heads = nn.ModuleList([
            nn.Linear(self.num_electrodes * (len(config.FREQ_BANDS) + 1), self.num_electrodes)
            for _ in range(config.PREDICT_SIZE)
        ])

    def forward(self, x_dict, return_states=False):
        x_raw = x_dict['raw']
        x_phase = x_dict['phase']
        x_amplitude = x_dict['amplitude']
        x_inst_freq = x_dict['inst_freq']

        batch_size = x_raw.shape[0]
        context_size = x_raw.shape[1]
        device = x_raw.device

        # Process each timestep
        sequence = []
        for t in range(context_size):
            # Raw features
            raw_t = x_raw[:, t, :]
            raw_features = self.raw_projection(raw_t)

            # Phase features
            phase_t = x_phase[:, :, t, :].reshape(batch_size, -1)
            phase_features = self.phase_projection(phase_t)

            # Amplitude features
            amplitude_t = x_amplitude[:, :, t, :].reshape(batch_size, -1)
            amplitude_features = self.amplitude_projection(amplitude_t)

            # Frequency features
            freq_t = x_inst_freq[:, :, t, :].reshape(batch_size, -1)
            freq_features = self.freq_projection(freq_t)

            # Combine features
            combined_features = torch.cat([
                raw_features, phase_features, amplitude_features, freq_features
            ], dim=1)

            # Apply feature fusion
            fused_features = self.feature_fusion(combined_features)
            sequence.append(fused_features)

        # Stack into sequence
        sequence_tensor = torch.stack(sequence, dim=1)  # [batch, time, features]

        # Apply positional encoding
        sequence_tensor = self.pos_encoder(sequence_tensor)

        # Pass through transformer encoder
        transformer_output = self.transformer_encoder(sequence_tensor)

        # Take the output of the last timestep
        final_encoding = transformer_output[:, -1, :]

        # Project to readout dimension
        final_hidden = self.encoder_proj(final_encoding)

        # Apply common readout
        common_features = self.readout_common(final_hidden)

        # Decoder outputs
        main_output = self.main_decoder(common_features)
        gamma_outputs = [decoder(common_features) for decoder in self.gamma_decoders]

        # Combine all outputs
        all_outputs_stacked = torch.stack([main_output] + gamma_outputs, dim=1)
        timestep_input_reshaped = all_outputs_stacked.reshape(batch_size, -1)

        # Predict each timestep
        predictions = torch.zeros(batch_size, self.predict_size, self.num_electrodes, device=device)
        for t_pred in range(self.predict_size):
            predictions[:, t_pred, :] = self.timestep_heads[t_pred](timestep_input_reshaped)

        # Optionally return states for visualization
        states = transformer_output if return_states else None
        return (predictions, states) if return_states else predictions


class PhysicsInformedNN(nn.Module):
    """Physics-informed neural network for gamma oscillation prediction."""
    def __init__(self, config):
        super().__init__()
        self.config = config

        # Input dimensions
        self.num_electrodes = config.NUM_ELECTRODES
        self.num_freq_bands = len(config.FREQ_BANDS)
        self.predict_size = config.PREDICT_SIZE
        self.sampling_rate = config.SAMPLING_RATE

        # Enhanced feature projections
        self.raw_projection = nn.Linear(self.num_electrodes, config.HIDDEN_SIZE // 4)
        self.phase_projection = nn.Linear(self.num_freq_bands * self.num_electrodes, config.HIDDEN_SIZE // 4)
        self.amplitude_projection = nn.Linear(self.num_freq_bands * self.num_electrodes, config.HIDDEN_SIZE // 4)
        self.freq_projection = nn.Linear(self.num_freq_bands * self.num_electrodes, config.HIDDEN_SIZE // 4)

        # Feature fusion
        self.feature_fusion = nn.Sequential(
            nn.Linear(config.HIDDEN_SIZE, config.HIDDEN_SIZE),
            nn.LayerNorm(config.HIDDEN_SIZE),
            nn.GELU(),
            nn.Dropout(config.DROPOUT),
            nn.Linear(config.HIDDEN_SIZE, config.HIDDEN_SIZE),
            nn.LayerNorm(config.HIDDEN_SIZE),
            nn.GELU()
        )

        # LSTM for sequential processing
        self.lstm = nn.LSTM(
            input_size=config.HIDDEN_SIZE,
            hidden_size=config.READOUT_HIDDEN,
            num_layers=2,
            batch_first=True,
            dropout=config.DROPOUT
        )

        # Physics-informed layers (oscillatory modeling)
        self.oscillator_params = nn.Parameter(torch.randn(self.num_electrodes, 3))  # [freq, damping, phase]

        # Readout layers
        readout_layers = []
        current_dim = config.READOUT_HIDDEN
        for i in range(config.READOUT_LAYERS - 1):
            next_dim = current_dim // 2
            readout_layers.extend([
                nn.Linear(current_dim, next_dim),
                nn.LayerNorm(next_dim),
                nn.GELU(),
                nn.Dropout(0.1)
            ])
            current_dim = next_dim

        self.readout_common = nn.Sequential(*readout_layers)
        self.final_readout_dim = current_dim

        # Frequency band-specific decoders for gamma oscillations
        self.freq_band_decoders = nn.ModuleList([
            nn.Linear(self.final_readout_dim, self.num_electrodes)
            for _ in range(len(config.FREQ_BANDS))
        ])

        # Main decoder
        self.main_decoder = nn.Linear(self.final_readout_dim, self.num_electrodes)

        # Time-specific prediction heads
        self.timestep_heads = nn.ModuleList([
            nn.Linear(self.num_electrodes * (len(config.FREQ_BANDS) + 1), self.num_electrodes)
            for _ in range(config.PREDICT_SIZE)
        ])

    def oscillator_term(self, time_steps, electrode_idx):
        """Calculate oscillatory terms based on physics."""
        # Extract parameters for this electrode
        freq = torch.abs(self.oscillator_params[electrode_idx, 0]) * 100 + 30  # 30-130 Hz (gamma range)
        damping = torch.sigmoid(self.oscillator_params[electrode_idx, 1]) * 0.5  # 0-0.5 damping factor
        phase = self.oscillator_params[electrode_idx, 2] * torch.pi  # 0-π phase

        # Convert frequency to angular frequency
        omega = 2 * torch.pi * freq / self.sampling_rate

        # Calculate oscillatory terms
        t = time_steps.float()
        oscillation = torch.exp(-damping * t) * torch.sin(omega * t + phase)

        return oscillation

    def forward(self, x_dict, return_states=False):
        x_raw = x_dict['raw']
        x_phase = x_dict['phase']
        x_amplitude = x_dict['amplitude']
        x_inst_freq = x_dict['inst_freq']

        batch_size = x_raw.shape[0]
        context_size = x_raw.shape[1]
        device = x_raw.device

        # Process each timestep
        sequence = []
        for t in range(context_size):
            # Raw features
            raw_t = x_raw[:, t, :]
            raw_features = self.raw_projection(raw_t)

            # Phase features
            phase_t = x_phase[:, :, t, :].reshape(batch_size, -1)
            phase_features = self.phase_projection(phase_t)

            # Amplitude features
            amplitude_t = x_amplitude[:, :, t, :].reshape(batch_size, -1)
            amplitude_features = self.amplitude_projection(amplitude_t)

            # Frequency features
            freq_t = x_inst_freq[:, :, t, :].reshape(batch_size, -1)
            freq_features = self.freq_projection(freq_t)

            # Combine features
            combined_features = torch.cat([
                raw_features, phase_features, amplitude_features, freq_features
            ], dim=1)

            # Apply feature fusion
            fused_features = self.feature_fusion(combined_features)
            sequence.append(fused_features)

        # Stack into sequence
        sequence_tensor = torch.stack(sequence, dim=1)  # [batch, time, features]

        # Pass through LSTM
        lstm_out, (h_n, c_n) = self.lstm(sequence_tensor)

        # Take the final hidden state
        final_hidden = h_n[-1]

        # Apply common readout
        common_features = self.readout_common(final_hidden)

        # Decoder outputs
        main_output = self.main_decoder(common_features)
        band_outputs = [decoder(common_features) for decoder in self.freq_band_decoders]

        # Combine all outputs
        all_outputs_stacked = torch.stack([main_output] + band_outputs, dim=1)
        timestep_input_reshaped = all_outputs_stacked.reshape(batch_size, -1)

        # Predict each timestep using neural and physics components
        predictions = torch.zeros(batch_size, self.predict_size, self.num_electrodes, device=device)

        # Generate time steps for physics-based oscillations
        time_steps = torch.arange(0, self.predict_size, device=device).unsqueeze(0).repeat(batch_size, 1)

        for t_pred in range(self.predict_size):
            # Neural prediction component
            neural_pred = self.timestep_heads[t_pred](timestep_input_reshaped)

            # Physics-based oscillatory component
            physics_component = torch.zeros(batch_size, self.num_electrodes, device=device)
            for e_idx in range(self.num_electrodes):
                # Calculate oscillatory term for this electrode
                oscillation = self.oscillator_term(time_steps[:, t_pred], e_idx)
                physics_component[:, e_idx] = oscillation

            # Combine neural and physics components
            predictions[:, t_pred, :] = neural_pred + 0.3 * physics_component

        # Optionally return states for visualization
        states = lstm_out if return_states else None
        return (predictions, states) if return_states else predictions

class CausalConv1d(nn.Module):
    def __init__(self, in_channels, out_channels, kernel_size, dilation=1):
        super().__init__()
        padding = (kernel_size - 1) * dilation
        self.conv = nn.Conv1d(in_channels, out_channels, kernel_size,
                             padding=padding, dilation=dilation)

    def forward(self, x):
        # x: [batch, channels, time]
        out = self.conv(x)
        # Remove padding from the end
        return out[:, :, :-self.conv.padding[0]]

class TCNBlock(nn.Module):
    def __init__(self, in_channels, out_channels, kernel_size, dilation, dropout=0.2):
        super().__init__()
        self.conv1 = CausalConv1d(in_channels, out_channels, kernel_size, dilation)
        self.norm1 = nn.BatchNorm1d(out_channels)
        self.relu1 = nn.ReLU()
        self.dropout1 = nn.Dropout(dropout)

        self.conv2 = CausalConv1d(out_channels, out_channels, kernel_size, dilation)
        self.norm2 = nn.BatchNorm1d(out_channels)
        self.relu2 = nn.ReLU()
        self.dropout2 = nn.Dropout(dropout)

        self.skip = nn.Conv1d(in_channels, out_channels, 1) if in_channels != out_channels else nn.Identity()

    def forward(self, x):
        residual = self.skip(x)

        out = self.conv1(x)
        out = self.norm1(out)
        out = self.relu1(out)
        out = self.dropout1(out)

        out = self.conv2(out)
        out = self.norm2(out)
        out = self.relu2(out)
        out = self.dropout2(out)

        return out + residual

class EnhancedTCNModel(nn.Module):
    """TCN model optimized for gamma oscillation prediction."""
    def __init__(self, config):
        super().__init__()
        self.config = config

        # Input dimensions
        self.num_electrodes = config.NUM_ELECTRODES
        self.num_freq_bands = len(config.FREQ_BANDS)
        self.predict_size = config.PREDICT_SIZE

        # Enhanced feature projections
        self.raw_projection = nn.Linear(self.num_electrodes, config.HIDDEN_SIZE // 4)
        self.phase_projection = nn.Linear(self.num_freq_bands * self.num_electrodes, config.HIDDEN_SIZE // 4)
        self.amplitude_projection = nn.Linear(self.num_freq_bands * self.num_electrodes, config.HIDDEN_SIZE // 4)
        self.freq_projection = nn.Linear(self.num_freq_bands * self.num_electrodes, config.HIDDEN_SIZE // 4)

        # Feature fusion
        self.feature_fusion = nn.Sequential(
            nn.Linear(config.HIDDEN_SIZE, config.HIDDEN_SIZE),
            nn.LayerNorm(config.HIDDEN_SIZE),
            nn.GELU(),
            nn.Dropout(config.DROPOUT),
            nn.Linear(config.HIDDEN_SIZE, config.HIDDEN_SIZE),
            nn.LayerNorm(config.HIDDEN_SIZE),
            nn.GELU()
        )

        # TCN Layers with increasing dilation
        layers = []
        num_levels = 4
        kernel_size = 3
        channels = config.HIDDEN_SIZE

        for i in range(num_levels):
            dilation = 2 ** i
            layers.append(TCNBlock(channels, channels, kernel_size, dilation, dropout=config.DROPOUT))

        self.tcn = nn.Sequential(*layers)

        # Readout projection
        self.readout_proj = nn.Linear(channels, config.READOUT_HIDDEN)

        # Common readout layers
        readout_layers = []
        current_dim = config.READOUT_HIDDEN
        for i in range(config.READOUT_LAYERS - 1):
            next_dim = current_dim // 2
            readout_layers.extend([
                nn.Linear(current_dim, next_dim),
                nn.LayerNorm(next_dim),
                nn.GELU(),
                nn.Dropout(0.1)
            ])
            current_dim = next_dim

        self.readout_common = nn.Sequential(*readout_layers)
        self.final_readout_dim = current_dim

        # Main decoder
        self.main_decoder = nn.Linear(self.final_readout_dim, self.num_electrodes)

        # Gamma band-specific decoders
        self.gamma_decoders = nn.ModuleList([
            nn.Linear(self.final_readout_dim, self.num_electrodes)
            for _ in config.FREQ_BANDS
        ])

        # Time-specific prediction heads
        self.timestep_heads = nn.ModuleList([
            nn.Linear(self.num_electrodes * (len(config.FREQ_BANDS) + 1), self.num_electrodes)
            for _ in range(config.PREDICT_SIZE)
        ])

    def forward(self, x_dict, return_states=False):
        x_raw = x_dict['raw']
        x_phase = x_dict['phase']
        x_amplitude = x_dict['amplitude']
        x_inst_freq = x_dict['inst_freq']

        batch_size = x_raw.shape[0]
        context_size = x_raw.shape[1]
        device = x_raw.device

        # Process each timestep
        sequence = []
        for t in range(context_size):
            # Raw features
            raw_t = x_raw[:, t, :]
            raw_features = self.raw_projection(raw_t)

            # Phase features
            phase_t = x_phase[:, :, t, :].reshape(batch_size, -1)
            phase_features = self.phase_projection(phase_t)

            # Amplitude features
            amplitude_t = x_amplitude[:, :, t, :].reshape(batch_size, -1)
            amplitude_features = self.amplitude_projection(amplitude_t)

            # Frequency features
            freq_t = x_inst_freq[:, :, t, :].reshape(batch_size, -1)
            freq_features = self.freq_projection(freq_t)

            # Combine features
            combined_features = torch.cat([
                raw_features, phase_features, amplitude_features, freq_features
            ], dim=1)

            # Apply feature fusion
            fused_features = self.feature_fusion(combined_features)
            sequence.append(fused_features)

        # Stack into sequence
        sequence_tensor = torch.stack(sequence, dim=1)  # [batch, time, channels]

        # Transpose for TCN [batch, channels, time]
        sequence_tensor = sequence_tensor.transpose(1, 2)

        # Apply TCN
        tcn_out = self.tcn(sequence_tensor)

        # Get final timestep output [batch, channels, 1]
        final_output = tcn_out[:, :, -1]

        # Project to readout dimension
        readout_input = self.readout_proj(final_output)

        # Apply common readout
        common_features = self.readout_common(readout_input)

        # Decoder outputs
        main_output = self.main_decoder(common_features)
        gamma_outputs = [decoder(common_features) for decoder in self.gamma_decoders]

        # Combine all outputs
        all_outputs_stacked = torch.stack([main_output] + gamma_outputs, dim=1)
        timestep_input_reshaped = all_outputs_stacked.reshape(batch_size, -1)

        # Predict each timestep
        predictions = torch.zeros(batch_size, self.predict_size, self.num_electrodes, device=device)
        for t_pred in range(self.predict_size):
            predictions[:, t_pred, :] = self.timestep_heads[t_pred](timestep_input_reshaped)

        # Optionally return states for visualization
        states = tcn_out.transpose(1, 2) if return_states else None
        return (predictions, states) if return_states else predictions

class MorletWaveletLayer(nn.Module):
    """Morlet wavelet transform layer for time-frequency analysis."""
    def __init__(self, num_wavelets, input_dim, device=None):
        super().__init__()
        self.num_wavelets = num_wavelets
        self.input_dim = input_dim

        # Learnable wavelet parameters
        self.scales = nn.Parameter(torch.linspace(1, 50, num_wavelets))
        self.shifts = nn.Parameter(torch.zeros(num_wavelets))

        # Morlet wavelet central frequency
        self.omega0 = 5.0

        # Generate time grid once
        t = torch.linspace(-5, 5, input_dim)
        self.register_buffer('t', t)


    def morlet_wavelet(self, t, scale, shift):
        """Compute Morlet wavelet."""
        t_new = (t - shift) / scale
        wavelet = torch.exp(1j * self.omega0 * t_new) * torch.exp(-0.5 * t_new * t_new)
        return wavelet

    def forward(self, x):
        """
        Apply wavelet transform.
        x: [batch_size, time_steps]
        """
        batch_size = x.shape[0]

        # Initialize output
        output = torch.zeros(batch_size, self.num_wavelets, 2, device=x.device)

        # Apply wavelet transform for each wavelet
        for i in range(self.num_wavelets):
            # Generate wavelet
            wavelet = self.morlet_wavelet(self.t, self.scales[i], self.shifts[i])

            # Apply convolution
            real_part = F.conv1d(x.unsqueeze(1),
                                wavelet.real.view(1, 1, -1),
                                padding=self.input_dim//2)
            imag_part = F.conv1d(x.unsqueeze(1),
                                wavelet.imag.view(1, 1, -1),
                                padding=self.input_dim//2)

            # Get amplitude and phase
            amplitude = torch.sqrt(real_part**2 + imag_part**2)
            phase = torch.atan2(imag_part, real_part)

            # Use central point
            output[:, i, 0] = amplitude.squeeze()[:, self.input_dim//2]
            output[:, i, 1] = phase.squeeze()[:, self.input_dim//2]

        return output.reshape(batch_size, -1)

class WaveletNeuralNetwork(nn.Module):
    """Wavelet-based neural network for gamma oscillation prediction."""
    def __init__(self, config):
        super().__init__()
        self.config = config

        # Input dimensions
        self.num_electrodes = config.NUM_ELECTRODES
        self.num_freq_bands = len(config.FREQ_BANDS)
        self.predict_size = config.PREDICT_SIZE
        self.context_size = config.CONTEXT_SIZE

        # Wavelet transform modules for each electrode
        self.wavelet_layers = nn.ModuleList([
            MorletWaveletLayer(10, self.context_size) for _ in range(self.num_electrodes)
        ])

        # Input projection
        self.input_dim = 10 * 2 * self.num_electrodes  # num_wavelets * 2 (mag/phase) * electrodes

        # Enhanced feature projections for other inputs
        self.phase_projection = nn.Linear(self.num_freq_bands * self.num_electrodes, config.HIDDEN_SIZE // 4)
        self.amplitude_projection = nn.Linear(self.num_freq_bands * self.num_electrodes, config.HIDDEN_SIZE // 4)
        self.freq_projection = nn.Linear(self.num_freq_bands * self.num_electrodes, config.HIDDEN_SIZE // 4)

        # Projection for wavelet features
        self.wavelet_projection = nn.Linear(self.input_dim, config.HIDDEN_SIZE // 4)

        # Feature fusion
        self.feature_fusion = nn.Sequential(
            nn.Linear(config.HIDDEN_SIZE, config.HIDDEN_SIZE),
            nn.LayerNorm(config.HIDDEN_SIZE),
            nn.GELU(),
            nn.Dropout(config.DROPOUT),
            nn.Linear(config.HIDDEN_SIZE, config.HIDDEN_SIZE),
            nn.LayerNorm(config.HIDDEN_SIZE),
            nn.GELU()
        )

        # Temporal processing with GRU
        self.gru = nn.GRU(
            input_size=config.HIDDEN_SIZE,
            hidden_size=config.READOUT_HIDDEN,
            num_layers=2,
            batch_first=True,
            dropout=config.DROPOUT,
            bidirectional=True
        )

        # Bidirectional projection
        self.bidirectional_proj = nn.Linear(config.READOUT_HIDDEN * 2, config.READOUT_HIDDEN)

        # Common readout layers
        readout_layers = []
        current_dim = config.READOUT_HIDDEN
        for i in range(config.READOUT_LAYERS - 1):
            next_dim = current_dim // 2
            readout_layers.extend([
                nn.Linear(current_dim, next_dim),
                nn.LayerNorm(next_dim),
                nn.GELU(),
                nn.Dropout(0.1)
            ])
            current_dim = next_dim

        self.readout_common = nn.Sequential(*readout_layers)
        self.final_readout_dim = current_dim

        # Main decoder
        self.main_decoder = nn.Linear(self.final_readout_dim, self.num_electrodes)

        # Gamma band-specific decoders
        self.gamma_decoders = nn.ModuleList([
            nn.Linear(self.final_readout_dim, self.num_electrodes)
            for _ in config.FREQ_BANDS
        ])

        # Time-specific prediction heads
        self.timestep_heads = nn.ModuleList([
            nn.Linear(self.num_electrodes * (len(config.FREQ_BANDS) + 1), self.num_electrodes)
            for _ in range(config.PREDICT_SIZE)
        ])

    def forward(self, x_dict, return_states=False):
        x_raw = x_dict['raw']
        x_phase = x_dict['phase']
        x_amplitude = x_dict['amplitude']
        x_inst_freq = x_dict['inst_freq']

        batch_size = x_raw.shape[0]
        context_size = x_raw.shape[1]
        device = x_raw.device

        # Apply wavelet transform to each electrode signal
        wavelet_features_all = []
        for e in range(self.num_electrodes):
            wavelet_features = self.wavelet_layers[e](x_raw[:, :, e])
            wavelet_features_all.append(wavelet_features)

        # Concatenate wavelet features for all electrodes
        wavelet_features_concat = torch.cat(wavelet_features_all, dim=1)

        # Process standard features for each timestep
        sequence = []
        for t in range(context_size):
            # Get phase features
            phase_t = x_phase[:, :, t, :].reshape(batch_size, -1)
            phase_features = self.phase_projection(phase_t)

            # Get amplitude features
            amplitude_t = x_amplitude[:, :, t, :].reshape(batch_size, -1)
            amplitude_features = self.amplitude_projection(amplitude_t)

            # Get frequency features
            freq_t = x_inst_freq[:, :, t, :].reshape(batch_size, -1)
            freq_features = self.freq_projection(freq_t)

            # Project wavelet features
            wavelet_projection = self.wavelet_projection(wavelet_features_concat)

            # Combine all features
            combined_features = torch.cat([
                wavelet_projection, phase_features, amplitude_features, freq_features
            ], dim=1)

            # Apply feature fusion
            fused_features = self.feature_fusion(combined_features)
            sequence.append(fused_features)

        # Stack into sequence
        sequence_tensor = torch.stack(sequence, dim=1)  # [batch, time, features]

        # Process with GRU
        gru_out, h_n = self.gru(sequence_tensor)

        # Use final hidden state from both directions
        h_forward = h_n[-2, :, :]
        h_backward = h_n[-1, :, :]
        bidirectional_hidden = torch.cat([h_forward, h_backward], dim=1)
        final_hidden = self.bidirectional_proj(bidirectional_hidden)

        # Apply common readout
        common_features = self.readout_common(final_hidden)

        # Decoder outputs
        main_output = self.main_decoder(common_features)
        gamma_outputs = [decoder(common_features) for decoder in self.gamma_decoders]

        # Combine all outputs
        all_outputs_stacked = torch.stack([main_output] + gamma_outputs, dim=1)
        timestep_input_reshaped = all_outputs_stacked.reshape(batch_size, -1)

        # Predict each timestep
        predictions = torch.zeros(batch_size, self.predict_size, self.num_electrodes, device=device)
        for t_pred in range(self.predict_size):
            predictions[:, t_pred, :] = self.timestep_heads[t_pred](timestep_input_reshaped)

        # Optionally return states for visualization
        states = gru_out if return_states else None
        return (predictions, states) if return_states else predictions


def get_models(config, device=None):
    if device is None:
        device = torch.device('cuda' if torch.cuda.is_available() else 'cpu')

    models = {
        "RNN": EnhancedRNNModel(config).to(device),
        "LSTM": EnhancedLSTMModel(config).to(device),
        "GRU": EnhancedGRUModel(config).to(device),
        "Transformer": EnhancedTransformerModel(config).to(device),
        "PINN": PhysicsInformedNN(config).to(device),
        "TCN": EnhancedTCNModel(config).to(device),
        "WaveletNN": WaveletNeuralNetwork(config).to(device)
    }

    return models


def main(data_path=None):
    """
    Execute the full pipeline from data loading to comparative model evaluation.

    Args:
        data_path: Path to the dataset directory. If None, the function will attempt to locate it.
    """
    try:
        print("\n" + "="*80)
        print("NEURAL ARCHITECTURE PERFORMANCE COMPARISON FOR GAMMA OSCILLATION PREDICTION")
        print("="*80 + "\n")

        # Create directory for results
        os.makedirs(config.OUTPUT_DIR, exist_ok=True)

        # Step 1: Find dataset path
        if data_path is None:
            # Try to locate the dataset automatically
            if os.path.exists("/kaggle/input/lfp-grating"):
                data_path = "/kaggle/input/lfp-grating"
            elif os.path.exists("/content/lfp_grating"):
                data_path = "/content/lfp_grating"
            else:
                try:
                    import kagglehub
                    print("Attempting to download dataset via kagglehub...")
                    data_path = kagglehub.dataset_download("arunramponnambalam/lfp-grating")
                except:
                    print("kagglehub not available. Looking for local path...")

                    local_path_try = "./lfp-grating"
                    if os.path.exists(local_path_try):
                        data_path = local_path_try
                    else:
                        data_path = input("Enter path to lfp-grating dataset directory: ")

        if not os.path.exists(data_path):
            raise FileNotFoundError(f"Data path not found: {data_path}")

        print(f"Using dataset at: {data_path}")

        # Step 2: Load and preprocess data
        print("\nLoading electrode positions...")
        positions = load_electrode_positions(data_path)

        print("\nSelecting electrodes...")
        channel_nums, selected_indices = select_electrodes(positions, config.NUM_ELECTRODES)

        print("\nLoading LFP data...")
        lfp_data = load_lfp_data(data_path, channel_nums, max_rows=500000)

        print("\nExtracting windows...")
        windows = extract_windows(lfp_data,
                                 context_size=config.CONTEXT_SIZE,
                                 predict_size=config.PREDICT_SIZE,
                                 stride=config.STRIDE,
                                 max_windows=config.MAX_SEQUENCES)

        if windows.shape[0] == 0:
            raise ValueError("No windows extracted. Cannot proceed.")

        print("\nPreprocessing data...")
        train_loader, val_loader, test_loader, scaler = preprocess_data(
            windows,
            context_size=config.CONTEXT_SIZE,
            predict_size=config.PREDICT_SIZE
        )

        # Step 3: Comparative evaluation of different models
        print("\n" + "="*80)
        print("STARTING MODEL COMPARISON")
        print("="*80)

        # Create directory for model comparison
        comparison_dir = os.path.join(config.OUTPUT_DIR, 'model_comparison')
        os.makedirs(comparison_dir, exist_ok=True)

        # Select which models to run
        models_to_run = ["RNN", "LSTM", "GRU", "Transformer", "PINN", "TCN", "WaveletNN"]
        print(f"Models to evaluate: {', '.join(models_to_run)}")

        # Get all models
        all_models = get_models(config, device)

        # Dictionary to store results
        results = {}

        # Train and evaluate each model
        for model_name in models_to_run:
            if model_name not in all_models:
                print(f"Model {model_name} not found. Skipping.")
                continue

            try:
                model = all_models[model_name]
                print(f"\n{'-'*80}\nTraining {model_name} model\n{'-'*80}")

                # Track parameter count
                param_count = sum(p.numel() for p in model.parameters() if p.requires_grad)
                print(f"Model has {param_count:,} trainable parameters")

                # Create criterion
                criterion = EnhancedGammaLoss(config)

                # Create optimizer
                optimizer = torch.optim.AdamW(
                    model.parameters(), lr=config.LEARNING_RATE, weight_decay=config.WEIGHT_DECAY, amsgrad=True
                )

                # Use the same scheduler
                scheduler = ReduceLROnPlateau(
                    optimizer, mode='min', factor=0.5, patience=7, min_lr=1e-6, verbose=True
                )

                # Training variables
                best_val_loss = float('inf')
                patience_counter = 0
                history = {'train_loss': [], 'val_loss': [], 'train_metrics': [], 'val_metrics': [], 'test_metrics': None}

                # Training loop
                for epoch in range(config.EPOCHS):
                    print(f"Epoch {epoch+1}/{config.EPOCHS}")

                    # Train for one epoch
                    train_metrics = train_epoch(
                        model, train_loader, optimizer, criterion, scheduler, device, config, epoch, config.CLIP_GRAD_NORM
                    )

                    # Evaluate on validation set
                    val_metrics, _, _, _, _ = evaluate(model, val_loader, criterion, device)

                    # Update scheduler
                    scheduler.step(val_metrics['loss'])

                    # Store history
                    history['train_loss'].append(train_metrics['loss'])
                    history['val_loss'].append(val_metrics['loss'])
                    history['train_metrics'].append(train_metrics)
                    history['val_metrics'].append(val_metrics)

                    # Print progress
                    print(f"  Train loss: {train_metrics['loss']:.6f}, Val loss: {val_metrics['loss']:.6f}")

                    # Check for improvement
                    if val_metrics['loss'] < best_val_loss:
                        best_val_loss = val_metrics['loss']
                        patience_counter = 0
                        # Save best model
                        torch.save(
                            {
                                'epoch': epoch,
                                'model_state_dict': model.state_dict(),
                                'optimizer_state_dict': optimizer.state_dict(),
                                'val_metrics': val_metrics
                            },
                            os.path.join(comparison_dir, f'best_{model_name}_model.pt')
                        )
                        print(f"  New best model saved! Loss: {best_val_loss:.6f}")
                    else:
                        patience_counter += 1
                        print(f"  No improvement for {patience_counter}/{config.PATIENCE} epochs")
                        if patience_counter >= config.PATIENCE:
                            print(f"  Early stopping after {epoch+1} epochs")
                            break

                # Final evaluation on test set if provided
                if test_loader:
                    print(f"Evaluating {model_name} on test set...")
                    test_metrics, _, _, _, _ = evaluate(model, test_loader, criterion, device)
                    history['test_metrics'] = test_metrics
                    print(f"  Test loss: {test_metrics['loss']:.6f}")

                # Store results
                results[model_name] = {
                    'history': history,
                    'param_count': param_count,
                    'best_val_loss': min(history['val_loss']),
                    'test_metrics': history['test_metrics'] if test_loader else None
                }

                print(f"{model_name} training complete!")

            except Exception as e:
                print(f"Error training {model_name}: {e}")
                traceback.print_exc()

        # Print performance comparison
        print("\n\n" + "="*80)
        print("MODEL PERFORMANCE COMPARISON")
        print("="*80)

        # Create results table
        headers = ["Model", "Params", "Best Val Loss", "Val MSE", "Val Corr", "Val Spectrum Corr"]

        if test_loader:
            headers.extend(["Test MSE", "Test Corr", "Test Spectrum Corr"])

        print("\n{:<15} {:<12} {:<15} {:<10} {:<10} {:<10}".format(*headers[:6]), end="")
        if test_loader:
            print(" {:<10} {:<10} {:<10}".format(*headers[6:]), end="")
        print()
        print("-" * (15 + 12 + 15 + 10*3 + (30 if test_loader else 0)))

        for model_name, result in results.items():
            # Get best validation metrics
            val_metrics = result['history']['val_metrics']
            best_idx = result['history']['val_loss'].index(min(result['history']['val_loss']))
            best_val_metrics = val_metrics[best_idx]

            row = [
                model_name,
                f"{result['param_count']:,}",
                f"{result['best_val_loss']:.6f}",
                f"{best_val_metrics.get('mse', float('nan')):.6f}",
                f"{best_val_metrics.get('corr', float('nan')):.4f}",
                f"{best_val_metrics.get('spectrum_corr', float('nan')):.4f}"
            ]

            print("{:<15} {:<12} {:<15} {:<10} {:<10} {:<10}".format(*row), end="")

            # Add test metrics if available
            if test_loader and result['test_metrics']:
                test_metrics = result['test_metrics']
                test_row = [
                    f"{test_metrics.get('mse', float('nan')):.6f}",
                    f"{test_metrics.get('corr', float('nan')):.4f}",
                    f"{test_metrics.get('spectrum_corr', float('nan')):.4f}"
                ]
                print(" {:<10} {:<10} {:<10}".format(*test_row), end="")
            print()

        # Create detailed table for gamma band metrics if available
        if test_loader:
            print("\n\nGAMMA BAND CORRELATION METRICS (Test Set)")
            print("="*80)

            # Get band names
            band_names = [f"{low}-{high}Hz" for low, high in config.FREQ_BANDS]

            # Print header
            print("{:<15}".format("Model"), end="")
            for band in band_names:
                print(" {:<10}".format(band), end="")
            print(" {:<10}".format("Average"))
            print("-" * (15 + 11 * (len(band_names) + 1)))

            # Print each model's gamma band metrics
            for model_name, result in results.items():
                if result['test_metrics']:
                    test_metrics = result['test_metrics']
                    gamma_metrics = test_metrics.get('gamma_metrics', {})

                    print("{:<15}".format(model_name), end="")

                    if gamma_metrics:
                        band_correlations = []
                        for band in config.FREQ_BANDS:
                            corr = gamma_metrics.get(band, {}).get('corr', float('nan'))
                            band_correlations.append(corr)
                            print(" {:<10.4f}".format(corr), end="")

                        # Calculate average
                        avg_corr = np.nanmean([c for c in band_correlations if not np.isnan(c)])
                        print(" {:<10.4f}".format(avg_corr))
                    else:
                        print(" No gamma band metrics available")

        # Save results to file
        import json

        # Convert results to serializable format
        serializable_results = {}
        for model_name, result in results.items():
            serializable_results[model_name] = {
                'param_count': result['param_count'],
                'best_val_loss': float(result['best_val_loss']),
                'train_loss': [float(x) for x in result['history']['train_loss']],
                'val_loss': [float(x) for x in result['history']['val_loss']]
            }

            # Add test metrics if available
            if result['test_metrics']:
                test_metrics = result['test_metrics']
                serializable_results[model_name]['test_metrics'] = {
                    'loss': float(test_metrics['loss']),
                    'mse': float(test_metrics.get('mse', float('nan'))),
                    'corr': float(test_metrics.get('corr', float('nan'))),
                    'spectrum_corr': float(test_metrics.get('spectrum_corr', float('nan')))
                }

                # Add gamma band metrics if available
                if 'gamma_metrics' in test_metrics:
                    gamma_results = {}
                    for band, metrics in test_metrics['gamma_metrics'].items():
                        # Need to convert tuple keys to strings
                        band_str = f"{band[0]}-{band[1]}"
                        gamma_results[band_str] = {
                            'corr': float(metrics.get('corr', float('nan'))),
                            'mse': float(metrics.get('mse', float('nan'))),
                            'r2': float(metrics.get('r2', float('nan')))
                        }
                    serializable_results[model_name]['test_metrics']['gamma_metrics'] = gamma_results

        with open(os.path.join(comparison_dir, 'model_comparison_results.json'), 'w') as f:
            json.dump(serializable_results, f, indent=2)

        print(f"\nResults saved to {os.path.join(comparison_dir, 'model_comparison_results.json')}")
        return results

    except Exception as e_main:
        print(f"Error in main function: {e_main}")
        traceback.print_exc()
        return None

if __name__ == "__main__":
    # Execute the full pipeline
    results = main()
    print("\nComparative evaluation complete!")


# %% [code]
```
