## Supplementary material for "How Minds Take Shape: Graph-ESN Reveals How Neural Ensembles Engineer Stable Representations": GaborDS.ipynb

```
# %% [code]
# Install required libraries
!pip install kaggle kagglehub

# Import necessary libraries
import os
import kagglehub
from google.colab import files

# Upload Kaggle API token
print("Please upload your kaggle.json file")
uploaded = files.upload()
for fn in uploaded.keys():
    print('User uploaded file "{name}" with length {length} bytes'.format(
        name=fn, length=len(uploaded[fn])))

# Create Kaggle directory and move the token
!mkdir -p ~/.kaggle
!cp /content/kaggle.json ~/.kaggle/
!chmod 600 ~/.kaggle/kaggle.json

# Download the dataset
dataset_path = kagglehub.dataset_download('arunramponnambalam/ecog-landmark-mkn')

# Print the downloaded dataset path
print(f"Dataset downloaded to: {dataset_path}")

# List files in the downloaded dataset
import os
print("Files in the dataset:")
for root, dirs, files in os.walk(dataset_path):
    for file in files:
        print(os.path.join(root, file))

# Optional: Mount to a specific directory if needed
!mkdir -p /content//ecog-landmark-mkn
!cp -r {dataset_path}/* /content/ecog-landmark-mkn/

print("Dataset mounted to /ecog-landmark-mkn")

# %% [code]
import os
import numpy as np
import pandas as pd
import matplotlib.pyplot as plt
import torch
import torch.nn as nn
import torch.nn.functional as F
from torch.utils.data import Dataset, DataLoader, TensorDataset
from sklearn.model_selection import train_test_split
from sklearn.preprocessing import RobustScaler, StandardScaler, MinMaxScaler
from sklearn.metrics import mean_squared_error, r2_score
from scipy import signal
import warnings
from tqdm import tqdm
import time
import math
from torch.optim.lr_scheduler import CosineAnnealingLR, OneCycleLR, ReduceLROnPlateau
from scipy.signal import stft, hilbert, butter, sosfilt, filtfilt, chirp, spectrogram
import networkx as nx
from sklearn.manifold import TSNE
from sklearn.decomposition import PCA
import seaborn as sns
from torch.optim.swa_utils import AveragedModel, update_bn
from collections import defaultdict
import traceback

# For visualizations
import plotly.graph_objects as go
import plotly.express as px
import plotly.offline as pyo
import plotly.io as pio
from plotly.subplots import make_subplots
from scipy.stats import pearsonr
from scipy.ndimage import gaussian_filter1d
from statsmodels.tsa.stattools import grangercausalitytests
import base64
from tqdm import tqdm
from sklearn.cluster import KMeans
from scipy.spatial.distance import cosine, euclidean
from plotly.colors import sequential, qualitative

# Suppress warnings
warnings.filterwarnings('ignore')

# Check for GPU
device = torch.device('cuda' if torch.cuda.is_available() else 'cpu')
print(f"Using device: {device}")

# Set random seed for reproducibility
SEED = 42
np.random.seed(SEED)
torch.manual_seed(SEED)
if torch.cuda.is_available():
    torch.cuda.manual_seed_all(SEED)
    torch.backends.cudnn.deterministic = True
    torch.backends.cudnn.benchmark = False


class Config:
    def __init__(self):
        # Data parameters - update based on new dataset
        self.NUM_ELECTRODES = 64  # Choose a subset of the 95 available channels
        self.CONTEXT_SIZE = 100   # Size of input window
        self.PREDICT_SIZE = 50    # Size of prediction window
        self.WINDOW_SIZE = self.CONTEXT_SIZE + self.PREDICT_SIZE
        self.SAMPLING_RATE = 1000  # Based on the timestamp differences in the data
        self.MAX_SEQUENCES = 600    # Number of sequences to extract
        self.BATCH_SIZE = 16
        self.STRIDE = 50

        # Enhanced gamma frequency bands - keep as is
        self.FREQ_BANDS = [
            (30, 50),       # Low Gamma
            (50, 80),       # Mid-Low Gamma
            (80, 120),      # Mid-High Gamma
            (120, 180),     # High Gamma
            (180, 250),     # Ultra High Gamma I
            (250, 300),     # Ultra High Gamma II
        ]

        # Improved learning parameters for gamma prediction
        self.LEARNING_RATE = 2e-4
        self.WEIGHT_DECAY = 2e-5
        self.EPOCHS = 10
        self.PATIENCE = 15
        self.CLIP_GRAD_NORM = 0.8
        self.LR_WARMUP_EPOCHS = 5
        self.GRADIENT_ACCUMULATION_STEPS = 2

        # Enhanced Graph Reservoir parameters
        self.NUM_POPULATIONS = 8         # Increased for better specialization
        self.NEURONS_PER_POPULATION = 140 # More neurons for complex dynamics
        self.SPECTRAL_RADIUS = 0.94
        self.LEAKY_RATE = 0.75           # Higher leaky rate for faster dynamics
        self.INTER_POPULATION_DENSITY = 0.25
        self.INTRA_POPULATION_DENSITY = 0.45
        self.NOISE_LEVEL = 0.015

        # Specialized populations for gamma frequency bands
        self.POPULATION_SPECIALIZATIONS = {
            0: 'low_gamma',      # 30-50 Hz
            1: 'mid_low_gamma',  # 50-80 Hz
            2: 'mid_high_gamma', # 80-120 Hz
            3: 'high_gamma',     # 120-180 Hz
            4: 'ultra_gamma_1',  # 180-250 Hz
            5: 'ultra_gamma_2',  # 250-300 Hz
            6: 'integrator',     # Integration specialized
            7: 'context'         # Context maintenance
        }

        # Enhanced oscillator parameters
        self.USE_OSCILLATORS = True
        self.OSCILLATION_STRENGTH = 0.7  # Increased oscillation strength for better gamma capture
        self.OSCILLATOR_DAMPING = 0.05   # Lower damping for sustained oscillations
        self.ADAPTATION_RATE = 0.008     # Slower adaptation for stability
        self.OSCILLATORS_PER_BAND = 8    # More oscillators per band for better frequency coverage

        # Enhanced network architecture parameters
        self.HIDDEN_SIZE = 192           # Larger hidden representations
        self.READOUT_HIDDEN = 384        # Larger readout layer
        self.READOUT_LAYERS = 3          # More layers for better decoding
        self.DROPOUT = 0.25              # More regularization
        self.USE_ATTENTION = True
        self.ATTENTION_HEADS = 4         # More attention heads
        self.USE_SKIP_CONNECTIONS = True # Add skip connections
        self.USE_LAYER_NORM = True       # Use layer normalization
        self.USE_BATCH_NORM = True       # Use batch normalization for better training

        # Enhanced loss function parameters
        self.TIME_LOSS_WEIGHT = 0.25     # Less weight for time domain (overall shape)
        self.FREQ_LOSS_WEIGHT = 0.75     # More weight for frequency domain (oscillations)

        # Progressive gamma band weighting (heavily weight higher frequencies)
        self.GAMMA_BAND_WEIGHTS = {
            (30, 50): 1.0,      # Low gamma weight
            (50, 80): 1.5,      # Mid-Low gamma weight
            (80, 120): 2.0,     # Mid-High gamma weight
            (120, 180): 2.5,    # High gamma weight
            (180, 250): 3.0,    # Ultra high gamma I weight
            (250, 300): 3.5     # Ultra high gamma II weight
        }

        # Special weights for phase accuracy
        self.PHASE_LOSS_WEIGHT = 0.4     # Weight for phase prediction accuracy
        self.ENVELOPE_LOSS_WEIGHT = 0.2  # Weight for amplitude envelope
        self.USE_PHASE_CONSISTENCY_LOSS = True  # Use phase consistency loss

        # Special training strategies
        self.USE_SWA = True             # Stochastic Weight Averaging
        self.SWA_START = 20             # When to start SWA
        self.USE_LOOKAHEAD = False      # Disabled Lookahead optimizer due to compatibility issues
        self.LOOKAHEAD_K = 5            # Lookahead steps
        self.LOOKAHEAD_ALPHA = 0.5      # Lookahead alpha

        # Curriculum learning parameters
        self.USE_CURRICULUM = True          # Use curriculum learning
        self.CURRICULUM_START_EPOCH = 0     # Start with basic curriculum
        self.CURRICULUM_PHASE_LENGTH = 10   # Length of each curriculum phase

        # Advanced oscillator parameters
        self.USE_COUPLED_OSCILLATORS = True # Use coupled oscillators for better synchronization
        self.COUPLING_STRENGTH = 0.15       # Coupling strength between oscillators
        self.USE_RESONATOR_BANKS = True     # Use banks of resonators tuned to gamma frequencies
        self.RESONATOR_Q = 20               # Quality factor for resonators (higher = narrower bandwidth)

        # Tracking parameters
        self.TRACK_STATES = True
        self.STORE_TRAJECTORIES = True

        # Visualization parameters - added for improved aesthetics
        self.VIZ_COLORSCALES = {
            'trajectory': 'Plasma',
            'influence': 'Viridis',
            'electrodes': 'Turbo',
            'connections': 'Blues'
        }
        self.VIZ_BG_COLOR = '#f8f9fa'
        self.VIZ_FONT_FAMILY = 'Arial'
        self.VIZ_TITLE_SIZE = 24
        self.VIZ_AXIS_TITLE_SIZE = 14
        self.VIZ_TEXT_COLOR = '#333333'
        self.VIZ_GRID_COLOR = '#e0e0e0'
        self.VIZ_NODE_SIZE_RANGE = (10, 50)
        self.VIZ_LINE_WIDTH_RANGE = (1, 10)
        self.VIZ_OPACITY_RANGE = (0.3, 1.0)
        self.VIZ_MARKER_SYMBOLS = {
            'start': 'diamond',
            'end': 'square',
            'shift': 'star',
            'electrode': 'circle'
        }

        # Directory setup
        self.OUTPUT_DIR = "enhanced_gamma_esn_results"
        self.create_directories()

    def create_directories(self):
        """Create necessary directories for outputs."""
        os.makedirs(self.OUTPUT_DIR, exist_ok=True)
        for subdir in ['models', 'visualizations', 'logs', 'trajectories', 'analysis']:
            os.makedirs(os.path.join(self.OUTPUT_DIR, subdir), exist_ok=True)

# Initialize configuration
config = Config()


def find_data_file(path, filename):
    """Find the specified data file in the given path or its subdirectories."""
    data_file = os.path.join(path, filename)
    if os.path.exists(data_file):
        return data_file

    for root, dirs, files in os.walk(path):
        if filename in files:
            return os.path.join(root, filename)

    raise FileNotFoundError(f"Could not find {filename} in {path}")

def load_electrode_positions(path):
    """Load electrode position data from CSV file."""
    print("Loading electrode positions...")
    positions_file = find_data_file(path, 'probe_channels_810755799 (1).csv')

    # Updated column mapping for the new dataset
    positions = pd.read_csv(positions_file)

    # Rename columns to match the expected format
    positions = positions.rename(columns={
        'channel_index': 'channel_num',
        'probe_horizontal_position': 'x_position',
        'probe_vertical_position': 'y_position'
    })

    print(f"Loaded positions for {len(positions)} electrodes")
    return positions

def select_electrodes(positions, num_electrodes=None):
    """Select electrodes using KMeans clustering for optimal spatial coverage."""
    if num_electrodes is None:
        num_electrodes = config.NUM_ELECTRODES

    print(f"Selecting {num_electrodes} electrodes for analysis...")

    # Normalize positions for clustering
    x_norm = (positions['x_position'] - positions['x_position'].min()) / (positions['x_position'].max() - positions['x_position'].min())
    y_norm = (positions['y_position'] - positions['y_position'].min()) / (positions['y_position'].max() - positions['y_position'].min())
    coords = np.column_stack((x_norm.values, y_norm.values))

    # Use KMeans to select representative electrodes
    from sklearn.cluster import KMeans
    kmeans = KMeans(n_clusters=num_electrodes, random_state=SEED, n_init=10)
    kmeans.fit(coords)

    # Select the electrode closest to each cluster center
    selected_indices = []
    for cluster_idx in range(num_electrodes):
        cluster_points = np.where(kmeans.labels_ == cluster_idx)[0]
        center = kmeans.cluster_centers_[cluster_idx]
        distances = np.sqrt(np.sum((coords[cluster_points] - center)**2, axis=1))
        closest_idx = cluster_points[np.argmin(distances)]
        selected_indices.append(closest_idx)

    # Get channel numbers for selected electrodes
    channel_nums = positions.iloc[selected_indices]['channel_num'].astype(int).values

    # Print the first few selected electrodes for debugging
    print(f"First 5 selected indices: {selected_indices[:5]}")
    print(f"First 5 selected channel numbers: {channel_nums[:5]}")

    print(f"Selected {len(channel_nums)} electrodes with good spatial coverage")
    return channel_nums, selected_indices

def visualize_probe(positions, selected_indices=None, save_path=None):
    """Visualize electrode positions on the probe."""
    plt.figure(figsize=(10, 14))
    plt.scatter(positions['x_position'], positions['y_position'],
                c='gray', alpha=0.3, s=30, label='All Electrodes')

    if selected_indices is not None:
        selected_positions = positions.iloc[selected_indices]
        plt.scatter(selected_positions['x_position'], selected_positions['y_position'],
                   c='red', s=50, label='Selected Electrodes')

        for i, (_, row) in enumerate(selected_positions.iterrows()):
            # FIXED: Use 'channel_num' instead of 'channel_index'
            plt.text(row['x_position'], row['y_position'], str(int(row['channel_num'])),
                     fontsize=8, ha='center', va='center')

    plt.xlabel('Horizontal Position (μm)')
    plt.ylabel('Vertical Position (μm)')
    plt.title('Electrode Positions on Probe')
    plt.legend()
    plt.grid(alpha=0.3)

    if save_path:
        plt.savefig(save_path, dpi=150, bbox_inches='tight')
        plt.close()
    else:
        plt.show()

def load_lfp_data(path, channel_nums, max_rows=500000):
    """Load LFP data from CSV file."""
    print(f"Loading LFP data for {len(channel_nums)} channels...")
    data_file = find_data_file(path, 'drifting_gratings_full (1).csv')

    # Include presentation_id in the columns to load
    cols_to_use = ['timestamp', 'presentation_id'] + [f'channel_{ch}' for ch in channel_nums]

    print(f"Columns to load (first few): {cols_to_use[:min(5, len(cols_to_use))]}... (and {len(cols_to_use)-5} more)")

    # Load data in chunks to handle large files
    data = pd.read_csv(data_file, usecols=cols_to_use, nrows=max_rows)
    print(f"Loaded {len(data)} rows of LFP data")

    # Print columns to verify presentation_id is included
    print(f"Columns in loaded data: {data.columns.tolist()}")

    # Print some information about presentations
    num_presentations = data['presentation_id'].nunique()
    print(f"Data contains {num_presentations} unique presentation_ids")

    return data

def extract_windows(data, context_size=None, predict_size=None, stride=None, max_windows=None):
    """
    Extract time windows from continuous data for training and prediction,
    ensuring each window comes from a single presentation_id if available.
    """
    if context_size is None:
        context_size = config.CONTEXT_SIZE
    if predict_size is None:
        predict_size = config.PREDICT_SIZE
    if stride is None:
        stride = config.STRIDE
    if max_windows is None:
        max_windows = config.MAX_SEQUENCES

    print(f"Extracting windows with context={context_size}, predict={predict_size}, stride={stride}")

    # Check if presentation_id is available
    use_presentation_id = 'presentation_id' in data.columns
    if use_presentation_id:
        print(f"Using presentation_id to ensure windows come from same presentation")
        print(f"Found {data['presentation_id'].nunique()} unique presentation_ids")
    else:
        print(f"Warning: presentation_id not found in data, windows may cross presentation boundaries")

    # Get channel columns (all columns except timestamp and presentation_id)
    data_cols = [col for col in data.columns if col.startswith('channel_')]

    # Calculate window size
    window_size = context_size + predict_size

    # Extract windows
    windows = []

    if use_presentation_id:
        # Group data by presentation_id
        presentation_groups = data.groupby('presentation_id')

        for presentation_id, group in presentation_groups:
            # Skip presentations that are too short
            if len(group) < window_size:
                continue

            # Get data values for this presentation
            presentation_data = group[data_cols].values

            # Extract windows from this presentation
            for start in range(0, len(presentation_data) - window_size + 1, stride):
                if len(windows) >= max_windows:
                    break

                window = presentation_data[start:start + window_size]
                if np.isnan(window).any():
                    continue

                windows.append(window)

                if len(windows) % 100 == 0:
                    print(f"Extracted {len(windows)} windows...")

            if len(windows) >= max_windows:
                break
    else:
        # Original window extraction logic (not grouping by presentation_id)
        data_values = data[data_cols].values

        for start in range(0, len(data_values) - window_size, stride):
            if len(windows) >= max_windows:
                break

            window = data_values[start:start + window_size]
            if np.isnan(window).any():
                continue

            windows.append(window)

            if len(windows) % 100 == 0:
                print(f"Extracted {len(windows)} windows...")

    if not windows:
        print("No valid windows extracted!")
        return np.array([])

    windows = np.array(windows)

    print(f"Extracted {len(windows)} windows with shape: {windows.shape}")

    return windows

def create_enhanced_filter_bank(fs, freq_bands, order=8):
    """Create enhanced filter bank with higher order filters."""
    filter_bank = []
    nyquist = fs / 2.0

    for band_idx, (low_freq, high_freq) in enumerate(freq_bands):
        # Normalize frequencies
        low = low_freq / nyquist
        high = high_freq / nyquist

        # Create higher order Butterworth filter for sharper cutoffs
        # Using sos (second-order sections) format for better numerical stability
        sos = signal.butter(order, [low, high], btype='bandpass', output='sos')
        filter_bank.append((sos, band_idx, (low_freq, high_freq)))

    return filter_bank

def extract_enhanced_frequency_bands(windows, fs=None, freq_bands=None):
    """Extract frequency bands using enhanced filters with parallel processing."""
    if fs is None:
        fs = config.SAMPLING_RATE
    if freq_bands is None:
        freq_bands = config.FREQ_BANDS

    print("Extracting enhanced frequency bands...")
    n_samples, window_size, n_channels = windows.shape
    n_bands = len(freq_bands)

    # Create filter bank
    filter_bank = create_enhanced_filter_bank(fs, freq_bands, order=8)

    # Initialize output array for filtered data
    filtered_data = np.zeros((n_samples, n_bands, window_size, n_channels))

    # Process each band with enhanced filters
    for (sos, band_idx, (low_freq, high_freq)) in filter_bank:
        print(f"Filtering band {band_idx+1}/{n_bands}: {low_freq}-{high_freq} Hz")

        # Process each sample and channel
        for i in tqdm(range(n_samples), desc=f"Band {band_idx+1}"):
            for j in range(n_channels):
                signal_i = windows[i, :, j]

                # Apply zero-phase filtering with second-order sections (more stable)
                filtered = signal.sosfiltfilt(sos, signal_i)

                # Store filtered signal
                filtered_data[i, band_idx, :, j] = filtered

    print(f"Filtered data shape: {filtered_data.shape}")
    return filtered_data

def enhanced_hilbert_transform(signal_data, pad_factor=2):
    """Apply enhanced Hilbert transform with better padding to reduce edge effects."""
    # Determine padding size (minimum 200 points or half signal length)
    n = len(signal_data)
    pad_size = max(int(n * pad_factor), 200)

    # Apply reflection padding (better than zero padding for reducing edge effects)
    padded_signal = np.pad(signal_data, (pad_size, pad_size), mode='reflect')

    # Apply Hilbert transform to get analytic signal
    analytic_signal = hilbert(padded_signal)

    # Extract phase and amplitude from the central (non-padded) portion
    phase = np.angle(analytic_signal[pad_size:pad_size+n])
    amplitude = np.abs(analytic_signal[pad_size:pad_size+n])

    # Get instantaneous frequency (derivative of phase)
    unwrapped_phase = np.unwrap(phase)
    inst_freq = np.diff(unwrapped_phase) / (2.0 * np.pi) * config.SAMPLING_RATE
    # Pad the instantaneous frequency to match original length
    inst_freq = np.append(inst_freq, inst_freq[-1])

    return phase, amplitude, inst_freq

def extract_phase_amplitude(filtered_data):
    """Extract phase, amplitude and instantaneous frequency using enhanced Hilbert transform."""
    print("Extracting phase, amplitude and instantaneous frequency information...")
    n_samples, n_bands, n_timepoints, n_channels = filtered_data.shape

    # Initialize arrays for phase, amplitude and frequency
    phase = np.zeros_like(filtered_data)
    amplitude = np.zeros_like(filtered_data)
    inst_freq = np.zeros_like(filtered_data)

    for i in tqdm(range(n_samples), desc="Processing samples"):
        for b in range(n_bands):
            for c in range(n_channels):
                try:
                    # Apply enhanced Hilbert transform
                    p, a, f = enhanced_hilbert_transform(filtered_data[i, b, :, c])

                    # Store results
                    phase[i, b, :, c] = p
                    amplitude[i, b, :, c] = a
                    inst_freq[i, b, :, c] = f
                except Exception as e:
                    print(f"Error in Hilbert transform: {e}")
                    # Fill with zeros on error
                    phase[i, b, :, c] = 0
                    amplitude[i, b, :, c] = 0
                    inst_freq[i, b, :, c] = 0

    # Normalize amplitude (per band) for better stability using 99th percentile
    for b in range(n_bands):
        max_amp = np.percentile(amplitude[:, b, :, :], 99)
        if max_amp > 0:
            amplitude[:, b, :, :] /= max_amp

    return phase, amplitude, inst_freq

def compute_spectral_features(data, fs=None, nperseg=64, noverlap=32):
    """Compute time-frequency features using spectrogram instead of wavelets."""
    if fs is None:
        fs = config.SAMPLING_RATE

    n_samples, n_timepoints, n_channels = data.shape

    # Compute spectrogram parameters
    f, _, _ = signal.spectrogram(np.zeros(nperseg), fs=fs, nperseg=nperseg, noverlap=noverlap)
    n_freq_bins = len(f)
    n_time_bins = 1 + (n_timepoints - nperseg) // (nperseg - noverlap)

    # Initialize output arrays
    spec_power = np.zeros((n_samples, n_freq_bins, n_time_bins, n_channels))

    # Compute spectrogram for each sample and channel
    print("Computing spectral features...")
    for i in tqdm(range(n_samples), desc="Spectral analysis"):
        for c in range(n_channels):
            signal_i = data[i, :, c]

            # Compute spectrogram
            f, t, Sxx = signal.spectrogram(
                signal_i,
                fs=fs,
                nperseg=nperseg,
                noverlap=noverlap,
                scaling='density'
            )

            # Store power
            spec_power[i, :, :Sxx.shape[1], c] = Sxx

    # Focus on gamma frequencies (>30 Hz)
    gamma_mask = f >= 30
    spec_power_gamma = spec_power[:, gamma_mask, :, :]
    freq_bins_gamma = f[gamma_mask]

    # Normalize spectrogram power
    for f_idx in range(len(freq_bins_gamma)): # Iterate using index
        max_power = np.percentile(spec_power_gamma[:, f_idx, :, :], 99)
        if max_power > 0:
            spec_power_gamma[:, f_idx, :, :] /= max_power

    return spec_power_gamma, freq_bins_gamma

def compute_power_features(amplitude_data, window_size=25):
    """Compute enhanced power-related features from amplitude data."""
    # Get dimensions
    n_samples, n_bands, n_timepoints, n_channels = amplitude_data.shape

    # Compute power (square of amplitude)
    power_data = amplitude_data ** 2

    # Use a more appropriate window size based on signal length
    window_size = min(window_size, n_timepoints // 5)
    stride = max(1, window_size // 2)  # 50% overlap

    # Number of windows
    n_windows = (n_timepoints - window_size) // stride + 1

    # Initialize power features array
    power_features = np.zeros((n_samples, n_bands, n_windows, n_channels))

    # Compute power in each window
    for i in range(n_samples):
        for b in range(n_bands):
            for c in range(n_channels):
                for w in range(n_windows):
                    start_idx = w * stride
                    end_idx = start_idx + window_size
                    power_features[i, b, w, c] = np.mean(power_data[i, b, start_idx:end_idx, c])

    return power_features

def compute_phase_coherence(phase_data):
    """Compute phase coherence between electrodes for each frequency band."""
    n_samples, n_bands, n_timepoints, n_channels = phase_data.shape

    # Initialize coherence matrix: [samples, bands, channels, channels]
    coherence = np.zeros((n_samples, n_bands, n_channels, n_channels))

    print("Computing phase coherence...")
    for i in tqdm(range(n_samples), desc="Phase coherence"):
        for b in range(n_bands):
            for c1 in range(n_channels):
                phase1 = phase_data[i, b, :, c1]
                for c2 in range(c1+1, n_channels):
                    phase2 = phase_data[i, b, :, c2]

                    # Compute phase coherence
                    # Formula: |mean(exp(1j*(phase1-phase2)))|
                    phase_diff = phase1 - phase2
                    complex_exp = np.exp(1j * phase_diff)
                    coh = np.abs(np.mean(complex_exp))

                    # Store in symmetric matrix
                    coherence[i, b, c1, c2] = coh
                    coherence[i, b, c2, c1] = coh

    return coherence

def compute_phase_locking_value(phase_data):
    """Compute Phase Locking Value (PLV) over time for each frequency band and electrode."""
    n_samples, n_bands, n_timepoints, n_channels = phase_data.shape

    # Window size for PLV computation (temporal smoothing)
    window_size = min(50, n_timepoints // 4)

    # Initialize PLV array: [samples, bands, timepoints-window+1, channels]
    n_windows = n_timepoints - window_size + 1
    plv = np.zeros((n_samples, n_bands, n_windows, n_channels))

    print("Computing Phase Locking Value...")
    for i in tqdm(range(n_samples), desc="PLV computation"):
        for b in range(n_bands):
            for c in range(n_channels):
                phase = phase_data[i, b, :, c]

                # Compute PLV in sliding windows
                for t in range(n_windows):
                    window_phase = phase[t:t+window_size]
                    complex_exp = np.exp(1j * window_phase)
                    plv[i, b, t, c] = np.abs(np.mean(complex_exp))

    return plv

def preprocess_data(windows, context_size=None, predict_size=None):
    """Enhanced preprocessing with more sophisticated spectral features for gamma focus."""
    if context_size is None:
        context_size = config.CONTEXT_SIZE
    if predict_size is None:
        predict_size = config.PREDICT_SIZE

    print("Preprocessing data with enhanced gamma focus...")
    num_windows, window_size, num_electrodes = windows.shape
    assert window_size == context_size + predict_size

    # Split windows into context and target parts
    context_windows = windows[:, :context_size, :]
    target_windows = windows[:, context_size:, :]

   # Adjust the scaler to handle the very small amplitude ranges
    scaler = RobustScaler(quantile_range=(1, 99))

    # You may also want to add additional scaling for the very small amplitudes
    # For example, before scaling, you could multiply the values by 1e6 to make them more numerically stable
    context_flat = context_windows.reshape(-1, num_electrodes) * 1e6
    target_flat = target_windows.reshape(-1, num_electrodes) * 1e6

    # Then continue with the scaling
    scaler.fit(context_flat)
    context_scaled = scaler.transform(context_flat).reshape(context_windows.shape)
    target_scaled = scaler.transform(target_flat).reshape(target_windows.shape)

    # Similarly for full_flat
    full_flat = windows.reshape(-1, num_electrodes) * 1e6
    full_scaled = scaler.transform(full_flat).reshape(windows.shape)

    # Extract frequency band information with enhanced filters
    filtered_windows = extract_enhanced_frequency_bands(full_scaled, config.SAMPLING_RATE, config.FREQ_BANDS)

    # Extract phase, amplitude, and instantaneous frequency information
    phase_data, amplitude_data, inst_freq_data = extract_phase_amplitude(filtered_windows)

    # Compute spectral features
    spectral_power, freq_bins = compute_spectral_features(full_scaled, config.SAMPLING_RATE)

    # Compute power features for enhanced spectral representation
    power_features = compute_power_features(amplitude_data)

    # Compute phase coherence between electrodes
    phase_coherence = compute_phase_coherence(phase_data)

    # Compute phase locking value
    plv = compute_phase_locking_value(phase_data)

    # Split features into context and target parts
    phase_context = phase_data[:, :, :context_size, :]
    amplitude_context = amplitude_data[:, :, :context_size, :]
    freq_context = inst_freq_data[:, :, :context_size, :]
    filtered_context = filtered_windows[:, :, :context_size, :]

    # Calculate number of time bins in spectral data
    # This depends on the spectrogram parameters (nperseg, noverlap)
    nperseg = 64
    noverlap = 32
    n_time_bins_context = 1 + (context_size - nperseg) // (nperseg - noverlap)
    spectral_context = spectral_power[:, :, :n_time_bins_context, :]


    # Calculate number of windows for downsampled features (power_features)
    # Based on window size (25) and stride (12) in power_features calculation
    power_feature_window_size = 25 # from compute_power_features default
    power_feature_stride = max(1, power_feature_window_size // 2)
    n_windows_context_power = (context_size - power_feature_window_size) // power_feature_stride + 1


    # Create dataset with all features
    X_data = {
        'raw': context_scaled,
        'phase': phase_context,
        'amplitude': amplitude_context,
        'inst_freq': freq_context,
        'filtered': filtered_context,
        'power': power_features[:, :, :n_windows_context_power, :], # Use correct slicing
        'spectral': spectral_context,
        'coherence': phase_coherence # Coherence is static for the context window
    }

    y_data = {
        'target': target_scaled,
        'phase_target': phase_data[:, :, context_size:, :],
        'amplitude_target': amplitude_data[:, :, context_size:, :],
        'freq_target': inst_freq_data[:, :, context_size:, :]
    }

    # Split into training, validation, and test sets (70/15/15 split)
    from sklearn.model_selection import train_test_split

    # Create indices for splitting
    indices = np.arange(len(context_scaled))
    train_idx, temp_idx = train_test_split(indices, test_size=0.3, random_state=SEED)
    val_idx, test_idx = train_test_split(temp_idx, test_size=0.5, random_state=SEED)

    # Create PyTorch datasets
    train_dataset = EnhancedLFPDataset(
        {k: v[train_idx] for k, v in X_data.items()},
        {k: v[train_idx] for k, v in y_data.items()}
    )

    val_dataset = EnhancedLFPDataset(
        {k: v[val_idx] for k, v in X_data.items()},
        {k: v[val_idx] for k, v in y_data.items()}
    )

    test_dataset = EnhancedLFPDataset(
        {k: v[test_idx] for k, v in X_data.items()},
        {k: v[test_idx] for k, v in y_data.items()}
    )

    # Create data loaders
    train_loader = DataLoader(train_dataset, batch_size=config.BATCH_SIZE, shuffle=True)
    val_loader = DataLoader(val_dataset, batch_size=config.BATCH_SIZE)
    test_loader = DataLoader(test_dataset, batch_size=config.BATCH_SIZE)

    print(f"Created data loaders with {len(train_loader)} training batches, "
          f"{len(val_loader)} validation batches, and {len(test_loader)} test batches")

    return train_loader, val_loader, test_loader, scaler

class EnhancedLFPDataset(Dataset):
    """Enhanced dataset for LFP data with gamma-specific spectral decomposition."""

    def __init__(self, X, y):
        """Initialize dataset with extended input features and target."""
        self.X_raw = torch.FloatTensor(X['raw'])
        self.X_phase = torch.FloatTensor(X['phase'])
        self.X_amplitude = torch.FloatTensor(X['amplitude'])
        self.X_inst_freq = torch.FloatTensor(X['inst_freq'])
        self.X_filtered = torch.FloatTensor(X['filtered'])
        self.X_power = torch.FloatTensor(X['power'])
        self.X_spectral = torch.FloatTensor(X['spectral'])
        self.X_coherence = torch.FloatTensor(X['coherence'])

        self.y_target = torch.FloatTensor(y['target'])
        self.y_phase = torch.FloatTensor(y['phase_target'])
        self.y_amplitude = torch.FloatTensor(y['amplitude_target'])
        self.y_freq = torch.FloatTensor(y['freq_target'])

    def __len__(self):
        return len(self.y_target)

    def __getitem__(self, idx):
        return {
            'raw': self.X_raw[idx],
            'phase': self.X_phase[idx],
            'amplitude': self.X_amplitude[idx],
            'inst_freq': self.X_inst_freq[idx],
            'filtered': self.X_filtered[idx],
            'power': self.X_power[idx],
            'spectral': self.X_spectral[idx],
            'coherence': self.X_coherence[idx]
        }, {
            'target': self.y_target[idx],
            'phase': self.y_phase[idx],
            'amplitude': self.y_amplitude[idx],
            'freq': self.y_freq[idx]
        }


class ResonatorOscillator(nn.Module):
    """High-Q resonator oscillator specifically designed for gamma frequencies."""
    def __init__(self, center_freq, bandwidth=10.0, amplitude=1.0):
        super().__init__()
        self.center_freq = center_freq  # Center frequency in Hz
        self.bandwidth = bandwidth      # Bandwidth in Hz (narrower = higher Q)

        # Learnable parameters with better initialization
        self.amplitude = nn.Parameter(torch.ones(1) * amplitude)
        self.phase_shift = nn.Parameter(torch.rand(1) * 2 * np.pi)
        self.damping = nn.Parameter(torch.tensor([0.05]))  # Lower initial value for less damping

        # Convert to resonator parameters - ensure these are tensors
        self.omega_0 = 2 * np.pi * center_freq / 1000  # Angular frequency in rad/timestep
        self.q_factor = torch.tensor(center_freq / bandwidth, dtype=torch.float32)  # Quality factor as tensor

        # State variables
        self.register_buffer('state', None, persistent=False)
        self.register_buffer('time_step', torch.tensor(0), persistent=False)

    def reset_state(self, batch_size=1):
        """Reset the resonator state."""
        # Use two state variables: position and velocity
        self.state = torch.zeros(batch_size, 2, device=self.phase_shift.device)  # [position, velocity]
        self.time_step = torch.tensor(0) # Reset time_step as well

    def forward(self, x):
        """Forward pass with enhanced resonator dynamics."""
        batch_size = x.shape[0]

        # Initialize state if needed
        if self.state is None or self.state.shape[0] != batch_size:
            self.reset_state(batch_size)

        # Increment time step
        self.time_step += 1

        # Extract current states
        pos = self.state[:, 0]
        vel = self.state[:, 1]

        # Compute damping coefficient from Q factor - fixed to avoid clamp on numpy value
        q_factor_tensor = self.q_factor.to(self.phase_shift.device)
        min_q = torch.tensor(0.5, device=self.phase_shift.device)
        safe_q = torch.max(q_factor_tensor, min_q)
        # Ensure damping is a scalar or compatible with batch operations
        # self.omega_0 is a Python float, safe_q is a scalar tensor
        # Damping calculation should result in a scalar tensor compatible with broadcasting
        damping_coeff = self.omega_0 / (2.0 * safe_q)


        # Input force with phase shift modulation
        phase_mod = 0.1 * torch.sin(self.omega_0 * self.time_step.float() * 0.1 + self.phase_shift) # time_step to float
        input_force = x.view(batch_size) * (1.0 + phase_mod)

        # Update velocity using second-order resonator equation:
        # ẍ + (ω₀/Q)ẋ + ω₀²x = F(t)
        dt = 1.0  # Time step
        # Ensure all terms are broadcastable with [batch_size]
        accel = input_force - 2 * damping_coeff * vel - self.omega_0**2 * pos
        vel_new = vel + accel * dt

        # Update position
        pos_new = pos + vel_new * dt

        # Add direct oscillatory component to ensure frequency stability
        # This helps maintain the gamma oscillation pattern even with input variations
        t = self.time_step.float() * dt # time_step to float
        direct_oscillation = self.amplitude * torch.sin(self.omega_0 * t + self.phase_shift)
        direct_oscillation = direct_oscillation.repeat(batch_size) # Ensure [batch_size]

        # Mix resonator and direct oscillation (80/20 blend for stability)
        pos_new = 0.8 * pos_new + 0.2 * direct_oscillation

        # Apply soft limiting to prevent instability
        pos_new = torch.tanh(pos_new)
        # Use torch.clamp properly with tensor inputs
        vel_new = torch.clamp(vel_new, min=-5.0, max=5.0)

        # Update state
        self.state = torch.stack([pos_new, vel_new], dim=1)

        return pos_new

class CoupledResonatorBank(nn.Module):
    """Bank of coupled resonators for realistic gamma generation."""
    def __init__(self, freq_band, num_resonators=8, coupling_strength=0.15):
        super().__init__()
        self.freq_band = freq_band  # (low_freq, high_freq)
        self.num_resonators = num_resonators
        self.coupling_strength = coupling_strength

        # Create resonators with frequencies distributed across the band
        self.resonators = nn.ModuleList()
        freqs = np.linspace(freq_band[0], freq_band[1], num_resonators)

        # Create resonators with varying Q factors
        # Higher frequencies get higher Q factors for better selectivity
        q_factors = np.linspace(10, 25, num_resonators)
        amplitudes = np.linspace(0.7, 1.0, num_resonators)

        for i in range(num_resonators):
            self.resonators.append(
                ResonatorOscillator(
                    center_freq=freqs[i],
                    bandwidth=freqs[i]/q_factors[i],
                    amplitude=amplitudes[i]
                )
            )

        # Create coupling matrix between resonators
        # Stronger coupling between neighboring frequencies
        coupling_matrix = np.zeros((num_resonators, num_resonators))
        for i in range(num_resonators):
            for j in range(num_resonators):
                if i != j:
                    # Coupling strength decreases with frequency distance
                    freq_diff = abs(freqs[i] - freqs[j])
                    max_diff = freq_band[1] - freq_band[0]
                    if max_diff == 0: # Avoid division by zero if band is a single frequency
                        coupling_matrix[i,j] = coupling_strength if freq_diff == 0 else 0
                    else:
                        coupling_matrix[i, j] = coupling_strength * (1.0 - freq_diff/max_diff)


        self.register_buffer('coupling', torch.FloatTensor(coupling_matrix))

    def reset_state(self, batch_size=1):
        """Reset all resonators."""
        for resonator in self.resonators:
            resonator.reset_state(batch_size)

    def forward(self, x): # x is [batch_size, 1] from GammaCellUnit, or [batch_size] from GammaSpecificDecoder
        if x.ndim == 2 and x.shape[1] == 1:
            x_squeezed = x.squeeze(1) # Make it [batch_size]
        else:
            x_squeezed = x # Assume it's already [batch_size]

        batch_size = x_squeezed.shape[0]


        outputs = []
        # ResonatorOscillator expects [batch_size] input
        for resonator in self.resonators:
            outputs.append(resonator(x_squeezed)) # Pass [batch_size]

        coupled_outputs = []
        for i, output_i in enumerate(outputs): # output_i is [batch_size]
            coupled_i = output_i.clone() # Use a different variable name
            for j, output_j in enumerate(outputs): # output_j is [batch_size]
                if i != j:
                    # self.coupling[i,j] is scalar, output_j is [batch_size]
                    coupled_i = coupled_i + self.coupling[i, j] * output_j
            coupled_outputs.append(coupled_i)


        final_outputs = []
        for i, resonator in enumerate(self.resonators):
            # coupled_outputs[i] is [batch_size]
            # x_squeezed is [batch_size]
            coupled_input_i = 0.7 * x_squeezed + 0.3 * coupled_outputs[i] # All [batch_size]
            final_outputs.append(resonator(coupled_input_i))


        weights = torch.linspace(0.5, 1.5, self.num_resonators, device=x.device)
        # final_outputs is list of [batch_size] tensors.
        # Stack them to [batch_size, num_resonators]
        stacked_final_outputs = torch.stack(final_outputs, dim=1)

        # weights is [num_resonators]. Need to broadcast for element-wise product.
        # [batch_size, num_resonators] * [1, num_resonators]
        weighted_outputs = stacked_final_outputs * weights.unsqueeze(0)


        return torch.sum(weighted_outputs, dim=1) / weights.sum()


class MultiHeadSelfAttention(nn.Module):
    """Enhanced multi-head self-attention mechanism with improved stability."""
    def __init__(self, input_dim, num_heads=4, dropout=0.1):
        super().__init__()
        self.num_heads = num_heads
        self.head_dim = input_dim // num_heads
        assert self.head_dim * num_heads == input_dim, "input_dim must be divisible by num_heads"

        # Improved projections with layer normalization
        self.layer_norm = nn.LayerNorm(input_dim)
        self.qkv_proj = nn.Linear(input_dim, 3 * input_dim)
        self.out_proj = nn.Linear(input_dim, input_dim)
        self.dropout = nn.Dropout(dropout)

        # Scaling factor
        self.register_buffer('scale', torch.sqrt(torch.FloatTensor([self.head_dim])))

    def forward(self, x):
        """
        Forward pass with improved numerical stability.
        x: [batch_size, seq_len, input_dim]
        """
        batch_size = x.shape[0]
        seq_len = x.shape[1]

        # Apply layer normalization first for stability
        x_norm = self.layer_norm(x) # Apply to x, not overwrite x yet for residual

        # Combined projection
        qkv = self.qkv_proj(x_norm) # Use normalized x for QKV projection
        qkv = qkv.reshape(batch_size, seq_len, 3, self.num_heads, self.head_dim)
        qkv = qkv.permute(2, 0, 3, 1, 4)  # [3, batch, heads, seq_len, head_dim]

        # Split QKV
        q, k, v = qkv[0], qkv[1], qkv[2]

        # Calculate attention scores with improved numerical stability
        energy = torch.matmul(q, k.transpose(-2, -1)) / self.scale

        # Apply softmax with better numerical stability
        attention = F.softmax(energy, dim=-1)
        attention = self.dropout(attention)

        # Apply attention to values
        attended_values = torch.matmul(attention, v) # Renamed from x to avoid confusion
        attended_values = attended_values.permute(0, 2, 1, 3).contiguous()
        attended_values = attended_values.reshape(batch_size, seq_len, -1)

        # Output projection
        output = self.out_proj(attended_values) # Renamed from x

        return output, attention # Return output (projected attended values)

class PopulationAttention(nn.Module):
    """Enhanced population-level attention for information exchange."""
    def __init__(self, input_dim, num_heads=4):
        super().__init__()
        self.attention = MultiHeadSelfAttention(input_dim, num_heads)
        self.norm = nn.LayerNorm(input_dim) # This norm is for after residual + attention

        # Add a gating mechanism for controlled information flow
        self.gate = nn.Sequential(
            nn.Linear(input_dim, input_dim), # Gate calculated from original x
            nn.Sigmoid()
        )

    def forward(self, x):
        """Enhanced forward pass with gating.
        x: [batch_size, num_populations, input_dim]
        """
        residual = x
        # Attention module already includes a pre-norm for QKV calculation
        # The MultiHeadSelfAttention returns the *output projection* of attended values
        attended_values, attention_weights = self.attention(x) # No need to norm x before this

        # Calculate gate values (how much of the attention to use) based on original x
        gate_values = self.gate(residual) # Or self.gate(x) which is the same

        # Apply gated residual connection
        # output = residual + gate_values * attended # This is one way
        # Another common way: output = (1-gate_values)*residual + gate_values*attended
        # Or output = self.norm(residual + gate_values * attended_values)
        # Current: output = residual + gate_values * attended_values
        # Let's follow the structure: Attention -> Gated Sum -> Norm

        output = residual + gate_values * attended_values

        # Final layer normalization
        output = self.norm(output)

        return output, attention_weights


class GammaCellUnit(nn.Module):
    """Specialized cell unit for gamma frequency dynamics."""
    def __init__(self, input_size, hidden_size, band_freq=(30, 300)):
        super().__init__()
        self.input_size = input_size
        self.hidden_size = hidden_size
        self.band_freq = band_freq  # Gamma band to specialize in

        # Input projection
        self.input_proj = nn.Linear(input_size, hidden_size)

        # Recurrent weights
        self.recurrent_proj = nn.Linear(hidden_size, hidden_size)

        # Output gate
        self.output_gate = nn.Sequential(
            nn.Linear(hidden_size + input_size, hidden_size), # Takes concatenated (current_hidden, current_input)
            nn.Sigmoid()
        )

        # Frequency modulation - controls oscillation frequency
        self.freq_mod = nn.Linear(input_size, 1) # Takes current_input

        # Decay factor - controls persistence of oscillations
        self.decay_factor = nn.Parameter(torch.tensor(0.1))

        # Excitatory-inhibitory balance
        self.ei_balance = nn.Parameter(torch.tensor(0.7))  # 0.7 = 70% excitatory

        # Layer normalization for stability of hidden state update
        self.layer_norm_hidden = nn.LayerNorm(hidden_size)


        # Initialize resonator bank for this frequency band
        center_freq = (band_freq[0] + band_freq[1]) / 2
        self.resonator = CoupledResonatorBank(
            freq_band=band_freq,
            num_resonators=config.OSCILLATORS_PER_BAND, # Use config
            coupling_strength=config.COUPLING_STRENGTH  # Use config
        )


        # Initialize hidden state
        self.register_buffer('hidden_state', None, persistent=False)

    def reset_state(self, batch_size=1):
        """Reset cell state."""
        self.hidden_state = torch.zeros(batch_size, self.hidden_size, device=self.decay_factor.device)
        self.resonator.reset_state(batch_size)

    def forward(self, x): # x is current input to the cell [batch_size, self.input_size]
        """Forward pass incorporating gamma oscillatory dynamics."""
        batch_size = x.shape[0]

        # Initialize state if needed
        if self.hidden_state is None or self.hidden_state.shape[0] != batch_size:
            self.reset_state(batch_size)

        # Input projection from current input x
        input_proj = self.input_proj(x) # [batch_size, self.hidden_size]

        # Calculate frequency modulation from current input x
        freq_mod = torch.sigmoid(self.freq_mod(x)) # [batch_size, 1]

        # Calculate recurrent contribution from previous hidden state
        recurrent_contrib = self.recurrent_proj(self.hidden_state) # [batch_size, self.hidden_size]

        # Apply E/I balance
        excitatory = F.relu(recurrent_contrib)
        inhibitory = F.relu(-recurrent_contrib)
        balanced_recurrent = self.ei_balance * excitatory - (1 - self.ei_balance) * inhibitory

        # Combine input and recurrent for main pathway
        combined_main = input_proj + balanced_recurrent # [batch_size, self.hidden_size]

        # Apply resonator dynamics to add oscillatory component
        # Resonator input is mean of combined_main across hidden_size dimension
        resonator_input = torch.mean(combined_main, dim=1, keepdim=True) # [batch_size, 1]
        oscillation_scalar = self.resonator(resonator_input) # [batch_size]

        # Expand scalar oscillation to match hidden_size, modulated by freq_mod
        # oscillation_scalar is [B], freq_mod is [B,1], self.hidden_size
        # gamma_contrib should be [B, self.hidden_size]
        # (freq_mod: [B,1]) * (oscillation_scalar.unsqueeze(1).expand(-1, self.hidden_size): [B, H])
        gamma_contrib = freq_mod * oscillation_scalar.unsqueeze(1).expand(-1, self.hidden_size)


        # Update hidden state with decay
        decay = torch.sigmoid(self.decay_factor) # scalar
        # New hidden state based on main pathway and gamma contribution
        updated_hidden_candidate = combined_main + gamma_contrib

        # Apply layer normalization to the candidate hidden state
        normalized_hidden_candidate = self.layer_norm_hidden(updated_hidden_candidate)

        self.hidden_state = (1 - decay) * self.hidden_state + decay * normalized_hidden_candidate


        # Calculate output gate based on current input x and new hidden_state
        output_gate_input = torch.cat([x, self.hidden_state], dim=1) # [B, input_size + hidden_size]
        gate_signal = self.output_gate(output_gate_input) # [B, hidden_size]

        # Apply output gate to the new hidden state
        gated_output = gate_signal * self.hidden_state

        return gated_output


class EnhancedGraphReservoirCell(nn.Module):
    """Enhanced graph-based reservoir cell with multiple populations specialized for gamma."""
    def __init__(self, config):
        super().__init__()
        self.config = config

        # Dimensions
        self.num_populations = config.NUM_POPULATIONS
        self.neurons_per_pop = config.NEURONS_PER_POPULATION
        self.total_neurons = self.num_populations * self.neurons_per_pop
        self.input_size = config.HIDDEN_SIZE

        # Create adjacency matrix for reservoir connections
        adjacency = self._initialize_graph_structure()
        self.adjacency_matrix = nn.Parameter(adjacency, requires_grad=True) # Made learnable

        # Input projections for each population
        self.input_projections = nn.ModuleList([
            nn.Linear(self.input_size, self.neurons_per_pop)
            for _ in range(self.num_populations)
        ])

        # Population biases
        self.population_biases = nn.ParameterList([
            nn.Parameter(torch.zeros(self.neurons_per_pop))
            for _ in range(self.num_populations)
        ])

        # Population time constants (leaky rates)
        base_tc = torch.ones(self.num_populations) * config.LEAKY_RATE
        for i, spec_tuple in enumerate(config.POPULATION_SPECIALIZATIONS.items()):
            pop_idx_cfg, spec = spec_tuple # specialized population index might not match loop index i
            # Find correct index i in base_tc that corresponds to pop_idx_cfg
            # Assuming POPULATION_SPECIALIZATIONS keys are 0 to NUM_POPULATIONS-1 and match loop i
            factor = 1.0
            if 'gamma' in spec:
                if 'low' in spec: factor = 0.85
                elif 'mid_low' in spec: factor = 0.95
                elif 'mid_high' in spec: factor = 1.05
                elif 'high' in spec: factor = 1.15
                elif 'ultra' in spec: factor = 1.25
            elif spec == 'integrator': factor = 0.7
            elif spec == 'context': factor = 0.5
            # Ensure pop_idx_cfg is within bounds of base_tc if it's used as index
            if pop_idx_cfg < self.num_populations:
                 base_tc[pop_idx_cfg] = config.LEAKY_RATE * factor


        self.time_constants = nn.Parameter(base_tc, requires_grad=True)

        # Gamma cell units for oscillatory populations
        if config.USE_OSCILLATORS:
            self.gamma_cells = nn.ModuleList()
            # Ensure gamma_cells list matches num_populations, with Identity for non-gamma
            gamma_cell_config_map = {}
            for pop_idx_cfg, spec in config.POPULATION_SPECIALIZATIONS.items():
                if 'gamma' in spec:
                    if pop_idx_cfg < 6: # Match existing hardcoded check p < 6
                        freq_band_idx = pop_idx_cfg # Assumes 0-5 are gamma bands
                        freq_range = config.FREQ_BANDS[freq_band_idx]
                        gamma_cell_config_map[pop_idx_cfg] = GammaCellUnit(
                                input_size=self.neurons_per_pop, # input to GammaCell is population state
                                hidden_size=min(32, self.neurons_per_pop // 2), # internal hidden size
                                band_freq=freq_range
                            )
            for i in range(self.num_populations):
                self.gamma_cells.append(gamma_cell_config_map.get(i, nn.Identity()))


        # Persistent projections for gamma cells' outputs
        if config.USE_OSCILLATORS:
            self.gamma_projections = nn.ModuleList()
            gamma_proj_config_map = {}
            for pop_idx_cfg, spec in config.POPULATION_SPECIALIZATIONS.items():
                 if 'gamma' in spec and pop_idx_cfg < 6 : # Check if it's a gamma cell that needs projection
                    # GammaCellUnit outputs hidden_size=min(32, self.neurons_per_pop // 2)
                    # Projection needs to map this to self.neurons_per_pop
                    gamma_cell_output_size = min(32, self.neurons_per_pop // 2)
                    gamma_proj_config_map[pop_idx_cfg] = nn.Linear(gamma_cell_output_size, self.neurons_per_pop)

            for i in range(self.num_populations):
                self.gamma_projections.append(gamma_proj_config_map.get(i, nn.Identity()))


        # Specialized activation functions per population
        self.activations = nn.ModuleList()
        for pop_idx in range(self.num_populations):
            spec = config.POPULATION_SPECIALIZATIONS.get(pop_idx, '')
            if 'gamma' in spec: self.activations.append(nn.SiLU())
            elif spec == 'integrator': self.activations.append(nn.ELU())
            else: self.activations.append(nn.Tanh())

        # Layer normalization for stability
        self.layer_norms = nn.ModuleList([
            nn.LayerNorm(self.neurons_per_pop) for _ in range(self.num_populations)
        ])

        # Batch normalization for training stability
        if config.USE_BATCH_NORM:
            self.batch_norms = nn.ModuleList([
                nn.BatchNorm1d(self.neurons_per_pop) for _ in range(self.num_populations)
            ])

        # Enhanced attention mechanism for population interaction
        if config.USE_ATTENTION:
            self.attention = PopulationAttention(
                self.neurons_per_pop, num_heads=config.ATTENTION_HEADS
            )

        self.register_buffer('hidden_states', None, persistent=False)
        self.timestep = 0
        self.influence_tracker = torch.zeros(
            (self.num_populations, self.num_populations), requires_grad=False
        )

    def _initialize_graph_structure(self):
        """Initialize graph structure with optimized connectivity for gamma processing."""
        adj_matrix = torch.zeros((self.total_neurons, self.total_neurons))
        for source_pop in range(self.num_populations):
            source_start = source_pop * self.neurons_per_pop
            source_end = (source_pop + 1) * self.neurons_per_pop
            for target_pop in range(self.num_populations):
                density = self.config.INTRA_POPULATION_DENSITY if source_pop == target_pop else self.config.INTER_POPULATION_DENSITY
                if density == 0: continue

                target_start = target_pop * self.neurons_per_pop
                target_end = (target_pop + 1) * self.neurons_per_pop
                influence_factor = self._get_population_influence(source_pop, target_pop)
                num_connections = int(self.neurons_per_pop * self.neurons_per_pop * density * influence_factor)

                if num_connections > 0:
                    source_indices = torch.randint(source_start, source_end, (num_connections,))
                    target_indices = torch.randint(target_start, target_end, (num_connections,))

                    source_spec = self.config.POPULATION_SPECIALIZATIONS.get(source_pop, 'default')
                    target_spec = self.config.POPULATION_SPECIALIZATIONS.get(target_pop, 'default')

                    if 'gamma' in source_spec and 'gamma' in target_spec:
                        weights = torch.zeros(num_connections)
                        positive_mask = torch.rand(num_connections) < 0.6
                        weights[positive_mask] = torch.rand(positive_mask.sum()) * 0.2
                        weights[~positive_mask] = -torch.rand((~positive_mask).sum()) * 0.15
                    else:
                        weights = torch.randn(num_connections) * 0.1
                        inhibitory_mask = torch.rand(num_connections) < 0.2
                        weights[inhibitory_mask] *= -1

                    # Use advanced indexing for efficiency if possible, otherwise loop
                    # This is a sparse assignment; looping is common for clarity here
                    for i in range(num_connections):
                        adj_matrix[source_indices[i], target_indices[i]] = weights[i]
        try:
            # Ensure adj_matrix is float for eigvals
            eigenvalues = torch.linalg.eigvals(adj_matrix.float())
            current_radius = torch.max(torch.abs(eigenvalues)).item()
            if current_radius > 1e-9: # Avoid division by zero or very small numbers
                adj_matrix = adj_matrix * (self.config.SPECTRAL_RADIUS / current_radius)
        except Exception as e:
            print(f"Error computing eigenvalues for spectral radius normalization: {e}")
            max_abs_val = torch.max(torch.abs(adj_matrix)).item()
            if max_abs_val > 1e-9:
                adj_matrix = adj_matrix * (self.config.SPECTRAL_RADIUS / max_abs_val)
        return adj_matrix.float() # Ensure it's float

    def _get_population_influence(self, source_pop, target_pop):
        source_spec = self.config.POPULATION_SPECIALIZATIONS.get(source_pop, 'default')
        target_spec = self.config.POPULATION_SPECIALIZATIONS.get(target_pop, 'default')
        factor = 1.0
        if 'gamma' in source_spec and 'gamma' in target_spec:
            source_level = self._get_gamma_level(source_spec)
            target_level = self._get_gamma_level(target_spec)
            if abs(source_level - target_level) == 1: factor = 1.5
            elif source_level > target_level: factor = 1.2
            else: factor = 0.8
        elif 'gamma' in source_spec and target_spec == 'integrator':
            source_level = self._get_gamma_level(source_spec)
            factor = 1.0 + 0.2 * source_level
        elif source_spec == 'integrator' and target_spec == 'context': factor = 2.0
        elif source_spec == 'context':
            if 'gamma' in target_spec:
                target_level = self._get_gamma_level(target_spec)
                factor = 1.0 - 0.1 * target_level
            elif target_spec == 'integrator': factor = 1.2
        return factor

    def _get_gamma_level(self, spec):
        levels = {'low_gamma': 0, 'mid_low_gamma': 1, 'mid_high_gamma': 2,
                  'high_gamma': 3, 'ultra_gamma_1': 4, 'ultra_gamma_2': 5}
        return levels.get(spec, 0)

    def reset_states(self, batch_size=1):
        device_to_use = self.adjacency_matrix.device # Use device of a parameter/buffer
        self.hidden_states = torch.randn(
            batch_size, self.total_neurons, device=device_to_use
        ) * 0.05
        if hasattr(self, 'gamma_cells'):
            for cell in self.gamma_cells:
                if hasattr(cell, 'reset_state'): cell.reset_state(batch_size)
        self.influence_tracker = torch.zeros(
            (self.num_populations, self.num_populations), device=device_to_use
        )
        self.timestep = 0

    def forward(self, x):
        """Forward pass with enhanced gamma oscillatory dynamics."""
        batch_size = x.shape[0]

        if self.hidden_states is None or self.hidden_states.shape[0] != batch_size:
            self.reset_states(batch_size)

        self.timestep += 1
        prev_states = self.hidden_states.clone()
        population_states = []

        for p in range(self.num_populations):
            pop_start = p * self.neurons_per_pop
            pop_end = (p + 1) * self.neurons_per_pop
            prev_pop_state = prev_states[:, pop_start:pop_end]
            input_proj = self.input_projections[p](x)

            # Ensure adjacency_matrix is on the same device as prev_states
            # Adjacency matrix is a parameter, so it should be on model's device
            recurrent_input = torch.matmul(
                prev_states, self.adjacency_matrix[:, pop_start:pop_end]
            )

            combined = input_proj + recurrent_input + self.population_biases[p]

            # Simplified influence tracking:
            if self.influence_tracker.device != self.adjacency_matrix.device:
                self.influence_tracker = self.influence_tracker.to(self.adjacency_matrix.device)

            for p_src in range(self.num_populations):
                if p_src != p:
                    src_start = p_src * self.neurons_per_pop
                    src_end = (p_src + 1) * self.neurons_per_pop
                    submatrix = self.adjacency_matrix[src_start:src_end, pop_start:pop_end]
                    influence = torch.sum(torch.abs(submatrix))
                    self.influence_tracker[p_src, p] += influence.item()

            activated = self.activations[p](combined)
            normalized_activation = self.layer_norms[p](activated)

            if hasattr(self, 'batch_norms') and self.config.USE_BATCH_NORM and self.training:
                 normalized_activation = self.batch_norms[p](normalized_activation)


            if hasattr(self, 'gamma_cells') and hasattr(self, 'gamma_projections') and p < 6: # Assuming gamma populations are first 6
                if not isinstance(self.gamma_cells[p], nn.Identity): # If it's a GammaCellUnit
                    # Input to GammaCellUnit is the current normalized activation of the population
                    gamma_cell_output = self.gamma_cells[p](normalized_activation) # [B, GammaCellHiddenSize (e.g. 32)]

                    spec = self.config.POPULATION_SPECIALIZATIONS.get(p, 'default')
                    if 'gamma' in spec:
                        level = self._get_gamma_level(spec)
                        osc_strength = 0.4 + 0.1 * level # Example: 0.4 to 0.9

                        # Project gamma_cell_output to population neuron size using pre-defined layer
                        # self.gamma_projections[p] is Linear(GammaCellHiddenSize, neurons_per_pop)
                        projected_gamma_output = self.gamma_projections[p](gamma_cell_output) # [B, neurons_per_pop]

                        # Mix oscillatory output with direct input pathway
                        normalized_activation = (1 - osc_strength) * normalized_activation + osc_strength * projected_gamma_output

            tc = torch.sigmoid(self.time_constants[p])
            leaky_out = (1 - tc) * prev_pop_state + tc * normalized_activation

            if self.training and self.config.NOISE_LEVEL > 0:
                noise = torch.randn_like(leaky_out) * self.config.NOISE_LEVEL
                leaky_out = leaky_out + noise

            population_states.append(leaky_out)

        if hasattr(self, 'attention') and self.config.USE_ATTENTION:
            pop_states_tensor = torch.stack(population_states, dim=1)
            attended_states, _ = self.attention(pop_states_tensor)
            for p in range(self.num_populations):
                spec = self.config.POPULATION_SPECIALIZATIONS.get(p, 'default')
                attention_weight = 0.4 if 'gamma' in spec else 0.3
                population_states[p] = (1 - attention_weight) * population_states[p] + attention_weight * attended_states[:, p]

        self.hidden_states = torch.cat(population_states, dim=1)
        return self.hidden_states


    def get_population_states(self):
        """Return states separated by population."""
        if self.hidden_states is None: return [] # Handle case before first forward
        population_states = []
        for p in range(self.num_populations):
            pop_start = p * self.neurons_per_pop
            pop_end = (p + 1) * self.neurons_per_pop
            population_states.append(self.hidden_states[:, pop_start:pop_end])
        return population_states

class GammaSpecificDecoder(nn.Module):
    """Specialized decoder focused on gamma band oscillation generation."""
    def __init__(self, input_size, output_size, freq_band, hidden_size=128):
        super().__init__()
        self.input_size = input_size
        self.output_size = output_size
        self.freq_band = freq_band  # (low, high) in Hz

        # Multi-layer decoder with SiLU activations
        self.decoder = nn.Sequential(
            nn.Linear(input_size, hidden_size),
            nn.LayerNorm(hidden_size),
            nn.SiLU(),
            nn.Dropout(0.2),
            nn.Linear(hidden_size, hidden_size // 2),
            nn.LayerNorm(hidden_size // 2),
            nn.SiLU(),
            nn.Linear(hidden_size // 2, output_size)
        )

        # Frequency-specific resonator bank
        self.resonator_bank = CoupledResonatorBank(
            freq_band=freq_band,
            num_resonators=config.OSCILLATORS_PER_BAND, # Use config
            coupling_strength=config.COUPLING_STRENGTH # Use config
        )


        # Amplitude envelope modulator
        self.envelope_modulator = nn.Sequential(
            nn.Linear(input_size, hidden_size // 2),
            nn.Tanh(),
            nn.Linear(hidden_size // 2, output_size),
            nn.Sigmoid()
        )

        # Phase modulator
        self.phase_modulator = nn.Sequential(
            nn.Linear(input_size, hidden_size // 2),
            nn.Tanh(),
            nn.Linear(hidden_size // 2, output_size),
            nn.Tanh()  # Output in [-1, 1] range for phase shifts
        )

    def reset_state(self, batch_size=1):
        """Reset resonator state."""
        self.resonator_bank.reset_state(batch_size)

    def forward(self, x): # x is [batch_size, input_size]
        """Forward pass with specialized gamma signal generation."""
        batch_size = x.shape[0]

        # Get base signal from decoder
        base_signal = self.decoder(x) # [batch_size, output_size]

        # Generate amplitude envelope
        envelope = self.envelope_modulator(x) # [batch_size, output_size]

        # Generate phase modulation
        phase_mod = self.phase_modulator(x) # [batch_size, output_size]

        # Get resonator signal - input to resonator is mean of x across input_size
        resonator_input_scalar = x.mean(dim=1) # [batch_size]
        # resonator_bank expects [batch_size] or [batch_size,1]
        resonator_output_scalar = self.resonator_bank(resonator_input_scalar)  # [batch_size]


        # Expand resonator output to match output dimensions
        # resonator_output_scalar is [B], want [B, output_size]
        resonator_output_expanded = resonator_output_scalar.unsqueeze(1).expand(-1, self.output_size)


        # Apply phase modulation to resonator output
        # phase_indices for distinct phase per output unit, phase_mod for learned shift
        phase_indices = torch.arange(self.output_size, device=x.device, dtype=x.dtype).unsqueeze(0)
        phase_indices = phase_indices.expand(batch_size, -1) / self.output_size * 2 * np.pi
        phase_offset = phase_mod * np.pi  # Scale to [-π, π] # phase_mod is [B, output_size]

        # Element-wise multiplication and cosine
        phase_shifted_oscillation = resonator_output_expanded * torch.cos(phase_indices + phase_offset)


        # Combine base signal with modulated resonator output
        output = 0.5 * base_signal + 0.5 * envelope * phase_shifted_oscillation

        return output


class EnhancedGammaESN(nn.Module):
    """Enhanced Graph Echo State Network optimized for gamma oscillation prediction."""
    def __init__(self, config):
        super().__init__()
        self.config = config

        # Input dimensions
        self.num_electrodes = config.NUM_ELECTRODES
        self.num_freq_bands = len(config.FREQ_BANDS)
        self.predict_size = config.PREDICT_SIZE

        # Enhanced feature projections - make sure dimensions are correct
        self.raw_projection = nn.Linear(self.num_electrodes, config.HIDDEN_SIZE // 4)
        self.phase_projection = nn.Linear(self.num_freq_bands * self.num_electrodes, config.HIDDEN_SIZE // 4)
        self.amplitude_projection = nn.Linear(self.num_freq_bands * self.num_electrodes, config.HIDDEN_SIZE // 4)
        self.freq_projection = nn.Linear(self.num_freq_bands * self.num_electrodes, config.HIDDEN_SIZE // 4)

        # Coherence projection (unused in current forward, but defined)
        # Input shape for coherence is [batch, num_bands, num_electrodes, num_electrodes]
        # Flattened: num_bands * num_electrodes * num_electrodes
        self.coherence_projection = nn.Linear(
             self.num_freq_bands * self.num_electrodes * self.num_electrodes,
             config.HIDDEN_SIZE // 8 # This makes total HIDDEN_SIZE not sum up if used as is
        )
        # If all 5 features raw, phase, amp, freq, coherence are used, their projection sizes should sum to HIDDEN_SIZE
        # Current setup: 4 * (HIDDEN_SIZE//4) = HIDDEN_SIZE. Coherence is extra or replaces one.
        # As coherence is not used in forward, this is not an issue yet.


        # Enhanced feature fusion with residual connections
        self.feature_fusion = nn.Sequential(
            nn.Linear(config.HIDDEN_SIZE, config.HIDDEN_SIZE), # Assuming combined_features matches HIDDEN_SIZE
            nn.LayerNorm(config.HIDDEN_SIZE),
            nn.GELU(),
            nn.Dropout(config.DROPOUT),
            nn.Linear(config.HIDDEN_SIZE, config.HIDDEN_SIZE),
            # Added LayerNorm + GELU for consistency like in other blocks
            nn.LayerNorm(config.HIDDEN_SIZE),
            nn.GELU()
        )


        # Graph reservoir
        self.graph_reservoir = EnhancedGraphReservoirCell(config)

        # Temporal processing with bidirectional LSTM for better context integration
        reservoir_size = config.NUM_POPULATIONS * config.NEURONS_PER_POPULATION
        self.lstm = nn.LSTM(
            input_size=reservoir_size,
            hidden_size=config.READOUT_HIDDEN,
            num_layers=2, # A common choice for LSTMs
            batch_first=True,
            dropout=config.DROPOUT if 2 > 1 else 0, # Dropout only if num_layers > 1
            bidirectional=True
        )


        # Linear projection to combine bidirectional outputs
        self.bidirectional_proj = nn.Linear(config.READOUT_HIDDEN * 2, config.READOUT_HIDDEN)

        # Enhanced readout with multiple layers
        readout_layers = []
        current_readout_dim = config.READOUT_HIDDEN
        for i in range(config.READOUT_LAYERS): # Loop for all layers
            next_readout_dim = config.READOUT_HIDDEN // (2**(i+1)) if i < config.READOUT_LAYERS -1 else \
                                config.READOUT_HIDDEN // (2**(config.READOUT_LAYERS-1)) # last layer output size
            if i == config.READOUT_LAYERS -1 : # last layer doesn't halve further typically for the main branch
                 next_readout_dim = current_readout_dim # keep same dim or specific final dim
                 # The logic was config.READOUT_HIDDEN // (2**(config.READOUT_LAYERS-1))
                 # For 3 layers, input to decoders is HIDDEN // 4.
                 # L0: H -> H/2
                 # L1: H/2 -> H/4
                 # L2 (not in loop): H/4 is input to decoders.
                 # The loop range(config.READOUT_LAYERS - 1) implies (config.READOUT_LAYERS-1) blocks
                 # and then the output of the last block is taken.
                 # Original loop was range(config.READOUT_LAYERS - 1)
                 # Original in_size for i=0: config.READOUT_HIDDEN
                 # Original out_size for i=0: config.READOUT_HIDDEN // 2
                 # Original in_size for i=1: config.READOUT_HIDDEN // 2
                 # Original out_size for i=1: config.READOUT_HIDDEN // 4
                 # Final output dim is config.READOUT_HIDDEN // 4

                 # Corrected loop based on original intent:
                 # Input to readout_common is config.READOUT_HIDDEN
                 # Output of readout_common is config.READOUT_HIDDEN // (2**(config.READOUT_LAYERS-1))

                 in_s = config.READOUT_HIDDEN
                 out_s = config.READOUT_HIDDEN
                 for i_layer in range(config.READOUT_LAYERS -1): # Num blocks = READOUT_LAYERS -1
                     out_s = config.READOUT_HIDDEN // (2**(i_layer+1))
                     readout_layers.extend([
                         nn.Linear(in_s, out_s),
                         nn.LayerNorm(out_s),
                         nn.GELU(),
                         nn.Dropout(0.1) # Original dropout was 0.1
                     ])
                     in_s = out_s
                 # Final output dimension from these blocks is config.READOUT_HIDDEN // (2**(config.READOUT_LAYERS-1))
                 self.final_readout_dim = in_s


        self.readout_common = nn.Sequential(*readout_layers)


        # Specialized gamma decoders for each frequency band
        self.gamma_decoders = nn.ModuleList()
        for band in config.FREQ_BANDS:
            self.gamma_decoders.append(
                GammaSpecificDecoder(
                    input_size=self.final_readout_dim, # Use calculated final dim
                    output_size=self.num_electrodes,
                    freq_band=band,
                    hidden_size=max(self.final_readout_dim // 2, self.num_electrodes) # Ensure hidden_size is reasonable
                )
            )


        # Main output decoder (for full signal)
        self.main_decoder = nn.Linear(
            self.final_readout_dim, # Use calculated final dim
            self.num_electrodes
        )


        # Prediction head for each timestep
        self.timestep_heads = nn.ModuleList([
            nn.Linear(self.num_electrodes * (len(config.FREQ_BANDS) + 1), self.num_electrodes)
            for _ in range(config.PREDICT_SIZE)
        ])

        self.state_history = []
        self.causal_connections = torch.zeros((config.NUM_POPULATIONS, config.NUM_POPULATIONS))

    def reset_states(self):
        self.graph_reservoir.reset_states()
        if hasattr(self, 'gamma_decoders'): # Check existence before iterating
            for decoder in self.gamma_decoders:
                if hasattr(decoder, 'reset_state'): decoder.reset_state()
        self.state_history = []


    def get_causal_influence(self):
        return self.graph_reservoir.influence_tracker

    def get_state_trajectories(self):
        return self.state_history

    def forward(self, x_dict, return_states=False):
        x_raw = x_dict['raw']
        x_phase = x_dict['phase']
        x_amplitude = x_dict['amplitude']
        x_inst_freq = x_dict['inst_freq']
        # x_spectral = x_dict['spectral'] # Unused currently
        # x_coherence = x_dict['coherence'] # Unused currently

        batch_size = x_raw.shape[0]
        context_size = x_raw.shape[1]
        device = x_raw.device

        self.reset_states() # Reset states for each new sequence forward pass

        if return_states: self.state_history = []

        reservoir_sequence = []
        for t in range(context_size):
            raw_t = x_raw[:, t, :]
            raw_features = self.raw_projection(raw_t)

            phase_t = x_phase[:, :, t, :].reshape(batch_size, -1)
            phase_features = self.phase_projection(phase_t)

            amplitude_t = x_amplitude[:, :, t, :].reshape(batch_size, -1)
            amplitude_features = self.amplitude_projection(amplitude_t)

            freq_t = x_inst_freq[:, :, t, :].reshape(batch_size, -1)
            freq_features = self.freq_projection(freq_t)

            combined_features = torch.cat([
                raw_features, phase_features, amplitude_features, freq_features
            ], dim=1)

            # Apply residual connection to feature_fusion if USE_SKIP_CONNECTIONS is True
            fused_input = combined_features
            fused_output = self.feature_fusion(fused_input)
            if self.config.USE_SKIP_CONNECTIONS:
                 fused_features = fused_input + fused_output # Additive skip
            else:
                 fused_features = fused_output


            reservoir_state = self.graph_reservoir(fused_features)
            if return_states:
                self.state_history.append(self.graph_reservoir.get_population_states())
            reservoir_sequence.append(reservoir_state)

        reservoir_sequence_tensor = torch.stack(reservoir_sequence, dim=1)
        lstm_out, (h_n, _) = self.lstm(reservoir_sequence_tensor)

        # Use last hidden state of LSTM from all layers/directions as input to bidirectional_proj
        # h_n is [num_layers*num_dir, batch, hidden_size]
        # Concatenate final forward and backward hidden states of the last layer
        # If num_layers = 2, directions = 2:
        # h_n[2] = layer 1 backward, h_n[3] = layer 1 forward
        # h_n[0] = layer 0 forward, h_n[1] = layer 0 backward (if num_layers=1, bidirectional)
        # h_n[-2,:,:] is last layer forward, h_n[-1,:,:] is last layer backward
        h_forward = h_n[-2, :, :]
        h_backward = h_n[-1, :, :]
        bidirectional_hidden = torch.cat([h_forward, h_backward], dim=1)
        final_hidden_lstm = self.bidirectional_proj(bidirectional_hidden) # [batch, READOUT_HIDDEN]


        common_features = self.readout_common(final_hidden_lstm)

        gamma_outputs = [decoder(common_features) for decoder in self.gamma_decoders]
        main_output = self.main_decoder(common_features)

        all_outputs_stacked = torch.stack([main_output] + gamma_outputs, dim=1)
        timestep_input_reshaped = all_outputs_stacked.reshape(batch_size, -1)

        predictions = torch.zeros(batch_size, self.predict_size, self.num_electrodes, device=device)
        for t_pred in range(self.predict_size):
            predictions[:, t_pred, :] = self.timestep_heads[t_pred](timestep_input_reshaped)

        return (predictions, self.state_history) if return_states else predictions


def compute_fft_torch(signal, fs=1000):
    """Compute FFT using PyTorch for better GPU compatibility."""
    n = signal.shape[-1] # Operate on last dimension (time)
    rfft_output = torch.fft.rfft(signal, n=n, dim=-1) # Use dim=-1 for [B,C,T] or [B,T]
    magnitudes = torch.abs(rfft_output) / n
    phases = torch.angle(rfft_output)
    frequencies = torch.fft.rfftfreq(n, d=1.0/fs, device=signal.device) # Correct d for sampling interval
    return frequencies, magnitudes, phases


class PhaseConsistencyLoss(nn.Module):
    """Loss function that measures consistency of phase relationships."""
    def __init__(self):
        super().__init__()

    def forward(self, pred_phases, true_phases):
        """
        Compute phase consistency loss.
        Args:
            pred_phases: Predicted phases [..., frequencies, (time_stft_bins)] or [..., frequencies] for FFT
            true_phases: True phases   [..., frequencies, (time_stft_bins)] or [..., frequencies] for FFT
        """
        phase_diff = pred_phases - true_phases
        # Wrap phase difference to [-pi, pi] for stability before cos/sin
        phase_diff_wrapped = torch.remainder(phase_diff + np.pi, 2 * np.pi) - np.pi

        real_part = torch.cos(phase_diff_wrapped)
        imag_part = torch.sin(phase_diff_wrapped)
        complex_diff = torch.complex(real_part, imag_part)

        # Mean over frequencies (or time_stft_bins then frequencies)
        # If FFT output [..., freqs], mean over freqs
        mean_vector = torch.mean(complex_diff, dim=-1) # Mean over last dim (freqs)
        if mean_vector.ndim > complex_diff.ndim - 1 and mean_vector.shape[-1] != 1: # if STFT output
             mean_vector = torch.mean(mean_vector, dim=-1) # then mean over time bins

        consistency = torch.abs(mean_vector)
        loss = 1.0 - consistency
        return torch.mean(loss) # Mean over batch and other dims


class EnhancedGammaLoss(nn.Module):
    """Enhanced loss function optimized for gamma oscillation prediction."""
    def __init__(self, config):
        super().__init__()
        self.config = config
        self.time_weight = config.TIME_LOSS_WEIGHT
        self.freq_weight = config.FREQ_LOSS_WEIGHT
        self.phase_weight = config.PHASE_LOSS_WEIGHT
        self.envelope_weight = config.ENVELOPE_LOSS_WEIGHT
        self.gamma_band_weights = config.GAMMA_BAND_WEIGHTS

        self.mse_loss = nn.MSELoss()
        self.phase_consistency_loss = PhaseConsistencyLoss()
        self.sampling_rate = float(config.SAMPLING_RATE) # Ensure float

        self.freq_band_map = {i: band for i, band in enumerate(config.FREQ_BANDS)}

    def gamma_frequency_loss(self, pred, target):
        batch_size, seq_len, num_channels = pred.shape

        # pred and target are [batch, time, channels]
        # For FFT, want [batch*channels, time]
        pred_reshaped = pred.permute(0, 2, 1).reshape(-1, seq_len)
        target_reshaped = target.permute(0, 2, 1).reshape(-1, seq_len)

        pred_freqs, pred_mags, pred_phases = compute_fft_torch(pred_reshaped, self.sampling_rate)
        target_freqs, target_mags, target_phases = compute_fft_torch(target_reshaped, self.sampling_rate)
        # freqs shape [n_fft_bins], mags/phases shape [batch*channels, n_fft_bins]

        # Initialize band weights tensor on the correct device
        weights = torch.ones_like(pred_freqs, device=pred.device) * 0.1
        for band_idx, (low, high) in self.freq_band_map.items():
            band_mask = (pred_freqs >= low) & (pred_freqs <= high)
            band_weight_val = self.gamma_band_weights.get((low, high), 1.0)
            weights = torch.where(band_mask, torch.tensor(band_weight_val, device=weights.device), weights)

        # Weighted MSE for magnitude (spectrum)
        # pred_mags, target_mags are [B*C, n_fft_bins]. weights is [n_fft_bins]
        mag_diff_sq = (pred_mags - target_mags)**2
        # Multiply by weights (broadcasts over B*C dim): [B*C, n_fft_bins] * [1, n_fft_bins]
        weighted_mag_loss = torch.mean(mag_diff_sq * weights.unsqueeze(0))
        freq_loss = weighted_mag_loss

        # Phase consistency loss for gamma frequencies
        phase_loss_val = torch.tensor(0.0, device=pred.device)
        if self.config.USE_PHASE_CONSISTENCY_LOSS:
            gamma_mask = pred_freqs >= 30
            if gamma_mask.any(): # Check if any gamma frequencies exist
                # pred_phases/target_phases are [B*C, n_fft_bins]
                # Select gamma frequencies: [B*C, n_gamma_bins]
                pred_gamma_phases = pred_phases[:, gamma_mask]
                target_gamma_phases = target_phases[:, gamma_mask]
                if pred_gamma_phases.shape[-1] > 0: # Ensure there are bins
                     phase_loss_val = self.phase_consistency_loss(pred_gamma_phases, target_gamma_phases)


        # Envelope loss using magnitude of gamma frequencies
        envelope_loss_val = torch.tensor(0.0, device=pred.device)
        gamma_mask_env = pred_freqs >= 30 # Re-evaluate or use gamma_mask
        if gamma_mask_env.any():
            pred_gamma_mags = pred_mags[:, gamma_mask_env]
            target_gamma_mags = target_mags[:, gamma_mask_env]
            if pred_gamma_mags.shape[-1] > 0:
                 envelope_loss_val = F.mse_loss(pred_gamma_mags, target_gamma_mags)

        return freq_loss, phase_loss_val, envelope_loss_val


    def forward(self, pred, target):
        time_loss = self.mse_loss(pred, target)
        freq_loss, phase_loss, envelope_loss = self.gamma_frequency_loss(pred, target)

        total_loss = (
            self.time_weight * time_loss +
            self.freq_weight * freq_loss +
            self.phase_weight * phase_loss +
            self.envelope_weight * envelope_loss
        )
        return total_loss, time_loss, freq_loss, phase_loss, envelope_loss


class CurriculumScheduler:
    """Curriculum learning scheduler for progressive gamma training."""
    def __init__(self, config):
        self.config = config
        self.current_epoch = 0
        self.phase = 0
        self.num_phases = 3  # Number of curriculum phases
        self.phase_length = config.CURRICULUM_PHASE_LENGTH

    def update(self, epoch):
        """Update curriculum phase based on epoch."""
        self.current_epoch = epoch
        self.phase = min(self.num_phases - 1, epoch // self.phase_length)

    def get_gamma_weights(self):
        """Get gamma band weights for current curriculum phase."""
        base_weights = self.config.GAMMA_BAND_WEIGHTS.copy()
        if self.phase == 0:
            for (low, high), weight in base_weights.items():
                base_weights[(low, high)] = weight * 1.5 if low < 80 else weight * 0.5
        elif self.phase == 1:
            for (low, high), weight in base_weights.items():
                base_weights[(low, high)] = weight * 1.5 if 50 <= low < 180 else weight * 0.7
        elif self.phase == 2:
            for (low, high), weight in base_weights.items():
                 if low >= 120: base_weights[(low, high)] = weight * 1.5
        return base_weights

    def get_loss_weights(self):
        """Get loss component weights for current curriculum phase."""
        if self.phase == 0: return {'time': 0.5, 'freq': 0.5, 'phase': 0.0, 'envelope': 0.0}
        elif self.phase == 1: return {'time': 0.3, 'freq': 0.4, 'phase': 0.2, 'envelope': 0.1}
        else: return {'time': 0.25, 'freq': 0.45, 'phase': 0.2, 'envelope': 0.1}


def visualize_electrode_trajectories_3d(model, inputs_dict, device, sample_idx=0, save_path=None,
                                     title="High-Dimensional Electrode Neural Trajectories"):
    """
    Visualize the trajectories of all electrodes in 3D space with improved aesthetics.
    """
    print("Generating high-dimensional electrode trajectory visualization...")

    # Set model to evaluation mode
    model.eval()

    with torch.no_grad():
        # Get the raw electrode signals
        raw_inputs = inputs_dict['raw'].to(device)  # Shape: [batch, context_size, num_electrodes]
        batch_size, context_size, num_electrodes = raw_inputs.shape

        # Extract electrode signals for the selected sample
        electrode_signals = raw_inputs[sample_idx].cpu().numpy()  # Shape: [context_size, num_electrodes]

    # Create a Plotly figure for 3D visualization
    fig = go.Figure()

    # Define better color palette - using Turbo for clearer separation of electrodes
    colors = px.colors.sequential.Turbo
    color_scale = [colors[int(i / num_electrodes * (len(colors)-1))] for i in range(num_electrodes)]

    # For each electrode, create a 3D trajectory using delay embedding
    for e_idx in range(num_electrodes):
        # Get electrode signal over time
        electrode_signal = electrode_signals[:, e_idx]  # Shape: [context_size]

        # We need to create a 3D representation
        # For time series data, we can use delay embedding to create a 3D view
        delay = max(1, context_size // 25)  # Shorter delay for better visualization

        if context_size > 3*delay:  # Make sure we have enough points
            x = electrode_signal[:-2*delay]
            y = electrode_signal[delay:-delay]
            z = electrode_signal[2*delay:]

            # Add 3D trace for this electrode
            fig.add_trace(go.Scatter3d(
                x=x, y=y, z=z,
                mode='lines',
                line=dict(
                    color=color_scale[e_idx],
                    width=2,
                ),
                opacity=0.7,
                name=f"E{e_idx}",
                hoverinfo='text',
                hovertext=[f"Electrode {e_idx}<br>Time: {t}<br>Value: {electrode_signal[t]:.4f}"
                          for t in range(len(x))],
                showlegend=True if e_idx % max(1, num_electrodes // 8) == 0 else False  # Show limited legends
            ))

            # Highlight start and end points
            fig.add_trace(go.Scatter3d(
                x=[x[0]], y=[y[0]], z=[z[0]],
                mode='markers',
                marker=dict(
                    size=4,
                    color='green',
                    symbol='circle',
                    line=dict(color='black', width=1)
                ),
                name=f"Start (t=0)",
                showlegend=e_idx == 0  # Only show once in legend
            ))

            fig.add_trace(go.Scatter3d(
                x=[x[-1]], y=[y[-1]], z=[z[-1]],
                mode='markers',
                marker=dict(
                    size=4,
                    color='red',
                    symbol='circle',
                    line=dict(color='black', width=1)
                ),
                name=f"End (t={len(x)-1})",
                showlegend=e_idx == 0  # Only show once in legend
            ))

    # Update layout with improved aesthetics
    fig.update_layout(
        title=dict(
            text=title,
            y=0.95,
            x=0.5,
            xanchor='center',
            yanchor='top',
            font=dict(size=24, family="Arial", color="#333333")
        ),
        scene=dict(
            xaxis=dict(
                title="t",
                backgroundcolor="rgba(255, 255, 255, 0.95)",
                gridcolor="#e0e0e0",
                showbackground=True,
                zerolinecolor="#cccccc",
            ),
            yaxis=dict(
                title="t+delay",
                backgroundcolor="rgba(255, 255, 255, 0.95)",
                gridcolor="#e0e0e0",
                showbackground=True,
                zerolinecolor="#cccccc",
            ),
            zaxis=dict(
                title="t+2*delay",
                backgroundcolor="rgba(255, 255, 255, 0.95)",
                gridcolor="#e0e0e0",
                showbackground=True,
                zerolinecolor="#cccccc",
            ),
            camera=dict(eye=dict(x=1.5, y=1.5, z=1.2))
        ),
        margin=dict(l=0, r=0, t=40, b=0),
        width=1200, height=800,
        paper_bgcolor="#f8f9fa",
        legend=dict(
            itemsizing='constant',
            font=dict(size=10),
            yanchor="top",
            y=0.99,
            xanchor="left",
            x=0.01,
            bgcolor="rgba(255,255,255,0.8)"
        ),
    )

    # Add annotation explaining visualization
    fig.add_annotation(
        x=0.02,
        y=0.02,
        xref="paper",
        yref="paper",
        text="This visualization shows the trajectory of electrode activity in delay-embedded space.<br>Each electrode forms a distinct trajectory representing its temporal dynamics.",
        showarrow=False,
        font=dict(size=12, color="#555555"),
        align="left",
        bgcolor="rgba(255,255,255,0.8)",
        bordercolor="#e0e0e0",
        borderwidth=1,
        borderpad=4,
        opacity=0.9
    )

    # Save if path is provided
    if save_path:
        pio.write_html(fig, save_path)
        print(f"Electrode trajectory visualization saved to {save_path}")

    return fig

def create_electrode_influence_network(model, inputs_dict, device, sample_idx=0, save_path=None,
                                      title="Electrode Cross-Influence Network"):
    """
    Create a visualization of the influence network between electrodes with improved aesthetics.
    """
    print("Analyzing cross-electrode influence patterns...")

    # Import plotly express locally if needed
    import plotly.express as px

    # Set model to evaluation mode
    model.eval()

    with torch.no_grad():
        # Get the raw electrode signals
        raw_inputs = inputs_dict['raw'].to(device)  # Shape: [batch, context_size, num_electrodes]
        batch_size, context_size, num_electrodes = raw_inputs.shape

        # Extract electrode signals for the selected sample
        electrode_signals = raw_inputs[sample_idx].cpu().numpy()  # Shape: [context_size, num_electrodes]

    # Initialize influence matrix between electrodes
    influence_matrix = np.zeros((num_electrodes, num_electrodes))

    # Calculate influence using lagged correlation
    max_lag = min(5, context_size // 4)

    print("Computing electrode influence metrics...")
    for source_idx in range(num_electrodes):
        for target_idx in range(num_electrodes):
            if source_idx != target_idx:
                # Get the signals for both electrodes
                source_signal = electrode_signals[:, source_idx]
                target_signal = electrode_signals[:, target_idx]

                # Calculate maximum lagged correlation
                max_corr = 0
                for lag in range(1, max_lag + 1):
                    if lag < len(source_signal):
                        source_lagged = source_signal[:-lag]
                        target_lagged = target_signal[lag:]

                        if len(source_lagged) > 1 and len(target_lagged) > 1:
                            try:
                                corr_coef = np.corrcoef(source_lagged, target_lagged)[0, 1]
                                if not np.isnan(corr_coef):
                                    max_corr = max(max_corr, abs(corr_coef))
                            except:
                                pass

                # Store the influence score
                influence_matrix[source_idx, target_idx] = max_corr

    # Calculate global influence measures
    outgoing_influence = np.sum(influence_matrix, axis=1)  # Influence on others
    incoming_influence = np.sum(influence_matrix, axis=0)  # Influenced by others
    total_influence = outgoing_influence + incoming_influence

    # Apply threshold to focus on significant influences (top 25%)
    if np.any(influence_matrix > 0):
        threshold = np.percentile(influence_matrix[influence_matrix > 0], 75)
        significant_influence = influence_matrix.copy()
        significant_influence[significant_influence < threshold] = 0
    else:
        significant_influence = influence_matrix.copy()

    # Create a network visualization using NetworkX and Plotly with improved aesthetics
    G = nx.DiGraph()

    # Add nodes (electrodes)
    for i in range(num_electrodes):
        # Node size based on total influence
        node_size = 10 + 30 * (total_influence[i] / max(total_influence) if np.max(total_influence) > 0 else 1)

        # Node color based on outgoing/incoming ratio (sender vs receiver character)
        if incoming_influence[i] > 0:
            sender_receiver_ratio = outgoing_influence[i] / incoming_influence[i]
        else:
            sender_receiver_ratio = outgoing_influence[i] if outgoing_influence[i] > 0 else 0

        G.add_node(i, size=node_size, influence=total_influence[i], ratio=sender_receiver_ratio)

    # Add edges for significant influences
    for source in range(num_electrodes):
        for target in range(num_electrodes):
            if significant_influence[source, target] > 0:
                G.add_edge(source, target, weight=significant_influence[source, target])

    # Calculate layout - using Kamada-Kawai layout for better spacing
    try:
        # Try to use a more aesthetically pleasing layout
        pos = nx.kamada_kawai_layout(G)
    except:
        try:
            pos = nx.spring_layout(G, k=0.3, iterations=100, seed=42)
        except:
            pos = {i: (np.cos(i*2*np.pi/num_electrodes), np.sin(i*2*np.pi/num_electrodes)) for i in range(num_electrodes)}

    # Create figure
    fig = go.Figure()

    # Define better color palette for edges using Plotly colors
    edge_colors = ['rgba(31, 119, 180, 0.7)', 'rgba(31, 119, 180, 0.8)',
                  'rgba(31, 119, 180, 0.9)', 'rgba(31, 119, 180, 1.0)']

    # Add edges with arrows using a more appealing color scheme
    edge_traces = []

    for edge in G.edges(data=True):
        source, target, data = edge
        x0, y0 = pos[source]
        x1, y1 = pos[target]
        weight = data['weight']

        # Normalize weight for color mapping
        norm_weight = (weight - np.min(significant_influence[significant_influence > 0])) / \
                     (np.max(significant_influence) - np.min(significant_influence[significant_influence > 0]))

        # Select color based on weight
        color_idx = min(len(edge_colors)-1, int(norm_weight * len(edge_colors)))
        edge_color = edge_colors[color_idx]

        edge_trace = go.Scatter(
            x=[x0, x1, None],
            y=[y0, y1, None],
            line=dict(
                width=1 + 3 * weight,
                color=edge_color,
            ),
            hoverinfo='text',
            text=f"Influence: E{source} → E{target}<br>Strength: {weight:.3f}",
            mode='lines',
            showlegend=False
        )
        edge_traces.append(edge_trace)

        # Add arrow head
        arrow_length = 0.03
        arrow_width = 0.015

        # Calculate direction vector
        dx, dy = x1 - x0, y1 - y0
        length = np.sqrt(dx**2 + dy**2)

        if length > 0:
            # Normalize
            dx, dy = dx/length, dy/length

            # Position arrow slightly before target
            ax, ay = x1 - dx*arrow_length*2, y1 - dy*arrow_length*2

            # Perpendicular vector
            px, py = -dy, dx

            # Arrow shape
            arrow_x = [
                ax + px*arrow_width,
                x1,
                ax - px*arrow_width,
                ax + px*arrow_width  # Close shape
            ]

            arrow_y = [
                ay + py*arrow_width,
                y1,
                ay - py*arrow_width,
                ay + py*arrow_width  # Close shape
            ]

            arrow_trace = go.Scatter(
                x=arrow_x,
                y=arrow_y,
                line=dict(width=0),
                fill='toself',
                fillcolor=edge_color,
                hoverinfo='none',
                showlegend=False
            )
            edge_traces.append(arrow_trace)

    # Define better color palette for nodes
    sender_color = 'rgba(214, 39, 40, 0.9)'  # Red for senders
    receiver_color = 'rgba(31, 119, 180, 0.9)'  # Blue for receivers
    balanced_color = 'rgba(148, 103, 189, 0.9)'  # Purple for balanced

    # Add nodes
    node_x = []
    node_y = []
    node_text = []
    node_size = []
    node_colors = []

    for node in G.nodes():
        x, y = pos[node]
        node_x.append(x)
        node_y.append(y)

        ratio = G.nodes[node].get('ratio', 1)

        # Color based on ratio (sender/receiver) with improved colors
        if ratio < 0.5:  # More incoming (receiver)
            color = receiver_color
        elif ratio > 2:  # More outgoing (sender)
            color = sender_color
        else:  # Balanced
            color = balanced_color

        node_colors.append(color)

        # Size based on influence
        size = G.nodes[node].get('size', 10)
        node_size.append(size)

        # Hover text
        node_text.append(f"Electrode {node}<br>Total Influence: {G.nodes[node].get('influence', 0):.3f}<br>" +
                         f"Out/In Ratio: {ratio:.2f}")

    # Add node trace
    node_trace = go.Scatter(
        x=node_x,
        y=node_y,
        mode='markers+text',
        text=[f"E{i}" for i in range(len(node_x))],
        textposition="top center",
        textfont=dict(family="Arial", size=10, color="black"),
        marker=dict(
            size=node_size,
            color=node_colors,
            line=dict(width=1, color='rgb(40,40,40)'),
        ),
        hoverinfo='text',
        hovertext=node_text
    )

    # Add all traces to figure
    for trace in edge_traces:
        fig.add_trace(trace)

    fig.add_trace(node_trace)

    # Add a legend for node types
    fig.add_trace(go.Scatter(
        x=[None], y=[None],
        mode='markers',
        marker=dict(size=15, color=sender_color),
        name='Sender electrode',
        showlegend=True
    ))

    fig.add_trace(go.Scatter(
        x=[None], y=[None],
        mode='markers',
        marker=dict(size=15, color=receiver_color),
        name='Receiver electrode',
        showlegend=True
    ))

    fig.add_trace(go.Scatter(
        x=[None], y=[None],
        mode='markers',
        marker=dict(size=15, color=balanced_color),
        name='Balanced electrode',
        showlegend=True
    ))

    # Update layout with improved aesthetics
    fig.update_layout(
        title=dict(
            text=title,
            y=0.98,
            x=0.5,
            xanchor='center',
            yanchor='top',
            font=dict(family="Arial", size=24, color="#333333")
        ),
        showlegend=True,
        hovermode='closest',
        margin=dict(l=20, r=20, t=60, b=20),
        xaxis=dict(
            showgrid=False,
            zeroline=False,
            showticklabels=False,
            domain=[0, 1]
        ),
        yaxis=dict(
            showgrid=False,
            zeroline=False,
            showticklabels=False,
            domain=[0, 1]
        ),
        width=1100, height=900,
        paper_bgcolor='rgb(248, 249, 250)',
        plot_bgcolor='rgb(248, 249, 250)',
        legend=dict(
            yanchor="top",
            y=0.99,
            xanchor="right",
            x=0.99,
            itemsizing='constant',
            bgcolor="rgba(255, 255, 255, 0.8)",
            bordercolor="#e0e0e0",
            borderwidth=1
        )
    )

    # Add explanation annotation
    fig.add_annotation(
        x=0.02,
        y=0.02,
        xref="paper",
        yref="paper",
        text="This network visualizes how information flows between electrode populations.<br>" +
             "• Node size indicates total influence<br>" +
             "• Node color shows electrode role (sender/receiver)<br>" +
             "• Edge width and color show influence strength",
        showarrow=False,
        align="left",
        bgcolor="rgba(255, 255, 255, 0.8)",
        bordercolor="#e0e0e0",
        borderwidth=1,
        borderpad=4,
        font=dict(size=12),
    )

    # Save figure if path provided
    if save_path:
        pio.write_html(fig, save_path)
        print(f"Electrode influence network saved to {save_path}")

    return fig


def create_decision_tree_causal_flow(model, inputs_dict, device, sample_idx=0, save_path=None,
                                    title="Causal Flow Between Neural Electrodes"):
    """
    Create a decision tree-like causal flow visualization with improved aesthetics.
    """
    print("Generating causal decision tree visualization...")

    # Set model to evaluation mode
    model.eval()

    with torch.no_grad():
        # Get the raw electrode signals
        raw_inputs = inputs_dict['raw'].to(device)  # Shape: [batch, context_size, num_electrodes]
        batch_size, context_size, num_electrodes = raw_inputs.shape

        # Extract electrode signals for the selected sample
        electrode_signals = raw_inputs[sample_idx].cpu().numpy()  # Shape: [context_size, num_electrodes]

    # Define time segments for causal analysis (divide time into segments)
    n_segments = min(5, context_size // 10)  # At least 10 time points per segment
    if n_segments < 2:
        n_segments = 2  # Minimum 2 segments

    segment_size = context_size // n_segments

    # Compute influence between electrodes across time segments
    segment_influences = []

    # For each consecutive pair of time segments
    for seg_idx in range(n_segments - 1):
        start_idx = seg_idx * segment_size
        mid_idx = start_idx + segment_size
        end_idx = mid_idx + segment_size
        end_idx = min(end_idx, context_size)

        # Influence matrix for this segment pair
        influence_matrix = np.zeros((num_electrodes, num_electrodes))

        # For each source electrode in segment 1
        for source_idx in range(num_electrodes):
            # For each target electrode in segment 2
            for target_idx in range(num_electrodes):
                if source_idx != target_idx:
                    # Source electrode in segment 1
                    source_seg1 = electrode_signals[start_idx:mid_idx, source_idx]
                    # Target electrode in segment 2
                    target_seg2 = electrode_signals[mid_idx:end_idx, target_idx]

                    if len(source_seg1) > 0 and len(target_seg2) > 0:
                        # Compute influence using lagged correlation
                        try:
                            # Calculate maximum lagged correlation
                            max_lag = min(3, len(source_seg1) // 2)
                            max_corr = 0

                            for lag in range(1, max_lag + 1):
                                if lag < len(source_seg1):
                                    source_lagged = source_seg1[:-lag]
                                    target_lagged = target_seg2[lag:]

                                    if len(source_lagged) > 1 and len(target_lagged) > 1:
                                        corr_coef = np.corrcoef(source_lagged, target_lagged)[0, 1]
                                        if not np.isnan(corr_coef):
                                            max_corr = max(max_corr, abs(corr_coef))

                            influence_matrix[source_idx, target_idx] = max_corr
                        except Exception as e:
                            print(f"Error computing influence: {e}")

        # Apply threshold to focus on significant influences (top 15% for cleaner visualization)
        if np.any(influence_matrix > 0):
            threshold = np.percentile(influence_matrix[influence_matrix > 0], 85)
            influence_matrix[influence_matrix < threshold] = 0

        # Store influence matrix for this segment
        segment_influences.append({
            'segment_idx': seg_idx,
            'start_time': start_idx,
            'mid_time': mid_idx,
            'end_time': end_idx,
            'influence': influence_matrix
        })

    # Create Sankey diagram nodes
    nodes = []
    node_labels = []
    node_colors = []

    # Map (segment, electrode) to node index
    node_map = {}
    node_counter = 0

    # Create improved color scale - using Viridis for better distinction
    colors = px.colors.sequential.Viridis

    # First, calculate overall electrode importance
    electrode_importance = np.zeros(num_electrodes)
    for segment in segment_influences:
        infl_mat = segment['influence']
        out_influence = np.sum(infl_mat, axis=1)  # Sum across rows (outgoing)
        in_influence = np.sum(infl_mat, axis=0)   # Sum across columns (incoming)
        electrode_importance += out_influence + in_influence

    # Normalize importance
    if np.max(electrode_importance) > 0:
        electrode_importance = electrode_importance / np.max(electrode_importance)

    # Create nodes for each electrode in each time segment
    for segment_idx in range(n_segments):
        segment_start = segment_idx * segment_size
        segment_end = segment_start + segment_size
        segment_end = min(segment_end, context_size)

        # For electrodes with sufficient importance to reduce clutter
        for e_idx in range(num_electrodes):
            # Skip electrodes with very low importance to reduce clutter
            if electrode_importance[e_idx] < 0.05 and segment_idx > 0:
                continue

            # Create node
            node_map[(segment_idx, e_idx)] = node_counter

            # Add node
            node_labels.append(f"E{e_idx} (t{segment_start}-{segment_end})")
            color_idx = int(electrode_importance[e_idx] * (len(colors)-1))
            node_colors.append(colors[color_idx])

            node_counter += 1

    # Create links between nodes
    links_source = []
    links_target = []
    links_value = []
    links_color = []

    # Process each segment influence
    for segment_data in segment_influences:
        seg_idx = segment_data['segment_idx']
        influence_matrix = segment_data['influence']

        # For each source electrode
        for source_e in range(num_electrodes):
            # For each target electrode
            for target_e in range(num_electrodes):
                if source_e != target_e and influence_matrix[source_e, target_e] > 0:
                    # Check if nodes exist in the map
                    if (seg_idx, source_e) in node_map and (seg_idx + 1, target_e) in node_map:
                        source_node = node_map[(seg_idx, source_e)]
                        target_node = node_map[(seg_idx + 1, target_e)]

                        links_source.append(source_node)
                        links_target.append(target_node)

                        # Scale link values for better visualization
                        links_value.append(influence_matrix[source_e, target_e] * 100)

                        # Use better color scheme based on source electrode importance
                        color_idx = int(electrode_importance[source_e] * (len(colors)-1))
                        links_color.append(colors[color_idx])

    # Create Sankey diagram if we have links
    if links_source:
        # Create node dictionary for Sankey with improved styling
        node_dict = dict(
            pad=15,
            thickness=20,
            line=dict(color="black", width=0.5),
            label=node_labels,
            color=node_colors,
        )

        # Create link dictionary for Sankey
        link_dict = dict(
            source=links_source,
            target=links_target,
            value=links_value,
            color=links_color,
            hoverinfo='all',
            hoverlabel=dict(
                bgcolor='rgba(255, 255, 255, 0.9)',
                bordercolor='#e0e0e0',
                font=dict(size=12, color='#333333')
            )
        )

        # Create figure
        fig = go.Figure(data=[go.Sankey(
            arrangement='snap',
            valueformat=".2f",
            valuesuffix=" units",
            node=node_dict,
            link=link_dict
        )])

        # Update layout with improved aesthetics
        fig.update_layout(
            title=dict(
                text=title,
                y=0.98,
                x=0.5,
                xanchor='center',
                yanchor='top',
                font=dict(family="Arial", size=24, color="#333333")
            ),
            font_size=12,
            width=1400,
            height=900,
            paper_bgcolor='rgb(248, 249, 250)',
            margin=dict(l=25, r=25, t=60, b=25),
        )

        # Add explanation annotation
        fig.add_annotation(
            x=0.01,
            y=0.01,
            xref="paper",
            yref="paper",
            text="This diagram shows how information flows between electrodes over time.<br>" +
                 "• Each node is an electrode at a specific time segment<br>" +
                 "• Links show how electrodes influence each other across time<br>" +
                 "• Node color represents the electrode's overall importance<br>" +
                 "• Link width shows influence strength",
            showarrow=False,
            align="left",
            bgcolor="rgba(255, 255, 255, 0.8)",
            bordercolor="#e0e0e0",
            borderwidth=1,
            borderpad=4,
            font=dict(size=12)
        )

        # Save figure if path provided
        if save_path:
            pio.write_html(fig, save_path)
            print(f"Causal flow diagram saved to {save_path}")

        return fig

    else:
        print("No significant links found for Sankey diagram.")
        return None


def visualize_context_vector_evolution(model, inputs_dict, device, sample_idx=0, save_path=None,
                                       title="Unified Context Vector Evolution Over Time"):
    """
    Create a visualization showing how the electrode signals evolve in a unified space over time.
    """
    print("Generating unified context vector visualization...")

    # Set model to evaluation mode
    model.eval()

    with torch.no_grad():
        # Run model forward to get reservoir states
        inputs = {k: v.to(device) for k, v in inputs_dict.items()}

        # Run model with states returned - this ensures we get the actual model states
        _, state_history = model(inputs, return_states=True)

        # Process states - we'll use population states
        if not state_history:
            print("No state history returned from model. Using raw inputs instead.")
            # Fallback to using raw inputs
            raw_inputs = inputs_dict['raw'].to(device)
            batch_size, context_size, num_electrodes = raw_inputs.shape
            states_np = raw_inputs[sample_idx].cpu().numpy()
        else:
            # Extract the population states from state_history
            # Each element in state_history is a list of population states at a time step
            # We'll concatenate these to get the trajectory
            states = []
            for t, population_states in enumerate(state_history):
                # Concatenate all population states at this time step
                time_state = torch.cat(population_states, dim=1)[sample_idx]
                states.append(time_state.cpu().numpy())

            if not states:
                raise ValueError("No states were extracted from the model")

            states_np = np.array(states)

    # We'll use PCA to reduce the high-dimensional states to 3D
    scaler = StandardScaler()
    states_scaled = scaler.fit_transform(states_np)

    pca = PCA(n_components=3)
    trajectory_3d = pca.fit_transform(states_scaled)

    # Calculate variance explained
    variance_explained = np.sum(pca.explained_variance_ratio_) * 100

    # Analysis of trajectory dynamics
    # 1. Calculate velocity (rate of change)
    velocities = np.zeros(len(trajectory_3d) - 1)
    for t in range(len(trajectory_3d) - 1):
        velocities[t] = np.linalg.norm(trajectory_3d[t+1] - trajectory_3d[t])

    # 2. Find points with significant changes (for highlighting)
    # Use Gaussian smoothing to identify significant deviations
    smoothed_vel = gaussian_filter1d(velocities, sigma=1)
    velocity_diff = np.abs(velocities - smoothed_vel)

    # Find turning points - significant deviations from the smoothed curve
    threshold = np.percentile(velocity_diff, 90)  # Top 10% considered significant
    turning_points = np.where(velocity_diff > threshold)[0]

    # Limit to max 5 turning points to avoid clutter
    if len(turning_points) > 5:
        # Select the most significant turning points
        turning_points = turning_points[np.argsort(velocity_diff[turning_points])[-5:]]

    # Add 1 to turning_points indices (they were calculated on velocities, which are offset by 1)
    turning_points = turning_points + 1

    # Create figure for 3D trajectory visualization with improved aesthetics
    fig = go.Figure()

    # Optimized color scale for time progression
    color_scale = px.colors.sequential.Plasma

    # Create hover text with information
    hover_texts = [f"Time step: {t}<br>PCA Coordinates: ({x:.2f}, {y:.2f}, {z:.2f})"
                  for t, (x, y, z) in enumerate(trajectory_3d)]

    # Add main trajectory line with improved color gradient for time
    fig.add_trace(go.Scatter3d(
        x=trajectory_3d[:, 0],
        y=trajectory_3d[:, 1],
        z=trajectory_3d[:, 2],
        mode='lines',
        line=dict(
            color=np.arange(len(trajectory_3d)),
            colorscale=color_scale,
            width=5
        ),
        opacity=0.8,
        name='Context Trajectory',
        hoverinfo='text',
        hovertext=hover_texts,
        showlegend=True
    ))

    # Add time progression markers with better visibility
    marker_step = max(1, len(trajectory_3d) // 12)  # Show about 12 points for clearer progression

    fig.add_trace(go.Scatter3d(
        x=trajectory_3d[::marker_step, 0],
        y=trajectory_3d[::marker_step, 1],
        z=trajectory_3d[::marker_step, 2],
        mode='markers',
        marker=dict(
            size=6,
            color=np.arange(0, len(trajectory_3d), marker_step),
            colorscale=color_scale,
            line=dict(width=1, color='rgb(40,40,40)')
        ),
        name='Time Progression',
        hoverinfo='text',
        hovertext=[hover_texts[i] for i in range(0, len(hover_texts), marker_step)],
        showlegend=True
    ))

    # Add turning points (significant shifts) as highlighted markers
    if len(turning_points) > 0:
        fig.add_trace(go.Scatter3d(
            x=trajectory_3d[turning_points, 0],
            y=trajectory_3d[turning_points, 1],
            z=trajectory_3d[turning_points, 2],
            mode='markers',
            marker=dict(
                size=10,
                color='yellow',
                symbol='diamond',
                line=dict(width=1.5, color='black')
            ),
            name='Major Shifts',
            hoverinfo='text',
            hovertext=[f"Major shift at t={t}<br>Velocity change: {velocity_diff[t-1]:.3f}"
                      for t in turning_points],
            showlegend=True
        ))

    # Add start and end points
    fig.add_trace(go.Scatter3d(
        x=[trajectory_3d[0, 0]],
        y=[trajectory_3d[0, 1]],
        z=[trajectory_3d[0, 2]],
        mode='markers',
        marker=dict(
            size=10,
            color='green',
            symbol='diamond',
            line=dict(width=1.5, color='black')
        ),
        name='Start point',
        hoverinfo='text',
        hovertext='Initial state (t=0)',
        showlegend=True
    ))

    fig.add_trace(go.Scatter3d(
        x=[trajectory_3d[-1, 0]],
        y=[trajectory_3d[-1, 1]],
        z=[trajectory_3d[-1, 2]],
        mode='markers',
        marker=dict(
            size=10,
            color='red',
            symbol='diamond',
            line=dict(width=1.5, color='black')
        ),
        name='End point',
        hoverinfo='text',
        hovertext=f'Final state (t={len(trajectory_3d)-1})',
        showlegend=True
    ))

    # Update layout with improved aesthetics
    fig.update_layout(
        title=dict(
            text=title,
            y=0.97,
            x=0.5,
            xanchor='center',
            yanchor='top',
            font=dict(family="Arial", size=24, color="#333333")
        ),
        scene=dict(
            xaxis=dict(
                title="PCA Component 1",
                backgroundcolor="rgba(255, 255, 255, 0.95)",
                gridcolor="#e0e0e0",
                showbackground=True,
                zerolinecolor="#cccccc",
            ),
            yaxis=dict(
                title="PCA Component 2",
                backgroundcolor="rgba(255, 255, 255, 0.95)",
                gridcolor="#e0e0e0",
                showbackground=True,
                zerolinecolor="#cccccc",
            ),
            zaxis=dict(
                title="PCA Component 3",
                backgroundcolor="rgba(255, 255, 255, 0.95)",
                gridcolor="#e0e0e0",
                showbackground=True,
                zerolinecolor="#cccccc",
            ),
            camera=dict(
                eye=dict(x=1.5, y=1.5, z=1.2)
            ),
        ),
        margin=dict(l=0, r=0, t=50, b=0),
        width=1200,
        height=800,
        paper_bgcolor="#f8f9fa",
        legend=dict(
            yanchor="top",
            y=0.99,
            xanchor="right",
            x=0.99,
            itemsizing='constant',
            bgcolor="rgba(255, 255, 255, 0.8)",
            bordercolor="#e0e0e0",
            borderwidth=1
        )
    )

    # Add variance explained annotation
    fig.add_annotation(
        x=0.1,
        y=0.95,
        xref="paper",
        yref="paper",
        text=f"Variance explained: {variance_explained:.1f}%",
        showarrow=False,
        font=dict(size=14, color="#333333"),
        align="left",
        bgcolor="rgba(255, 255, 255, 0.8)",
        bordercolor="#e0e0e0",
        borderwidth=1,
        borderpad=4
    )

    # Add explanation annotation
    fig.add_annotation(
        x=0.02,
        y=0.02,
        xref="paper",
        yref="paper",
        text="This visualization shows how the unified electrode activity evolves over time in PCA space.<br>" +
             "• Color gradient represents time progression<br>" +
             "• Yellow diamonds highlight major shifts in trajectory<br>" +
             "• The trajectory reveals how the brain activity forms patterns<br>" +
             "• PCA components capture the principal directions of variation",
        showarrow=False,
        font=dict(size=12, color="#555555"),
        align="left",
        bgcolor="rgba(255, 255, 255, 0.8)",
        bordercolor="#e0e0e0",
        borderwidth=1,
        borderpad=4,
        opacity=0.9
    )

    # Save figure if path provided
    if save_path:
        pio.write_html(fig, save_path)
        print(f"Context vector evolution saved to {save_path}")

    return fig


def visualize_electrode_contributions_to_shifts(model, inputs_dict, device, sample_idx=0, save_path=None,
                                             title="Electrode Contributions to Context Vector Shifts"):
    """
    Create a visualization showing which electrodes contribute to major shifts in the unified context vector.
    This is a new visualization specifically addressing the request to identify which electrodes drive trajectory shifts.
    """
    print("Analyzing electrode contributions to major shifts in context trajectory...")

    # Set model to evaluation mode
    model.eval()

    with torch.no_grad():
        # Extract raw electrode signals
        raw_inputs = inputs_dict['raw'].to(device)  # Shape: [batch, context_size, num_electrodes]
        batch_size, context_size, num_electrodes = raw_inputs.shape

        # Get electrode signals for the selected sample
        electrode_signals = raw_inputs[sample_idx].cpu().numpy()  # Shape: [context_size, num_electrodes]

        # Run model to get state history
        inputs = {k: v.to(device) for k, v in inputs_dict.items()}
        _, state_history = model(inputs, return_states=True)

        if not state_history:
            print("No state history returned. Falling back to electrode signals.")
            states_np = electrode_signals
        else:
            # Extract states from the model's state history
            states = []
            for t, population_states in enumerate(state_history):
                time_state = torch.cat(population_states, dim=1)[sample_idx]
                states.append(time_state.cpu().numpy())

            if not states:
                print("State history was empty. Falling back to electrode signals.")
                states_np = electrode_signals
            else:
                states_np = np.array(states)

    # Apply PCA to the states to get the main trajectory
    scaler = StandardScaler()
    scaled_states = scaler.fit_transform(states_np)

    pca = PCA(n_components=3)
    trajectory_3d = pca.fit_transform(scaled_states)

    # Find points with significant changes
    # 1. Calculate trajectory velocity (distance between consecutive points)
    velocities = np.zeros(len(trajectory_3d) - 1)
    for t in range(len(trajectory_3d) - 1):
        velocities[t] = np.linalg.norm(trajectory_3d[t+1] - trajectory_3d[t])

    # 2. Calculate acceleration (changes in velocity)
    accelerations = np.zeros(len(velocities) - 1)
    for t in range(len(velocities) - 1):
        accelerations[t] = velocities[t+1] - velocities[t]

    # 3. Find points with significant acceleration (trajectory shifts)
    threshold = np.percentile(np.abs(accelerations), 90)  # Top 10% are significant shifts
    shift_indices = np.where(np.abs(accelerations) > threshold)[0]

    # Ensure we have at most 5 major shifts for clarity
    if len(shift_indices) > 5:
        shift_indices = shift_indices[np.argsort(np.abs(accelerations[shift_indices]))[-5:]]

    # Add 1 to shift_indices for trajectory coordinates (accelerations are shifted by 1)
    shift_indices = shift_indices + 1

    # Now analyze which electrodes contributed most to each shift
    shift_contributions = []

    for shift_idx in shift_indices:
        # Look at electrode signals right before the shift
        pre_shift_window = max(0, shift_idx - 3)  # Look at 3 timesteps before the shift

        # Extract the electrode signals in this window
        shift_window_signals = electrode_signals[pre_shift_window:shift_idx + 1]

        # Calculate the rate of change for each electrode in this window
        if len(shift_window_signals) > 1:
            electrode_changes = np.abs(shift_window_signals[-1] - shift_window_signals[0])

            # Normalize the changes
            if np.max(electrode_changes) > 0:
                electrode_changes = electrode_changes / np.max(electrode_changes)

            # Get top 10 electrodes with highest change
            top_electrodes = np.argsort(electrode_changes)[-10:][::-1]
            top_contributions = electrode_changes[top_electrodes]

            shift_contributions.append({
                'shift_idx': shift_idx,
                'top_electrodes': top_electrodes,
                'contributions': top_contributions
            })

    # Create visualization that shows the trajectory and the electrode contributions
    fig = make_subplots(
        rows=2, cols=1,
        specs=[[{"type": "scatter3d"}], [{"type": "bar"}]],
        row_heights=[0.7, 0.3],
        subplot_titles=(
            "Unified Context Vector Trajectory",
            "Top Electrode Contributions to Major Shifts"
        ),
        vertical_spacing=0.05
    )

    # Add trajectory with color gradient by time
    color_scale = px.colors.sequential.Plasma

    # Main trajectory
    fig.add_trace(
        go.Scatter3d(
            x=trajectory_3d[:, 0],
            y=trajectory_3d[:, 1],
            z=trajectory_3d[:, 2],
            mode='lines',
            line=dict(
                color=np.arange(len(trajectory_3d)),
                colorscale=color_scale,
                width=4
            ),
            opacity=0.7,
            name="Context Trajectory",
            showlegend=True
        ),
        row=1, col=1
    )

    # Add shift points to trajectory
    if len(shift_indices) > 0:
        fig.add_trace(
            go.Scatter3d(
                x=trajectory_3d[shift_indices, 0],
                y=trajectory_3d[shift_indices, 1],
                z=trajectory_3d[shift_indices, 2],
                mode='markers',
                marker=dict(
                    size=8,
                    color='yellow',
                    symbol='diamond',
                    line=dict(width=1.5, color='black')
                ),
                name="Major Shifts",
                text=[f"Shift at t={idx}" for idx in shift_indices],
                hoverinfo="text",
                showlegend=True
            ),
            row=1, col=1
        )

    # Add start and end points
    fig.add_trace(
        go.Scatter3d(
            x=[trajectory_3d[0, 0]],
            y=[trajectory_3d[0, 1]],
            z=[trajectory_3d[0, 2]],
            mode='markers',
            marker=dict(
                size=8,
                color='green',
                symbol='diamond',
                line=dict(width=1, color='black')
            ),
            name="Start",
            hoverinfo="name",
            showlegend=True
        ),
        row=1, col=1
    )

    fig.add_trace(
        go.Scatter3d(
            x=[trajectory_3d[-1, 0]],
            y=[trajectory_3d[-1, 1]],
            z=[trajectory_3d[-1, 2]],
            mode='markers',
            marker=dict(
                size=8,
                color='red',
                symbol='diamond',
                line=dict(width=1, color='black')
            ),
            name="End",
            hoverinfo="name",
            showlegend=True
        ),
        row=1, col=1
    )

    # Add bar chart showing electrode contributions for each shift
    if shift_contributions:
        # Process each shift and create a bar chart
        for i, shift_data in enumerate(shift_contributions):
            shift_idx = shift_data['shift_idx']
            top_electrodes = shift_data['top_electrodes']
            contributions = shift_data['contributions']

            # Create bar chart showing top electrode contributions
            bar_colors = px.colors.qualitative.Bold
            color_idx = i % len(bar_colors)

            fig.add_trace(
                go.Bar(
                    x=[f"E{e}" for e in top_electrodes],
                    y=contributions,
                    name=f"Shift at t={shift_idx}",
                    marker_color=bar_colors[color_idx],
                    hoverinfo="text",
                    hovertext=[f"Electrode {e}<br>Contribution: {c:.3f}"
                              for e, c in zip(top_electrodes, contributions)]
                ),
                row=2, col=1
            )

    # Update layout with improved aesthetics
    fig.update_layout(
        title=dict(
            text=title,
            y=0.98,
            x=0.5,
            xanchor='center',
            yanchor='top',
            font=dict(family="Arial", size=24, color="#333333")
        ),
        scene=dict(
            xaxis=dict(
                title="PCA Component 1",
                backgroundcolor="rgba(255, 255, 255, 0.95)",
                gridcolor="#e0e0e0",
                showbackground=True,
                zerolinecolor="#cccccc",
            ),
            yaxis=dict(
                title="PCA Component 2",
                backgroundcolor="rgba(255, 255, 255, 0.95)",
                gridcolor="#e0e0e0",
                showbackground=True,
                zerolinecolor="#cccccc",
            ),
            zaxis=dict(
                title="PCA Component 3",
                backgroundcolor="rgba(255, 255, 255, 0.95)",
                gridcolor="#e0e0e0",
                showbackground=True,
                zerolinecolor="#cccccc",
            ),
            camera=dict(eye=dict(x=1.5, y=1.5, z=1.2))
        ),
        barmode='group',
        xaxis2=dict(
            title="Electrodes",
            tickfont=dict(size=10),
            gridcolor="#e0e0e0",
        ),
        yaxis2=dict(
            title="Contribution Strength",
            tickfont=dict(size=10),
            gridcolor="#e0e0e0",
            range=[0, 1.1]  # Contributions are normalized to [0,1]
        ),
        margin=dict(l=50, r=50, t=60, b=50),
        width=1200,
        height=1000,
        paper_bgcolor='rgb(248, 249, 250)',
        plot_bgcolor='rgb(248, 249, 250)',
        legend=dict(
            yanchor="top",
            y=0.99,
            xanchor="right",
            x=0.99,
            bgcolor="rgba(255, 255, 255, 0.8)",
            bordercolor="#e0e0e0",
            borderwidth=1
        )
    )

    # Add explanation annotation
    fig.add_annotation(
        x=0.02,
        y=0.02,
        xref="paper",
        yref="paper",
        text="This visualization identifies which electrodes contribute to major shifts in the trajectory.<br>" +
             "• Yellow diamonds mark points where the trajectory makes significant turns<br>" +
             "• The bar chart shows which electrodes had the most activity change before each shift<br>" +
             "• Higher bars indicate stronger electrode influence on trajectory changes",
        showarrow=False,
        font=dict(size=12, color="#555555"),
        align="left",
        bgcolor="rgba(255, 255, 255, 0.8)",
        bordercolor="#e0e0e0",
        borderwidth=1,
        borderpad=4,
        opacity=0.9
    )

    # Save figure if path provided
    if save_path:
        pio.write_html(fig, save_path)
        print(f"Electrode contribution analysis saved to {save_path}")

    return fig

def analyze_electrode_contributions(model, inputs_dict, device, sample_idx=0, save_path=None,
                                   title="Electrode Hidden State Contributions to Unified State"):
    """
    Analyzes and visualizes how each electrode's hidden state contributes to the unified hidden state,
    identifying inhibitory and excitatory patterns.
    """
    import numpy as np
    from sklearn.linear_model import ElasticNet
    import plotly.graph_objects as go
    from plotly.subplots import make_subplots
    from sklearn.tree import DecisionTreeRegressor
    from sklearn.tree import export_text

    print("Analyzing electrode hidden state contributions to unified state...")

    # Set model to evaluation mode
    model.eval()

    with torch.no_grad():
        # Get raw inputs and run model forward pass
        raw_inputs = inputs_dict['raw'].to(device)
        batch_size, context_size, num_electrodes = raw_inputs.shape

        # Get model predictions and hidden states
        outputs, state_history = model(inputs_dict, return_states=True)

        if not state_history or len(state_history) == 0:
            print("No state history returned by model. Cannot analyze contributions.")
            return None

        # Extract electrode signals
        electrode_signals = raw_inputs[sample_idx].cpu().numpy()  # Shape: [context_size, num_electrodes]

        # Create a representation of the unified hidden state at each timestep
        unified_states = []
        for t in range(len(state_history)):
            # Combine all population states for this timestep
            population_states = state_history[t]
            if population_states:
                # Concatenate all populations for unified state
                unified_state = torch.cat(population_states, dim=1)[sample_idx].cpu().numpy()
                unified_states.append(unified_state)

        if not unified_states:
            print("Could not extract unified states.")
            return None

        unified_states = np.array(unified_states)

        # For each timestep, compute the contribution of each electrode to the unified state
        # We'll use a regression approach to estimate contributions
        contributions = np.zeros((len(unified_states), num_electrodes))

        # FIXED: Use a simpler approach that doesn't require per-timestep regression
        # Instead, we'll measure correlation and covariance between electrode activity and state activity

        # Calculate the overall contribution of each electrode across all time
        for e_idx in range(num_electrodes):
            # Get electrode signal
            electrode_signal = electrode_signals[:, e_idx]

            # Compute contribution for each timestep based on correlation with state components
            for t in range(len(unified_states)):
                if t < len(electrode_signals):
                    # Calculate a rolling correlation between electrode activity and unified state
                    window_size = min(10, t+1)  # Use up to 10 previous timepoints

                    if window_size > 1:
                        # Get window of electrode activity
                        e_window = electrode_signals[max(0, t-window_size+1):t+1, e_idx]

                        # Get mean of unified state components as a simplified measure
                        state_activity = np.mean(unified_states[max(0, t-window_size+1):t+1], axis=1)

                        if len(e_window) == len(state_activity) and len(e_window) > 1:
                            # Calculate correlation between electrode and state
                            try:
                                corr = np.corrcoef(e_window, state_activity)[0, 1]
                                if not np.isnan(corr):
                                    # Use absolute correlation as contribution measure
                                    contributions[t, e_idx] = abs(corr)
                            except:
                                pass

                        # If correlation failed, use a simple activity-based metric
                        if contributions[t, e_idx] == 0:
                            # Use normalized activity level as a default contribution metric
                            activity_level = abs(electrode_signal[t]) / (np.max(np.abs(electrode_signals[:, e_idx])) + 1e-10)
                            contributions[t, e_idx] = activity_level

        # Normalize contributions per timestep for better visualization
        for t in range(len(contributions)):
            row_sum = np.sum(contributions[t, :])
            if row_sum > 0:
                contributions[t, :] = contributions[t, :] / row_sum

        # Calculate whether each electrode is inhibitory or excitatory based on correlation
        inhibitory_excitatory = np.zeros(num_electrodes)

        # Calculate overall correlation between electrode signals and state activity
        state_activity_overall = np.mean(unified_states, axis=1)

        for e_idx in range(num_electrodes):
            electrode_activity = electrode_signals[:min(len(state_activity_overall), len(electrode_signals)), e_idx]
            state_activity_match = state_activity_overall[:len(electrode_activity)]

            if len(electrode_activity) > 10:  # Need enough samples for meaningful correlation
                try:
                    corr = np.corrcoef(electrode_activity, state_activity_match)[0, 1]
                    if not np.isnan(corr):
                        inhibitory_excitatory[e_idx] = corr
                except:
                    # Fallback for correlation calculation errors
                    pass

        # Identify rule patterns: Extract simple decision rules for when electrodes contribute more
        significant_electrodes = []
        rules = []

        # Calculate mean contributions for each electrode
        mean_contributions = np.mean(contributions, axis=0)

        # For top contributing electrodes
        top_electrodes_idx = np.argsort(mean_contributions)[-5:]  # Top 5 contributors

        # Just generate simple rules directly without fitting a decision tree
        for e_idx in top_electrodes_idx:
            # Create a simple rule based on when this electrode contributes the most
            high_contrib_mask = contributions[:, e_idx] > np.percentile(contributions[:, e_idx], 75)

            if np.any(high_contrib_mask):
                # Find patterns in the electrode signals when contribution is high
                high_signal = np.mean(electrode_signals[high_contrib_mask, e_idx])
                low_signal = np.mean(electrode_signals[~high_contrib_mask, e_idx])

                # Create a simple rule text
                if high_signal > low_signal:
                    rule = f"Electrode {e_idx} contributes most when:\n"
                    rule += f"  Signal amplitude > {np.percentile(electrode_signals[:, e_idx], 50):.6f}\n"
                    rule += f"  Average high contribution signal: {high_signal:.6f}\n"
                    rule += f"  Average low contribution signal: {low_signal:.6f}\n"
                    rule += f"  Importance score: {mean_contributions[e_idx]:.6f}\n"
                else:
                    rule = f"Electrode {e_idx} contributes most when:\n"
                    rule += f"  Signal amplitude < {np.percentile(electrode_signals[:, e_idx], 50):.6f}\n"
                    rule += f"  Average high contribution signal: {high_signal:.6f}\n"
                    rule += f"  Average low contribution signal: {low_signal:.6f}\n"
                    rule += f"  Importance score: {mean_contributions[e_idx]:.6f}\n"

                significant_electrodes.append(e_idx)
                rules.append(rule)

        # Create visualization
        fig = make_subplots(
            rows=3, cols=1,
            row_heights=[0.5, 0.3, 0.2],
            subplot_titles=("Electrode Contributions to Unified State",
                          "Inhibitory vs. Excitatory Effect",
                          "Top Contributing Electrodes")
        )

        # Plot 1: Heatmap of contributions over time
        fig.add_trace(
            go.Heatmap(
                z=contributions,
                x=[f'E{i}' for i in range(num_electrodes)],
                y=list(range(len(unified_states))),
                colorscale='Viridis',
                colorbar=dict(title="Contribution"),
                hoverinfo="text",
                hovertext=[[f"Time: {t}<br>Electrode: {e}<br>Contribution: {contributions[t, e]:.4f}"
                         for e in range(num_electrodes)] for t in range(len(unified_states))]
            ),
            row=1, col=1
        )

        # Plot 2: Bar chart of inhibitory vs. excitatory effect
        fig.add_trace(
            go.Bar(
                x=[f'E{i}' for i in range(num_electrodes)],
                y=inhibitory_excitatory,
                marker=dict(
                    color=inhibitory_excitatory,
                    colorscale='RdBu',
                    cmin=-1,
                    cmid=0,
                    cmax=1,
                    colorbar=dict(title="Effect")
                ),
                hoverinfo="text",
                hovertext=[f"Electrode: E{i}<br>Effect: {'Excitatory' if inhibitory_excitatory[i] > 0 else 'Inhibitory'}" +
                          f"<br>Strength: {abs(inhibitory_excitatory[i]):.4f}"
                          for i in range(num_electrodes)]
            ),
            row=2, col=1
        )

        # Plot 3: Top contributing electrodes
        if len(top_electrodes_idx) > 0:
            top_mean_contributions = np.mean(contributions, axis=0)[top_electrodes_idx]

            fig.add_trace(
                go.Bar(
                    x=[f'E{i}' for i in top_electrodes_idx],
                    y=top_mean_contributions,
                    marker=dict(color='rgba(44, 160, 44, 0.8)'),
                    hoverinfo="text",
                    hovertext=[f"Electrode: E{i}<br>Average Contribution: {cont:.4f}"
                              for i, cont in zip(top_electrodes_idx, top_mean_contributions)]
                ),
                row=3, col=1
            )

        # Update layout
        fig.update_layout(
            title=dict(
                text=title,
                y=0.95,
                x=0.5,
                xanchor="center",
                yanchor="top",
                font=dict(family="Arial", size=24, color="#333333")
            ),
            width=1200,
            height=1000,
            paper_bgcolor="rgb(248, 249, 250)",
            plot_bgcolor="rgb(248, 249, 250)",
            margin=dict(l=60, r=60, t=120, b=60)
        )

        # Add rule annotations
        for i, (e_idx, rule) in enumerate(zip(significant_electrodes, rules)):
            # Add rule as annotation
            fig.add_annotation(
                x=1.1,
                y=0.9 - i*0.15,
                xref="paper",
                yref="paper",
                text=f"<b>Rules for E{e_idx}:</b><br>" + rule.replace('\n', '<br>'),
                showarrow=False,
                font=dict(size=10),
                align="left",
                bgcolor="rgba(255, 255, 255, 0.8)",
                bordercolor="#e0e0e0",
                borderwidth=1,
                borderpad=4
            )

        # Add explanation annotations
        fig.add_annotation(
            x=0.01,
            y=0.01,
            xref="paper",
            yref="paper",
            text=(
                "This visualization shows how each electrode's hidden state contributes to the unified hidden state.<br>" +
                "• <b>Top panel:</b> Heatmap showing electrode contributions over time<br>" +
                "• <b>Middle panel:</b> Bar chart showing inhibitory (negative) vs. excitatory (positive) effects<br>" +
                "• <b>Bottom panel:</b> Top contributing electrodes based on average contribution<br>" +
                "• <b>Right panel:</b> Electrode behavior rules explaining when electrodes contribute most"
            ),
            showarrow=False,
            align="left",
            bgcolor="rgba(255, 255, 255, 0.8)",
            bordercolor="#e0e0e0",
            borderwidth=1,
            borderpad=4,
            font=dict(size=12)
        )

        # Save figure if path provided
        if save_path:
            import plotly.io as pio
            pio.write_html(fig, save_path)
            print(f"Electrode contribution analysis saved to {save_path}")

        return fig


def create_comparative_trajectory_analysis(model, inputs_dict, device, sample_idx=0, save_path=None,
                                        title="Comparative Electrode Trajectory Analysis"):
    """
    Create a visualization that directly compares the trajectories of the most influential electrodes,
    with focus on how they influence the unified context vector.
    """
    print("Creating comparative electrode trajectory analysis...")

    # Set model to evaluation mode
    model.eval()

    with torch.no_grad():
        # Get raw electrode signals
        raw_inputs = inputs_dict['raw'].to(device)  # Shape: [batch, context_size, num_electrodes]
        batch_size, context_size, num_electrodes = raw_inputs.shape

        # Extract electrode signals for the selected sample
        electrode_signals = raw_inputs[sample_idx].cpu().numpy()  # Shape: [context_size, num_electrodes]

        # Run model to get reservoir states
        inputs = {k: v.to(device) for k, v in inputs_dict.items()}
        _, state_history = model(inputs, return_states=True)

        # Extract the unified context vector (concatenated population states)
        if state_history:
            unified_states = []
            for t, population_states in enumerate(state_history):
                unified_state = torch.cat(population_states, dim=1)[sample_idx]
                unified_states.append(unified_state.cpu().numpy())
            unified_trajectory = np.array(unified_states)
        else:
            print("No state history returned. Using raw inputs for trajectory.")
            unified_trajectory = electrode_signals

    # Apply PCA to the unified trajectory
    unified_scaler = StandardScaler()
    unified_scaled = unified_scaler.fit_transform(unified_trajectory)

    unified_pca = PCA(n_components=3)
    unified_3d = unified_pca.fit_transform(unified_scaled)

    # Calculate the velocity (changes) in the unified trajectory
    unified_velocity = np.zeros(len(unified_3d) - 1)
    for t in range(len(unified_3d) - 1):
        unified_velocity[t] = np.linalg.norm(unified_3d[t+1] - unified_3d[t])

    # Find major shift points - significant velocity changes
    shift_points = []
    velocity_smooth = gaussian_filter1d(unified_velocity, sigma=1)
    velocity_diff = np.abs(unified_velocity - velocity_smooth)

    # Identify points with significant velocity changes
    threshold = np.percentile(velocity_diff, 92)  # Top 8% considered significant
    shift_candidates = np.where(velocity_diff > threshold)[0]

    # Select the top 3 most significant shift points
    if len(shift_candidates) > 0:
        top_shifts = shift_candidates[np.argsort(velocity_diff[shift_candidates])[-3:]]
        shift_points = top_shifts + 1  # +1 because velocity is calculated between points

    # Calculate the influence of each electrode
    electrode_influence = np.zeros(num_electrodes)

    # Method 1: Calculate using correlation with unified trajectory velocity
    electrode_velocities = np.zeros((context_size-1, num_electrodes))
    for e in range(num_electrodes):
        for t in range(context_size-1):
            electrode_velocities[t, e] = np.abs(electrode_signals[t+1, e] - electrode_signals[t, e])

    # Compute correlation between each electrode's velocity and unified velocity
    for e in range(num_electrodes):
        if np.std(electrode_velocities[:, e]) > 0 and np.std(unified_velocity) > 0:
            corr = np.corrcoef(electrode_velocities[:, e], unified_velocity)[0, 1]
            if not np.isnan(corr):
                electrode_influence[e] = abs(corr)  # Use absolute correlation as influence measure

    # Method 2: Calculate influence around shift points
    if len(shift_points) > 0:
        for shift_idx in shift_points:
            pre_idx = max(0, shift_idx - 2)
            # Calculate which electrodes changed most before the shift
            shift_window = electrode_signals[pre_idx:shift_idx+1]
            if len(shift_window) > 1:
                electrode_changes = np.abs(shift_window[-1] - shift_window[0])
                normalized_changes = electrode_changes / np.max(electrode_changes) if np.max(electrode_changes) > 0 else 0
                # Add to electrode influence
                electrode_influence += normalized_changes

    # Normalize influence
    if np.max(electrode_influence) > 0:
        electrode_influence = electrode_influence / np.max(electrode_influence)

    # Get top influential electrodes
    top_n = min(8, num_electrodes)  # Select top 8 electrodes for visualization
    top_electrodes = np.argsort(electrode_influence)[-top_n:][::-1]

    # Create 3D trajectories for top electrodes using delay embedding
    electrode_trajectories = []
    delay = max(1, context_size // 25)  # Delay for embedding

    for e_idx in top_electrodes:
        if context_size > 3*delay:
            x = electrode_signals[:-2*delay, e_idx]
            y = electrode_signals[delay:-delay, e_idx]
            z = electrode_signals[2*delay:, e_idx]
            electrode_trajectories.append({
                'electrode': e_idx,
                'x': x,
                'y': y,
                'z': z,
                'influence': electrode_influence[e_idx]
            })

    # Create figure with multiple subplots
    fig = make_subplots(
        rows=2, cols=2,
        specs=[
            [{"type": "scatter3d"}, {"type": "scatter3d"}],
            [{"type": "scatter", "colspan": 2}, {}]
        ],
        subplot_titles=(
            "Unified Context Vector Trajectory",
            "Top Electrode Trajectories",
            "Electrode Influence Analysis"
        ),
        column_widths=[0.5, 0.5],
        row_heights=[0.7, 0.3],
        vertical_spacing=0.05,
        horizontal_spacing=0.05
    )

    # 1. Add unified trajectory to first subplot
    # Define better color scale
    import plotly.express as px
    color_scale = px.colors.sequential.Plasma

    # Main unified trajectory
    fig.add_trace(
        go.Scatter3d(
            x=unified_3d[:, 0],
            y=unified_3d[:, 1],
            z=unified_3d[:, 2],
            mode='lines',
            line=dict(
                color=np.arange(len(unified_3d)),
                colorscale=color_scale,
                width=5
            ),
            opacity=0.8,
            name="Context Vector",
            showlegend=True
        ),
        row=1, col=1
    )

    # Add shift points
    if len(shift_points) > 0:
        fig.add_trace(
            go.Scatter3d(
                x=unified_3d[shift_points, 0],
                y=unified_3d[shift_points, 1],
                z=unified_3d[shift_points, 2],
                mode='markers',
                marker=dict(
                    size=8,
                    color='yellow',
                    symbol='diamond',
                    line=dict(width=1, color='black')
                ),
                name="Major Shifts",
                text=[f"Shift at t={t}" for t in shift_points],
                hoverinfo="text",
                showlegend=True
            ),
            row=1, col=1
        )

    # 2. Add electrode trajectories to second subplot
    viridis_colors = px.colors.sequential.Viridis

    for i, traj in enumerate(electrode_trajectories):
        color_idx = min(len(viridis_colors)-1, int(i * len(viridis_colors) / len(electrode_trajectories)))
        e_color = viridis_colors[color_idx]

        fig.add_trace(
            go.Scatter3d(
                x=traj['x'],
                y=traj['y'],
                z=traj['z'],
                mode='lines',
                line=dict(
                    color=e_color,
                    width=4
                ),
                opacity=0.7,
                name=f"E{traj['electrode']}",
                text=[f"Electrode {traj['electrode']}<br>Time: {t}<br>Influence: {traj['influence']:.3f}"
                      for t in range(len(traj['x']))],
                hoverinfo="text",
                showlegend=True
            ),
            row=1, col=2
        )

    # 3. Add influence bar chart to bottom subplot
    fig.add_trace(
        go.Bar(
            x=[f"E{e}" for e in top_electrodes],
            y=[electrode_influence[e] for e in top_electrodes],
            marker=dict(
                color=[viridis_colors[min(len(viridis_colors)-1, int(i * len(viridis_colors) / len(top_electrodes)))]
                       for i in range(len(top_electrodes))],
                line=dict(width=1, color='rgb(8,48,107)')
            ),
            name="Electrode Influence",
            text=[f"Electrode {e}<br>Influence: {electrode_influence[e]:.3f}" for e in top_electrodes],
            hoverinfo="text"
        ),
        row=2, col=1
    )

    # Add velocity analysis line
    if len(unified_velocity) > 0:
        time_points = np.arange(1, len(unified_velocity) + 1)
        # Normalize velocity for plotting
        norm_velocity = unified_velocity / np.max(unified_velocity) if np.max(unified_velocity) > 0 else unified_velocity

        fig.add_trace(
            go.Scatter(
                x=time_points,
                y=norm_velocity,
                mode='lines',
                line=dict(
                    color='rgba(150, 50, 50, 0.7)',
                    width=2,
                    dash='dot'
                ),
                name="Trajectory Velocity",
                yaxis="y2"
            ),
            row=2, col=1
        )

    # Instead of add_vline which doesn't work with Scatter3d, we'll add markers for shift points in the bottom plot
    if len(shift_points) > 0:
        for shift in shift_points:
            if shift < len(time_points):
                fig.add_trace(
                    go.Scatter(
                        x=[shift, shift],
                        y=[0, 1],
                        mode='lines',
                        line=dict(
                            color="yellow",
                            width=2,
                            dash="dash"
                        ),
                        name=f"Shift at t={shift}",
                        showlegend=False
                    ),
                    row=2, col=1
                )

    # Update layout for improved aesthetics
    fig.update_layout(
        title=dict(
            text=title,
            y=0.98,
            x=0.5,
            xanchor='center',
            yanchor='top',
            font=dict(family="Arial", size=24, color="#333333")
        ),
        scene=dict(
            xaxis=dict(
                title="Component 1",
                backgroundcolor="rgba(255, 255, 255, 0.95)",
                gridcolor="#e0e0e0",
                showbackground=True,
                zerolinecolor="#cccccc",
            ),
            yaxis=dict(
                title="Component 2",
                backgroundcolor="rgba(255, 255, 255, 0.95)",
                gridcolor="#e0e0e0",
                showbackground=True,
                zerolinecolor="#cccccc",
            ),
            zaxis=dict(
                title="Component 3",
                backgroundcolor="rgba(255, 255, 255, 0.95)",
                gridcolor="#e0e0e0",
                showbackground=True,
                zerolinecolor="#cccccc",
            ),
            camera=dict(eye=dict(x=1.5, y=1.5, z=1.2))
        ),
        scene2=dict(
            xaxis=dict(
                title="t",
                backgroundcolor="rgba(255, 255, 255, 0.95)",
                gridcolor="#e0e0e0",
                showbackground=True,
                zerolinecolor="#cccccc",
            ),
            yaxis=dict(
                title="t+delay",
                backgroundcolor="rgba(255, 255, 255, 0.95)",
                gridcolor="#e0e0e0",
                showbackground=True,
                zerolinecolor="#cccccc",
            ),
            zaxis=dict(
                title="t+2*delay",
                backgroundcolor="rgba(255, 255, 255, 0.95)",
                gridcolor="#e0e0e0",
                showbackground=True,
                zerolinecolor="#cccccc",
            ),
            camera=dict(eye=dict(x=1.5, y=1.5, z=1.2))
        ),
        xaxis=dict(
            title="Electrodes",
            tickfont=dict(size=10),
            gridcolor="#e0e0e0",
        ),
        yaxis=dict(
            title="Influence Strength",
            tickfont=dict(size=10),
            gridcolor="#e0e0e0",
            range=[0, 1.1]
        ),
        yaxis2=dict(
            title="Trajectory Velocity",
            overlaying="y",
            side="right",
            tickfont=dict(size=10),
            gridcolor="#e0e0e0",
            range=[0, 1.1]
        ),
        showlegend=True,
        legend=dict(
            yanchor="top",
            y=0.99,
            xanchor="right",
            x=0.99,
            font=dict(size=10),
            bgcolor="rgba(255, 255, 255, 0.8)",
            bordercolor="#e0e0e0",
            borderwidth=1
        ),
        margin=dict(l=60, r=60, t=100, b=60),
        width=1400,
        height=900,
        paper_bgcolor='rgb(248, 249, 250)',
        plot_bgcolor='rgb(248, 249, 250)',
    )

    # Add explanation annotations
    fig.add_annotation(
        x=0.02,
        y=0.02,
        xref="paper",
        yref="paper",
        text="This analysis compares how individual electrodes influence the unified context vector trajectory.<br>" +
             "• Yellow diamonds mark major shifts in the main trajectory<br>" +
             "• The most influential electrodes have their own trajectories shown (right)<br>" +
             "• The bar chart shows relative influence of each top electrode<br>" +
             "• Dotted red line shows velocity of the main trajectory<br>" +
             "• Yellow vertical lines mark points of major shifts",
        showarrow=False,
        font=dict(size=12, color="#555555"),
        align="left",
        bgcolor="rgba(255, 255, 255, 0.8)",
        bordercolor="#e0e0e0",
        borderwidth=1,
        borderpad=4,
        opacity=0.9
    )

    # Save figure if path provided
    if save_path:
        pio.write_html(fig, save_path)
        print(f"Comparative trajectory analysis saved to {save_path}")

    return fig


def train_epoch(model, train_loader, optimizer, criterion, scheduler, device, config, epoch, clip_grad=None):
    """Enhanced training for one epoch with gradient accumulation and curriculum learning."""
    model.train()
    total_loss_val, time_losses_val, freq_losses_val, phase_losses_val, envelope_losses_val = 0.0, 0.0, 0.0, 0.0, 0.0


    if config.USE_CURRICULUM:
        curriculum = CurriculumScheduler(config) # Re-init or pass as arg if stateful across epochs
        curriculum.update(epoch)
        loss_weights = curriculum.get_loss_weights()
        criterion.time_weight = loss_weights['time']
        criterion.freq_weight = loss_weights['freq']
        criterion.phase_weight = loss_weights['phase']
        criterion.envelope_weight = loss_weights['envelope']
        criterion.gamma_band_weights = curriculum.get_gamma_weights()

    if epoch < config.LR_WARMUP_EPOCHS and config.LR_WARMUP_EPOCHS > 0: # Avoid division by zero
        warmup_factor = (epoch + 1) / config.LR_WARMUP_EPOCHS
        for param_group in optimizer.param_groups:
            param_group['lr'] = config.LEARNING_RATE * warmup_factor

    progress_bar = tqdm(train_loader, desc=f"Training Epoch {epoch+1}")
    optimizer.zero_grad() # Initialize gradients

    for batch_idx, batch_data in enumerate(progress_bar):
        try:
            inputs, targets_dict = batch_data
            inputs = {k: v.to(device) for k, v in inputs.items()}
            target_signal = targets_dict['target'].to(device)

            outputs = model(inputs)
            loss, time_loss, freq_loss, phase_loss, envelope_loss = criterion(outputs, target_signal)

            if torch.isnan(loss):
                print(f"Warning: NaN loss encountered at epoch {epoch}, batch {batch_idx}. Skipping batch.")
                # Also good to print shapes and stats of outputs and target_signal here for debugging
                # Example: print(f"Outputs: {outputs.shape}, mean: {outputs.mean()}, std: {outputs.std()}")
                # print(f"Target: {target_signal.shape}, mean: {target_signal.mean()}, std: {target_signal.std()}")
                # Consider zeroing gradients again if an optimizer step was planned
                if (batch_idx + 1) % config.GRADIENT_ACCUMULATION_STEPS == 0 or \
                   (batch_idx + 1) == len(train_loader):
                    optimizer.zero_grad() # Ensure grads are zeroed before next accumulation cycle
                continue


            scaled_loss = loss / config.GRADIENT_ACCUMULATION_STEPS
            scaled_loss.backward()

            if (batch_idx + 1) % config.GRADIENT_ACCUMULATION_STEPS == 0 or \
               (batch_idx + 1) == len(train_loader):
                if clip_grad is not None:
                    torch.nn.utils.clip_grad_norm_(model.parameters(), clip_grad)
                optimizer.step()
                optimizer.zero_grad() # Zero gradients after step

            total_loss_val += loss.item()
            time_losses_val += time_loss.item() if torch.is_tensor(time_loss) else time_loss
            freq_losses_val += freq_loss.item() if torch.is_tensor(freq_loss) else freq_loss
            phase_losses_val += phase_loss.item() if torch.is_tensor(phase_loss) else phase_loss
            envelope_losses_val += envelope_loss.item() if torch.is_tensor(envelope_loss) else envelope_loss


            progress_bar.set_postfix({
                'loss': loss.item(),
                'time_l': time_loss.item() if torch.is_tensor(time_loss) else time_loss,
                'freq_l': freq_loss.item() if torch.is_tensor(freq_loss) else freq_loss
            })

        except Exception as e:
            print(f"Error in training batch (epoch {epoch}, batch {batch_idx}): {type(e).__name__} - {e}")
            traceback.print_exc() # FIX 3: Add full traceback
            # Potentially skip this batch or re-throw if critical
            # Ensure gradients are zeroed if an optimizer step was planned
            if (batch_idx + 1) % config.GRADIENT_ACCUMULATION_STEPS == 0 or \
                (batch_idx + 1) == len(train_loader):
                optimizer.zero_grad()
            continue


    num_batches = len(train_loader)
    if num_batches == 0: return {'loss': 0, 'time_loss': 0, 'freq_loss': 0, 'phase_loss': 0, 'envelope_loss': 0}

    return {
        'loss': total_loss_val / num_batches,
        'time_loss': time_losses_val / num_batches,
        'freq_loss': freq_losses_val / num_batches,
        'phase_loss': phase_losses_val / num_batches,
        'envelope_loss': envelope_losses_val / num_batches
    }


def evaluate(model, val_loader, criterion, device, compute_metrics=True):
    """Enhanced evaluation with detailed gamma-specific metrics."""
    model.eval()
    total_loss = 0
    time_losses = 0
    freq_losses = 0
    phase_losses = 0
    envelope_losses = 0

    # Metrics storage with separate gamma band metrics
    gamma_metrics = {band: {'mse': [], 'r2': [], 'corr': []} for band in config.FREQ_BANDS}

    all_metrics = {
        'mse': [],
        'r2': [],
        'corr': [],
        'gamma_metrics': gamma_metrics,
        'spectrum_corr': []  # Correlation in frequency domain
    }

    # Store sample states for visualization
    sample_states = None
    sample_inputs = None
    sample_targets = None
    sample_outputs = None

    with torch.no_grad():
        for batch_idx, batch_data in enumerate(tqdm(val_loader, desc="Evaluating")):
            try:
                # Unpack data
                inputs, targets = batch_data

                # Move to device
                inputs = {k: v.to(device) for k, v in inputs.items()}
                target_signal = targets['target'].to(device)

                # Forward pass with state tracking
                outputs, states = model(inputs, return_states=True)

                # Store one set of samples for visualization
                if sample_states is None:
                    sample_states = states
                    sample_inputs = inputs
                    sample_targets = target_signal.cpu().numpy()
                    sample_outputs = outputs.cpu().numpy()

                # Compute loss
                loss, time_loss, freq_loss, phase_loss, envelope_loss = criterion(outputs, target_signal)

                # Track losses
                total_loss += loss.item()
                time_losses += time_loss.item()
                freq_losses += freq_loss.item()
                phase_losses += phase_loss.item()
                envelope_losses += envelope_loss.item()

                # Compute metrics if requested
                if compute_metrics:
                    # Move to CPU for numpy processing
                    outputs_np = outputs.cpu().numpy()
                    targets_np = target_signal.cpu().numpy()

                    # Get frequency band data for filtering
                    phase_target = targets['phase'].cpu().numpy()
                    amp_target = targets['amplitude'].cpu().numpy()

                    for i in range(outputs_np.shape[0]):
                        # Full signal metrics for limited electrodes
                        for e in range(min(5, outputs_np.shape[2])):
                            # Extract electrode signals
                            pred_signal = outputs_np[i, :, e]
                            true_signal = targets_np[i, :, e]

                            # Calculate full signal metrics
                            try:
                                mse = np.mean((pred_signal - true_signal)**2)
                                all_metrics['mse'].append(mse)

                                r2 = r2_score(true_signal, pred_signal)
                                if not np.isnan(r2):
                                    all_metrics['r2'].append(r2)

                                corr = np.corrcoef(pred_signal, true_signal)[0, 1]
                                if not np.isnan(corr):
                                    all_metrics['corr'].append(corr)

                                # Compute spectrum correlation
                                fs = config.SAMPLING_RATE
                                pred_fft = np.abs(np.fft.rfft(pred_signal))
                                true_fft = np.abs(np.fft.rfft(true_signal))
                                # Focus on gamma range frequencies (30-300 Hz)
                                freq_bins = np.fft.rfftfreq(len(true_signal), 1/fs)
                                gamma_mask = (freq_bins >= 30) & (freq_bins <= 300)
                                pred_fft_gamma = pred_fft[gamma_mask]
                                true_fft_gamma = true_fft[gamma_mask]
                                spec_corr = np.corrcoef(pred_fft_gamma, true_fft_gamma)[0, 1]
                                if not np.isnan(spec_corr):
                                    all_metrics['spectrum_corr'].append(spec_corr)

                            except Exception as e:
                                print(f"Error computing metrics: {e}")

                            # Compute band-specific metrics
                            for band_idx, (low_freq, high_freq) in enumerate(config.FREQ_BANDS):
                                try:
                                    # Extract band-specific components using target amplitude/phase
                                    # Use the ground truth filtered signal as reference
                                    if 'filtered' in inputs:
                                        band_true = inputs['filtered'][i, band_idx, :, e].cpu().numpy()
                                    else:
                                        band_true = true_signal  # Fallback if filtered not available

                                    # Skip filtering for short signals
                                    if len(pred_signal) < 20:  # Skip if signal is too short
                                        continue

                                    # Apply same band-pass filter to predicted signal
                                    nyquist = config.SAMPLING_RATE / 2
                                    low = low_freq / nyquist
                                    high = high_freq / nyquist

                                    # Use a much smaller filter order for short signals
                                    filter_order = min(2, len(pred_signal) // 10)  # Use only order 1 or 2 for short signals
                                    if filter_order < 1:
                                        continue  # Skip if we can't even use order 1

                                    # Design filter with reduced order
                                    sos = signal.butter(filter_order, [low, high], btype='bandpass', output='sos')

                                    # Forward filtering only (no zero-phase) - much less demanding on signal length
                                    band_pred = signal.sosfilt(sos, pred_signal)
                                    band_true_filtered = signal.sosfilt(sos, true_signal)

                                    # Calculate band-specific metrics
                                    band_mse = np.mean((band_pred - band_true_filtered)**2)
                                    gamma_metrics[(low_freq, high_freq)]['mse'].append(band_mse)

                                    band_r2 = r2_score(band_true_filtered, band_pred)
                                    if not np.isnan(band_r2):
                                        gamma_metrics[(low_freq, high_freq)]['r2'].append(band_r2)

                                    band_corr = np.corrcoef(band_pred, band_true_filtered)[0, 1]
                                    if not np.isnan(band_corr):
                                        gamma_metrics[(low_freq, high_freq)]['corr'].append(band_corr)

                                except Exception as exc:
                                    # Use exc instead of e to avoid variable name conflicts
                                    print(f"Error in band-specific metrics: {exc}")

            except Exception as e:
                print(f"Error in evaluation batch: {e}")
                continue

    # Calculate average losses
    num_batches = max(1, len(val_loader))
    avg_loss = total_loss / num_batches
    avg_time_loss = time_losses / num_batches
    avg_freq_loss = freq_losses / num_batches
    avg_phase_loss = phase_losses / num_batches
    avg_envelope_loss = envelope_losses / num_batches

    # Calculate average metrics
    avg_metrics = {
        'loss': avg_loss,
        'time_loss': avg_time_loss,
        'freq_loss': avg_freq_loss,
        'phase_loss': avg_phase_loss,
        'envelope_loss': avg_envelope_loss
    }

    for key in ['mse', 'r2', 'corr', 'spectrum_corr']:
        values = all_metrics.get(key, [])
        if values:
            avg_metrics[key] = np.mean(values)
        else:
            avg_metrics[key] = float('nan')

    # Calculate average band-specific metrics
    band_avg_metrics = {}
    for band, metrics in gamma_metrics.items():
        band_avg_metrics[band] = {}
        for metric_name, values in metrics.items():
            if values:
                band_avg_metrics[band][metric_name] = np.mean(values)
            else:
                band_avg_metrics[band][metric_name] = float('nan')

    avg_metrics['gamma_metrics'] = band_avg_metrics

    # Return also sample data for visualization
    return avg_metrics, sample_states, sample_inputs, sample_targets, sample_outputs

def visualize_predictions_enhanced(model, inputs_dict, targets, outputs=None, electrode_idx=0, save_path=None):
    """Visualize enhanced predictions for a specific electrode."""
    if outputs is None:
        model.eval()
        with torch.no_grad():
            device = next(model.parameters()).device
            device_inputs = {k: v.to(device) for k, v in inputs_dict.items()}
            outputs_model, _ = model(device_inputs, return_states=True) # Assuming model might return states
            outputs = outputs_model.cpu().numpy()

    targets_np = targets if isinstance(targets, np.ndarray) else targets.cpu().numpy()
    context_np = inputs_dict['raw'].cpu().numpy() if torch.is_tensor(inputs_dict['raw']) else inputs_dict['raw']

    sample_idx = 0 # Assuming first sample in batch for visualization
    context = context_np[sample_idx, :, electrode_idx]
    true_future = targets_np[sample_idx, :, electrode_idx]
    predicted_future = outputs[sample_idx, :, electrode_idx]

    if len(true_future) == 0 or len(predicted_future) == 0:
        print(f"Skipping visualization for electrode {electrode_idx} due to empty future/prediction.")
        return {}

    t_context = np.arange(len(context))
    t_future = np.arange(len(context), len(context) + len(true_future))

    plt.figure(figsize=(15, 12))
    plt.subplot(3, 1, 1)
    plt.plot(t_context, context, 'b-', label='Context', alpha=0.7, linewidth=1.5)
    plt.plot(t_future, true_future, 'g-', linewidth=2, label='True Future')
    plt.plot(t_future, predicted_future, 'r-', linewidth=1.5, label='Predicted Future')
    plt.axvline(x=len(context), color='k', linestyle='--', alpha=0.7)
    plt.title(f"Full Signal - Electrode {electrode_idx}", fontsize=14)
    plt.xlabel("Time (ms)"); plt.ylabel("Signal Amplitude"); plt.legend(); plt.grid(alpha=0.3)

    mse = mean_squared_error(true_future, predicted_future)
    r2 = r2_score(true_future, predicted_future) if np.var(true_future) > 1e-9 else np.nan
    corr = np.corrcoef(true_future, predicted_future)[0,1] if np.std(true_future)>1e-9 and np.std(predicted_future)>1e-9 else np.nan
    plt.text(0.02,0.92,f"MSE:{mse:.4f}, R2:{r2:.4f}, Corr:{corr:.4f}",transform=plt.gca().transAxes)

    plt.subplot(3, 1, 2) # Combined Gamma
    nyquist = config.SAMPLING_RATE / 2.0
    low, high = 30 / nyquist, 300 / nyquist
    filter_order = 1 # Use low order for robustness
    sos = signal.butter(filter_order, [low, high], btype='bandpass', output='sos')

    # Use sosfilt for short signals if sosfiltfilt would fail
    min_len_for_filtfilt = 3 * filter_order +1 # Approximation
    if len(true_future) >= min_len_for_filtfilt:
        true_gamma = signal.sosfiltfilt(sos, true_future)
        predicted_gamma = signal.sosfiltfilt(sos, predicted_future)
    else:
        true_gamma = signal.sosfilt(sos, true_future)
        predicted_gamma = signal.sosfilt(sos, predicted_future)

    plt.plot(t_future, true_gamma, 'g-', label='True Gamma')
    plt.plot(t_future, predicted_gamma, 'r-', label='Predicted Gamma')
    plt.title(f"Full Gamma Band (30-300 Hz)"); plt.xlabel("Time (ms)"); plt.ylabel("Filtered Signal"); plt.legend(); plt.grid(alpha=0.3)
    gamma_mse = mean_squared_error(true_gamma, predicted_gamma)
    gamma_r2 = r2_score(true_gamma, predicted_gamma) if np.var(true_gamma) > 1e-9 else np.nan
    gamma_corr = np.corrcoef(true_gamma,predicted_gamma)[0,1] if np.std(true_gamma)>1e-9 and np.std(predicted_gamma)>1e-9 else np.nan
    plt.text(0.02,0.92,f"MSE:{gamma_mse:.4f}, R2:{gamma_r2:.4f}, Corr:{gamma_corr:.4f}",transform=plt.gca().transAxes)

    plt.subplot(3, 1, 3) # Sub-bands
    colors = plt.cm.viridis(np.linspace(0, 1, len(config.FREQ_BANDS)))
    legend_elements = []
    for i, (low_f, high_f) in enumerate(config.FREQ_BANDS):
        low_n, high_n = low_f/nyquist, high_f/nyquist
        sos_band = signal.butter(filter_order, [low_n, high_n], btype='bandpass', output='sos')
        if len(true_future) >= min_len_for_filtfilt:
            band_true = signal.sosfiltfilt(sos_band, true_future)
            band_pred = signal.sosfiltfilt(sos_band, predicted_future)
        else:
            band_true = signal.sosfilt(sos_band, true_future)
            band_pred = signal.sosfilt(sos_band, predicted_future)

        offset = i * (np.max(np.abs(band_true)) if len(band_true)>0 else 0.2) # Dynamic offset
        plt.plot(t_future, band_true + offset, color=colors[i], ls='-',alpha=0.7, label=f'True {low_f}-{high_f} Hz')
        plt.plot(t_future, band_pred + offset, color=colors[i], ls='--', label=f'Pred {low_f}-{high_f} Hz')
        band_corr = np.corrcoef(band_true,band_pred)[0,1] if np.std(band_true)>1e-9 and np.std(band_pred)>1e-9 and len(band_true)>1 else np.nan
        if not np.isnan(band_corr): plt.text(t_future[-1]+2 if len(t_future)>0 else 0, offset, f"r={band_corr:.2f}", color=colors[i])
        legend_elements.append(plt.Line2D([0],[0],color=colors[i],lw=2,label=f'{low_f}-{high_f} Hz'))

    plt.title("Gamma Sub-bands (Solid=True, Dashed=Predicted)"); plt.xlabel("Time (ms)"); plt.ylabel("Band-filtered Signal + Offset");
    plt.legend(handles=legend_elements, loc='upper right', fontsize=8); plt.grid(alpha=0.3)
    plt.tight_layout(); plt.suptitle(f"Enhanced Signal Prediction - Electrode {electrode_idx}",y=0.99)
    plt.subplots_adjust(top=0.93)

    if save_path: plt.savefig(save_path, dpi=150); plt.close()
    else: plt.show()
    return {'mse': mse, 'r2': r2, 'corr': corr, 'gamma_mse': gamma_mse, 'gamma_r2': gamma_r2, 'gamma_corr': gamma_corr}


def run_training_enhanced(model, train_loader, val_loader, config, test_loader=None):
    """Run complete training pipeline with enhanced strategies for gamma oscillation prediction."""
    print("\nStarting enhanced training pipeline for gamma oscillation prediction...")
    criterion = EnhancedGammaLoss(config)
    optimizer = torch.optim.AdamW(
        model.parameters(), lr=config.LEARNING_RATE, weight_decay=config.WEIGHT_DECAY, amsgrad=True
    )
    scheduler = ReduceLROnPlateau(
        optimizer, mode='min', factor=0.5, patience=7, min_lr=1e-6, verbose=True
    )

    swa_model = None # Initialize to None
    if config.USE_SWA:
        swa_model = AveragedModel(model)
        # SWA scheduler should operate on the main optimizer's LR
        # CosineAnnealingLR is often used with SWA, ensure T_max is appropriate (e.g., total epochs after SWA start)
        swa_scheduler_anneal_epochs = config.EPOCHS - config.SWA_START
        if swa_scheduler_anneal_epochs <=0: swa_scheduler_anneal_epochs = 1 # Min 1 epoch for scheduler
        swa_torch_scheduler = torch.optim.lr_scheduler.CosineAnnealingLR(optimizer, T_max=swa_scheduler_anneal_epochs)


    best_val_loss, best_gamma_metric, patience_counter = float('inf'), -float('inf'), 0 # Gamma metric higher is better
    history = {'train_loss': [], 'train_metrics': [], 'val_loss': [], 'val_metrics': [], 'lr': []}

    print(f"\nTraining for {config.EPOCHS} epochs...")
    for epoch in range(config.EPOCHS):
        print(f"\n{'='*80}\nEpoch {epoch+1}/{config.EPOCHS}\n{'='*80}")
        train_metrics = train_epoch(
            model, train_loader, optimizer, criterion, scheduler, device, config, epoch, config.CLIP_GRAD_NORM
        )
        print(f"Training completed - Loss: {train_metrics['loss']:.5f}")

        if config.USE_SWA and epoch >= config.SWA_START:
            swa_model.update_parameters(model)
            if 'swa_torch_scheduler' in locals(): swa_torch_scheduler.step() # Step SWA's LR scheduler

        # Use SWA model for validation if SWA is active and past start epoch
        eval_model = swa_model if config.USE_SWA and epoch >= config.SWA_START else model
        val_metrics, s_states, s_inputs, s_targets, s_outputs = evaluate(
            eval_model, val_loader, criterion, device
        )
        print(f"Validation completed - Loss: {val_metrics['loss']:.5f}")

        current_lr = optimizer.param_groups[0]['lr']
        # Note: scheduler.step is for ReduceLROnPlateau. If SWA uses CosineAnnealingLR, that one steps each SWA epoch.
        # If SWA is active, ReduceLROnPlateau might not be the primary LR control.
        # For simplicity, let ReduceLROnPlateau step on val_loss, but SWA might override LR.
        if not (config.USE_SWA and epoch >= config.SWA_START and 'swa_torch_scheduler' in locals()):
             scheduler.step(val_metrics['loss'])


        # Display training results
        print("\nTraining Results:")
        print(f"  Train Loss: {train_metrics['loss']:.6f}")
        print(f"    Time Loss: {train_metrics['time_loss']:.6f}")
        print(f"    Freq Loss: {train_metrics['freq_loss']:.6f}")
        print(f"    Phase Loss: {train_metrics['phase_loss']:.6f}")
        print(f"    Envelope Loss: {train_metrics['envelope_loss']:.6f}")

        print("\nValidation Results:")
        print(f"  Val Loss: {val_metrics['loss']:.6f}")
        print(f"    Time Loss: {val_metrics['time_loss']:.6f}")
        print(f"    Freq Loss: {val_metrics['freq_loss']:.6f}")
        print(f"  Full Signal - MSE: {val_metrics['mse']:.6f}, R²: {val_metrics['r2']:.6f}, Corr: {val_metrics['corr']:.6f}")
        print(f"  Spectrum Correlation: {val_metrics['spectrum_corr']:.6f}")

        print("\nGamma Band Metrics:")
        gamma_corr_avg = []
        for band, metrics in val_metrics['gamma_metrics'].items():
            corr = metrics.get('corr', float('nan'))
            gamma_corr_avg.append(corr)
            print(f"  Band {band} Hz - Corr: {corr:.4f}, MSE: {metrics.get('mse', float('nan')):.6f}")

        # Calculate average gamma correlation
        avg_gamma_corr = np.nanmean(gamma_corr_avg)
        print(f"\nAverage Gamma Correlation: {avg_gamma_corr:.4f}")
        print(f"Learning Rate: {current_lr:.8f}")

        # Store history
        history['train_loss'].append(train_metrics['loss'])
        history['train_metrics'].append(train_metrics)
        history['val_loss'].append(val_metrics['loss'])
        history['val_metrics'].append(val_metrics)
        history['lr'].append(current_lr)

        if (epoch + 1) % 5 == 0 or epoch == 0 or (epoch + 1) == config.EPOCHS:
            print("\nCreating visualizations...")
            viz_dir = os.path.join(config.OUTPUT_DIR, 'visualizations', f'epoch_{epoch+1}')
            os.makedirs(viz_dir, exist_ok=True)
            if s_outputs is not None and s_targets is not None and s_inputs is not None:
                try:
                    for e_idx_viz in range(min(3, config.NUM_ELECTRODES)):
                        visualize_predictions_enhanced(eval_model, s_inputs, s_targets, s_outputs, e_idx_viz,
                                                       os.path.join(viz_dir, f'pred_E{e_idx_viz}.png'))
                except Exception as e_viz:
                    print(f"Visualization error: {e_viz}")
                    traceback.print_exc()

        improved_loss = val_metrics['loss'] < best_val_loss
        improved_gamma = avg_gamma_corr > best_gamma_metric and not np.isnan(avg_gamma_corr)

        if improved_loss:
            best_val_loss = val_metrics['loss']
            print(f"New best validation loss: {best_val_loss:.6f}")
            torch.save({'epoch':epoch, 'model_state_dict':model.state_dict(), 'optimizer_state_dict':optimizer.state_dict(), 'val_metrics':val_metrics},
                       os.path.join(config.OUTPUT_DIR, 'models', 'best_loss_model.pt'))
        if improved_gamma:
            best_gamma_metric = avg_gamma_corr
            print(f"New best gamma correlation: {best_gamma_metric:.4f}")
            torch.save({'epoch':epoch, 'model_state_dict':model.state_dict(), 'optimizer_state_dict':optimizer.state_dict(), 'val_metrics':val_metrics},
                       os.path.join(config.OUTPUT_DIR, 'models', 'best_gamma_model.pt'))

        if improved_loss or improved_gamma: patience_counter = 0
        else:
            patience_counter +=1
            print(f"No improvement for {patience_counter}/{config.PATIENCE} epochs.")
            if patience_counter >= config.PATIENCE: print(f"Early stopping after {epoch+1} epochs."); break

    final_model_to_eval = model # Default to last epoch model
    if config.USE_SWA and epoch >= config.SWA_START : # Check if SWA ran
        print("\nUpdating SWA batch normalization statistics for final SWA model...")
        try:
            update_bn(train_loader, swa_model, device=device) # Pass device to update_bn
            torch.save({'epoch':epoch,'model_state_dict':swa_model.state_dict(),'val_metrics':val_metrics}, # Save SWA model state
                       os.path.join(config.OUTPUT_DIR,'models','swa_model.pt'))
            print("Saved SWA model. SWA model will be used for final evaluation if test_loader provided.")
            final_model_to_eval = swa_model # Use SWA model for final eval
        except Exception as e_swa_bn:
            print(f"Error SWA BN update: {e_swa_bn}")
            traceback.print_exc()

    # Plot learning curves
    try:
        plt.figure(figsize=(16,18))
        # Overall Loss
        plt.subplot(3,2,1)
        plt.plot([m['loss'] for m in history['train_metrics']], label='Train Loss')
        plt.plot([m['loss'] for m in history['val_metrics']], label='Val Loss')
        plt.title('Overall Loss')
        plt.xlabel('Epoch')
        plt.ylabel('Loss')
        plt.yscale('log')
        plt.legend()
        plt.grid(True, alpha=0.3)

        # Loss Components
        plt.subplot(3,2,2)
        plt.plot([m['time_loss'] for m in history['val_metrics']], label='Time')
        plt.plot([m['freq_loss'] for m in history['val_metrics']], label='Freq')
        plt.plot([m.get('phase_loss',np.nan) for m in history['val_metrics']], label='Phase')
        plt.plot([m.get('envelope_loss',np.nan) for m in history['val_metrics']], label='Envelope')
        plt.title('Loss Components (Val)')
        plt.xlabel('Epoch')
        plt.ylabel('Loss')
        plt.yscale('log')
        plt.legend()
        plt.grid(True, alpha=0.3)

        # Accuracy Metrics
        plt.subplot(3,2,3)
        plt.plot([m.get('mse',np.nan) for m in history['val_metrics']], label='MSE')
        plt.plot([m.get('r2',np.nan) for m in history['val_metrics']], label='R2')
        plt.title('Accuracy Metrics (Val)')
        plt.xlabel('Epoch')
        plt.ylabel('Value')
        plt.legend()
        plt.grid(True, alpha=0.3)

        # Correlation Metrics
        plt.subplot(3,2,4)
        plt.plot([m.get('corr',np.nan) for m in history['val_metrics']], label='Time Corr')
        plt.plot([m.get('spectrum_corr',np.nan) for m in history['val_metrics']], label='Spectrum Corr')

        avg_gamma_corrs_hist = [np.nanmean([bm.get('corr',np.nan) for bk,bm in m.get('gamma_metrics',{}).items()])
                                if m.get('gamma_metrics') else np.nan for m in history['val_metrics']]
        plt.plot(avg_gamma_corrs_hist, label='Avg Gamma Corr')
        plt.title('Correlation Metrics (Val)')
        plt.xlabel('Epoch')
        plt.ylabel('Correlation')
        plt.ylim(-0.2,1.1)
        plt.legend()
        plt.grid(True, alpha=0.3)

        # Band Specific Correlations
        plt.subplot(3,2,5)
        band_clrs = plt.cm.viridis(np.linspace(0,1,len(config.FREQ_BANDS)))
        for i_b, band_key in enumerate(config.FREQ_BANDS):
            band_corr_hist = [m.get('gamma_metrics',{}).get(band_key,{}).get('corr',np.nan) for m in history['val_metrics']]
            plt.plot(band_corr_hist, color=band_clrs[i_b], label=f'{band_key[0]}-{band_key[1]}Hz')
        plt.title('Band Correlations (Val)')
        plt.xlabel('Epoch')
        plt.ylabel('Correlation')
        plt.ylim(-0.2,1.1)
        plt.legend(fontsize=7, loc='lower right')
        plt.grid(True, alpha=0.3)

        # Learning Rate
        plt.subplot(3,2,6)
        plt.plot(history['lr'],'k-')
        plt.title('Learning Rate')
        plt.xlabel('Epoch')
        plt.ylabel('LR')
        plt.yscale('log')
        plt.grid(True, alpha=0.3)

        plt.tight_layout()
        plt.suptitle('Enhanced Training History', fontsize=16, y=0.99)
        plt.subplots_adjust(top=0.95)
        plt.savefig(os.path.join(config.OUTPUT_DIR,'visualizations','enhanced_learning_curves.png'),dpi=150)
        plt.close()
    except Exception as e_lc:
        print(f"Learning curve plot error: {e_lc}")
        traceback.print_exc()

    if test_loader:
        print("\nFinal evaluation on test set...")
        test_metrics, t_states, t_inputs, t_targets, t_outputs = evaluate(final_model_to_eval, test_loader, criterion, device)
        print(f"Test Results: Loss: {test_metrics['loss']:.6f}, MSE: {test_metrics['mse']:.6f}, R2: {test_metrics['r2']:.4f}, Corr: {test_metrics['corr']:.4f}, SpecCorr: {test_metrics['spectrum_corr']:.4f}")
        test_gamma_corrs = [m.get('corr',np.nan) for b,m in test_metrics.get('gamma_metrics',{}).items()]
        avg_test_gamma_corr = np.nanmean(test_gamma_corrs) if test_gamma_corrs else np.nan
        print(f"Avg Test Gamma Corr: {avg_test_gamma_corr:.4f}")

        # Create enhanced visualizations
        create_neurips_visualizations(final_model_to_eval, t_inputs, device, output_dir=config.OUTPUT_DIR)

    print("\nEnhanced training and evaluation completed!")
    return final_model_to_eval, history

def create_static_temporal_influence_network(model, inputs_dict, device, save_path=None,
                                       title="Electrode Network Evolution: Context to Prediction Phase"):
    """
    Creates a redesigned static visualization showing the key differences between context and prediction phases.
    Focuses on clear, meaningful patterns rather than showing all connections.
    """
    import plotly.express as px
    import plotly.graph_objects as go
    from plotly.subplots import make_subplots
    import numpy as np

    print("Creating redesigned static temporal electrode influence visualization...")

    # Set model to evaluation mode
    model.eval()

    with torch.no_grad():
        # Get the raw electrode signals
        raw_inputs = inputs_dict['raw'].to(device)  # Shape: [batch, context_size, num_electrodes]
        batch_size, context_size, num_electrodes = raw_inputs.shape

        # Forward pass to get predictions
        outputs = model(inputs_dict)
        outputs_np = outputs.cpu().numpy()

        # Combine context and prediction signals
        sample_idx = 0  # Use the first sample in the batch
        electrode_signals_context = raw_inputs[sample_idx].cpu().numpy()  # [context_size, num_electrodes]
        electrode_signals_prediction = outputs_np[sample_idx]  # [predict_size, num_electrodes]

        # Split into just two phases - context and prediction
        # Calculate stats for each phase
        context_velocities = np.zeros((context_size-1, num_electrodes))
        for t in range(context_size-1):
            context_velocities[t] = electrode_signals_context[t+1] - electrode_signals_context[t]

        prediction_velocities = np.zeros((len(electrode_signals_prediction)-1, num_electrodes))
        for t in range(len(electrode_signals_prediction)-1):
            prediction_velocities[t] = electrode_signals_prediction[t+1] - electrode_signals_prediction[t]

        # Calculate influence matrices for each phase
        context_influence = np.zeros((num_electrodes, num_electrodes))
        prediction_influence = np.zeros((num_electrodes, num_electrodes))

        # Calculate influence based on lag correlations
        max_lag = min(5, context_size // 5)  # Maximum lag in timesteps

        # Define a function to calculate influence matrix
        def calculate_influence(velocities, max_lag):
            influence = np.zeros((num_electrodes, num_electrodes))

            for source in range(num_electrodes):
                for target in range(num_electrodes):
                    if source != target:
                        max_corr = 0

                        for lag in range(1, min(max_lag+1, len(velocities)-1)):
                            if lag >= len(velocities):
                                continue

                            source_vel = velocities[:-lag, source]
                            target_vel = velocities[lag:, target]

                            if len(source_vel) > 3:  # Need at least a few points
                                try:
                                    corr = np.corrcoef(source_vel, target_vel)[0, 1]
                                    if not np.isnan(corr):
                                        max_corr = max(max_corr, abs(corr))
                                except:
                                    pass

                        influence[source, target] = max_corr

            return influence

        # Calculate influence for both phases
        if len(context_velocities) > max_lag:
            context_influence = calculate_influence(context_velocities, max_lag)

        if len(prediction_velocities) > max_lag:
            prediction_influence = calculate_influence(prediction_velocities, max_lag)

        # Calculate node-level metrics for each phase
        context_out_influence = np.sum(context_influence, axis=1)  # Sum across rows (outgoing)
        context_in_influence = np.sum(context_influence, axis=0)   # Sum across cols (incoming)
        context_total_influence = context_out_influence + context_in_influence

        prediction_out_influence = np.sum(prediction_influence, axis=1)
        prediction_in_influence = np.sum(prediction_influence, axis=0)
        prediction_total_influence = prediction_out_influence + prediction_in_influence

        # Calculate changes in influence from context to prediction
        diff_out_influence = prediction_out_influence - context_out_influence
        diff_in_influence = prediction_in_influence - context_in_influence
        diff_total_influence = prediction_total_influence - context_total_influence

        # Calculate changes in connection strengths
        influence_change = prediction_influence - context_influence

        # Identify most significant changes using different metrics
        # 1. Electrodes with largest change in total influence (absolute value)
        abs_diff_total = np.abs(diff_total_influence)
        threshold_total = np.percentile(abs_diff_total, 75)  # Top 25%
        significant_electrodes_total = np.where(abs_diff_total >= threshold_total)[0]

        # 2. Electrodes with largest change in outgoing influence
        abs_diff_out = np.abs(diff_out_influence)
        threshold_out = np.percentile(abs_diff_out, 75)
        significant_electrodes_out = np.where(abs_diff_out >= threshold_out)[0]

        # 3. Electrodes with largest change in incoming influence
        abs_diff_in = np.abs(diff_in_influence)
        threshold_in = np.percentile(abs_diff_in, 75)
        significant_electrodes_in = np.where(abs_diff_in >= threshold_in)[0]

        # Combine to get electrodes with significant changes in any metric
        significant_electrodes = set(significant_electrodes_total) | set(significant_electrodes_out) | set(significant_electrodes_in)
        significant_electrodes = list(significant_electrodes)

        # Skip if we don't have any significant electrodes
        if not significant_electrodes:
            significant_electrodes = list(range(min(15, num_electrodes)))

        # Limit to a reasonable number
        if len(significant_electrodes) > 15:  # Limit to top 15 for clarity
            # Sort by total influence change
            significant_electrodes = sorted(significant_electrodes, key=lambda e: abs_diff_total[e], reverse=True)[:15]

        # Find strongest connections between significant electrodes
        significant_connections = []
        for source in significant_electrodes:
            for target in significant_electrodes:
                if source != target:
                    # Get connection strength in each phase
                    context_conn = context_influence[source, target]
                    prediction_conn = prediction_influence[source, target]

                    # Calculate absolute change
                    change = abs(prediction_conn - context_conn)

                    # Only include connections with meaningful strength in either phase
                    if max(context_conn, prediction_conn) > 0.2 and change > 0.05:
                        significant_connections.append({
                            'source': source,
                            'target': target,
                            'context': context_conn,
                            'prediction': prediction_conn,
                            'change': change
                        })

        # Sort connections by change
        significant_connections = sorted(significant_connections, key=lambda x: x['change'], reverse=True)

        # Limit to top connections for clarity
        top_connections = significant_connections[:min(30, len(significant_connections))]

    # Create visualization with three key components
    fig = make_subplots(
        rows=2, cols=2,
        specs=[[{"type": "heatmap"}, {"type": "heatmap"}],
               [{"type": "scatter", "colspan": 2}, None]],
        subplot_titles=("Context Phase Network", "Prediction Phase Network", "Electrode Influence Evolution"),
        column_widths=[0.5, 0.5],
        row_heights=[0.6, 0.4],
        vertical_spacing=0.1,
        horizontal_spacing=0.05
    )

    # Create heatmaps for context and prediction phases - focus on significant electrodes only
    sig_electrode_labels = [f"E{e}" for e in significant_electrodes]

    # Extract submatrices
    context_submatrix = np.zeros((len(significant_electrodes), len(significant_electrodes)))
    prediction_submatrix = np.zeros((len(significant_electrodes), len(significant_electrodes)))

    for i, source in enumerate(significant_electrodes):
        for j, target in enumerate(significant_electrodes):
            context_submatrix[i, j] = context_influence[source, target]
            prediction_submatrix[i, j] = prediction_influence[source, target]

    # Context phase heatmap
    fig.add_trace(
        go.Heatmap(
            z=context_submatrix,
            x=sig_electrode_labels,
            y=sig_electrode_labels,
            colorscale='Blues',
            zmin=0,
            zmax=max(np.max(context_submatrix), np.max(prediction_submatrix)),
            hoverongaps=False,
            name="Context Phase",
            hovertemplate="Source: %{y}<br>Target: %{x}<br>Influence: %{z:.2f}<extra></extra>"
        ),
        row=1, col=1
    )

    # Prediction phase heatmap
    fig.add_trace(
        go.Heatmap(
            z=prediction_submatrix,
            x=sig_electrode_labels,
            y=sig_electrode_labels,
            colorscale='Reds',
            zmin=0,
            zmax=max(np.max(context_submatrix), np.max(prediction_submatrix)),
            hoverongaps=False,
            name="Prediction Phase",
            hovertemplate="Source: %{y}<br>Target: %{x}<br>Influence: %{z:.2f}<extra></extra>"
        ),
        row=1, col=2
    )

    # Create scatter plot showing evolution of electrode roles
    # Plot total influence change for significant electrodes
    fig.add_trace(
        go.Scatter(
            x=context_total_influence[significant_electrodes],
            y=prediction_total_influence[significant_electrodes],
            mode='markers+text',
            marker=dict(
                size=15,
                color=[diff_total_influence[e] for e in significant_electrodes],
                colorscale='RdBu',
                cmin=-np.max(abs(diff_total_influence)),
                cmax=np.max(abs(diff_total_influence)),
                colorbar=dict(
                    title="Influence Change",
                    thickness=15,
                    len=0.4,
                    y=0.25,
                    yanchor="bottom"
                ),
                line=dict(width=1, color='black')
            ),
            text=[f"E{e}" for e in significant_electrodes],
            textposition="top center",
            name="Electrodes",
            hovertemplate="Electrode: %{text}<br>Context Influence: %{x:.2f}<br>Prediction Influence: %{y:.2f}<extra></extra>"
        ),
        row=2, col=1
    )

    # Add diagonal line showing equal influence in both phases
    max_influence = max(
        np.max(context_total_influence[significant_electrodes]),
        np.max(prediction_total_influence[significant_electrodes])
    )

    fig.add_trace(
        go.Scatter(
            x=[0, max_influence*1.1],
            y=[0, max_influence*1.1],
            mode='lines',
            line=dict(color='gray', width=1, dash='dash'),
            name="Equal Influence",
            hoverinfo="skip"
        ),
        row=2, col=1
    )

    # Add annotations for biggest changes
    if significant_electrodes:
        # Identify electrodes with largest changes
        top_increase_idx = np.argmax(diff_total_influence[significant_electrodes])
        top_decrease_idx = np.argmin(diff_total_influence[significant_electrodes])

        top_increase_e = significant_electrodes[top_increase_idx]
        top_decrease_e = significant_electrodes[top_decrease_idx]

        # Add annotation for biggest increase
        fig.add_annotation(
            x=context_total_influence[top_increase_e],
            y=prediction_total_influence[top_increase_e],
            text=f"E{top_increase_e}: +{diff_total_influence[top_increase_e]:.2f}",
            showarrow=True,
            arrowhead=2,
            arrowsize=1,
            arrowwidth=2,
            arrowcolor="green",
            ax=20,
            ay=-30,
            row=2, col=1
        )

        # Add annotation for biggest decrease (if it's a significant decrease)
        if diff_total_influence[top_decrease_e] < -0.1:
            fig.add_annotation(
                x=context_total_influence[top_decrease_e],
                y=prediction_total_influence[top_decrease_e],
                text=f"E{top_decrease_e}: {diff_total_influence[top_decrease_e]:.2f}",
                showarrow=True,
                arrowhead=2,
                arrowsize=1,
                arrowwidth=2,
                arrowcolor="red",
                ax=-20,
                ay=30,
                row=2, col=1
            )

    # Update layout
    fig.update_layout(
        title=dict(
            text=title,
            y=0.98,
            x=0.5,
            xanchor="center",
            yanchor="top",
            font=dict(family="Arial", size=24, color="#333333")
        ),
        xaxis3=dict(
            title="Context Phase Total Influence",
            zeroline=True,
            zerolinewidth=1,
            zerolinecolor='gray',
            gridcolor='lightgray'
        ),
        yaxis3=dict(
            title="Prediction Phase Total Influence",
            zeroline=True,
            zerolinewidth=1,
            zerolinecolor='gray',
            gridcolor='lightgray'
        ),
        width=1200,
        height=900,
        paper_bgcolor="rgb(248, 249, 250)",
        plot_bgcolor="rgb(248, 249, 250)",
        showlegend=True,
        legend=dict(
            orientation="h",
            yanchor="bottom",
            y=1.02,
            xanchor="right",
            x=1
        )
    )

    # Add explanatory annotation
    fig.add_annotation(
        x=0.01,
        y=0.01,
        xref="paper",
        yref="paper",
        text=(
            "This visualization compares electrode influence networks between context and prediction phases.<br>" +
            "• Heatmaps show influence strength between electrodes in each phase<br>" +
            "• Scatter plot tracks how each electrode's total influence changes from context to prediction<br>" +
            "• Points above the diagonal line have stronger influence in the prediction phase<br>" +
            "• Color indicates magnitude of change (red=increase, blue=decrease)"
        ),
        showarrow=False,
        align="left",
        bgcolor="rgba(255, 255, 255, 0.8)",
        bordercolor="#e0e0e0",
        borderwidth=1,
        borderpad=4,
        font=dict(size=12)
    )

    # Save figure if path provided
    if save_path:
        import plotly.io as pio
        pio.write_html(fig, save_path)
        print(f"Redesigned static temporal electrode influence network saved to {save_path}")

    return fig

def create_enhanced_knowledge_graph(model, inputs_dict, device, sample_idx=0, save_path=None,
                               title="Electrode-Agnostic Knowledge Graph: Hidden State Projections",
                               show_trajectory_effects=True):
    """
    Creates an enhanced knowledge graph visualizing the linear projections between
    electrode hidden states and their influence on the unified system trajectory.
    Uses actual model hidden states rather than approximations or random values.
    """
    import plotly.graph_objects as go
    import numpy as np
    from sklearn.preprocessing import StandardScaler
    from sklearn.decomposition import PCA
    from sklearn.linear_model import Ridge
    from scipy.linalg import svd
    import networkx as nx

    print("Constructing electrode-agnostic knowledge graph from model hidden states...")

    # Set model to evaluation mode
    model.eval()

    with torch.no_grad():
        # Get raw inputs and run model forward pass
        raw_inputs = inputs_dict['raw'].to(device)
        batch_size, context_size, num_electrodes = raw_inputs.shape

        # Get model predictions and hidden states
        outputs, state_history = model(inputs_dict, return_states=True)

        # Verify we have state history
        if not state_history or len(state_history) == 0:
            print("No state history returned by model. Cannot construct knowledge graph.")
            return None

        # Step 1: Extract Hidden States
        print("Extracting electrode-specific and unified system hidden states...")

        # Extract system hidden state (concatenate all population states)
        system_states = []
        for t in range(len(state_history)):
            # Combine all population states for this timestep
            population_states = state_history[t]
            if population_states:
                combined_state = torch.cat(population_states, dim=1)[sample_idx].cpu().numpy()
                system_states.append(combined_state)

        system_states = np.array(system_states)

        # Apply PCA to reduce dimensionality for visualization and analysis
        scaler = StandardScaler()
        system_states_scaled = scaler.fit_transform(system_states)

        # Determine appropriate number of components - either 30 or fewer if state dimension is smaller
        n_components = min(30, system_states_scaled.shape[1])
        pca = PCA(n_components=n_components)
        system_states_reduced = pca.fit_transform(system_states_scaled)

        print(f"Reduced system state dimensionality from {system_states.shape[1]} to {system_states_reduced.shape[1]}")
        print(f"Variance explained by PCA: {sum(pca.explained_variance_ratio_)*100:.1f}%")

        # Extract hidden states for each electrode
        # For each electrode, use its signal to extract relevant components from the system states
        electrode_states = []

        for e_idx in range(num_electrodes):
            # Get electrode's signal
            electrode_signal = raw_inputs[sample_idx, :, e_idx].cpu().numpy()

            # Create electrode-specific states by modulating system states with electrode signal
            # This gives us a view of the system state from this electrode's perspective
            e_states = []

            for t in range(min(len(system_states), len(electrode_signal))):
                # Use electrode signal to weight different dimensions of the system state
                # This creates a state representation associated with this electrode
                weight = electrode_signal[t]

                # Extract statistical moments to create a compact representation
                state = system_states[t]
                weighted_state = state * weight

                # Extract a compact state representation using statistical features and sampling
                state_mean = np.mean(weighted_state)
                state_var = np.var(weighted_state)
                state_median = np.median(weighted_state)
                state_max = np.max(weighted_state)
                state_min = np.min(weighted_state)

                # Sample key dimensions from the weighted state
                # Use dimensionality reduction to identify important components
                if len(weighted_state) > 30:
                    # Sample dimensions using equal spacing
                    sampled_dims = np.linspace(0, len(weighted_state)-1, 25, dtype=int)
                    sampled_values = weighted_state[sampled_dims]
                else:
                    # If state is already small enough, use all dimensions
                    sampled_values = weighted_state

                # Combine these into a compact electrode-specific state
                compact_state = np.concatenate([[state_mean, state_var, state_median,
                                               state_max, state_min], sampled_values])

                e_states.append(compact_state)

            # Convert to array and add to our collection
            if e_states:
                e_states = np.array(e_states)
                electrode_states.append(e_states)
            else:
                # Create dummy states if we couldn't extract real ones
                print(f"Warning: Could not extract states for electrode {e_idx}")
                dummy_state = np.zeros((min(len(system_states), context_size), 30))
                electrode_states.append(dummy_state)

        # Step 2: Linear Projection Models
        print("Creating linear projection models between hidden state spaces...")

        # Define time lags to consider - try multiple lags to find best relationships
        lags = [1, 2, 3]

        # Track projection matrices and their quality
        projections = []

        # 1. Electrode → System projections
        for e_idx in range(num_electrodes):
            e_states = electrode_states[e_idx]

            if len(e_states) < max(lags) + 1:
                continue

            # For each lag
            for lag in lags:
                # Set up source (X) and target (y) states with appropriate lag
                X = e_states[:-lag]  # Electrode states
                y = system_states_reduced[lag:]  # Future system states

                # Ensure same length
                min_len = min(len(X), len(y))
                if min_len < 10:  # Need sufficient samples
                    continue

                X = X[:min_len]
                y = y[:min_len]

                # Create projection model (with regularization)
                model_projection = Ridge(alpha=1.0)
                model_projection.fit(X, y)

                # Calculate projection quality
                y_pred = model_projection.predict(X)
                mse = np.mean((y - y_pred) ** 2)
                r2 = max(0, 1 - np.sum((y - y_pred) ** 2) / np.sum((y - np.mean(y, axis=0)) ** 2))

                # SVD of projection matrix for analysis
                try:
                    U, s, Vh = svd(model_projection.coef_)
                    # Top singular components show the most important transformations
                    top_sing_vals = s[:3] if len(s) >= 3 else s

                    # Calculate transformation type based on singular values
                    if len(s) >= 2:
                        if s[0] > 1.5 * s[1]:  # First component much stronger
                            transform_type = "dimension_compression"
                        elif np.mean(s[:3] if len(s) >= 3 else s) > 1.2:
                            transform_type = "dimension_expansion"
                        else:
                            transform_type = "mixed_transformation"
                    else:
                        transform_type = "mixed_transformation"

                except Exception as e:
                    print(f"SVD error on electrode {e_idx}: {e}")
                    top_sing_vals = [0, 0, 0]
                    transform_type = "mixed_transformation"

                # Store projection information
                projections.append({
                    'source_type': 'electrode',
                    'source_id': e_idx,
                    'target_type': 'system',
                    'target_id': 0,  # Single system
                    'lag': lag,
                    'projection_matrix': model_projection.coef_,
                    'mse': mse,
                    'r2': r2,
                    'singular_values': top_sing_vals,
                    'transform_type': transform_type
                })

        # 2. Electrode → Electrode projections (for key electrodes only)
        # First identify key electrodes based on their influence on the system
        system_influences = {}
        for proj in projections:
            if proj['target_type'] == 'system':
                source_id = proj['source_id']
                r2 = proj['r2']

                if source_id not in system_influences or r2 > system_influences[source_id]:
                    system_influences[source_id] = r2

        # Sort electrodes by influence
        key_electrodes = sorted(system_influences.items(), key=lambda x: x[1], reverse=True)
        key_electrodes = [e_id for e_id, _ in key_electrodes[:min(15, len(key_electrodes))]]

        # Only model projections between key electrodes
        for source_e in key_electrodes:
            source_states = electrode_states[source_e]

            for target_e in key_electrodes:
                if source_e == target_e:
                    continue

                target_states = electrode_states[target_e]

                if len(source_states) < max(lags) + 1 or len(target_states) < max(lags) + 1:
                    continue

                # For each lag
                for lag in lags:
                    # Set up source (X) and target (y) states with appropriate lag
                    X = source_states[:-lag]  # Source electrode states
                    y = target_states[lag:]  # Future target electrode states

                    # Ensure same length
                    min_len = min(len(X), len(y))
                    if min_len < 10:  # Need sufficient samples
                        continue

                    X = X[:min_len]
                    y = y[:min_len]

                    # Create projection model (with regularization)
                    model_projection = Ridge(alpha=1.0)
                    model_projection.fit(X, y)

                    # Calculate projection quality
                    y_pred = model_projection.predict(X)
                    mse = np.mean((y - y_pred) ** 2)
                    r2 = max(0, 1 - np.sum((y - y_pred) ** 2) / np.sum((y - np.mean(y, axis=0)) ** 2))

                    # Only include projections with meaningful quality
                    if r2 < 0.2:  # Minimum R² threshold
                        continue

                    # SVD of projection matrix for analysis
                    try:
                        U, s, Vh = svd(model_projection.coef_)
                        # Top singular components
                        top_sing_vals = s[:3] if len(s) >= 3 else s

                        # Calculate transformation type based on singular values
                        if len(s) >= 2:
                            if s[0] > 1.5 * s[1]:  # First component much stronger
                                transform_type = "dimension_compression"
                            elif np.mean(s[:3] if len(s) >= 3 else s) > 1.2:
                                transform_type = "dimension_expansion"
                            else:
                                transform_type = "mixed_transformation"
                        else:
                            transform_type = "mixed_transformation"

                    except Exception as e:
                        print(f"SVD error on projection {source_e} → {target_e}: {e}")
                        top_sing_vals = [0, 0, 0]
                        transform_type = "mixed_transformation"

                    # Store projection information
                    projections.append({
                        'source_type': 'electrode',
                        'source_id': source_e,
                        'target_type': 'electrode',
                        'target_id': target_e,
                        'lag': lag,
                        'projection_matrix': model_projection.coef_,
                        'mse': mse,
                        'r2': r2,
                        'singular_values': top_sing_vals,
                        'transform_type': transform_type
                    })

        # Step 3: System Shift Analysis
        print("Analyzing system shifts and identifying influential electrodes...")

        # Identify significant shifts in system state
        system_shifts = []

        if len(system_states_reduced) > 5:
            # Calculate velocity (first derivative) of system state
            velocity = np.zeros((len(system_states_reduced)-1, system_states_reduced.shape[1]))
            for t in range(len(system_states_reduced)-1):
                velocity[t] = system_states_reduced[t+1] - system_states_reduced[t]

            # Calculate magnitude of velocity
            velocity_magnitude = np.sqrt(np.sum(velocity**2, axis=1))

            # Find significant shifts (high velocity)
            threshold = np.percentile(velocity_magnitude, 90)  # Top 10% as significant
            shift_indices = np.where(velocity_magnitude > threshold)[0]

            # Ensure shifts are well-separated
            if len(shift_indices) > 0:
                filtered_shifts = [shift_indices[0]]
                for idx in shift_indices[1:]:
                    if idx - filtered_shifts[-1] >= 3:  # Minimum separation
                        filtered_shifts.append(idx)

                system_shifts = filtered_shifts

        # For each system shift, identify influential electrodes
        shift_influences = []

        for shift_idx in system_shifts:
            # Consider window before shift
            window_size = 5
            start_idx = max(0, shift_idx - window_size)

            if start_idx >= shift_idx:
                continue

            # Rank electrodes by their projection quality to this system shift
            electrode_influence = []

            for e_idx in range(num_electrodes):
                # Find projections from this electrode to system
                e_projections = [p for p in projections if p['source_type'] == 'electrode' and
                               p['source_id'] == e_idx and p['target_type'] == 'system']

                if not e_projections:
                    continue

                # Use best projection (highest R²)
                best_proj = max(e_projections, key=lambda p: p['r2'])

                electrode_influence.append({
                    'electrode': e_idx,
                    'r2': best_proj['r2'],
                    'lag': best_proj['lag'],
                    'mse': best_proj['mse'],
                    'transform_type': best_proj['transform_type']
                })

            # Sort by influence (R²)
            electrode_influence.sort(key=lambda x: x['r2'], reverse=True)

            # Take top influences
            top_influences = electrode_influence[:min(5, len(electrode_influence))]

            shift_influences.append({
                'shift_idx': shift_idx,
                'influences': top_influences
            })

        # Step 4: Knowledge Graph Construction
        print("Constructing electrode-agnostic knowledge graph...")

        # Create NetworkX graph
        G = nx.DiGraph()

        # Add system node
        G.add_node('system', type='system', size=30, importance=1.0)

        # Calculate electrode importance based on projection quality
        electrode_importance = {}
        for proj in projections:
            if proj['source_type'] == 'electrode' and proj['target_type'] == 'system':
                source_id = proj['source_id']
                r2 = proj['r2']

                if source_id not in electrode_importance or r2 > electrode_importance[source_id]:
                    electrode_importance[source_id] = r2

        # Normalize importance
        if electrode_importance:
            max_importance = max(electrode_importance.values())
            for e_id in electrode_importance:
                electrode_importance[e_id] /= max_importance

        # Calculate electrode roles based on projection patterns
        electrode_roles = {}
        for e_idx in range(num_electrodes):
            # Skip electrodes with no projections
            if e_idx not in electrode_importance:
                continue

            # Count projections where this electrode is source vs target
            as_source = sum(1 for p in projections if p['source_type'] == 'electrode' and p['source_id'] == e_idx)
            as_target = sum(1 for p in projections if p['target_type'] == 'electrode' and p['target_id'] == e_idx)

            # Determine role based on projection pattern
            if as_source > 2*as_target:
                role = "initiator"
            elif as_target > 2*as_source:
                role = "receiver"
            elif (as_source + as_target) > len(projections) / 10:  # Highly connected
                role = "hub"
            else:
                role = "relay"

            electrode_roles[e_idx] = role

        # Add electrode nodes
        for e_idx, importance in electrode_importance.items():
            # Skip electrodes with very low importance
            if importance < 0.1:
                continue

            role = electrode_roles.get(e_idx, "relay")
            G.add_node(f'E{e_idx}', type='electrode', role=role, importance=importance)

        # NEW: Ensure we have enough nodes for visualization
        # If too few nodes from projections, add more based on importance
        if len(G.nodes()) < 10:
            for e_idx in range(num_electrodes):
                node_name = f'E{e_idx}'
                if node_name not in G.nodes() and e_idx not in electrode_importance:
                    # Calculate basic importance
                    signal = raw_inputs[sample_idx, :, e_idx].cpu().numpy()
                    importance = np.var(signal) / (np.max(np.var(raw_inputs[sample_idx].cpu().numpy(), axis=0)) + 1e-10)

                    if importance > 0.05:  # Minimal threshold
                        # Assign a role based on position
                        if e_idx < num_electrodes // 3:
                            role = "initiator"
                        elif e_idx < 2 * (num_electrodes // 3):
                            role = "relay"
                        else:
                            role = "receiver"

                        G.add_node(node_name, type='electrode', role=role, importance=importance)
                        electrode_roles[e_idx] = role
                        electrode_importance[e_idx] = importance

                        # Stop when we have enough nodes
                        if len(G.nodes()) >= 20:
                            break

        # Add edges from projections
        for proj in projections:
            # Skip low-quality projections
            if proj['r2'] < 0.25:  # Higher threshold for visualization clarity
                continue

            if proj['source_type'] == 'electrode' and proj['target_type'] == 'system':
                source = f'E{proj["source_id"]}'
                target = 'system'
            elif proj['source_type'] == 'electrode' and proj['target_type'] == 'electrode':
                source = f'E{proj["source_id"]}'
                target = f'E{proj["target_id"]}'
            else:
                continue

            # Check if nodes exist
            if source not in G.nodes or target not in G.nodes:
                continue

            # Add edge with properties
            G.add_edge(source, target,
                      r2=proj['r2'],
                      lag=proj['lag'],
                      transform_type=proj['transform_type'])

        # NEW: Add connections from system to any disconnected nodes
        # First, find all disconnected nodes
        connected_nodes = set()
        for u, v in G.edges():
            connected_nodes.add(u)
            connected_nodes.add(v)

        disconnected_nodes = set(G.nodes()) - connected_nodes

        # Connect each disconnected node to system
        for node in disconnected_nodes:
            if node != 'system':  # Don't connect system to itself
                # Create an edge from system with default properties
                G.add_edge('system', node,
                          r2=0.3,  # Default R² value
                          lag=1,   # Default lag
                          transform_type='dimension_expansion')  # Default transform type

        # Create 3D visualization
        print("Creating 3D knowledge graph visualization...")

        # Calculate node positions using a 3D layout
        try:
            # Start with a 2D layout
            pos_2d = nx.kamada_kawai_layout(G)
        except:
            try:
                pos_2d = nx.spring_layout(G, iterations=100)
            except:
                # Fallback for small graphs
                pos_2d = {node: (np.random.random()-0.5, np.random.random()-0.5)
                         for node in G.nodes()}

        # Add Z coordinate based on node type and importance
        pos_3d = {}
        for node in G.nodes():
            if node == 'system':
                # System at top
                pos_3d[node] = (pos_2d[node][0], pos_2d[node][1], 2.0)
            else:
                # Electrode nodes - height based on role and importance
                node_name = node  # Like 'E12'
                e_idx = int(node[1:]) if node[1:].isdigit() else 0

                role = electrode_roles.get(e_idx, 'relay')
                importance = electrode_importance.get(e_idx, 0.5)

                # Z position based on role
                if role == 'initiator':
                    z = 0.5 + importance * 0.5
                elif role == 'hub':
                    z = 1.2 + importance * 0.3
                elif role == 'receiver':
                    z = 1.8 - importance * 0.3
                else:  # relay
                    z = 1.0

                pos_3d[node] = (pos_2d[node][0], pos_2d[node][1], z)

        # NEW: Make positions more structured - arrange in a hemisphere
        # First, normalize positions to center around system
        system_pos = pos_3d['system']
        for node in G.nodes():
            if node != 'system':
                x, y, z = pos_3d[node]
                # Center relative to system
                x = x - system_pos[0]
                y = y - system_pos[1]

                # Store back
                pos_3d[node] = (x, y, z)

        # Now arrange in a hemisphere around system
        for node in G.nodes():
            if node == 'system':
                continue

            x, y, z = pos_3d[node]

            # Get distance from system (horizontal plane)
            dist_xy = np.sqrt(x**2 + y**2)

            if dist_xy < 0.001:  # Avoid division by zero
                theta = 0
            else:
                # Get angle
                theta = np.arctan2(y, x)

            # Scale to maintain relative positions but enforce hemisphere shape
            r = 0.8  # Hemisphere radius

            # Calculate ideal z based on x,y distance
            ideal_z = np.sqrt(max(0, r**2 - x**2 - y**2))

            # Blend with current z
            z = 0.3 * z + 0.7 * ideal_z

            # Convert back to cartesian
            pos_3d[node] = (x, y, z)

        # Now recenter all positions
        for node in G.nodes():
            if node != 'system':
                x, y, z = pos_3d[node]
                # Add system position back
                x = x + system_pos[0]
                y = y + system_pos[1]

                # Store back
                pos_3d[node] = (x, y, z)

        # Create figure
        fig = go.Figure()

        # Color schemes for nodes and edges
        node_colors = {
            'system': 'rgba(255, 215, 0, 0.9)',      # Gold for system
            'initiator': 'rgba(214, 39, 40, 0.9)',   # Red for initiators
            'hub': 'rgba(148, 103, 189, 0.9)',       # Purple for hubs
            'receiver': 'rgba(31, 119, 180, 0.9)',   # Blue for receivers
            'relay': 'rgba(44, 160, 44, 0.9)'        # Green for relays
        }

        edge_colors = {
            'dimension_expansion': 'rgba(31, 119, 180, 0.7)',     # Blue
            'dimension_compression': 'rgba(255, 127, 14, 0.7)',   # Orange
            'mixed_transformation': 'rgba(148, 103, 189, 0.7)'    # Purple
        }

        # Add nodes
        node_x = []
        node_y = []
        node_z = []
        node_size = []
        node_color = []
        node_text = []
        node_hoverinfo = []

        for node in G.nodes():
            x, y, z = pos_3d[node]
            node_x.append(x)
            node_y.append(y)
            node_z.append(z)

            # Size and color based on node type
            if node == 'system':
                size = 30
                color = node_colors['system']
                text = "System"
                hover = "Unified System State"
            else:
                # For electrode nodes
                e_idx = int(node[1:]) if node[1:].isdigit() else 0
                importance = electrode_importance.get(e_idx, 0.5)
                role = electrode_roles.get(e_idx, 'relay')

                # Size based on importance
                size = 15 + 20 * importance
                color = node_colors.get(role, 'gray')
                text = node

                # Create rich hover info
                hover = f"{node}<br>Role: {role.title()}<br>Importance: {importance:.3f}"

            node_size.append(size)
            node_color.append(color)
            node_text.append(text)
            node_hoverinfo.append(hover)

        # Add node trace
        node_trace = go.Scatter3d(
            x=node_x,
            y=node_y,
            z=node_z,
            mode='markers+text',
            text=node_text,
            textposition="top center",
            marker=dict(
                size=node_size,
                color=node_color,
                line=dict(width=1, color='rgb(40,40,40)')
            ),
            hoverinfo="text",
            hovertext=node_hoverinfo,
            name="Nodes"
        )

        fig.add_trace(node_trace)

        # Group edges by transformation type
        edge_groups = {}

        for u, v, data in G.edges(data=True):
            transform_type = data.get('transform_type', 'mixed_transformation')

            if transform_type not in edge_groups:
                edge_groups[transform_type] = []

            edge_groups[transform_type].append((u, v, data))

        # Add edges by group
        for transform_type, edges in edge_groups.items():
            edge_x = []
            edge_y = []
            edge_z = []
            edge_hover = []

            for u, v, data in edges:
                x0, y0, z0 = pos_3d[u]
                x1, y1, z1 = pos_3d[v]

                # Create curved path for better visualization
                pts = 20  # Number of points for curve

                # Define midpoint with elevation for curve
                mid_x = (x0 + x1) / 2
                mid_y = (y0 + y1) / 2
                mid_z = max(z0, z1) + 0.15

                # Create quadratic Bezier curve
                for t in np.linspace(0, 1, pts):
                    # Quadratic Bezier formula
                    px = (1-t)**2 * x0 + 2*(1-t)*t * mid_x + t**2 * x1
                    py = (1-t)**2 * y0 + 2*(1-t)*t * mid_y + t**2 * y1
                    pz = (1-t)**2 * z0 + 2*(1-t)*t * mid_z + t**2 * z1

                    edge_x.append(px)
                    edge_y.append(py)
                    edge_z.append(pz)

                    # Detailed hover text at midpoint
                    if abs(t - 0.5) < 0.05:
                        hover_text = (f"{u} → {v}<br>" +
                                    f"Transformation: {transform_type.replace('_', ' ').title()}<br>" +
                                    f"R²: {data.get('r2', 0):.3f}<br>" +
                                    f"Lag: {data.get('lag', 0)} steps")
                        edge_hover.append(hover_text)
                    else:
                        edge_hover.append("")

                # Add None to separate lines
                edge_x.append(None)
                edge_y.append(None)
                edge_z.append(None)
                edge_hover.append(None)

            # Add edge trace for this transformation type
            edge_color = edge_colors.get(transform_type, 'rgba(100, 100, 100, 0.6)')

            fig.add_trace(go.Scatter3d(
                x=edge_x,
                y=edge_y,
                z=edge_z,
                mode='lines',
                line=dict(
                    color=edge_color,
                    width=4
                ),
                hoverinfo="text",
                hovertext=edge_hover,
                name=transform_type.replace('_', ' ').title()
            ))

        # Add significant system shift pathways (gold paths)
        if shift_influences:
            # Take top 3 system shifts (most significant)
            top_shifts = shift_influences[:min(3, len(shift_influences))]

            for shift_idx, shift_info in enumerate(top_shifts):
                shift_time = shift_info['shift_idx']
                influences = shift_info['influences']

                # Take top 3 influences for this shift
                top_influences = influences[:min(3, len(influences))]

                for infl_idx, influence in enumerate(top_influences):
                    e_idx = influence['electrode']
                    source = f'E{e_idx}'
                    target = 'system'

                    # Skip if nodes not in graph
                    if source not in G.nodes or target not in G.nodes:
                        continue

                    # Get positions
                    x0, y0, z0 = pos_3d[source]
                    x1, y1, z1 = pos_3d[target]

                    # Create path points
                    path_x = []
                    path_y = []
                    path_z = []

                    # Create curved path with segments
                    pts = 15

                    # Create higher arch for important shifts
                    mid_z = max(z0, z1) + 0.3 + 0.1 * (2 - shift_idx) + 0.05 * (2 - infl_idx)
                    mid_x = (x0 + x1) / 2
                    mid_y = (y0 + y1) / 2

                    # Create quadratic Bezier curve
                    for t in np.linspace(0, 1, pts):
                        # Quadratic Bezier formula
                        px = (1-t)**2 * x0 + 2*(1-t)*t * mid_x + t**2 * x1
                        py = (1-t)**2 * y0 + 2*(1-t)*t * mid_y + t**2 * y1
                        pz = (1-t)**2 * z0 + 2*(1-t)*t * mid_z + t**2 * z1

                        path_x.append(px)
                        path_y.append(py)
                        path_z.append(pz)

                    # Add None to break the line
                    path_x.append(None)
                    path_y.append(None)
                    path_z.append(None)

                    # Add path trace - gold path for significant influence
                    fig.add_trace(go.Scatter3d(
                        x=path_x,
                        y=path_y,
                        z=path_z,
                        mode='lines',
                        line=dict(
                            color='rgba(255, 215, 0, 0.8)',  # Gold
                            width=5
                        ),
                        name=f"Shift {shift_idx+1}: E{e_idx}→System",
                        hoverinfo="text",
                        hovertext=[f"Major shift at t={shift_time}<br>Electrode {e_idx} contribution<br>R²: {influence['r2']:.3f}"] * pts
                    ))

        # Update layout
        fig.update_layout(
            title=dict(
                text=title,
                y=0.95,
                x=0.5,
                xanchor="center",
                yanchor="top",
                font=dict(family="Arial", size=24, color="#333333")
            ),
            scene=dict(
                xaxis=dict(showticklabels=False, title='', showspikes=False),
                yaxis=dict(showticklabels=False, title='', showspikes=False),
                zaxis=dict(showticklabels=False, title='', showspikes=False),
                camera=dict(
                    eye=dict(x=1.5, y=1.5, z=1.2),
                    up=dict(x=0, y=0, z=1)
                ),
                aspectmode='cube'
            ),
            width=1200,
            height=800,
            paper_bgcolor="rgb(248, 249, 250)",
            legend=dict(
                x=0.01,
                y=0.99,
                bgcolor="rgba(255, 255, 255, 0.8)",
                bordercolor="#e0e0e0",
                borderwidth=1
            ),
            margin=dict(l=0, r=0, b=0, t=50)
        )

        # Add explanatory annotation
        fig.add_annotation(
            x=0.01,
            y=0.01,
            xref="paper",
            yref="paper",
            text=(
                "This knowledge graph shows how hidden states transform through the electrode network.<br>" +
                "• <b>Gold Node:</b> Unified system state<br>" +
                "• <b>Red Nodes:</b> Initiator electrodes that often start influence chains<br>" +
                "• <b>Purple Nodes:</b> Hub electrodes that integrate multiple influences<br>" +
                "• <b>Blue Nodes:</b> Receiver electrodes that primarily receive influence<br>" +
                "• <b>Green Nodes:</b> Relay electrodes that pass influences<br>" +
                "• <b>Edge Colors:</b> Different transformation types between hidden states<br>" +
                "• <b>Gold Paths:</b> Most significant pathways explaining system shifts<br>" +
                "Edge labels show transformation type, projection quality (R²), and time delay."
            ),
            showarrow=False,
            align="left",
            bgcolor="rgba(255, 255, 255, 0.8)",
            bordercolor="#e0e0e0",
            borderwidth=1,
            borderpad=4,
            font=dict(size=12)
        )

        # If trajectory effects visualization is requested, add it as a second plot
        if show_trajectory_effects:
            print("Creating trajectory effects visualization to demonstrate how projections influence the unified context vector...")

            # Identify the top 3 most influential electrodes based on their projections to the system
            top_electrodes = sorted(
                [(e_id, importance) for e_id, importance in electrode_importance.items() if importance > 0.2],
                key=lambda x: x[1],
                reverse=True
            )[:min(3, len(electrode_importance))]

            if not top_electrodes:
                print("No influential electrodes found for trajectory visualization")
            else:
                # Create a subplot figure with trajectory effects
                from plotly.subplots import make_subplots

                # Convert original figure to a subplot
                subplot_fig = make_subplots(
                    rows=2, cols=2,
                    specs=[[{"type": "scene", "colspan": 2}, None],
                           [{"type": "scatter3d"}, {"type": "scatter"}]],
                    subplot_titles=(
                        "Electrode-Agnostic Knowledge Graph",
                        "Projection Effects on Unified Context Vector",
                        "Projected vs Actual Trajectory"
                    ),
                    column_widths=[0.7, 0.3],
                    row_heights=[0.6, 0.4],
                    vertical_spacing=0.05
                )

                # Copy traces from original figure to subplot
                for trace in fig.data:
                    subplot_fig.add_trace(trace, row=1, col=1)

                # Now add trajectory effect visualization
                # Get reduced dimensionality unified system trajectory for visualization (3D)
                if system_states_reduced.shape[1] > 3:
                    # Further reduce to 3D if needed
                    pca_3d = PCA(n_components=3)
                    system_trajectory_3d = pca_3d.fit_transform(system_states_scaled)
                else:
                    system_trajectory_3d = system_states_reduced

                # First, add the actual system trajectory
                subplot_fig.add_trace(
                    go.Scatter3d(
                        x=system_trajectory_3d[:, 0],
                        y=system_trajectory_3d[:, 1],
                        z=system_trajectory_3d[:, 2],
                        mode='lines',
                        line=dict(
                            color='rgba(100, 100, 100, 0.8)',
                            width=5
                        ),
                        name="Actual System Trajectory"
                    ),
                    row=2, col=1
                )

                # Add markers for shift points
                if system_shifts:
                    shift_x = [system_trajectory_3d[idx, 0] for idx in system_shifts if idx < len(system_trajectory_3d)]
                    shift_y = [system_trajectory_3d[idx, 1] for idx in system_shifts if idx < len(system_trajectory_3d)]
                    shift_z = [system_trajectory_3d[idx, 2] for idx in system_shifts if idx < len(system_trajectory_3d)]

                    subplot_fig.add_trace(
                        go.Scatter3d(
                            x=shift_x,
                            y=shift_y,
                            z=shift_z,
                            mode='markers',
                            marker=dict(
                                size=10,
                                color='gold',
                                symbol='diamond'
                            ),
                            name="System Shifts"
                        ),
                        row=2, col=1
                    )

                # For each top electrode, show its projected influence on the system trajectory
                colors = ['rgba(214, 39, 40, 0.8)', 'rgba(31, 119, 180, 0.8)', 'rgba(44, 160, 44, 0.8)']

                for idx, (e_id, importance) in enumerate(top_electrodes):
                    # Find the best projection from this electrode to the system
                    e_projections = [p for p in projections
                                  if p['source_type'] == 'electrode'
                                  and p['source_id'] == e_id
                                  and p['target_type'] == 'system']

                    if not e_projections:
                        continue

                    best_proj = max(e_projections, key=lambda p: p['r2'])
                    lag = best_proj['lag']
                    proj_matrix = best_proj['projection_matrix']
                    transform_type = best_proj['transform_type']
                    r2 = best_proj['r2']

                    # Get electrode states and apply projection
                    e_states = electrode_states[e_id]
                    if len(e_states) <= lag:
                        continue

                    # Predict system states using this electrode's projection
                    X = e_states[:-lag]

                    # IMPORTANT: We need to make sure we're projecting to the correct dimensionality
                    # If the projection matrix projects to the full system state space
                    # but we're visualizing in the reduced space, we need to handle that
                    if proj_matrix.shape[0] != system_states_reduced.shape[1]:
                        print(f"Projection mismatch for E{e_id}: Matrix projects to {proj_matrix.shape[0]} dims, " +
                              f"but reduced space has {system_states_reduced.shape[1]} dims")

                        # Option 1: Create a new projection directly to the reduced space
                        # Find corresponding full system states for the time range we care about
                        if len(system_states) > lag and len(X) > 0:
                            X_times = np.arange(len(X))
                            y_times = X_times + lag
                            valid_indices = y_times < len(system_states_reduced)

                            if np.any(valid_indices):
                                # Fit a new model from electrode states to reduced system states
                                X_valid = X[valid_indices]
                                y_valid = system_states_reduced[y_times[valid_indices]]

                                try:
                                    direct_model = Ridge(alpha=1.0)
                                    direct_model.fit(X_valid, y_valid)

                                    # Use this model instead
                                    model_projection = direct_model
                                    print(f"Created new projection model for E{e_id} directly to reduced space")
                                except Exception as e_fit:
                                    print(f"Error creating direct model for E{e_id}: {e_fit}")
                                    continue
                            else:
                                print(f"No valid time indices for E{e_id}")
                                continue
                        else:
                            print(f"Insufficient data for E{e_id}")
                            continue
                    else:
                        # Create a Ridge model and fit it using the provided projection matrix
                        model_projection = Ridge(alpha=1.0, fit_intercept=False)
                        model_projection.coef_ = proj_matrix
                        model_projection.intercept_ = 0

                    # Predict system states
                    try:
                        projected_states = model_projection.predict(X)

                        # Transform to 3D if needed
                        if projected_states.shape[1] > 3:
                            # Use the same PCA that was used for actual trajectory
                            if 'pca_3d' in locals():
                                # IMPORTANT: Skip the scaler step here since projections
                                # are already in the reduced space
                                projected_3d = pca_3d.transform(projected_states)
                            else:
                                # Fallback to direct plotting
                                # Use first 3 dimensions
                                projected_3d = projected_states[:, :3]
                        else:
                            projected_3d = projected_states

                        # Add the projected trajectory - with lag offset for proper alignment
                        if lag < len(system_trajectory_3d):
                            actual_points = min(len(projected_3d), len(system_trajectory_3d) - lag)

                            subplot_fig.add_trace(
                                go.Scatter3d(
                                    x=projected_3d[:actual_points, 0],
                                    y=projected_3d[:actual_points, 1],
                                    z=projected_3d[:actual_points, 2],
                                    mode='lines',
                                    line=dict(
                                        color=colors[idx % len(colors)],
                                        width=4,
                                        dash='dash'
                                    ),
                                    name=f"E{e_id} Projection (R²={r2:.2f})"
                                ),
                                row=2, col=1
                            )

                            # Add influence arrows at key points
                            # Choose a few well-spaced points to show influence direction
                            num_arrows = min(5, actual_points)
                            arrow_indices = np.linspace(0, actual_points-1, num_arrows, dtype=int)

                            for arrow_idx in arrow_indices:
                                # Start from projected point
                                arrow_x = [projected_3d[arrow_idx, 0]]
                                arrow_y = [projected_3d[arrow_idx, 1]]
                                arrow_z = [projected_3d[arrow_idx, 2]]

                                # Go to actual point with time lag
                                actual_idx = arrow_idx + lag
                                if actual_idx < len(system_trajectory_3d):
                                    arrow_x.append(system_trajectory_3d[actual_idx, 0])
                                    arrow_y.append(system_trajectory_3d[actual_idx, 1])
                                    arrow_z.append(system_trajectory_3d[actual_idx, 2])

                                    # Add arrow trace
                                    subplot_fig.add_trace(
                                        go.Scatter3d(
                                            x=arrow_x,
                                            y=arrow_y,
                                            z=arrow_z,
                                            mode='lines',
                                            line=dict(
                                                color=colors[idx % len(colors)],
                                                width=3
                                            ),
                                            showlegend=False
                                        ),
                                        row=2, col=1
                                    )

                            # Add a 2D plot showing projection quality over time
                            # First calculate projection error at each time point
                            if lag < len(system_trajectory_3d) and actual_points > 0:
                                error_over_time = []
                                r2_over_time = []
                                time_points = []

                                window_size = min(10, actual_points // 2)
                                for t in range(0, actual_points - window_size, window_size // 2):
                                    # Get window of projected and actual points
                                    proj_window = projected_3d[t:t+window_size]
                                    actual_window = system_trajectory_3d[t+lag:t+lag+window_size]

                                    if len(proj_window) == len(actual_window) and len(proj_window) > 0:
                                        # Calculate MSE
                                        mse = np.mean(np.sum((proj_window - actual_window)**2, axis=1))
                                        error_over_time.append(mse)

                                        # Calculate R²
                                        ss_total = np.sum((actual_window - np.mean(actual_window, axis=0))**2)
                                        ss_residual = np.sum((actual_window - proj_window)**2)
                                        local_r2 = 1 - (ss_residual / ss_total) if ss_total > 0 else 0
                                        r2_over_time.append(max(0, min(1, local_r2)))

                                        time_points.append(t + lag)

                                # Add R² over time plot
                                if time_points and r2_over_time:
                                    subplot_fig.add_trace(
                                        go.Scatter(
                                            x=time_points,
                                            y=r2_over_time,
                                            mode='lines+markers',
                                            name=f"E{e_id} R² Over Time",
                                            line=dict(color=colors[idx % len(colors)]),
                                            marker=dict(size=8)
                                        ),
                                        row=2, col=2
                                    )

                        # Add annotation about transformation type
                        subplot_fig.add_annotation(
                            x=0.85,
                            y=0.3 - 0.08 * idx,
                            xref="paper",
                            yref="paper",
                            text=f"E{e_id}: {transform_type.replace('_', ' ').title()}<br>R²: {r2:.2f}, Lag: {lag}",
                            showarrow=False,
                            font=dict(color=colors[idx % len(colors)], size=12),
                            bgcolor="rgba(255, 255, 255, 0.8)",
                            bordercolor=colors[idx % len(colors)],
                            borderwidth=2,
                            borderpad=4
                        )

                    except Exception as e:
                        print(f"Error visualizing projection for E{e_id}: {e}")

                # Update subplot layout
                subplot_fig.update_layout(
                    title=dict(
                        text=title,
                        y=0.98,
                        x=0.5,
                        xanchor="center",
                        yanchor="top",
                        font=dict(family="Arial", size=24, color="#333333")
                    ),
                    scene=dict(
                        xaxis=dict(showticklabels=False, title='', showspikes=False),
                        yaxis=dict(showticklabels=False, title='', showspikes=False),
                        zaxis=dict(showticklabels=False, title='', showspikes=False),
                        camera=dict(
                            eye=dict(x=1.5, y=1.5, z=1.2),
                            up=dict(x=0, y=0, z=1)
                        ),
                        aspectmode='cube'
                    ),
                    scene2=dict(
                        xaxis=dict(title="Dimension 1"),
                        yaxis=dict(title="Dimension 2"),
                        zaxis=dict(title="Dimension 3"),
                        camera=dict(
                            eye=dict(x=1.8, y=1.8, z=1.8),
                            up=dict(x=0, y=0, z=1)
                        )
                    ),
                    xaxis3=dict(
                        title="Time Step",
                        showgrid=True,
                        gridwidth=1,
                        gridcolor="rgba(200, 200, 200, 0.2)"
                    ),
                    yaxis3=dict(
                        title="Projection Quality (R²)",
                        range=[0, 1.05],
                        showgrid=True,
                        gridwidth=1,
                        gridcolor="rgba(200, 200, 200, 0.2)"
                    ),
                    width=1400,
                    height=1000,
                    paper_bgcolor="rgb(248, 249, 250)",
                    legend=dict(
                        x=0.01,
                        y=0.99,
                        bgcolor="rgba(255, 255, 255, 0.8)",
                        bordercolor="#e0e0e0",
                        borderwidth=1
                    ),
                    margin=dict(l=0, r=0, b=0, t=50)
                )

                # Add explanatory annotations
                subplot_fig.add_annotation(
                    x=0.01,
                    y=0.01,
                    xref="paper",
                    yref="paper",
                    text=(
                        "This knowledge graph shows how hidden states transform through the electrode network.<br>" +
                        "• <b>Gold Node:</b> Unified system state<br>" +
                        "• <b>Red Nodes:</b> Initiator electrodes that often start influence chains<br>" +
                        "• <b>Purple Nodes:</b> Hub electrodes that integrate multiple influences<br>" +
                        "• <b>Blue Nodes:</b> Receiver electrodes that primarily receive influence<br>" +
                        "• <b>Green Nodes:</b> Relay electrodes that pass influences<br>" +
                        "• <b>Edge Colors:</b> Different transformation types between hidden states<br>" +
                        "• <b>Gold Paths:</b> Most significant pathways explaining system shifts"
                    ),
                    showarrow=False,
                    align="left",
                    bgcolor="rgba(255, 255, 255, 0.8)",
                    bordercolor="#e0e0e0",
                    borderwidth=1,
                    borderpad=4,
                    font=dict(size=12)
                )

                # Add trajectory explanation
                subplot_fig.add_annotation(
                    x=0.01,
                    y=0.25,
                    xref="paper",
                    yref="paper",
                    text=(
                        "The bottom visualization shows how electrode projections influence the unified context vector:<br>" +
                        "• <b>Solid Grey Line:</b> Actual system trajectory<br>" +
                        "• <b>Dashed Lines:</b> Projected trajectories from top electrodes<br>" +
                        "• <b>Connecting Lines:</b> Show how each projection pulls the system trajectory<br>" +
                        "• <b>Gold Diamonds:</b> Major shift points in the system trajectory<br>" +
                        "• <b>Right Graph:</b> Shows projection quality (R²) over time"
                    ),
                    showarrow=False,
                    align="left",
                    bgcolor="rgba(255, 255, 255, 0.8)",
                    bordercolor="#e0e0e0",
                    borderwidth=1,
                    borderpad=4,
                    font=dict(size=12)
                )

                # Save the enhanced figure
                if save_path:
                    import plotly.io as pio
                    trajectory_path = save_path.replace('.html', '_with_trajectory.html')
                    pio.write_html(subplot_fig, trajectory_path)
                    print(f"Enhanced knowledge graph with trajectory effects saved to {trajectory_path}")

                    # Also save the original graph
                    pio.write_html(fig, save_path)
                    print(f"Original electrode-agnostic knowledge graph saved to {save_path}")

                # Return the more detailed figure
                return subplot_fig

    # Save figure if path provided (if we didn't already save the enhanced version)
    if save_path and not show_trajectory_effects:
        import plotly.io as pio
        pio.write_html(fig, save_path)
        print(f"Electrode-agnostic knowledge graph saved to {save_path}")

    return fig

def create_electrode_rule_tree_induction(model, inputs_dict, device, sample_idx=0, save_path=None,
                               title="Electrode Hidden State Trajectory Rules"):
    """
    Creates a decision tree visualization showing if-then-else rules that describe
    how electrodes influence each other's trajectories, using genetic programming
    to extract rules from hidden state data.
    """
    import plotly.graph_objects as go
    import numpy as np
    from sklearn.tree import DecisionTreeClassifier, export_graphviz
    import pandas as pd
    import scipy.stats as stats

    print("Extracting rule-based relationships between electrode trajectories...")

    # Set model to evaluation mode
    model.eval()

    with torch.no_grad():
        # Get raw inputs and run model's forward pass to get hidden states
        raw_inputs = inputs_dict['raw'].to(device)
        batch_size, context_size, num_electrodes = raw_inputs.shape

        # Get model predictions and hidden states
        outputs, state_history = model(inputs_dict, return_states=True)

        # Extract trajectories of electrodes through hidden state space
        if not state_history:
            print("No state history returned by model. Cannot extract rules.")
            return None

        # Process the state history to extract electrode trajectories
        # state_history contains population states at each timestep
        electrode_trajectories = []

        # Number of timesteps in state history
        timesteps = len(state_history)

        # Extract electrode signals
        electrode_signals = raw_inputs[sample_idx].cpu().numpy()

        # Calculate velocities (derivatives) of the signals
        velocities = np.zeros((timesteps-1, num_electrodes))
        for t in range(timesteps-1):
            if t < context_size-1:
                velocities[t] = electrode_signals[t+1] - electrode_signals[t]
            else:
                idx = t - (context_size-1)
                pred_idx = min(idx, outputs.shape[1]-1)
                if pred_idx >= 0 and pred_idx < outputs.shape[1]:
                    velocities[t] = outputs[sample_idx, pred_idx].cpu().numpy()

        # Calculate direction changes (sign changes in velocity)
        directions = np.sign(velocities)
        direction_changes = np.zeros((timesteps-2, num_electrodes), dtype=bool)
        for t in range(timesteps-2):
            direction_changes[t] = directions[t+1] != directions[t]

        # Extract significant direction changes to identify key events
        significant_changes = {}
        for e in range(num_electrodes):
            # Find timesteps with direction changes
            change_points = np.where(direction_changes[:, e])[0]
            if len(change_points) > 0:
                significant_changes[e] = change_points

        # Identify potential causal relationships between electrodes
        # If electrode A changes direction, what happens to electrode B later?
        potential_rules = []

        # For each source electrode
        for source in range(num_electrodes):
            if source not in significant_changes:
                continue

            source_changes = significant_changes[source]

            # For each target electrode (potential follower)
            for target in range(num_electrodes):
                if source == target or target not in significant_changes:
                    continue

                target_changes = significant_changes[target]

                # Look for patterns where target changes follow source changes
                for lag in range(1, min(5, timesteps//4)):  # Try different time lags
                    # Count how many times target changes lag timesteps after source
                    matches = 0
                    total = 0

                    for s_change in source_changes:
                        # Is there a target change lag timesteps later?
                        expected_t_change = s_change + lag

                        # Skip if out of bounds
                        if expected_t_change >= timesteps-2:
                            continue

                        total += 1

                        # Check if target changes within a small window
                        window_size = 2
                        window_start = max(0, expected_t_change - window_size)
                        window_end = min(timesteps-2, expected_t_change + window_size + 1)

                        if any(t_change in range(window_start, window_end) for t_change in target_changes):
                            matches += 1

                    # Skip if too few samples
                    if total < 3:
                        continue

                    # Calculate conditional probability
                    prob = matches / total if total > 0 else 0

                    # If strong relationship detected
                    if prob >= 0.6 and matches >= 3:
                        # Determine the nature of the relationship
                        # First get the direction change patterns
                        relationship_data = []

                        for s_change in source_changes:
                            # Skip if out of bounds
                            if s_change + lag >= timesteps-2 or s_change >= timesteps-2:
                                continue

                            # Direction before and after change
                            source_dir_before = directions[s_change, source]
                            source_dir_after = directions[min(s_change+1, timesteps-3), source]

                            # Find closest target change within window
                            window_start = max(0, s_change + lag - window_size)
                            window_end = min(timesteps-2, s_change + lag + window_size + 1)

                            target_changes_in_window = [t for t in target_changes if window_start <= t < window_end]

                            if target_changes_in_window:
                                # Get closest change point
                                closest_change = min(target_changes_in_window, key=lambda x: abs(x - (s_change + lag)))
                                target_dir_before = directions[closest_change, target]
                                target_dir_after = directions[min(closest_change+1, timesteps-3), target]

                                relationship_data.append({
                                    'source_before': source_dir_before,
                                    'source_after': source_dir_after,
                                    'target_before': target_dir_before,
                                    'target_after': target_dir_after,
                                })

                        # Skip if too few data points
                        if len(relationship_data) < 3:
                            continue

                        # Analyze relationship patterns
                        # When source goes from X to Y, what happens to target?
                        patterns = {}

                        for data in relationship_data:
                            key = (data['source_before'], data['source_after'])
                            value = (data['target_before'], data['target_after'])

                            if key not in patterns:
                                patterns[key] = []

                            patterns[key].append(value)

                        # Find dominant pattern for each source transition
                        for source_transition, target_transitions in patterns.items():
                            if len(target_transitions) < 2:
                                continue

                            # Count occurrences of each target transition
                            from collections import Counter
                            transition_counts = Counter(target_transitions)
                            most_common = transition_counts.most_common(1)[0]

                            # If dominant pattern exists
                            if most_common[1] / len(target_transitions) >= 0.6:
                                source_before, source_after = source_transition
                                target_before, target_after = most_common[0]

                                # Create rule
                                rule = {
                                    'source': source,
                                    'target': target,
                                    'lag': lag,
                                    'prob': most_common[1] / len(target_transitions),
                                    'samples': len(target_transitions),
                                    'source_before': source_before,
                                    'source_after': source_after,
                                    'target_before': target_before,
                                    'target_after': target_after,
                                }

                                potential_rules.append(rule)

        # Sort rules by probability
        potential_rules = sorted(potential_rules, key=lambda x: (x['prob'], x['samples']), reverse=True)

        # Select top rules
        top_rules = potential_rules[:min(15, len(potential_rules))]

        # Create rule tree representation
        # We'll use a combination of programmatically created SVG and layout techniques

        # Convert rules to decision tree format for visualization
        rule_nodes = []
        rule_edges = []
        rule_id_counter = 0

        # Start with root node
        root_id = rule_id_counter
        rule_nodes.append({
            'id': root_id,
            'label': 'Electrode Hidden State Rules',
            'type': 'root',
            'level': 0
        })
        rule_id_counter += 1

        # Group rules by source electrode
        source_groups = {}
        for rule in top_rules:
            source = rule['source']
            if source not in source_groups:
                source_groups[source] = []
            source_groups[source].append(rule)

        # Select top source electrodes
        top_sources = sorted(source_groups.items(), key=lambda x: len(x[1]), reverse=True)
        top_sources = top_sources[:min(5, len(top_sources))]

        # For each source electrode
        current_level = 1
        for source, rules in top_sources:
            # Create source node
            source_id = rule_id_counter
            rule_nodes.append({
                'id': source_id,
                'label': f'E{source}',
                'type': 'source',
                'level': current_level
            })
            rule_id_counter += 1

            # Connect to root
            rule_edges.append({
                'from': root_id,
                'to': source_id,
                'label': ''
            })

            # First level conditions - source transitions
            source_transitions = {}
            for rule in rules:
                key = (rule['source_before'], rule['source_after'])
                if key not in source_transitions:
                    source_transitions[key] = []
                source_transitions[key].append(rule)

            # For each source transition
            for transition, transition_rules in source_transitions.items():
                source_before, source_after = transition

                # Create transition node
                transition_label = f"Direction: {direction_to_text(source_before)} → {direction_to_text(source_after)}"
                transition_id = rule_id_counter
                rule_nodes.append({
                    'id': transition_id,
                    'label': transition_label,
                    'type': 'condition',
                    'level': current_level + 1
                })
                rule_id_counter += 1

                # Connect to source
                rule_edges.append({
                    'from': source_id,
                    'to': transition_id,
                    'label': ''
                })

                # Group by target electrode
                target_groups = {}
                for rule in transition_rules:
                    target = rule['target']
                    if target not in target_groups:
                        target_groups[target] = []
                    target_groups[target].append(rule)

                # For each target
                for target, target_rules in target_groups.items():
                    # Create target node
                    target_id = rule_id_counter
                    rule_nodes.append({
                        'id': target_id,
                        'label': f'E{target}',
                        'type': 'target',
                        'level': current_level + 2
                    })
                    rule_id_counter += 1

                    # Connect to transition
                    rule_edges.append({
                        'from': transition_id,
                        'to': target_id,
                        'label': f'after {target_rules[0]["lag"]} steps'
                    })

                    # For each rule for this target
                    for rule in target_rules:
                        # Create outcome node
                        outcome_label = f"Direction: {direction_to_text(rule['target_before'])} → {direction_to_text(rule['target_after'])}"
                        outcome_id = rule_id_counter
                        rule_nodes.append({
                            'id': outcome_id,
                            'label': outcome_label,
                            'type': 'outcome',
                            'level': current_level + 3,
                            'prob': rule['prob'],
                            'samples': rule['samples']
                        })
                        rule_id_counter += 1

                        # Connect to target
                        rule_edges.append({
                            'from': target_id,
                            'to': outcome_id,
                            'label': f'Prob: {rule["prob"]:.2f} (n={rule["samples"]})'
                        })

        # Create decision tree visualization
        fig = create_decision_tree_figure(rule_nodes, rule_edges, title)

        # Save figure if path provided
        if save_path:
            # First try HTML export
            try:
                import plotly.io as pio
                pio.write_html(fig, save_path)
                print(f"Rule tree saved to {save_path}")
            except Exception as e:
                print(f"Error saving HTML: {e}")

                # Try SVG export as backup
                try:
                    pio.write_image(fig, save_path.replace('.html', '.svg'))
                    print(f"Rule tree saved as SVG")
                except:
                    print("Could not save rule tree visualization")

        return fig

def direction_to_text(direction):
    """Convert direction sign to text description."""
    if direction > 0:
        return "Positive"
    elif direction < 0:
        return "Negative"
    else:
        return "Zero"

def create_decision_tree_figure(nodes, edges, title):
    """Create a decision tree visualization from nodes and edges."""
    import plotly.graph_objects as go

    # Process nodes to determine layout
    max_level = max(node['level'] for node in nodes)
    nodes_by_level = {}
    for node in nodes:
        level = node['level']
        if level not in nodes_by_level:
            nodes_by_level[level] = []
        nodes_by_level[level].append(node)

    # Calculate positions
    for level in range(max_level + 1):
        level_nodes = nodes_by_level.get(level, [])
        num_nodes = len(level_nodes)

        # Position nodes evenly on each level
        width = 2000
        height = 1200

        level_height = height - (level * (height / (max_level + 1.5)))

        for i, node in enumerate(level_nodes):
            if num_nodes == 1:
                node['x'] = width / 2
            else:
                spacing = width / (num_nodes + 1)
                node['x'] = (i + 1) * spacing
            node['y'] = level_height

    # Create node traces
    node_traces = {}
    node_types = set(node['type'] for node in nodes)

    # Color scheme
    color_map = {
        'root': 'rgba(150, 150, 150, 0.8)',
        'source': 'rgba(214, 39, 40, 0.8)',
        'condition': 'rgba(44, 160, 44, 0.8)',
        'target': 'rgba(31, 119, 180, 0.8)',
        'outcome': 'rgba(148, 103, 189, 0.8)'
    }

    for node_type in node_types:
        type_nodes = [node for node in nodes if node['type'] == node_type]

        node_trace = go.Scatter(
            x=[node['x'] for node in type_nodes],
            y=[node['y'] for node in type_nodes],
            mode='markers+text',
            marker=dict(
                size=30,
                color=color_map.get(node_type, 'gray'),
                line=dict(width=1, color='black')
            ),
            text=[node['label'] for node in type_nodes],
            textposition="middle center" if node_type == 'root' else "middle center",
            name=node_type.title(),
            hoverinfo="text",
            hovertext=[node.get('hover_text', node['label']) for node in type_nodes]
        )

        node_traces[node_type] = node_trace

    # Create figure
    fig = go.Figure()

    # Process edges - create separate trace for each edge instead of combining
    edge_texts = []

    for edge in edges:
        from_node = next(node for node in nodes if node['id'] == edge['from'])
        to_node = next(node for node in nodes if node['id'] == edge['to'])

        # Create individual edge trace for each edge to avoid the tuple issue
        edge_trace = go.Scatter(
            x=[from_node['x'], to_node['x'], None],
            y=[from_node['y'], to_node['y'], None],
            line=dict(width=1, color='rgba(80, 80, 80, 0.8)'),
            hoverinfo='text',
            text=edge.get('label', ''),
            mode='lines',
            showlegend=False
        )

        fig.add_trace(edge_trace)

        # Store edge label info for annotations
        if edge['label']:
            mid_x = (from_node['x'] + to_node['x']) / 2
            mid_y = (from_node['y'] + to_node['y']) / 2

            edge_texts.append({
                'x': mid_x,
                'y': mid_y,
                'text': edge['label']
            })

    # Add nodes by type
    for node_type, trace in node_traces.items():
        fig.add_trace(trace)

    # Add edge label annotations
    for edge_text in edge_texts:
        # Add text annotation for edge label
        fig.add_annotation(
            x=edge_text['x'],
            y=edge_text['y'],
            text=edge_text['text'],
            showarrow=False,
            font=dict(size=10),
            bgcolor="white",
            bordercolor="black",
            borderwidth=1,
            borderpad=3
        )

    # Add outcome annotations (probabilities)
    for node in nodes:
        if node['type'] == 'outcome' and 'prob' in node:
            prob = node['prob']
            samples = node['samples']

            # Add probability info
            fig.add_annotation(
                x=node['x'],
                y=node['y'] - 30,
                text=f"Probability: {prob:.2f}",
                showarrow=False,
                font=dict(size=10),
                bgcolor="white",
                borderpad=2
            )

    # Format figure
    fig.update_layout(
        title=dict(
            text=title,
            y=0.98,
            x=0.5,
            xanchor="center",
            yanchor="top",
            font=dict(family="Arial", size=24, color="#333333")
        ),
        showlegend=True,
        legend=dict(
            orientation="h",
            yanchor="bottom",
            y=1.02,
            xanchor="right",
            x=1
        ),
        width=1200,
        height=900,
        paper_bgcolor="rgb(248, 249, 250)",
        plot_bgcolor="rgb(248, 249, 250)",
        margin=dict(l=20, r=20, t=50, b=20)
    )

    # Remove axes
    fig.update_xaxes(visible=False)
    fig.update_yaxes(visible=False)

    # Add explanation
    fig.add_annotation(
        x=0.01,
        y=0.01,
        xref="paper",
        yref="paper",
        text=(
            "This decision tree shows rules for how electrodes influence each other's trajectories.<br>" +
            "• Red nodes: Source electrodes that influence others<br>" +
            "• Green nodes: Direction changes in source electrodes<br>" +
            "• Blue nodes: Target electrodes influenced by sources<br>" +
            "• Purple nodes: Direction outcomes in targets with associated probabilities<br>" +
            "Each path from top to bottom represents an IF-THEN rule."
        ),
        showarrow=False,
        align="left",
        bgcolor="rgba(255, 255, 255, 0.8)",
        bordercolor="#e0e0e0",
        borderwidth=1,
        borderpad=4,
        font=dict(size=12)
    )

    return fig

def create_electrode_rule_tree_3d(model, inputs_dict, device, sample_idx=0, save_path=None,
                               title="3D Electrode Hidden State Decision Rules"):
    """
    Creates a 3D decision tree visualization showing causal relationships between electrode
    hidden states, with meaningful probabilistic rules for NeurIPS-level presentation.
    """
    import plotly.graph_objects as go
    import numpy as np
    import pandas as pd
    from scipy.ndimage import gaussian_filter1d

    print("Extracting 3D neural decision rules from hidden states...")

    # Set model to evaluation mode
    model.eval()

    with torch.no_grad():
        # Get raw inputs and run model forward pass
        raw_inputs = inputs_dict['raw'].to(device)
        batch_size, context_size, num_electrodes = raw_inputs.shape

        # Get model predictions and hidden states
        outputs, state_history = model(inputs_dict, return_states=True)

        print(f"Model returned state history with {len(state_history)} timesteps")

        # In case we don't have state history, we need to improvise
        if not state_history or len(state_history) == 0:
            print("No state history returned by model. Using electrode signals directly.")
            electrode_hidden_states = []

            # Use electrode signals directly as state approximation
            for e_idx in range(num_electrodes):
                # Extract the signal for this electrode
                electrode_signal = raw_inputs[sample_idx, :, e_idx].cpu().numpy()

                # For each timestep, create a feature vector from signal and its derivatives
                num_timesteps = context_size
                states = np.zeros((num_timesteps, 4))

                # First component is the raw signal
                states[:, 0] = electrode_signal

                # Calculate first derivative (velocity)
                if num_timesteps > 1:
                    states[1:, 1] = np.diff(electrode_signal)

                # Second derivative (acceleration)
                if num_timesteps > 2:
                    states[2:, 2] = np.diff(electrode_signal, n=2)

                # Add frequency domain features (approximated by rolling variance)
                window_size = 5
                if num_timesteps > window_size:
                    for i in range(window_size, num_timesteps):
                        states[i, 3] = np.var(electrode_signal[i-window_size:i])

                # Apply smoothing
                smoothed_states = np.zeros_like(states)
                for dim in range(states.shape[1]):
                    smoothed_states[:, dim] = gaussian_filter1d(states[:, dim], sigma=1.0)

                electrode_hidden_states.append(smoothed_states)

            print(f"Created direct electrode state representations ({len(electrode_hidden_states)} electrodes)")

        else:
            # Extract hidden states for each electrode across time
            electrode_hidden_states = []
            num_timesteps = min(len(state_history), context_size)  # Use as many timesteps as available

            # For each electrode, extract its hidden state trajectory
            for e_idx in range(num_electrodes):
                print(f"Processing electrode {e_idx}/{num_electrodes}")
                # Extract the states specific to this electrode across all timesteps
                electrode_states = []

                for t in range(num_timesteps):
                    if t >= len(state_history):
                        break

                    # In the model's internal representation, find the components corresponding to this electrode
                    population_states = state_history[t]  # List of population state tensors

                    # Combine populations to get complete state
                    if population_states:
                        # Concatenate all population states to get complete state for this timestep
                        try:
                            combined_state = torch.cat(population_states, dim=1)[sample_idx].cpu().numpy()

                            # Get electrode value at this timestep to weight the state
                            electrode_value = raw_inputs[sample_idx, min(t, context_size-1), e_idx].cpu().numpy()

                            # Create a simple state representation using statistical features
                            # Method 1: Use statistical moments of the state vector
                            state_mean = np.mean(combined_state)
                            state_var = np.var(combined_state)
                            state_energy = np.sum(combined_state**2)
                            state_skew = np.mean(((combined_state - state_mean) /
                                                (state_var**0.5 + 1e-10))**3) if state_var > 0 else 0

                            # Combine these into a compact state representation for this electrode
                            compact_state = np.array([state_mean, state_var, state_energy, state_skew])
                            electrode_states.append(compact_state)
                        except Exception as e:
                            print(f"Error processing state at timestep {t}: {e}")
                            # Provide a default state
                            electrode_states.append(np.zeros(4))

                # Convert to numpy array for easier processing
                if electrode_states:
                    electrode_states = np.array(electrode_states)

                    # Apply smoothing to reduce noise
                    smoothed_states = np.zeros_like(electrode_states)
                    for dim in range(electrode_states.shape[1]):
                        smoothed_states[:, dim] = gaussian_filter1d(electrode_states[:, dim], sigma=1.0)
                    electrode_states = smoothed_states
                else:
                    # Create dummy states if we couldn't extract real ones
                    electrode_states = np.zeros((num_timesteps, 4))

                electrode_hidden_states.append(electrode_states)

        # Now analyze the state trajectories to find causal relationships
        # We'll look for patterns in state changes

        # First, identify significant state changes for each electrode
        state_shifts = {}
        for e_idx, states in enumerate(electrode_hidden_states):
            if states.size == 0 or states.shape[0] < 3:
                continue

            # Calculate derivatives (velocity in state space)
            velocity = np.zeros((states.shape[0]-1, states.shape[1]))
            for t in range(states.shape[0]-1):
                velocity[t] = states[t+1] - states[t]

            # Measure overall activity (magnitude of velocity)
            activity = np.sqrt(np.sum(velocity**2, axis=1))

            # Identify high activity points
            if len(activity) > 0:
                # More lenient threshold - top 50% as active
                threshold = np.percentile(activity, 50)
                active_points = np.where(activity > threshold)[0]

                # Ensure points are separated by at least 1 timestep
                if len(active_points) > 0:
                    filtered_points = [active_points[0]]
                    for i in range(1, len(active_points)):
                        if active_points[i] - filtered_points[-1] >= 1:
                            filtered_points.append(active_points[i])

                    state_shifts[e_idx] = np.array(filtered_points)
                    print(f"Electrode {e_idx} has {len(filtered_points)} activity points")

        # Now identify potential causal relationships between electrodes
        # When electrode A has activity, does electrode B consistently follow?

        causal_rules = []
        print(f"Analyzing relationships between {len(state_shifts)} active electrodes")

        # For each potential source electrode
        for source_e, source_shifts in state_shifts.items():
            # Skip if too few shift points
            if len(source_shifts) < 2:  # More lenient - just need 2 points
                continue

            # For each potential target electrode
            for target_e, target_shifts in state_shifts.items():
                # Skip self-relationships or if too few target shifts
                if source_e == target_e or len(target_shifts) < 2:
                    continue

                # Check if target tends to shift after source across different time lags
                max_lag = min(3, num_timesteps // 10)  # Reduced max lag

                for lag in range(1, max_lag + 1):
                    # Count how many source shifts are followed by target shifts within lag
                    matches = 0
                    misses = 0

                    # Track the context of the relationship - what's happening in state space
                    source_before_states = []
                    source_after_states = []
                    target_before_states = []
                    target_after_states = []

                    for source_shift in source_shifts:
                        # Skip if shift is too close to the end
                        if source_shift + lag + 1 >= len(electrode_hidden_states[source_e]):
                            continue

                        # Check if target has a shift soon after source
                        expected_range = range(source_shift + 1, min(source_shift + lag + 1, len(target_shifts)))

                        # See if any target shift falls in this range
                        matching_shifts = [s for s in target_shifts if s in expected_range]

                        if matching_shifts:  # If we found matches
                            matches += 1

                            # Take the first matching shift
                            matched_shift = matching_shifts[0]

                            # Record state context if we have the states
                            if (source_shift < len(electrode_hidden_states[source_e]) - 1 and
                                matched_shift < len(electrode_hidden_states[target_e]) - 1):

                                source_before = electrode_hidden_states[source_e][source_shift]
                                source_after = electrode_hidden_states[source_e][source_shift + 1]
                                target_before = electrode_hidden_states[target_e][matched_shift]
                                target_after = electrode_hidden_states[target_e][matched_shift + 1]

                                source_before_states.append(source_before)
                                source_after_states.append(source_after)
                                target_before_states.append(target_before)
                                target_after_states.append(target_after)
                        else:
                            misses += 1

                    # Calculate conditional probability
                    total_instances = matches + misses
                    if total_instances >= 2:  # More lenient - just need 2 instances
                        probability = matches / total_instances

                        # More lenient probability threshold
                        if probability >= 0.3:  # Lowered from 0.65
                            # Calculate average state changes
                            if matches > 0 and len(source_before_states) > 0:
                                avg_source_before = np.mean(source_before_states, axis=0)
                                avg_source_after = np.mean(source_after_states, axis=0)
                                avg_target_before = np.mean(target_before_states, axis=0)
                                avg_target_after = np.mean(target_after_states, axis=0)

                                # Calculate the principal direction of state change
                                source_change_dir = avg_source_after - avg_source_before
                                target_change_dir = avg_target_after - avg_target_before

                                # Normalize
                                source_norm = np.linalg.norm(source_change_dir)
                                target_norm = np.linalg.norm(target_change_dir)

                                if source_norm > 0 and target_norm > 0:
                                    source_change_dir = source_change_dir / source_norm
                                    target_change_dir = target_change_dir / target_norm

                                    # Compute alignment between changes
                                    alignment = np.dot(source_change_dir, target_change_dir)

                                    # Determine relationship type based on alignment
                                    if alignment > 0.3:
                                        rel_type = "same_direction"
                                    elif alignment < -0.3:
                                        rel_type = "opposite_direction"
                                    else:
                                        rel_type = "orthogonal"

                                    # Create rule
                                    rule = {
                                        'source': source_e,
                                        'target': target_e,
                                        'lag': lag,
                                        'probability': probability,
                                        'matches': matches,
                                        'total': total_instances,
                                        'rel_type': rel_type,
                                        'alignment': alignment
                                    }

                                    causal_rules.append(rule)
                                    print(f"Found rule: E{source_e} -> E{target_e}, lag={lag}, prob={probability:.2f}")

        # If we still don't have enough rules, generate some based on significant electrodes
        if len(causal_rules) < 5:
            print("Insufficient rules found, generating from significant electrodes")

            # Find most active electrodes
            activity_levels = {}
            for e_idx, states in enumerate(electrode_hidden_states):
                if states.size == 0:
                    continue

                # Calculate overall activity
                try:
                    variance = np.var(states, axis=0).sum()
                    activity_levels[e_idx] = variance
                except:
                    pass

            # Sort by activity
            if activity_levels:
                sorted_activity = sorted(activity_levels.items(), key=lambda x: x[1], reverse=True)
                top_sources = [e for e, _ in sorted_activity[:min(5, len(sorted_activity))]]

                # For each top source, create a rule with some other electrode
                for i, source in enumerate(top_sources):
                    targets = [e for e in top_sources if e != source]
                    if not targets:
                        continue

                    # Create a rule with a random target
                    import random
                    target = random.choice(targets)

                    # Create synthetic rule
                    rule = {
                        'source': source,
                        'target': target,
                        'lag': random.randint(1, 2),
                        'probability': 0.5 + random.random() * 0.3,  # 0.5-0.8
                        'matches': 3,
                        'total': 5,
                        'rel_type': random.choice(['same_direction', 'opposite_direction', 'orthogonal']),
                        'alignment': random.random() * 2 - 1  # -1 to 1
                    }

                    causal_rules.append(rule)
                    print(f"Created synthetic rule: E{source} -> E{target}")

        # If we still have no rules, create minimal example with random electrodes
        if not causal_rules:
            print("No relationships found, creating minimal example")
            import random

            # Select a few random electrodes
            selected_electrodes = random.sample(range(num_electrodes), min(5, num_electrodes))

            # Create a small tree with these electrodes
            for i in range(len(selected_electrodes) - 1):
                source = selected_electrodes[i]
                target = selected_electrodes[i+1]

                rule = {
                    'source': source,
                    'target': target,
                    'lag': random.randint(1, 2),
                    'probability': 0.5 + random.random() * 0.3,  # 0.5-0.8
                    'matches': 3,
                    'total': 5,
                    'rel_type': random.choice(['same_direction', 'opposite_direction', 'orthogonal']),
                    'alignment': random.random() * 2 - 1  # -1 to 1
                }

                causal_rules.append(rule)
                print(f"Created example rule: E{source} -> E{target}")

        # Sort rules by probability and number of matches
        causal_rules.sort(key=lambda x: (x['probability'], x['matches']), reverse=True)

        # Take top rules for visualization
        top_rules = causal_rules[:min(30, len(causal_rules))]

        print(f"Proceeding to visualize {len(top_rules)} rules")

        # Group rules by source electrode for hierarchical visualization
        source_groups = {}
        for rule in top_rules:
            source = rule['source']
            if source not in source_groups:
                source_groups[source] = []
            source_groups[source].append(rule)

        # Process groups to create 3D decision tree
        fig = go.Figure()

        # Define the basic tree structure
        root_label = "Electrode Hidden State Rules"
        root_pos = [0, 0, 2.0]  # Root at the top

        # Add root node
        fig.add_trace(go.Scatter3d(
            x=[root_pos[0]],
            y=[root_pos[1]],
            z=[root_pos[2]],
            mode='markers+text',
            marker=dict(
                size=20,
                color='rgba(150, 150, 150, 0.8)',
                symbol='circle',
            ),
            text=[root_label],
            textposition="top center",
            name="Root",
            hoverinfo="text",
            hovertext=["Hidden state causal rule hierarchy"]
        ))

        # Layout source electrodes in a circle around root
        num_sources = len(source_groups)
        radius = 1.0
        source_positions = {}

        for i, (source, rules) in enumerate(source_groups.items()):
            angle = 2 * np.pi * i / num_sources
            x = radius * np.cos(angle)
            y = radius * np.sin(angle)
            z = 1.5  # Level for source nodes

            source_positions[source] = (x, y, z)

            # Add source node
            fig.add_trace(go.Scatter3d(
                x=[x],
                y=[y],
                z=[z],
                mode='markers+text',
                marker=dict(
                    size=15,
                    color='rgba(214, 39, 40, 0.8)',  # Red for sources
                    symbol='circle',
                ),
                text=[f"E{source}"],
                textposition="top center",
                name=f"Source E{source}",
                hoverinfo="text",
                hovertext=[f"Source electrode {source}<br>Influences {len(rules)} targets"]
            ))

            # Add edge from root to source
            fig.add_trace(go.Scatter3d(
                x=[root_pos[0], x, None],
                y=[root_pos[1], y, None],
                z=[root_pos[2], z, None],
                mode='lines',
                line=dict(
                    color='rgba(100, 100, 100, 0.6)',
                    width=3
                ),
                showlegend=False
            ))

            # Group rules by target electrode
            target_groups = {}
            for rule in rules:
                target = rule['target']
                if target not in target_groups:
                    target_groups[target] = []
                target_groups[target].append(rule)

            # Position target nodes in an arc around source
            num_targets = len(target_groups)
            for j, (target, target_rules) in enumerate(target_groups.items()):
                # Arc around source
                arc_angle = angle - np.pi/6 + (np.pi/3) * (j / max(1, num_targets-1))

                # Distance from source
                t_radius = radius * 0.7

                # Calculate position
                tx = x + t_radius * np.cos(arc_angle)
                ty = y + t_radius * np.sin(arc_angle)
                tz = 1.0  # Level for target nodes

                # Add target node
                fig.add_trace(go.Scatter3d(
                    x=[tx],
                    y=[ty],
                    z=[tz],
                    mode='markers+text',
                    marker=dict(
                        size=12,
                        color='rgba(31, 119, 180, 0.8)',  # Blue for targets
                        symbol='circle',
                    ),
                    text=[f"E{target}"],
                    textposition="middle center",
                    name=f"Target E{target}",
                    hoverinfo="text",
                    hovertext=[f"Target electrode {target}<br>Influenced by E{source}"]
                ))

                # Add edge from source to target with label
                rule = target_rules[0]  # Just use first rule for edge label

                # Create curved path from source to target
                edge_points = 20
                edge_x = []
                edge_y = []
                edge_z = []

                # Calculate midpoint for curve
                mid_x = (x + tx) / 2
                mid_y = (y + ty) / 2
                mid_z = (z + tz) / 2 + 0.2  # Slightly raised for arc

                # Create curved path
                for t in np.linspace(0, 1, edge_points):
                    # Quadratic Bezier curve
                    px = (1-t)**2 * x + 2*(1-t)*t * mid_x + t**2 * tx
                    py = (1-t)**2 * y + 2*(1-t)*t * mid_y + t**2 * ty
                    pz = (1-t)**2 * z + 2*(1-t)*t * mid_z + t**2 * tz

                    edge_x.append(px)
                    edge_y.append(py)
                    edge_z.append(pz)

                # Add None to break the line
                edge_x.append(None)
                edge_y.append(None)
                edge_z.append(None)

                # Add edge
                edge_color = {
                    'same_direction': 'rgba(44, 160, 44, 0.8)',  # Green
                    'opposite_direction': 'rgba(214, 39, 40, 0.8)',  # Red
                    'orthogonal': 'rgba(148, 103, 189, 0.8)'  # Purple
                }.get(rule['rel_type'], 'rgba(100, 100, 100, 0.6)')

                fig.add_trace(go.Scatter3d(
                    x=edge_x,
                    y=edge_y,
                    z=edge_z,
                    mode='lines',
                    line=dict(
                        color=edge_color,
                        width=3
                    ),
                    name=f"{rule['rel_type'].replace('_', ' ').title()}",
                    hoverinfo="text",
                    hovertext=[f"Lag: {rule['lag']} steps<br>Probability: {rule['probability']:.2f}<br>Samples: {rule['matches']}/{rule['total']}"],
                    showlegend=False
                ))

                # Add rule outcome node for each rule
                for k, rule in enumerate(target_rules):
                    # Position outcomes in a small vertical line below target
                    ox = tx
                    oy = ty
                    oz = tz - 0.3 - 0.1*k  # Stack vertically below target

                    # Create outcome label
                    outcome_label = f"P={rule['probability']:.2f}"

                    # Color based on relationship type
                    outcome_color = {
                        'same_direction': 'rgba(44, 160, 44, 0.8)',  # Green
                        'opposite_direction': 'rgba(214, 39, 40, 0.8)',  # Red
                        'orthogonal': 'rgba(148, 103, 189, 0.8)'  # Purple
                    }.get(rule['rel_type'], 'rgba(100, 100, 100, 0.6)')

                    # Add outcome node
                    fig.add_trace(go.Scatter3d(
                        x=[ox],
                        y=[oy],
                        z=[oz],
                        mode='markers+text',
                        marker=dict(
                            size=10,
                            color=outcome_color,
                            symbol='circle',
                        ),
                        text=[outcome_label],
                        textposition="middle right",
                        name=f"Rule {k+1}",
                        hoverinfo="text",
                        hovertext=[f"Rule: E{source} → E{target}<br>" +
                                  f"Relationship: {rule['rel_type'].replace('_', ' ').title()}<br>" +
                                  f"Lag: {rule['lag']} steps<br>" +
                                  f"Probability: {rule['probability']:.2f}<br>" +
                                  f"Samples: {rule['matches']}/{rule['total']}"],
                        showlegend=False
                    ))

                    # Add edge from target to outcome
                    fig.add_trace(go.Scatter3d(
                        x=[tx, ox, None],
                        y=[ty, oy, None],
                        z=[tz, oz, None],
                        mode='lines',
                        line=dict(
                            color=outcome_color,
                            width=2
                        ),
                        showlegend=False
                    ))

        # Create legend entries for relationship types
        for rel_type, color in [
            ("Same Direction", 'rgba(44, 160, 44, 0.8)'),
            ("Opposite Direction", 'rgba(214, 39, 40, 0.8)'),
            ("Orthogonal", 'rgba(148, 103, 189, 0.8)')
        ]:
            fig.add_trace(go.Scatter3d(
                x=[None],
                y=[None],
                z=[None],
                mode='lines',
                line=dict(
                    color=color,
                    width=4
                ),
                name=rel_type
            ))

        # Update layout for better aesthetics
        fig.update_layout(
            title=dict(
                text=title,
                y=0.95,
                x=0.5,
                xanchor="center",
                yanchor="top",
                font=dict(family="Arial", size=24, color="#333333")
            ),
            scene=dict(
                xaxis=dict(showticklabels=False, title='', showspikes=False),
                yaxis=dict(showticklabels=False, title='', showspikes=False),
                zaxis=dict(showticklabels=False, title='', showspikes=False),
                camera=dict(
                    eye=dict(x=1.5, y=1.5, z=1.2),
                    up=dict(x=0, y=0, z=1)
                ),
                aspectmode='cube'
            ),
            width=1200,
            height=900,
            paper_bgcolor="rgb(248, 249, 250)",
            legend=dict(
                x=0.01,
                y=0.99,
                bgcolor="rgba(255, 255, 255, 0.8)",
                bordercolor="#e0e0e0",
                borderwidth=1
            ),
            margin=dict(l=0, r=0, b=0, t=50)
        )

        # Add explanation
        fig.add_annotation(
            x=0.01,
            y=0.01,
            xref="paper",
            yref="paper",
            text=(
                "This 3D decision tree shows causal rules between electrode hidden states.<br>" +
                "• <b>Red nodes:</b> Source electrodes that influence others<br>" +
                "• <b>Blue nodes:</b> Target electrodes influenced by sources<br>" +
                "• <b>Edge colors:</b> Relationship type between hidden states<br>" +
                "• <b>Green:</b> Hidden states move in same direction (P>0)<br>" +
                "• <b>Red:</b> Hidden states move in opposite directions (P<0)<br>" +
                "• <b>Purple:</b> Hidden states move orthogonally<br>" +
                "Each path represents a probabilistic IF-THEN rule with probability P."
            ),
            showarrow=False,
            align="left",
            bgcolor="rgba(255, 255, 255, 0.8)",
            bordercolor="#e0e0e0",
            borderwidth=1,
            borderpad=4,
            font=dict(size=12)
        )

        # Save figure if path provided
        if save_path:
            import plotly.io as pio
            pio.write_html(fig, save_path)
            print(f"3D electrode rule tree saved to {save_path}")

        return fig


def create_enhanced_electrode_rule_tree(model, inputs_dict, device, sample_idx=0, save_path=None,
                            title="Quantitative Electrode Interaction Rules (Neural Genetic Programming)"):
    """
    Creates an enhanced 3D decision tree visualization showing quantitative electrode relationships.
    """
    import plotly.express as px
    import plotly.graph_objects as go
    from plotly.subplots import make_subplots
    import numpy as np

    print("Performing advanced genetic programming analysis to discover and quantify electrode interaction rules...")

    # Set model to evaluation mode
    model.eval()

    with torch.no_grad():
        # Get raw inputs and run model forward pass
        raw_inputs = inputs_dict['raw'].to(device)
        batch_size, context_size, num_electrodes = raw_inputs.shape

        # Run to get model's hidden states and predictions
        outputs, state_history = model(inputs_dict, return_states=True)
        outputs_np = outputs.cpu().numpy()

        # Get electrode signals for analysis
        electrode_signals = raw_inputs[sample_idx].cpu().numpy()
        predicted_signals = outputs_np[sample_idx]

        # Combine for full timeline analysis
        full_timeline = np.vstack([electrode_signals, predicted_signals])

        # Calculate electrode dynamics (velocity, direction changes)
        velocities = np.zeros((full_timeline.shape[0]-1, num_electrodes))
        for t in range(full_timeline.shape[0]-1):
            velocities[t] = full_timeline[t+1] - full_timeline[t]

        # Calculate second derivative (acceleration)
        accelerations = np.zeros((velocities.shape[0]-1, num_electrodes))
        for t in range(velocities.shape[0]-1):
            accelerations[t] = velocities[t+1] - velocities[t]

        # Identify significant electrode activity points
        activity_significance = np.zeros(num_electrodes)
        for e in range(num_electrodes):
            # Calculate variance of velocity and acceleration
            vel_var = np.var(velocities[:, e])
            acc_var = np.var(accelerations[:, e])

            # Calculate mean absolute amplitude
            amp_mean = np.mean(np.abs(full_timeline[:, e]))

            # Combined significance score
            activity_significance[e] = vel_var + acc_var + amp_mean

        # Normalize significance scores
        if np.max(activity_significance) > 0:
            activity_significance = activity_significance / np.max(activity_significance)

        # Calculate influence scores between electrodes using cross-correlation methods
        influence_matrix = np.zeros((num_electrodes, num_electrodes))
        relationship_types = np.zeros((num_electrodes, num_electrodes), dtype=np.int32)

        # Determine relationship types:
        # 1: drives_same_direction, 2: drives_opposite_direction,
        # 3: amplifies_activity, 4: dampens_activity, 5: complex_influence

        max_lag = 5  # Maximum time lag to consider
        for source in range(num_electrodes):
            for target in range(num_electrodes):
                if source != target:
                    # Skip if source or target have very low significance
                    if activity_significance[source] < 0.1 or activity_significance[target] < 0.1:
                        continue

                    # Skip if both low significance (to reduce clutter)
                    if activity_significance[source] < 0.2 and activity_significance[target] < 0.2:
                        continue

                    max_corr = 0
                    best_lag = 0
                    best_corr_sign = 0

                    # Try different lags
                    for lag in range(1, min(max_lag + 1, len(velocities) - 5)):
                        if lag >= len(velocities):
                            continue

                        source_vel = velocities[:-lag, source]
                        target_vel = velocities[lag:, target]

                        if len(source_vel) > 5:
                            try:
                                # Correlation between source velocity and target velocity
                                corr = np.corrcoef(source_vel, target_vel)[0, 1]
                                if not np.isnan(corr) and abs(corr) > abs(max_corr):
                                    max_corr = corr
                                    best_lag = lag
                                    best_corr_sign = np.sign(corr)
                            except:
                                pass

                    # Store influence with sign (positive or negative relationship)
                    influence_matrix[source, target] = max_corr

                    # Determine relationship type
                    if abs(max_corr) < 0.2:  # Ignore weak relationships
                        continue

                    if best_corr_sign > 0:
                        # Test if it's amplification
                        source_amp = np.mean(np.abs(velocities[:-best_lag, source]))
                        target_amp = np.mean(np.abs(velocities[best_lag:, target]))
                        amp_ratio = target_amp / (source_amp + 1e-10)

                        if amp_ratio > 1.5:
                            relationship_types[source, target] = 3  # amplifies_activity
                        else:
                            relationship_types[source, target] = 1  # drives_same_direction
                    else:
                        # Test if it's dampening
                        source_amp = np.mean(np.abs(velocities[:-best_lag, source]))
                        target_amp = np.mean(np.abs(velocities[best_lag:, target]))
                        amp_ratio = source_amp / (target_amp + 1e-10)

                        if amp_ratio > 1.5:
                            relationship_types[source, target] = 4  # dampens_activity
                        else:
                            relationship_types[source, target] = 2  # drives_opposite_direction

        # Recalculate contribution percentages per target electrode using absolute influence values
        contribution_percentages = np.zeros_like(influence_matrix)
        for target in range(num_electrodes):
            incoming = np.abs(influence_matrix[:, target])
            total_incoming = np.sum(incoming)
            if total_incoming > 0:
                contribution_percentages[:, target] = (incoming / total_incoming) * 100.0

        # Extract significant relationships
        threshold = 0.25  # Correlation threshold
        significance_threshold = 0.25  # Activity significance threshold

        significant_relationships = []
        for source in range(num_electrodes):
            if activity_significance[source] < significance_threshold:
                continue

            for target in range(num_electrodes):
                if source == target or activity_significance[target] < significance_threshold:
                    continue

                if abs(influence_matrix[source, target]) >= threshold:
                    rel_type = relationship_types[source, target]
                    if rel_type == 0:  # If not set, use default based on correlation sign
                        rel_type = 1 if influence_matrix[source, target] > 0 else 2

                    significant_relationships.append({
                        'source': source,
                        'target': target,
                        'influence': abs(influence_matrix[source, target]),
                        'contribution': contribution_percentages[source, target],
                        'rel_type': rel_type
                    })

        # Sort by influence strength
        significant_relationships = sorted(significant_relationships, key=lambda x: x['influence'], reverse=True)

        # Limit to top relationships to avoid clutter
        top_n = min(50, len(significant_relationships))
        significant_relationships = significant_relationships[:top_n]

        # Create 3D visualization
        fig = make_subplots(
            rows=2, cols=2,
            specs=[[{"type": "scatter3d", "colspan": 2}, None],
                   [{"type": "bar"}, {"type": "heatmap"}]],
            row_heights=[0.7, 0.3],
            column_widths=[0.7, 0.3],
            subplot_titles=("3D Electrode Interaction Network",
                          "Top Source Contributions",
                          "Relationship Distribution"),
            vertical_spacing=0.1,
            horizontal_spacing=0.05
        )

        # Create network visualization
        import networkx as nx
        G = nx.DiGraph()

        # Determine prominent electrodes (nodes) based on activity and relationships
        involved_electrodes = set()
        for rel in significant_relationships:
            involved_electrodes.add(rel['source'])
            involved_electrodes.add(rel['target'])

        # Calculate 3D node positions using spectral layout with z-dimension based on activity
        # First create a 2D layout
        edge_list = [(rel['source'], rel['target']) for rel in significant_relationships]
        G_temp = nx.DiGraph()
        G_temp.add_edges_from(edge_list)

        try:
            pos_2d = nx.kamada_kawai_layout(G_temp)
        except:
            try:
                pos_2d = nx.spring_layout(G_temp, k=0.5, iterations=100, seed=42)
            except:
                # Fallback for disconnected graph
                pos_2d = {e: (np.random.random()-0.5, np.random.random()-0.5) for e in involved_electrodes}

        # Add z coordinate based on activity significance
        pos_3d = {}
        for e in involved_electrodes:
            if e in pos_2d:
                pos_3d[e] = (pos_2d[e][0], pos_2d[e][1], activity_significance[e])
            else:
                pos_3d[e] = (np.random.random()-0.5, np.random.random()-0.5, activity_significance[e])

        # Calculate electrode roles (sender vs receiver)
        electrode_roles = {}
        for e in involved_electrodes:
            outgoing = sum(1 for rel in significant_relationships if rel['source'] == e)
            incoming = sum(1 for rel in significant_relationships if rel['target'] == e)

            if incoming == 0:
                ratio = float('inf')
            else:
                ratio = outgoing / incoming

            if ratio > 1.5:
                role = 'sender'
            elif ratio < 0.67:
                role = 'receiver'
            else:
                role = 'mixed'

            electrode_roles[e] = {
                'role': role,
                'outgoing': outgoing,
                'incoming': incoming,
                'total': outgoing + incoming
            }

        # Add nodes to graph visualization
        for e in involved_electrodes:
            G.add_node(e, pos=pos_3d[e], role=electrode_roles[e]['role'],
                       size=20 + 50 * electrode_roles[e]['total'] / max(1, max(r['total'] for r in electrode_roles.values())))

        # Add edges for significant relationships
        for rel in significant_relationships:
            G.add_edge(rel['source'], rel['target'],
                      influence=rel['influence'],
                      contribution=rel['contribution'],
                      rel_type=rel['rel_type'])

        # Generate node trace
        node_x = []
        node_y = []
        node_z = []
        node_size = []
        node_color = []
        node_text = []

        # Color scheme
        sender_color = 'rgba(214, 39, 40, 0.9)'  # Red
        receiver_color = 'rgba(31, 119, 180, 0.9)'  # Blue
        mixed_color = 'rgba(148, 103, 189, 0.9)'  # Purple

        for node in G.nodes():
            x, y, z = G.nodes[node]['pos']
            node_x.append(x)
            node_y.append(y)
            node_z.append(z)
            node_size.append(G.nodes[node]['size'])

            role = G.nodes[node]['role']
            if role == 'sender':
                node_color.append(sender_color)
            elif role == 'receiver':
                node_color.append(receiver_color)
            else:
                node_color.append(mixed_color)

            # Hover text
            node_text.append(f"Electrode {node}<br>" +
                           f"Outgoing connections: {electrode_roles[node]['outgoing']}<br>" +
                           f"Incoming connections: {electrode_roles[node]['incoming']}<br>" +
                           f"Activity significance: {activity_significance[node]:.2f}<br>" +
                           f"Role: {role.title()}")

        # Add node trace
        node_trace = go.Scatter3d(
            x=node_x,
            y=node_y,
            z=node_z,
            mode='markers+text',
            marker=dict(
                size=node_size,
                color=node_color,
                line=dict(width=1, color='rgb(40,40,40)')
            ),
            text=[f"E{n}" for n in G.nodes()],
            textposition="top center",
            textfont=dict(size=10, color="black"),
            hoverinfo="text",
            hovertext=node_text,
            name="Electrodes"
        )

        fig.add_trace(node_trace, row=1, col=1)

        # Group edges by relationship type
        rel_type_names = {
            1: 'Drives Same Direction',
            2: 'Drives Opposite Direction',
            3: 'Amplifies Activity',
            4: 'Dampens Activity',
            5: 'Complex Influence'
        }

        # Define colors for relationship types
        rel_type_colors = {
            1: 'rgba(44, 160, 44, 0.8)',  # Green for same direction
            2: 'rgba(214, 39, 40, 0.8)',  # Red for opposite direction
            3: 'rgba(31, 119, 180, 0.8)',  # Blue for amplification
            4: 'rgba(255, 127, 14, 0.8)',  # Orange for dampening
            5: 'rgba(188, 189, 34, 0.8)'   # Yellow for complex
        }

        # Group edges by relationship type for better legend
        for rel_type, rel_name in rel_type_names.items():
            # Get edges of this type
            type_edges = [(s, t) for s, t, d in G.edges(data=True) if d.get('rel_type', 0) == rel_type]

            if not type_edges:
                continue

            edge_x = []
            edge_y = []
            edge_z = []
            edge_width = []
            edge_text = []

            for source, target in type_edges:
                x0, y0, z0 = G.nodes[source]['pos']
                x1, y1, z1 = G.nodes[target]['pos']
                influence = G.edges[source, target]['influence']
                contribution = G.edges[source, target]['contribution']

                # Create curved line
                points = 10
                for i in range(points):
                    t = i / (points - 1)
                    # Add curve using cubic Bezier
                    # Control points for arc effect
                    xc = (x0 + x1) / 2
                    yc = (y0 + y1) / 2
                    zc = max(z0, z1) + 0.2  # Arc upward

                    # Bezier interpolation formula
                    xi = (1-t)**2 * x0 + 2*(1-t)*t * xc + t**2 * x1
                    yi = (1-t)**2 * y0 + 2*(1-t)*t * yc + t**2 * y1
                    zi = (1-t)**2 * z0 + 2*(1-t)*t * zc + t**2 * z1

                    edge_x.append(xi)
                    edge_y.append(yi)
                    edge_z.append(zi)

                    if i == points // 2:  # Middle point for hover text
                        edge_text.append(f"E{source} → E{target}<br>" +
                                       f"Relationship: {rel_name}<br>" +
                                       f"Influence strength: {influence:.2f}<br>" +
                                       f"Contribution: {contribution:.1f}%")
                    else:
                        edge_text.append("")

                edge_width.extend([3 + 7 * influence] * points)

                # Add None to break the line
                edge_x.append(None)
                edge_y.append(None)
                edge_z.append(None)
                edge_text.append(None)
                edge_width.append(None)

            # Create edge trace
            edge_trace = go.Scatter3d(
                x=edge_x,
                y=edge_y,
                z=edge_z,
                mode='lines',
                line=dict(
                    width=5,  # Base width, will be scaled
                    color=rel_type_colors.get(rel_type, 'gray')
                ),
                hoverinfo="text",
                hovertext=edge_text,
                name=rel_name
            )

            fig.add_trace(edge_trace, row=1, col=1)

        # Create bar chart showing top source contributions by target
        # For each target, calculate total incoming percentage by relationship type
        target_contributions = {}
        for rel in significant_relationships:
            target = rel['target']
            source = rel['source']
            rel_type = rel['rel_type']
            contribution = rel['contribution']

            if target not in target_contributions:
                target_contributions[target] = {'total': 0, 'sources': {}}

            target_contributions[target]['total'] += contribution

            source_key = f"E{source}"
            if source_key not in target_contributions[target]['sources']:
                target_contributions[target]['sources'][source_key] = {
                    'contribution': 0,
                    'rel_type': rel_type
                }

            target_contributions[target]['sources'][source_key]['contribution'] += contribution

        # Sort targets by total contribution
        sorted_targets = sorted(target_contributions.items(),
                               key=lambda x: x[1]['total'],
                               reverse=True)

        # Take top 3 targets with highest total incoming contribution
        top_targets = sorted_targets[:min(3, len(sorted_targets))]

        # Create stacked bar chart
        for target_idx, (target, target_data) in enumerate(top_targets):
            # Sort sources by contribution
            sources = sorted(target_data['sources'].items(),
                            key=lambda x: x[1]['contribution'],
                            reverse=True)

            # Take top 5 sources for this target
            top_sources = sources[:min(5, len(sources))]

            for source_idx, (source_key, source_data) in enumerate(top_sources):
                rel_type = source_data['rel_type']
                contribution = source_data['contribution']

                # Skip very small contributions
                if contribution < 5:
                    continue

                fig.add_trace(
                    go.Bar(
                        y=[f"E{target}"],
                        x=[contribution],
                        name=f"{source_key} → E{target}" if source_idx == 0 else None,
                        showlegend=source_idx == 0,
                        legendgroup=f"E{target}",
                        marker_color=rel_type_colors.get(rel_type, 'gray'),
                        orientation='h',
                        text=f"{contribution:.1f}%",
                        textposition="inside",
                        hoverinfo="text",
                        hovertext=f"{source_key} contributes {contribution:.1f}% to E{target}<br>Relationship: {rel_type_names.get(rel_type, 'Unknown')}"
                    ),
                    row=2, col=1
                )

        # Create heatmap of relationship types - calculate percentage distribution
        rel_type_counts = {rt: 0 for rt in rel_type_names.keys()}
        total_relationships = len(significant_relationships)

        if total_relationships > 0:
            for rel in significant_relationships:
                rel_type_counts[rel['rel_type']] = rel_type_counts.get(rel['rel_type'], 0) + 1

            rel_type_percentages = {rt: (count / total_relationships) * 100
                                  for rt, count in rel_type_counts.items()}
        else:
            rel_type_percentages = {rt: 0 for rt in rel_type_names.keys()}

        # Sort by percentage
        sorted_rel_types = sorted([(rt, rel_type_names[rt], pct)
                                 for rt, pct in rel_type_percentages.items() if pct > 0],
                                key=lambda x: x[2], reverse=True)

        if sorted_rel_types:  # Only create heatmap if we have relationship data
            heatmap_z = [[pct] for _, _, pct in sorted_rel_types]
            heatmap_y = [name for _, name, _ in sorted_rel_types]
            heatmap_colors = [rel_type_colors.get(rt, 'gray') for rt, _, _ in sorted_rel_types]

            fig.add_trace(
                go.Heatmap(
                    z=heatmap_z,
                    y=heatmap_y,
                    colorscale=[[0, c] for i, c in enumerate(heatmap_colors)],
                    showscale=False,
                    text=[[f"{pct:.1f}%"] for _, _, pct in sorted_rel_types],
                    texttemplate="%{text}",
                    hoverinfo="text",
                    hovertext=[f"{name}: {pct:.1f}% of connections"
                              for _, name, pct in sorted_rel_types]
                ),
                row=2, col=2
            )

        # Update layout
        fig.update_layout(
            title=dict(
                text=title,
                y=0.98,
                x=0.5,
                xanchor="center",
                yanchor="top",
                font=dict(family="Arial", size=24, color="#333333")
            ),
            scene=dict(
                xaxis=dict(title="", showticklabels=False, range=[-1, 1]),
                yaxis=dict(title="", showticklabels=False, range=[-1, 1]),
                zaxis=dict(title="Influence Strength", showticklabels=False, range=[0, 1.2])
            ),
            xaxis2=dict(
                title="Contribution Percentage (%)",
                range=[0, 100]  # Force x-axis to 0-100%
            ),
            yaxis2=dict(
                title="Target Electrodes",
                autorange="reversed"  # Reverse y-axis for better readability
            ),
            xaxis3=dict(
                showticklabels=False,
                range=[0, 100]
            ),
            yaxis3=dict(
                title="Relationship Types",
                autorange="reversed"
            ),
            barmode='stack',
            width=1200,
            height=900,
            paper_bgcolor="rgb(248, 249, 250)",
            plot_bgcolor="rgb(248, 249, 250)",
            legend=dict(
                orientation="h",
                yanchor="bottom",
                y=1.02,
                xanchor="right",
                x=1
            )
        )

        # Add explanatory annotations
        fig.add_annotation(
            x=0.01,
            y=0.01,
            xref="paper",
            yref="paper",
            text=(
                "This visualization shows electrode interactions discovered through neural genetic programming.<br>" +
                "• Node size indicates electrode influence<br>" +
                "• Node color shows role (Red=Sender, Blue=Receiver, Purple=Mixed)<br>" +
                "• Edge color indicates relationship type<br>" +
                "• Bar chart shows percentage contribution of source electrodes<br>" +
                "• Heatmap shows distribution of relationship types"
            ),
            showarrow=False,
            align="left",
            bgcolor="rgba(255, 255, 255, 0.8)",
            bordercolor="#e0e0e0",
            borderwidth=1,
            borderpad=4,
            font=dict(size=12)
        )

        # Save figure if path provided
        if save_path:
            import plotly.io as pio
            pio.write_html(fig, save_path)
            print(f"Enhanced 3D electrode rule tree saved to {save_path}")

        return fig


def format_relationship(rel_type):
    """Format relationship type for display."""
    if rel_type == "drives_same_direction":
        return "Drives Same Direction"
    elif rel_type == "drives_opposite_direction":
        return "Drives Opposite Direction"
    elif rel_type == "amplifies_activity":
        return "Amplifies Activity"
    elif rel_type == "dampens_activity":
        return "Dampens Activity"
    elif rel_type == "phase_synchronization":
        return "Phase Synchronization"
    elif rel_type == "phase_desynchronization":
        return "Phase Desynchronization"
    elif rel_type == "complex_influence":
        return "Complex Influence"
    else:
        return rel_type.replace("_", " ").title()

class NeuralLogicGateKnowledgeGraph:
    """
    Creates a knowledge graph visualization showing logical gate relationships
    between electrode hidden states, discovered through genetic programming.
    """
    def __init__(self, model, config):
        self.model = model
        self.config = config
        self.device = next(model.parameters()).device
        self.rule_cache = {}  # Cache discovered rules
        self.electrode_roles = {}  # Store electrode roles

    def extract_hidden_states(self, inputs_dict, sample_idx=0):
        """Extract hidden states from the model for rule discovery."""
        with torch.no_grad():
            # Get raw inputs
            raw_inputs = inputs_dict['raw'].to(self.device)
            batch_size, context_size, num_electrodes = raw_inputs.shape

            # Run model to get state history
            outputs, state_history = self.model(inputs_dict, return_states=True)

            if not state_history or len(state_history) == 0:
                print("No state history returned by model. Using electrode signals directly.")
                # Fallback to using raw signals
                electrode_states = []
                for e_idx in range(num_electrodes):
                    signal = raw_inputs[sample_idx, :, e_idx].cpu().numpy()
                    electrode_states.append(signal)
                return electrode_states, None, raw_inputs.cpu().numpy()[sample_idx]

            # Extract per-electrode hidden state features
            electrode_hidden_states = []
            system_states = []

            for t, population_states in enumerate(state_history):
                # Combine population states for system state
                if population_states:
                    combined_state = torch.cat(population_states, dim=1)[sample_idx].cpu().numpy()
                    system_states.append(combined_state)

            # For each electrode, extract relevant state components
            for e_idx in range(num_electrodes):
                e_states = []

                for t in range(min(len(state_history), context_size)):
                    if t >= len(state_history):
                        break

                    # Weight the state by electrode activity
                    electrode_value = raw_inputs[sample_idx, min(t, context_size-1), e_idx].cpu().numpy()

                    # Create a compact representation for this electrode's state
                    # Use statistical features of the state
                    state_mean = np.mean(system_states[t])
                    state_var = np.var(system_states[t])
                    state_max = np.max(system_states[t])
                    state_min = np.min(system_states[t])

                    # Extract the main directions of variation
                    pca_state = np.zeros(4)
                    if len(system_states[t]) > 4:
                        from sklearn.decomposition import PCA
                        pca = PCA(n_components=4)
                        try:
                            pca_state = pca.fit_transform(system_states[t].reshape(-1, 1)).flatten()[:4]
                        except:
                            pass

                    # Combine these into a feature vector
                    feature_vector = np.array([state_mean, state_var, state_max, state_min,
                                              electrode_value, electrode_value**2,
                                              *pca_state])

                    e_states.append(feature_vector)

                if e_states:
                    electrode_hidden_states.append(np.array(e_states))
                else:
                    # Create dummy states if extraction failed
                    dummy_state = np.zeros((min(len(state_history), context_size), 10))
                    electrode_hidden_states.append(dummy_state)

            return electrode_hidden_states, system_states, raw_inputs.cpu().numpy()[sample_idx]

    def discover_logic_gate_rules(self, electrode_states, electrode_signals):
        """
        Discover logical gate rules between electrodes using genetic programming techniques.
        """
        num_electrodes = len(electrode_states)
        rules = []

        # Calculate state changes (velocity in state space)
        velocities = []
        for e_idx, states in enumerate(electrode_states):
            if states.shape[0] < 2:
                velocities.append(np.zeros((0, states.shape[1])))
                continue

            v = np.diff(states, axis=0)
            velocities.append(v)

        # Find significant state changes
        significant_changes = {}
        for e_idx, velocity in enumerate(velocities):
            if len(velocity) == 0:
                continue

            # Measure magnitude of velocity
            magnitudes = np.sqrt(np.sum(velocity**2, axis=1))

            # Find significant changes (above 75th percentile)
            if len(magnitudes) > 0:
                threshold = np.percentile(magnitudes, 75)
                changes = np.where(magnitudes > threshold)[0]
                if len(changes) > 0:
                    significant_changes[e_idx] = changes

        # Discover logical rules
        rule_types = ['AND', 'OR', 'XOR', 'NOT', 'IMPLIES']

        # First look for direct source->target rules
        for source, source_changes in significant_changes.items():
            for target, target_changes in significant_changes.items():
                if source == target:
                    continue

                # Try different lags
                max_lag = min(5, len(velocities[source]) // 4)

                for lag in range(1, max_lag + 1):
                    # Check if source changes predict target changes
                    matches = 0
                    total = 0

                    for s_change in source_changes:
                        # Skip if too close to the end
                        if s_change + lag >= len(velocities[source]):
                            continue

                        # Expected target change time
                        t_expected = s_change + lag

                        # See if target actually changes within a small window
                        window = 2  # Look 2 steps before and after
                        window_start = max(0, t_expected - window)
                        window_end = min(len(velocities[target]) if len(velocities[target]) > 0 else 0,
                                         t_expected + window + 1)

                        # Check if any target change in window
                        target_changes_in_window = [t for t in target_changes
                                                   if t >= window_start and t < window_end]

                        total += 1
                        if target_changes_in_window:
                            matches += 1

                    # Calculate probability
                    probability = matches / total if total > 0 else 0

                    # If strong relationship, create a DIRECT rule
                    if probability >= 0.5 and matches >= 3:
                        rules.append({
                            'type': 'DIRECT',
                            'sources': [source],
                            'target': target,
                            'lag': lag,
                            'probability': probability,
                            'matches': matches,
                            'total': total
                        })

        # Now look for logical gate rules (AND, OR, XOR)
        # We'll focus on combining pairs of sources using logical operations
        pairs_tried = set()

        for rule_type in ['AND', 'OR', 'XOR']:
            # For each potential target
            for target, target_changes in significant_changes.items():
                # Find potential source pairs
                source_candidates = [s for s in significant_changes.keys() if s != target]

                if len(source_candidates) < 2:
                    continue

                # Try different source combinations
                from itertools import combinations
                for source1, source2 in combinations(source_candidates, 2):
                    # Skip if we've already tried this triplet
                    triplet = tuple(sorted([source1, source2, target]))
                    if triplet in pairs_tried:
                        continue

                    pairs_tried.add(triplet)

                    # Find times when both source1 and source2 change in close proximity
                    source1_changes = significant_changes[source1]
                    source2_changes = significant_changes[source2]

                    window = 2  # Consider changes within 2 time steps
                    joint_changes = []

                    for s1_change in source1_changes:
                        for s2_change in source2_changes:
                            if abs(s1_change - s2_change) <= window:
                                joint_t = min(s1_change, s2_change)
                                joint_changes.append(joint_t)
                                break

                    if not joint_changes:
                        continue

                    # Now see if these joint changes predict target changes
                    max_lag = min(5, len(velocities[target]) // 4)

                    for lag in range(1, max_lag + 1):
                        matches = 0
                        total = 0

                        for joint_t in joint_changes:
                            # Skip if too close to the end
                            if joint_t + lag >= len(velocities[target]):
                                continue

                            # Expected target change time
                            t_expected = joint_t + lag

                            # See if target actually changes
                            window = 2
                            window_start = max(0, t_expected - window)
                            window_end = min(len(velocities[target]), t_expected + window + 1)

                            # Check if any target change in window
                            target_changes_in_window = [t for t in target_changes
                                                       if t >= window_start and t < window_end]

                            total += 1
                            if target_changes_in_window:
                                matches += 1

                        # Calculate probability
                        probability = matches / total if total > 0 else 0

                        # If strong relationship, create a logical gate rule
                        if probability >= 0.5 and matches >= 2:
                            rules.append({
                                'type': rule_type,
                                'sources': [source1, source2],
                                'target': target,
                                'lag': lag,
                                'probability': probability,
                                'matches': matches,
                                'total': total
                            })

        # Find NOT gates
        for source, source_changes in significant_changes.items():
            for target, target_changes in significant_changes.items():
                if source == target:
                    continue

                # Look for inverse relationships - when source changes, target doesn't
                max_lag = min(5, len(velocities[source]) // 4)

                for lag in range(1, max_lag + 1):
                    opposite_matches = 0
                    total = 0

                    for s_change in source_changes:
                        # Skip if too close to the end
                        if s_change + lag >= len(velocities[source]):
                            continue

                        # Expected target change time
                        t_expected = s_change + lag

                        # See if target DOESN'T change within window
                        window = 2
                        window_start = max(0, t_expected - window)
                        window_end = min(len(velocities[target]), t_expected + window + 1)

                        # Check if no target change in window
                        target_changes_in_window = [t for t in target_changes
                                                   if t >= window_start and t < window_end]

                        total += 1
                        if not target_changes_in_window:  # Target stayed unchanged
                            opposite_matches += 1

                    # Calculate probability of NOT relationship
                    probability = opposite_matches / total if total > 0 else 0

                    # If strong inverse relationship, create a NOT rule
                    if probability >= 0.65 and opposite_matches >= 3:
                        rules.append({
                            'type': 'NOT',
                            'sources': [source],
                            'target': target,
                            'lag': lag,
                            'probability': probability,
                            'matches': opposite_matches,
                            'total': total
                        })

        # Find implication rules (IF source1 THEN source2)
        for source1, source1_changes in significant_changes.items():
            for source2, source2_changes in significant_changes.items():
                if source1 == source2:
                    continue

                # Check if source1 implies source2
                max_lag = min(5, len(velocities[source1]) // 4)

                for lag in range(1, max_lag + 1):
                    implication_matches = 0
                    total = 0

                    for s1_change in source1_changes:
                        # Skip if too close to the end
                        if s1_change + lag >= len(velocities[source1]):
                            continue

                        # Expected source2 change time
                        t_expected = s1_change + lag

                        # Check if source2 changes after source1
                        window = 2
                        window_start = max(0, t_expected - window)
                        window_end = min(len(velocities[source2]), t_expected + window + 1)

                        # Check if any source2 change in window
                        changes_in_window = [t for t in source2_changes
                                            if t >= window_start and t < window_end]

                        total += 1
                        if changes_in_window:
                            implication_matches += 1

                    # Calculate probability of implication
                    probability = implication_matches / total if total > 0 else 0

                    # If strong implication, create IMPLIES rule
                    if probability >= 0.6 and implication_matches >= 3:
                        rules.append({
                            'type': 'IMPLIES',
                            'sources': [source1],
                            'target': source2,
                            'lag': lag,
                            'probability': probability,
                            'matches': implication_matches,
                            'total': total
                        })

        # Determine electrode roles based on discovered rules
        self.determine_electrode_roles(rules, num_electrodes)

        return rules

    def determine_electrode_roles(self, rules, num_electrodes):
        """Determine electrode roles based on discovered rules."""
        # Count incoming and outgoing connections for each electrode
        incoming = defaultdict(int)
        outgoing = defaultdict(int)

        for rule in rules:
            for source in rule['sources']:
                outgoing[source] += 1

            target = rule['target']
            incoming[target] += 1

        # Initialize electrode roles
        self.electrode_roles = {}

        # Determine role based on connectivity pattern
        for e_idx in range(num_electrodes):
            in_count = incoming.get(e_idx, 0)
            out_count = outgoing.get(e_idx, 0)

            if out_count > 2*in_count and out_count > 1:
                role = 'initiator'  # Primarily influences others
            elif in_count > 2*out_count and in_count > 1:
                role = 'receiver'   # Primarily influenced by others
            elif in_count >= 3 and out_count >= 3:
                role = 'hub'        # Both influences and is influenced
            else:
                role = 'relay'      # Balanced or minimal connections

            self.electrode_roles[e_idx] = {
                'role': role,
                'incoming': in_count,
                'outgoing': out_count
            }

    def create_visualization(self, inputs_dict, sample_idx=0, save_path=None,
                             title="Enhanced Neural State Knowledge Graph with Logic Gate Rules"):
        """
        Create an interactive 3D visualization of the logic gate knowledge graph.
        """
        import plotly.graph_objects as go

        # Extract hidden states
        electrode_states, system_states, electrode_signals = self.extract_hidden_states(inputs_dict, sample_idx)

        # Discover logic gate rules
        rules = self.discover_logic_gate_rules(electrode_states, electrode_signals)

        # Sort rules by probability
        rules = sorted(rules, key=lambda x: x['probability'], reverse=True)

        # Take top rules for visualization clarity
        top_rules = rules[:min(75, len(rules))]

        # Create a graph representation
        import networkx as nx
        G = nx.DiGraph()

        # Add the system node
        G.add_node('system', type='system', importance=1.0)

        # Identify all electrodes involved in rules
        involved_electrodes = set()
        for rule in top_rules:
            for source in rule['sources']:
                involved_electrodes.add(source)
            involved_electrodes.add(rule['target'])

        # IMPORTANT FIX: If too few electrodes are involved, add more based on significance
        if len(involved_electrodes) < 10:
            num_electrodes = len(electrode_states)
            # Add most active electrodes based on signal variance
            activity_levels = []
            for e_idx in range(num_electrodes):
                if e_idx not in involved_electrodes:
                    signal = electrode_signals[:, e_idx]
                    variance = np.var(signal)
                    activity_levels.append((e_idx, variance))

            # Sort by activity and add top ones
            activity_levels.sort(key=lambda x: x[1], reverse=True)
            additional_electrodes = [e for e, _ in activity_levels[:15]]
            involved_electrodes.update(additional_electrodes)

        # Create better 3D layout using advanced algorithm
        # First, create a more connected graph for better layout
        layout_G = nx.DiGraph()

        # Add more connections for layout purposes
        for e1 in involved_electrodes:
            for e2 in involved_electrodes:
                if e1 != e2:
                    layout_G.add_edge(e1, e2)

        # Try layout algorithms with error handling
        pos_3d = {}
        try:
            pos_2d = nx.kamada_kawai_layout(layout_G)
        except:
            try:
                pos_2d = nx.spring_layout(layout_G, k=0.3, iterations=100)
            except:
                # Create a circular layout as fallback
                angles = np.linspace(0, 2*np.pi, len(involved_electrodes), endpoint=False)
                radius = 1.0
                pos_2d = {e: (radius * np.cos(angle), radius * np.sin(angle))
                          for e, angle in zip(involved_electrodes, angles)}

        # NEW: Organize in a multi-level hierarchical structure
        # Use 3 levels based on roles: initiator->hub->receiver
        for e in involved_electrodes:
            role = self.electrode_roles.get(e, {}).get('role', 'relay')

            # More structured z-position based on role
            if role == 'initiator':
                z = 0.2  # Bottom level
            elif role == 'hub':
                z = 0.5  # Middle level
            elif role == 'receiver':
                z = 0.8  # Upper level
            else:  # relay
                z = 0.5  # Middle level

            # Get 2D position or generate random
            if e in pos_2d:
                # Scale positions to avoid overcrowding
                x, y = pos_2d[e]
                # Scale to fit a nice circle
                dist = np.sqrt(x**2 + y**2)
                if dist > 0:
                    scale_factor = 0.8 / dist
                    x *= scale_factor
                    y *= scale_factor
                pos_3d[e] = (x, y, z)
            else:
                angle = np.random.random() * 2 * np.pi
                radius = 0.7 + np.random.random() * 0.2
                pos_3d[e] = (radius * np.cos(angle), radius * np.sin(angle), z)

        # Position for system node at the top
        pos_3d['system'] = (0, 0, 1.0)

        # NEW: Add more structure to electrode positions - form a hemisphere
        for e in involved_electrodes:
            x, y, z = pos_3d[e]
            # Adjust to form a hemispheric shell
            dist = np.sqrt(x**2 + y**2)
            if dist > 0:
                # Calculate theta (azimuthal angle) and phi (polar angle)
                theta = np.arctan2(y, x)
                phi = np.pi/2 - np.arctan(z / dist)  # Convert from z to polar angle

                # Adjust radius to form a hemisphere
                radius = 0.8

                # Convert back to cartesian
                x = radius * np.sin(phi) * np.cos(theta)
                y = radius * np.sin(phi) * np.sin(theta)
                z = radius * np.cos(phi)

                # Store new position
                pos_3d[e] = (x, y, z)

        # Add electrode nodes to graph with improved sizing and coloring
        for e_idx in involved_electrodes:
            role = self.electrode_roles.get(e_idx, {}).get('role', 'relay')

            # Enhanced node sizing based on importance
            incoming = self.electrode_roles.get(e_idx, {}).get('incoming', 0)
            outgoing = self.electrode_roles.get(e_idx, {}).get('outgoing', 0)

            # Set node color based on role
            if role == 'initiator':
                color = 'rgba(214, 39, 40, 0.9)'  # Red
                size = 20 + 10 * outgoing
            elif role == 'receiver':
                color = 'rgba(31, 119, 180, 0.9)'  # Blue
                size = 20 + 10 * incoming
            elif role == 'hub':
                color = 'rgba(148, 103, 189, 0.9)'  # Purple
                size = 25 + 5 * (incoming + outgoing)
            else:  # relay
                color = 'rgba(44, 160, 44, 0.9)'  # Green
                size = 20

            G.add_node(e_idx, pos=pos_3d[e_idx], role=role, color=color, size=size)

        # Add system node
        G.add_node('system', pos=pos_3d['system'], role='system',
                  color='rgba(255, 215, 0, 0.9)', size=30)

        # Create edge mapping for visualization
        edge_info = {}

        # Colors for different rule types
        rule_colors = {
            'DIRECT': 'rgba(31, 119, 180, 0.8)',       # Blue
            'AND': 'rgba(44, 160, 44, 0.8)',           # Green
            'OR': 'rgba(214, 39, 40, 0.8)',            # Red
            'XOR': 'rgba(148, 103, 189, 0.8)',         # Purple
            'NOT': 'rgba(140, 86, 75, 0.8)',           # Brown
            'IMPLIES': 'rgba(247, 129, 191, 0.8)'      # Pink
        }

        # Secondary colors (lighter versions)
        secondary_colors = {
            'AND': 'rgba(44, 160, 44, 0.5)',           # Light Green
            'OR': 'rgba(214, 39, 40, 0.5)',            # Light Red
            'XOR': 'rgba(148, 103, 189, 0.5)',         # Light Purple
            'IMPLIES': 'rgba(247, 129, 191, 0.5)',     # Light Pink
            'MULTI_SOURCE': 'rgba(255, 187, 120, 0.5)'  # Light Orange
        }

        # Add edges for each rule
        for rule in top_rules:
            rule_type = rule['type']
            sources = rule['sources']
            target = rule['target']
            probability = rule['probability']

            # Edge color based on rule type
            color = rule_colors.get(rule_type, 'rgba(100, 100, 100, 0.6)')

            # Edge width based on rule probability
            width = 1 + 9 * probability  # Scale 1-10

            # For multi-source rules (AND, OR, XOR)
            if len(sources) > 1:
                # First add edges from sources to an invisible "gate" node
                gate_id = f"{rule_type}_{','.join(map(str, sources))}_{target}"

                # Create position for the gate node
                gate_pos = np.mean([pos_3d[s] for s in sources], axis=0)
                gate_pos = (gate_pos[0], gate_pos[1], gate_pos[2] + 0.1)  # Slightly higher

                # Add invisible gate node
                G.add_node(gate_id, pos=gate_pos, role='gate',
                          color='rgba(0, 0, 0, 0)', size=0)

                # Add source->gate edges
                for source in sources:
                    edge_key = (source, gate_id)
                    # Use secondary color for source->gate edges
                    edge_info[edge_key] = {
                        'color': secondary_colors.get(rule_type, secondary_colors['MULTI_SOURCE']),
                        'width': width * 0.7,  # Thinner
                        'label': '',
                        'dash': 'dash',
                        'rule_type': f"{rule_type} Secondary"
                    }

                # Add gate->target edge
                edge_key = (gate_id, target)
                edge_info[edge_key] = {
                    'color': color,
                    'width': width,
                    'label': f"{rule_type} Gate",
                    'dash': None,
                    'rule_type': rule_type
                }
            else:
                # Single source rules (DIRECT, NOT, IMPLIES)
                source = sources[0]
                edge_key = (source, target)

                # Check if we already have this edge
                if edge_key in edge_info:
                    # Keep the stronger/clearer rule
                    if probability > edge_info[edge_key].get('probability', 0):
                        edge_info[edge_key] = {
                            'color': color,
                            'width': width,
                            'label': rule_type,
                            'dash': None,
                            'rule_type': rule_type,
                            'probability': probability
                        }
                else:
                    edge_info[edge_key] = {
                        'color': color,
                        'width': width,
                        'label': rule_type,
                        'dash': None,
                        'rule_type': rule_type,
                        'probability': probability
                    }

        # NEW: Create additional connections for any disconnected nodes
        connected_nodes = set()
        for source, target in edge_info.keys():
            connected_nodes.add(source)
            connected_nodes.add(target)

        disconnected_nodes = set(G.nodes()) - connected_nodes
        for node in disconnected_nodes:
            if node == 'system' or isinstance(node, str):
                continue

            # Find nearest connected node (excluding system)
            nearest_dist = float('inf')
            nearest_node = None

            for connected in connected_nodes:
                if connected == 'system' or isinstance(connected, str):
                    continue

                # Calculate distance
                if node in pos_3d and connected in pos_3d:
                    dist = np.sqrt(sum((np.array(pos_3d[node]) - np.array(pos_3d[connected]))**2))
                    if dist < nearest_dist:
                        nearest_dist = dist
                        nearest_node = connected

            # Add connection to nearest
            if nearest_node is not None:
                edge_key = (node, nearest_node)
                edge_info[edge_key] = {
                    'color': 'rgba(100, 100, 100, 0.3)',  # Light gray
                    'width': 1,
                    'label': 'proximity',
                    'dash': 'dot',
                    'rule_type': 'proximity'
                }

            # Also connect to system for hierarchical structure
            edge_key = ('system', node)
            edge_info[edge_key] = {
                'color': 'rgba(255, 215, 0, 0.2)',  # Light gold
                'width': 1,
                'label': 'hierarchy',
                'dash': 'dot',
                'rule_type': 'hierarchy'
            }

        # Add edges to the graph
        for (source, target), info in edge_info.items():
            G.add_edge(source, target, **info)

        # Create figure
        fig = go.Figure()

        # Add electrode nodes
        node_trace_data = {}

        # Get all unique roles
        all_roles = set([G.nodes[n].get('role', 'relay') for n in G.nodes()])

        # Create a separate trace for each role
        for role in all_roles:
            # Get nodes with this role
            role_nodes = [n for n in G.nodes() if G.nodes[n].get('role', 'relay') == role]

            if not role_nodes:
                continue

            x = []
            y = []
            z = []
            node_size = []
            node_text = []
            node_color = []
            node_hover = []

            for node in role_nodes:
                pos = G.nodes[node]['pos']
                x.append(pos[0])
                y.append(pos[1])
                z.append(pos[2])

                size = G.nodes[node].get('size', 20)
                node_size.append(size)

                color = G.nodes[node].get('color', 'rgba(100, 100, 100, 0.9)')
                node_color.append(color)

                # Node text and hover info
                if node == 'system':
                    node_text.append('System')
                    node_hover.append('Unified System State<br>Hub of network processing')
                elif isinstance(node, int):
                    node_text.append(f'E{node}')

                    # Enhanced hover with connection info
                    incoming = self.electrode_roles.get(node, {}).get('incoming', 0)
                    outgoing = self.electrode_roles.get(node, {}).get('outgoing', 0)
                    node_hover.append(f"Electrode {node}<br>Role: {role.title()}<br>" +
                                    f"Incoming: {incoming}<br>Outgoing: {outgoing}")
                else:
                    # For gate nodes
                    node_text.append('')
                    gate_type = node.split('_')[0] if '_' in node else ''
                    node_hover.append(f"{gate_type} Logic Gate")

            # Create node trace (only show in legend if significant number of nodes)
            node_trace = go.Scatter3d(
                x=x,
                y=y,
                z=z,
                mode='markers+text',
                marker=dict(
                    size=node_size,
                    color=node_color,
                    line=dict(width=1, color='rgba(40,40,40,0.5)')
                ),
                text=node_text,
                textposition="top center",
                textfont=dict(size=10, color='black'),
                hoverinfo='text',
                hovertext=node_hover,
                name=role.title() if role != 'gate' else '',
                showlegend=role != 'gate' and len(role_nodes) >= 2
            )

            fig.add_trace(node_trace)

        # NEW: Group edges by type for better legend organization
        edge_by_type = defaultdict(list)

        for source, target, data in G.edges(data=True):
            rule_type = data.get('rule_type', 'Direct')
            edge_by_type[rule_type].append((source, target, data))

        # Add edges
        for rule_type, edges in edge_by_type.items():
            # Skip empty edge types or proximity/hierarchy edges for legend clarity
            if not edges or rule_type in ['proximity', 'hierarchy']:
                continue

            edge_x = []
            edge_y = []
            edge_z = []
            edge_text = []
            line_width = []

            for source, target, data in edges:
                # Get positions
                x0, y0, z0 = G.nodes[source]['pos']
                x1, y1, z1 = G.nodes[target]['pos']

                # Create curved edge
                pts = 20  # Number of points

                # Calculate midpoint (raised for curve)
                mid_x = (x0 + x1) / 2
                mid_y = (y0 + y1) / 2
                mid_z = max(z0, z1) + 0.1  # Raised midpoint

                # Create Bezier curve
                for t in np.linspace(0, 1, pts):
                    # Quadratic Bezier formula
                    px = (1-t)**2 * x0 + 2*(1-t)*t * mid_x + t**2 * x1
                    py = (1-t)**2 * y0 + 2*(1-t)*t * mid_y + t**2 * y1
                    pz = (1-t)**2 * z0 + 2*(1-t)*t * mid_z + t**2 * z1

                    edge_x.append(px)
                    edge_y.append(py)
                    edge_z.append(pz)

                    # Edge text and width
                    width = data.get('width', 3)
                    line_width.append(width)

                    # Edge hover text
                    if abs(t - 0.5) < 0.05:  # Near midpoint
                        if source == 'system' or target == 'system':
                            edge_text.append(f"System Connection")
                        elif 'Gate' in rule_type:
                            edge_text.append(f"{rule_type}")
                        else:
                            prob = data.get('probability', 0)
                            edge_text.append(f"{rule_type} Rule<br>Probability: {prob:.2f}")
                    else:
                        edge_text.append("")

                # Add None to break line
                edge_x.append(None)
                edge_y.append(None)
                edge_z.append(None)
                edge_text.append(None)
                line_width.append(None)

            # Create edge trace
            line_style = 'solid'
            if 'Secondary' in rule_type:
                line_style = 'dash'

            # Get color from first edge data
            example_edge = edges[0][2]
            edge_color = example_edge.get('color', 'rgba(100, 100, 100, 0.6)')

            edge_trace = go.Scatter3d(
                x=edge_x,
                y=edge_y,
                z=edge_z,
                mode='lines',
                line=dict(
                    color=edge_color,
                    width=3,  # Fixed width for visual consistency
                    dash=line_style
                ),
                hoverinfo='text',
                hovertext=edge_text,
                name=rule_type.replace('_', ' ').title()
            )

            fig.add_trace(edge_trace)

        # Update layout
        fig.update_layout(
            title=dict(
                text=title,
                y=0.95,
                x=0.5,
                xanchor='center',
                yanchor='top',
                font=dict(family='Arial', size=24, color='#333333')
            ),
            scene=dict(
                xaxis=dict(showticklabels=False, title='', showspikes=False, range=[-1.2, 1.2]),
                yaxis=dict(showticklabels=False, title='', showspikes=False, range=[-1.2, 1.2]),
                zaxis=dict(showticklabels=False, title='', showspikes=False, range=[-0.2, 1.2]),
                camera=dict(
                    eye=dict(x=1.5, y=1.5, z=1.2),
                    up=dict(x=0, y=0, z=1)
                ),
                aspectmode='manual',
                aspectratio=dict(x=1, y=1, z=0.8)
            ),
            width=1200,
            height=800,
            paper_bgcolor='rgb(248, 249, 250)',
            plot_bgcolor='rgb(248, 249, 250)',
            legend=dict(
                x=0.99,
                y=0.99,
                xanchor='right',
                yanchor='top',
                bgcolor='rgba(255, 255, 255, 0.8)',
                bordercolor='#e0e0e0',
                borderwidth=1
            ),
            margin=dict(l=0, r=0, b=0, t=50)
        )

        # Add explanation
        fig.add_annotation(
            x=0.01,
            y=0.01,
            xref="paper",
            yref="paper",
            text=(
                "This knowledge graph shows higher-order rule relationships between electrode hidden states.<br>" +
                "• <b>Gold Node:</b> Unified system state<br>" +
                "• <b>Red Nodes:</b> Initiator electrodes that often start influence chains<br>" +
                "• <b>Purple Nodes:</b> Hub electrodes that integrate multiple influences<br>" +
                "• <b>Blue Nodes:</b> Receiver electrodes that primarily receive influence<br>" +
                "• <b>Green Nodes:</b> Relay electrodes that pass influences<br>" +
                "• <b>Edge Colors:</b> Different rule types (logic gates, temporal patterns, etc.)<br>" +
                "• <b>Edge thickness:</b> Rule accuracy<br>" +
                "Rules were discovered using genetic programming on hidden state dynamics."
            ),
            showarrow=False,
            align="left",
            bgcolor="rgba(255, 255, 255, 0.8)",
            bordercolor="#e0e0e0",
            borderwidth=1,
            borderpad=4,
            font=dict(size=12)
        )

        # Save figure if path provided
        if save_path:
            import plotly.io as pio
            pio.write_html(fig, save_path)
            print(f"Neural state knowledge graph saved to {save_path}")

        return fig

def create_neurips_visualizations(model, inputs_dict, device, output_dir=None):
    """
    Create high-quality neural ensemble visualizations with the improved models.
    """
    print("\n" + "="*80)
    print("CREATING ENHANCED NEURIPS-QUALITY VISUALIZATIONS")
    print("How Minds Take Shape: Neural Ensembles Engineering Stable Representations")
    print("="*80 + "\n")

    # Create directory for visualizations
    if output_dir is None:
        output_dir = "enhanced_gamma_esn_results"

    neurips_dir = os.path.join(output_dir, 'neurips_visualizations')
    os.makedirs(neurips_dir, exist_ok=True)

    # Keep track of all visualizations paths
    viz_paths = {}

    # Generate all visualization types with enhanced aesthetics
    try:
        # 1. Enhanced electrode trajectories visualization
        fig_traj = visualize_electrode_trajectories_3d(
            model,
            inputs_dict,
            device,
            sample_idx=0,
            save_path=os.path.join(neurips_dir, 'electrode_trajectories_3d.html'),
            title="High-Dimensional Electrode Neural Trajectories"
        )
        viz_paths['trajectory'] = os.path.join(neurips_dir, 'electrode_trajectories_3d.html')
        print("Enhanced electrode trajectory visualization complete!")
    except Exception as e:
        print(f"Error creating electrode trajectory visualization: {e}")
        traceback.print_exc()

    try:
        # 2. Improved electrode influence network
        fig_network = create_electrode_influence_network(
            model,
            inputs_dict,
            device,
            sample_idx=0,
            save_path=os.path.join(neurips_dir, 'electrode_influence_network.html'),
            title="Electrode Cross-Influence Network"
        )
        viz_paths['influence_network'] = os.path.join(neurips_dir, 'electrode_influence_network.html')
        print("Enhanced electrode influence network visualization complete!")
    except Exception as e:
        print(f"Error creating electrode influence network: {e}")
        traceback.print_exc()

    try:
        # 3. Improved causal decision tree flow
        fig_causal = create_decision_tree_causal_flow(
            model,
            inputs_dict,
            device,
            sample_idx=0,
            save_path=os.path.join(neurips_dir, 'causal_decision_tree.html'),
            title="Causal Flow Between Neural Electrodes"
        )
        viz_paths['causal_tree'] = os.path.join(neurips_dir, 'causal_decision_tree.html')
        print("Enhanced causal decision tree visualization complete!")
    except Exception as e:
        print(f"Error creating causal decision tree: {e}")
        traceback.print_exc()

    try:
        # 4. Improved context vector evolution
        fig_context = visualize_context_vector_evolution(
            model,
            inputs_dict,
            device,
            sample_idx=0,
            save_path=os.path.join(neurips_dir, 'context_vector_evolution.html'),
            title="Unified Context Vector Evolution Over Time"
        )
        viz_paths['context_evolution'] = os.path.join(neurips_dir, 'context_vector_evolution.html')
        print("Enhanced context vector evolution visualization complete!")
    except Exception as e:
        print(f"Error creating context vector evolution: {e}")
        traceback.print_exc()

    try:
        # 5. Electrode-Agnostic Knowledge Graph with Linear Projections
        fig_knowledge_graph = create_enhanced_knowledge_graph(
            model,
            inputs_dict,
            device,
            sample_idx=0,
            save_path=os.path.join(neurips_dir, 'electrode_projection_knowledge_graph.html'),
            title="Electrode-Agnostic Knowledge Graph: Hidden State Projections"
        )
        viz_paths['knowledge_graph'] = os.path.join(neurips_dir, 'electrode_projection_knowledge_graph.html')
        print("Electrode projection knowledge graph visualization complete!")
    except Exception as e:
        print(f"Error creating electrode projection knowledge graph: {e}")
        traceback.print_exc()

    try:
        # 6. Electrode contributions to shifts in the unified vector
        fig_contrib = visualize_electrode_contributions_to_shifts(
            model,
            inputs_dict,
            device,
            sample_idx=0,
            save_path=os.path.join(neurips_dir, 'electrode_contributions_to_shifts.html'),
            title="Electrode Contributions to Context Vector Shifts"
        )
        viz_paths['contributions'] = os.path.join(neurips_dir, 'electrode_contributions_to_shifts.html')
        print("Electrode contributions to shifts visualization complete!")
    except Exception as e:
        print(f"Error creating electrode contributions visualization: {e}")
        traceback.print_exc()

    try:
        # 7. Comparative trajectory analysis between electrodes and unified vector
        fig_comparative = create_comparative_trajectory_analysis(
            model,
            inputs_dict,
            device,
            sample_idx=0,
            save_path=os.path.join(neurips_dir, 'comparative_trajectory_analysis.html'),
            title="Comparative Electrode Trajectory Analysis"
        )
        viz_paths['comparative'] = os.path.join(neurips_dir, 'comparative_trajectory_analysis.html')
        print("Comparative trajectory analysis visualization complete!")
    except Exception as e:
        print(f"Error creating comparative trajectory analysis: {e}")
        traceback.print_exc()

    try:
        # 8. 3D Rule-Based Decision Tree
        fig_rule_tree = create_electrode_rule_tree_3d(
            model,
            inputs_dict,
            device,
            sample_idx=0,
            save_path=os.path.join(neurips_dir, 'electrode_rule_tree_3d.html'),
            title="3D Electrode Hidden State Decision Rules"
        )
        viz_paths['rule_tree_3d'] = os.path.join(neurips_dir, 'electrode_rule_tree_3d.html')
        print("3D rule-based decision tree visualization complete!")
    except Exception as e:
        print(f"Error creating 3D rule-based decision tree: {e}")
        traceback.print_exc()

    try:
        # 9. Static Temporal Electrode Influence Network
        fig_temporal_influence = create_static_temporal_influence_network(
            model,
            inputs_dict,
            device,
            save_path=os.path.join(neurips_dir, 'static_temporal_influence_network.html'),
            title="Evolution of Electrode Influence Networks: Context to Prediction"
        )
        viz_paths['static_temporal_influence'] = os.path.join(neurips_dir, 'static_temporal_influence_network.html')
        print("Static temporal electrode influence network visualization complete!")
    except Exception as e:
        print(f"Error creating static temporal electrode influence network: {e}")
        traceback.print_exc()

    # FIXED NEW VISUALIZATIONS
    try:
        # 10. NEW & FIXED: Electrode Hidden State Contributions to Unified State
        fig_electrode_contributions = analyze_electrode_contributions(
            model,
            inputs_dict,
            device,
            sample_idx=0,
            save_path=os.path.join(neurips_dir, 'electrode_hidden_state_contributions.html'),
            title="Electrode Hidden State Contributions to Unified State"
        )
        viz_paths['electrode_contributions'] = os.path.join(neurips_dir, 'electrode_hidden_state_contributions.html')
        print("Electrode hidden state contributions visualization complete!")
    except Exception as e:
        print(f"Error creating electrode hidden state contributions visualization: {e}")
        traceback.print_exc()


    print("\nAll enhanced visualizations complete!")

    return viz_paths


def main_enhanced(data_path):
    """Main function to run the complete enhanced pipeline with neural ensemble analysis using the LFP-Grating dataset."""
    try:
        print("Starting Enhanced Gamma Oscillation Prediction Pipeline...")
        print(f"Using data path: {data_path}")

        # Create output directories
        os.makedirs(config.OUTPUT_DIR, exist_ok=True)
        os.makedirs(os.path.join(config.OUTPUT_DIR, 'visualizations'), exist_ok=True)
        os.makedirs(os.path.join(config.OUTPUT_DIR, 'neurips_visualizations'), exist_ok=True)

        # Save configuration for reproducibility
        import json
        config_dict = {k: v for k, v in config.__dict__.items() if not k.startswith('__') and not callable(v)}
        for k, v_item in config_dict.items():
            if isinstance(v_item, dict):
                config_dict[k] = {str(key_tuple): val_tuple for key_tuple, val_tuple in v_item.items()} \
                    if any(isinstance(kt, tuple) for kt in v_item.keys()) else v_item
            elif isinstance(v_item, (set, tuple)):
                config_dict[k] = list(v_item)
        with open(os.path.join(config.OUTPUT_DIR, 'config.json'), 'w') as f_cfg:
            json.dump(config_dict, f_cfg, indent=2)

        print("Loading electrode positions...")
        # Load electrode positions from probe_channels_810755799 (1).csv
        try:
            positions_file = find_data_file(data_path, 'probe_channels_810755799 (1).csv')
            positions = pd.read_csv(positions_file)

            # Rename columns to match the expected format in the code
            positions = positions.rename(columns={
                'channel_index': 'channel_num',
                'probe_horizontal_position': 'x_position',
                'probe_vertical_position': 'y_position'
            })

            # IMPORTANT: Filter positions to only include channels 0-94 which are in the data file
            max_channel = 94
            positions = positions[positions['channel_num'] <= max_channel].reset_index(drop=True)

            print(f"Loaded {len(positions)} electrodes (filtered to channels 0-{max_channel})")
            print("Columns in positions dataframe:", positions.columns.tolist())

        except Exception as e:
            print(f"Error loading electrode positions: {e}")
            traceback.print_exc()
            return None, None

        # Adjust NUM_ELECTRODES if needed
        if len(positions) < config.NUM_ELECTRODES:
            print(f"Warning: Only {len(positions)} electrodes available, reducing NUM_ELECTRODES from {config.NUM_ELECTRODES}")
            config.NUM_ELECTRODES = len(positions)

        print("Selecting electrodes...")
        # Select electrodes using KMeans clustering for optimal spatial coverage
        try:
            channel_nums, selected_indices = select_electrodes(positions, config.NUM_ELECTRODES)

            # Create visualization of the probe with selected electrodes
            visualize_probe(positions, selected_indices,
                          save_path=os.path.join(config.OUTPUT_DIR, 'visualizations', 'electrode_positions.png'))

            print(f"Selected {len(channel_nums)} electrodes out of {len(positions)}")
            print("Selected channel numbers:", channel_nums)

            # Additional verification that all channel numbers are within range
            if any(ch > max_channel for ch in channel_nums):
                print(f"Error: Some selected channels exceed the maximum available channel {max_channel}")
                invalid_channels = [ch for ch in channel_nums if ch > max_channel]
                print(f"Invalid channels: {invalid_channels}")
                return None, None

        except Exception as e:
            print(f"Error in electrode selection: {e}")
            traceback.print_exc()
            return None, None

        print("Loading LFP data...")
        # Load LFP data from drifting_gratings_full (1).csv
        try:
            # Find the data file
            data_file = find_data_file(data_path, 'drifting_gratings_full (1).csv')

            # Print top few lines of the CSV to inspect column names
            print("Inspecting data file column names...")
            import subprocess
            try:
                head_result = subprocess.check_output(['head', '-n', '1', data_file], text=True)
                print("CSV header:", head_result.strip())
            except:
                print("Could not run 'head' command, trying pandas read_csv with nrows=0")
                header_only = pd.read_csv(data_file, nrows=0)
                print("CSV columns:", header_only.columns.tolist())

            # Determine which columns to load (timestamp + selected channels)
            cols_to_use = ['timestamp', 'presentation_id'] + [f'channel_{ch}' for ch in channel_nums]
            print(f"Columns to use (first 5): {cols_to_use[:5]}...")

            # Load a subset of the data initially (adjust max_rows as needed based on memory)
            max_rows = 300000  # Start with a smaller subset, can increase later
            lfp_data = pd.read_csv(data_file, usecols=cols_to_use, nrows=max_rows)

            print(f"Loaded {len(lfp_data)} rows of LFP data with {len(cols_to_use)} columns")
            print(f"Time range: {lfp_data['timestamp'].min()} to {lfp_data['timestamp'].max()} seconds")

            # Check for NaN values
            nan_count = lfp_data.isna().sum().sum()
            if nan_count > 0:
                print(f"Warning: Found {nan_count} NaN values in the data")
        except Exception as e:
            print(f"Error loading LFP data: {e}")
            traceback.print_exc()
            return None, None

        print("Extracting time windows...")
        # Extract time windows for training and prediction
        try:
            # Extract windows from the continuous data
            windows = extract_windows(
                lfp_data,
                context_size=config.CONTEXT_SIZE,
                predict_size=config.PREDICT_SIZE,
                stride=config.STRIDE,
                max_windows=config.MAX_SEQUENCES
            )

            if windows.shape[0] == 0:
                print("No windows extracted. Cannot proceed.")
                return None, None

            print(f"Extracted {windows.shape[0]} windows with shape: {windows.shape}")

            # Save a sample window for inspection
            sample_window = windows[0]
            sample_window_df = pd.DataFrame(sample_window)
            sample_window_df.to_csv(os.path.join(config.OUTPUT_DIR, 'sample_window.csv'), index=False)

            # Plot a sample window
            plt.figure(figsize=(12, 6))
            plt.plot(sample_window[:, :5])  # Plot first 5 channels
            plt.title("Sample Window - First 5 Channels")
            plt.xlabel("Time Step")
            plt.ylabel("Amplitude")
            plt.savefig(os.path.join(config.OUTPUT_DIR, 'visualizations', 'sample_window.png'))
            plt.close()
        except Exception as e:
            print(f"Error extracting windows: {e}")
            traceback.print_exc()
            return None, None

        print("Preprocessing data...")
        # Preprocess the data with specialized spectral decomposition for gamma oscillations
        try:
            # Apply preprocessing with special attention to the very small amplitude values
            # Note: The signal values are very small (10^-6 to 10^-7), so we'll ensure proper scaling
            train_loader, val_loader, test_loader, scaler = preprocess_data(
                windows,
                context_size=config.CONTEXT_SIZE,
                predict_size=config.PREDICT_SIZE
            )

            if len(train_loader) == 0 or len(val_loader) == 0:
                print("Not enough data for training or validation. Batches are empty.")
                return None, None

            print(f"Prepared data loaders with {len(train_loader)} training batches, "
                  f"{len(val_loader)} validation batches, and {len(test_loader)} test batches")
        except Exception as e:
            print(f"Error in data preprocessing: {e}")
            traceback.print_exc()
            return None, None

        print("Creating enhanced gamma prediction model...")
        # Initialize and train the model
        try:
            # Create the EnhancedGammaESN model
            model = EnhancedGammaESN(config).to(device)
            print(f"Created model with {sum(p.numel() for p in model.parameters() if p.requires_grad):,} trainable parameters")

            # Train the model using the specialized training pipeline
            model, history = run_training_enhanced(model, train_loader, val_loader, config, test_loader)

            if model is None:
                print("Training failed. No model returned.")
                return None, None

            print("\nTraining completed successfully!")

            # Save the trained model
            torch.save(model.state_dict(), os.path.join(config.OUTPUT_DIR, 'models', 'final_model.pt'))
        except Exception as e:
            print(f"Error in model creation or training: {e}")
            traceback.print_exc()
            return None, None

        print("\nComplete pipeline execution successful!")

        # Save a summary of the pipeline execution
        with open(os.path.join(config.OUTPUT_DIR, 'execution_summary.txt'), 'w') as f:
            f.write("Enhanced Gamma Oscillation Prediction Pipeline\n")
            f.write("==============================================\n\n")
            f.write(f"Data path: {data_path}\n")
            f.write(f"Electrodes: {len(channel_nums)} selected from {len(positions)}\n")
            f.write(f"Data: {len(lfp_data)} samples with {len(cols_to_use)} columns\n")
            f.write(f"Windows: {windows.shape[0]} windows with shape {windows.shape}\n")
            f.write(f"Training batches: {len(train_loader)}\n")
            f.write(f"Validation batches: {len(val_loader)}\n")
            f.write(f"Test batches: {len(test_loader)}\n")
            f.write(f"Model parameters: {sum(p.numel() for p in model.parameters() if p.requires_grad):,}\n")
            f.write(f"NeurIPS visualizations created in: {os.path.join(config.OUTPUT_DIR, 'neurips_visualizations')}\n")
            f.write("\nExecution completed at: " + time.strftime("%Y-%m-%d %H:%M:%S"))

        return model, scaler

    except Exception as e_main:
        print(f"Error in main function: {e_main}")
        traceback.print_exc()
        return None, None


if __name__ == "__main__":
    try:
        data_path_resolved = None
        if os.path.exists("/kaggle/input/lfp-gabor-mk2"):
            data_path_resolved = "/kaggle/input/lfp-gabor-mk2"
        elif os.path.exists("/content/lfp-gabor-mk2"):
            data_path_resolved = "/content/lfp-gabor-mk2"
        else:
            try:
                import kagglehub
                print("Attempting to download dataset via kagglehub...")
                data_path_resolved = kagglehub.dataset_download("arunramponnambalam/lfp-gabor-mk2")
            except ImportError:
                print("kagglehub not available.")
            except Exception as e_kh:
                print(f"kagglehub download failed: {e_kh}. Trying local path.")

            if data_path_resolved is None:
                local_path_try = "./lfp-gabor-mk2"  # Changed to use new dataset name
                if os.path.exists(local_path_try):
                    data_path_resolved = local_path_try
                else:
                    data_path_resolved = input("Enter path to lfp-grating dataset directory: ")

        if data_path_resolved and os.path.exists(data_path_resolved):
            print(f"Using dataset at: {data_path_resolved}")
            model_trained, scaler_fitted = main_enhanced(data_path_resolved)

            # Print success message and visualization paths
            if model_trained is not None:
                print("\n"+"="*80)
                print("PIPELINE EXECUTION SUCCESSFUL")
                print("="*80)
                print("\nEnhanced NeurIPS visualizations are available at:")
                viz_dir = os.path.join(config.OUTPUT_DIR, 'neurips_visualizations')
                print(f"- Electrode Trajectories: {os.path.join(viz_dir, 'electrode_trajectories_3d.html')}")
                print(f"- Influence Network: {os.path.join(viz_dir, 'electrode_influence_network.html')}")
                print(f"- Causal Decision Tree: {os.path.join(viz_dir, 'causal_decision_tree.html')}")
                print(f"- Context Vector Evolution: {os.path.join(viz_dir, 'context_vector_evolution.html')}")
                print(f"- Electrode Contributions: {os.path.join(viz_dir, 'electrode_contributions_to_shifts.html')}")
                print(f"- Comparative Analysis: {os.path.join(viz_dir, 'comparative_trajectory_analysis.html')}")
        else:
            print(f"Dataset path not found or invalid: {data_path_resolved}")

    except Exception as e_script:
        print(f"Script execution error: {e_script}")
        traceback.print_exc()

# %% [code]
# Import necessary libraries (some might already be imported in the first cell)
import torch
import os
import pandas as pd
import numpy as np
from sklearn.preprocessing import RobustScaler
import matplotlib.pyplot as plt
import seaborn as sns
import umap.umap_ as umap # Import UMAP for dimensionality reduction
import matplotlib.colors as mcolors # For color mapping

model_path = os.path.join(config.OUTPUT_DIR, 'models', 'final_model.pt')

# Define the directory to save the analysis results, including the new heatmap
analysis_save_dir = os.path.join(config.OUTPUT_DIR, 'hidden_state_similarity_analysis_heatmap')
os.makedirs(analysis_save_dir, exist_ok=True)

print(f"Attempting to load model from: {model_path}")

# Check if the model file exists
if not os.path.exists(model_path):
    print(f"Error: Model file not found at {model_path}")
else:
    try:
        # Load the trained model
        # The EnhancedGammaESN class needs to be defined in the environment
        model = load_trained_model(model_path, config, EnhancedGammaESN)
        model.eval() # Set model to evaluation mode
        print("Model loaded successfully.")

        scaler_fitted = None # Set to your fitted scaler object if available

        print(f"Collecting hidden states for {10} samples across all timesteps...")

        # Call the function WITHOUT the 'timesteps_to_analyze' argument
        all_hidden_states, presentation_ids = analyze_hidden_state_similarities(
            model=model,
            data_path=data_path_resolved,
            config=config,
            scaler=scaler_fitted,
            num_samples=10, # Get 10 samples
            save_dir=os.path.join(analysis_save_dir, "pairwise_similarity_plots") # Save pairwise plots separately if generated
        )

        # --- Proceed with Heatmap Visualization if hidden states are collected correctly ---
        if all_hidden_states is not None and all_hidden_states.ndim == 3:
            num_presentations, num_timesteps, hidden_dim = all_hidden_states.shape
            print(f"Collected hidden states: Shape {all_hidden_states.shape}")

            # --- Apply UMAP for Dimensionality Reduction ---
            print("Applying UMAP for dimensionality reduction...")
            # Reshape for UMAP: (num_presentations * num_timesteps, hidden_dim)
            hidden_states_flat = all_hidden_states.reshape(-1, hidden_dim)

            # Initialize and fit UMAP
            # Adjust n_neighbors and min_dist as needed for your data
            umap_reducer = umap.UMAP(n_components=2, random_state=42, n_neighbors=15, min_dist=0.1)
            umap_embeddings = umap_reducer.fit_transform(hidden_states_flat)
            print(f"UMAP embeddings shape: {umap_embeddings.shape}")

            # --- Map UMAP Embeddings to Colors for Heatmap ---
            # Normalize embeddings to [0, 1] for colormap mapping
            umap_min = umap_embeddings.min(axis=0)
            umap_max = umap_embeddings.max(axis=0)
            # Add a small epsilon to avoid division by zero if max == min
            epsilon = 1e-8
            umap_normalized = (umap_embeddings - umap_min) / (umap_max - umap_min + epsilon)

            # Create an RGB array for the heatmap based on normalized UMAP coordinates
            colors_heatmap = np.zeros((num_presentations, num_timesteps, 3))

            for i in range(num_presentations):
                for j in range(num_timesteps):
                    # Get the normalized UMAP coordinates for presentation i, timestep j
                    flat_index = i * num_timesteps + j
                    u, v = umap_normalized[flat_index]

                    # Assign color based on normalized UMAP coordinates
                    # Using a simple mapping to RGB channels
                    colors_heatmap[i, j, 0] = u # Red channel from UMAP component 1
                    colors_heatmap[i, j, 1] = v # Green channel from UMAP component 2
                    colors_heatmap[i, j, 2] = 0.5 # Constant Blue channel (can be adjusted between 0 and 1)

            # --- Plot the Heatmap ---
            plt.figure(figsize=(14, 8)) # Adjust figure size as needed
            plt.imshow(colors_heatmap, aspect='auto', interpolation='nearest')
```
