## Supplementary material for "How Minds Take Shape: Graph-ESN Reveals How Neural Ensembles Engineer Stable Representations": wded.pdf

Anonymous Author(s)

This supplementary material provides detailed implementation specifications, extended experimental results, and additional analysis for the Graph-ESN architecture presented in the main paper.

### 1 Detailed Implementation of Enhanced Graph-ESN Architecture

#### 1.1 Population Structure Specifications

The model consists of 8 specialized neural populations, each containing 140 neurons, for a total of 1,120 neurons. The populations are structured as follows:

| Population Index | Specialization | Frequency Range (Hz) | Activation Function | Leaky Rate |
|---|---|---|---|---|
| 0 | Low Gamma | 30-50 | SiLU | 0.85 |
| 1 | Mid-Low Gamma | 50-80 | SiLU | 0.95 |
| 2 | Mid-High Gamma | 80-120 | SiLU | 1.05 |
| 3 | High Gamma | 120-180 | SiLU | 1.15 |
| 4 | Ultra Gamma I | 180-250 | SiLU | 1.25 |
| 5 | Ultra Gamma II | 250-300 | SiLU | 1.25 |
| 6 | Integrator | - | ELU | 0.70 |
| 7 | Context | - | Tanh | 0.50 |

The connectivity matrix between populations follows the pattern:

$$A_{i,j} = \begin{cases} \rho_{\text{intra}} & \text{if } i = j \\ \rho_{\text{inter}} \cdot f_{\text{influence}}(i,j) & \text{if } i \neq j \end{cases} \tag{1}$$

where $\rho_{\text{intra}} = 0.45$ is the intra-population connection density, $\rho_{\text{inter}} = 0.25$ is the inter-population connection density, and $f_{\text{influence}}(i,j)$ is the influence factor determined by the relationship between populations $i$ and $j$.

#### 1.2 Resonator Bank Implementation

For each gamma frequency band, we implemented a bank of coupled resonators. Each bank consists of 8 resonators distributed across the frequency range with the following specifications:

| Resonator Index | Center Frequency (Hz) | Q-Factor | Amplitude | Coupling Strength |
|---|---|---|---|---|
| 0 | 31.4 | 10 | 0.70 | 0.15 |
| 1 | 34.3 | 12 | 0.75 | 0.15 |
| 2 | 37.1 | 14 | 0.80 | 0.15 |
| 3 | 40.0 | 16 | 0.85 | 0.15 |
| 4 | 42.9 | 18 | 0.90 | 0.15 |
| 5 | 45.7 | 20 | 0.95 | 0.15 |
| 6 | 48.6 | 22 | 1.00 | 0.15 |
| 7 | 50.0 | 25 | 1.00 | 0.15 |

The coupled resonator dynamics are implemented as:

$$\ddot{x}_i + \frac{\omega_i}{Q_i}\dot{x}_i + \omega_i^2 x_i = F_i(t) + \sum_{j \neq i} c_{ij}(x_j - x_i) \tag{2}$$

where $\omega_i = 2\pi f_i$ is the angular frequency, $Q_i$ is the quality factor, $F_i(t)$ is the input force, and $c_{ij}$ is the coupling strength between resonators $i$ and $j$.

#### 1.3 GammaCellUnit Detailed Implementation

The GammaCellUnit integrates resonator dynamics with neural processing:

```
class GammaCellUnit(nn.Module):
    def __init__(self, input_size, hidden_size, band_freq=(30, 300)):
        super().__init__()
        self.input_size = input_size
        self.hidden_size = hidden_size
        self.band_freq = band_freq

        # Input projection
        self.input_proj = nn.Linear(input_size, hidden_size)

        # Recurrent weights
        self.recurrent_proj = nn.Linear(hidden_size, hidden_size)

        # Output gate
        self.output_gate = nn.Sequential(
            nn.Linear(hidden_size + input_size, hidden_size),
            nn.Sigmoid()
        )
```

```
        # Frequency modulation
        self.freq_mod = nn.Linear(input_size, 1)

        # Decay factor
        self.decay_factor = nn.Parameter(torch.tensor(0.1))

        # Excitatory-inhibitory balance
        self.ei_balance = nn.Parameter(torch.tensor(0.7))

        # Layer normalization
        self.layer_norm_hidden = nn.LayerNorm(hidden_size)

        # Resonator bank
        self.resonator = CoupledResonatorBank(
            freq_band=band_freq,
            num_resonators=8,
            coupling_strength=0.15
        )

        # Initialize hidden state
        self.register_buffer('hidden_state', None, persistent=False)
```

The forward pass of the GammaCellUnit is:

```
def forward(self, x):
    batch_size = x.shape[0]

    # Initialize state if needed
    if self.hidden_state is None or self.hidden_state.shape[0] != batch_size:
        self.reset_state(batch_size)

    # Input projection
    input_proj = self.input_proj(x)

    # Calculate frequency modulation
    freq_mod = torch.sigmoid(self.freq_mod(x))

    # Calculate recurrent contribution
    recurrent_contrib = self.recurrent_proj(self.hidden_state)

    # Apply E/I balance
    excitatory = F.relu(recurrent_contrib)
    inhibitory = F.relu(-recurrent_contrib)
    balanced_recurrent = self.ei_balance * excitatory - (1 - self.ei_balance) * inhibito
```

```
    # Combine input and recurrent for main pathway
    combined_main = input_proj + balanced_recurrent

    # Apply resonator dynamics
    resonator_input = torch.mean(combined_main, dim=1, keepdim=True)
    oscillation_scalar = self.resonator(resonator_input)
    gamma_contrib = freq_mod * oscillation_scalar.unsqueeze(1).expand(-1, self.hidden_si

    # Update hidden state with decay
    decay = torch.sigmoid(self.decay_factor)
    updated_hidden_candidate = combined_main + gamma_contrib
    normalized_hidden_candidate = self.layer_norm_hidden(updated_hidden_candidate)
    self.hidden_state = (1 - decay) * self.hidden_state + decay * normalized_hidden_cand

    # Calculate output gate
    output_gate_input = torch.cat([x, self.hidden_state], dim=1)
    gate_signal = self.output_gate(output_gate_input)

    # Apply output gate to the new hidden state
    gated_output = gate_signal * self.hidden_state

    return gated_output
```

### 2 Detailed Data Preprocessing

#### 2.1 Electrode Selection Process

The electrode selection procedure uses KMeans clustering to ensure optimal spatial coverage:

```
def select_electrodes(positions, num_electrodes):
    # Normalize positions for clustering
    x_norm = (positions['x_position'] - positions['x_position'].min()) /
             (positions['x_position'].max() - positions['x_position'].min())
    y_norm = (positions['y_position'] - positions['y_position'].min()) /
             (positions['y_position'].max() - positions['y_position'].min())
    coords = np.column_stack((x_norm.values, y_norm.values))

    # Use KMeans to select representative electrodes
    kmeans = KMeans(n_clusters=num_electrodes, random_state=42, n_init=10)
    kmeans.fit(coords)

    # Select the electrode closest to each cluster center
    selected_indices = []
    for cluster_idx in range(num_electrodes):
```

```
        cluster_points = np.where(kmeans.labels_ == cluster_idx)[0]
        center = kmeans.cluster_centers_[cluster_idx]
        distances = np.sqrt(np.sum((coords[cluster_points] - center)**2, axis=1))
        closest_idx = cluster_points[np.argmin(distances)]
        selected_indices.append(closest_idx)

    # Get channel numbers for selected electrodes
    channel_nums = positions.iloc[selected_indices]['channel_num'].astype(int).values

    return channel_nums, selected_indices
```

#### 2.2 Enhanced Frequency Band Decomposition

The band decomposition uses higher-order Butterworth filters in second-order sections format:

```
def create_enhanced_filter_bank(fs, freq_bands, order=8):
    filter_bank = []
    nyquist = fs / 2.0

    for band_idx, (low_freq, high_freq) in enumerate(freq_bands):
        # Normalize frequencies
        low = low_freq / nyquist
        high = high_freq / nyquist

        # Create higher order Butterworth filter for sharper cutoffs
        sos = signal.butter(order, [low, high], btype='bandpass', output='sos')
        filter_bank.append((sos, band_idx, (low_freq, high_freq)))

    return filter_bank

def extract_enhanced_frequency_bands(windows, fs, freq_bands):
    n_samples, window_size, n_channels = windows.shape
    n_bands = len(freq_bands)

    # Create filter bank
    filter_bank = create_enhanced_filter_bank(fs, freq_bands, order=8)

    # Initialize output array for filtered data
    filtered_data = np.zeros((n_samples, n_bands, window_size, n_channels))

    # Process each band with enhanced filters
    for (sos, band_idx, (low_freq, high_freq)) in filter_bank:
        # Process each sample and channel
        for i in range(n_samples):
```

```
        for j in range(n_channels):
            signal_i = windows[i, :, j]

            # Apply zero-phase filtering with second-order sections
            filtered = signal.sosfiltfilt(sos, signal_i)

            # Store filtered signal
            filtered_data[i, band_idx, :, j] = filtered

    return filtered_data
```

### 3 Detailed Training Methodology

#### 3.1 Curriculum Learning Implementation

The curriculum learning approach progressively focuses on different aspects of gamma oscillations:

```
class CurriculumScheduler:
    def __init__(self, config):
        self.config = config
        self.current_epoch = 0
        self.phase = 0
        self.num_phases = 3
        self.phase_length = config.CURRICULUM_PHASE_LENGTH

    def update(self, epoch):
        self.current_epoch = epoch
        self.phase = min(self.num_phases - 1, epoch // self.phase_length)

    def get_gamma_weights(self):
        base_weights = self.config.GAMMA_BAND_WEIGHTS.copy()
        if self.phase == 0:
            # Phase 1: Focus on low gamma bands (30-80 Hz)
            for (low, high), weight in base_weights.items():
                base_weights[(low, high)] = weight * 1.5 if low < 80 else weight * 0.5
        elif self.phase == 1:
            # Phase 2: Focus on mid-range gamma (50-180 Hz)
            for (low, high), weight in base_weights.items():
                base_weights[(low, high)] = weight * 1.5 if 50 <= low < 180 else weight
        elif self.phase == 2:
            # Phase 3: Focus on higher gamma (120-300 Hz)
            for (low, high), weight in base_weights.items():
                if low >= 120: base_weights[(low, high)] = weight * 1.5
        return base_weights
```

```
    def get_loss_weights(self):
        if self.phase == 0:
            return {'time': 0.5, 'freq': 0.5, 'phase': 0.0, 'envelope': 0.0}
        elif self.phase == 1:
            return {'time': 0.3, 'freq': 0.4, 'phase': 0.2, 'envelope': 0.1}
        else:
            return {'time': 0.25, 'freq': 0.45, 'phase': 0.2, 'envelope': 0.1}
```

#### 3.2 Enhanced Loss Function Implementation

```
class EnhancedGammaLoss(nn.Module):
    def __init__(self, config):
        super().__init__()
        self.config = config
        self.time_weight = config.TIME_LOSS_WEIGHT
        self.freq_weight = config.FREQ_LOSS_WEIGHT
        self.phase_weight = config.PHASE_LOSS_WEIGHT
        self.envelope_weight = config.ENVELOPE_LOSS_WEIGHT
        self.gamma_band_weights = config.GAMMA_BAND_WEIGHTS

        self.mse_loss = nn.MSELoss()
        self.phase_consistency_loss = PhaseConsistencyLoss()
        self.sampling_rate = float(config.SAMPLING_RATE)

        self.freq_band_map = {i: band for i, band in enumerate(config.FREQ_BANDS)}

    def gamma_frequency_loss(self, pred, target):
        batch_size, seq_len, num_channels = pred.shape

        # Reshape for FFT
        pred_reshaped = pred.permute(0, 2, 1).reshape(-1, seq_len)
        target_reshaped = target.permute(0, 2, 1).reshape(-1, seq_len)

        # Compute FFT
        pred_freqs, pred_mags, pred_phases = compute_fft_torch(pred_reshaped, self.sampl
        target_freqs, target_mags, target_phases = compute_fft_torch(target_reshaped, se

        # Initialize band weights tensor
        weights = torch.ones_like(pred_freqs, device=pred.device) * 0.1
        for band_idx, (low, high) in self.freq_band_map.items():
            band_mask = (pred_freqs >= low) & (pred_freqs <= high)
            band_weight_val = self.gamma_band_weights.get((low, high), 1.0)
            weights = torch.where(band_mask, torch.tensor(band_weight_val, device=weight
```

```
        # Weighted MSE for magnitude
        mag_diff_sq = (pred_mags - target_mags)**2
        weighted_mag_loss = torch.mean(mag_diff_sq * weights.unsqueeze(0))
        freq_loss = weighted_mag_loss

        # Phase consistency loss for gamma frequencies
        phase_loss_val = torch.tensor(0.0, device=pred.device)
        gamma_mask = pred_freqs >= 30
        if gamma_mask.any():
            pred_gamma_phases = pred_phases[:, gamma_mask]
            target_gamma_phases = target_phases[:, gamma_mask]
            if pred_gamma_phases.shape[-1] > 0:
                phase_loss_val = self.phase_consistency_loss(pred_gamma_phases, target_g

        # Envelope loss
        envelope_loss_val = torch.tensor(0.0, device=pred.device)
        if gamma_mask.any():
            pred_gamma_mags = pred_mags[:, gamma_mask]
            target_gamma_mags = target_mags[:, gamma_mask]
            if pred_gamma_mags.shape[-1] > 0:
                envelope_loss_val = F.mse_loss(pred_gamma_mags, target_gamma_mags)

        return freq_loss, phase_loss_val, envelope_loss_val

    def forward(self, pred, target):
        time_loss = self.mse_loss(pred, target)
        freq_loss, phase_loss, envelope_loss = self.gamma_frequency_loss(pred, target)

        total_loss = (
            self.time_weight * time_loss +
            self.freq_weight * freq_loss +
            self.phase_weight * phase_loss +
            self.envelope_weight * envelope_loss
        )
        return total_loss, time_loss, freq_loss, phase_loss, envelope_loss
```

### 4 Extended Experimental Results

#### 4.1 Performance Metrics Across Gamma Bands

| Gamma Band | MSE | $R^2$ | Time Correlation | Spectral Correlation | Phase Consistency |
|---|---|---|---|---|---|
| Low (30-50 Hz) | 0.0215 | 0.7845 | 0.8857 | 0.9062 | 0.6352 |
| Mid-Low (50-80 Hz) | 0.0312 | 0.7593 | 0.8616 | 0.8845 | 0.5981 |
| Mid-High (80-120 Hz) | 0.0384 | 0.7245 | 0.8337 | 0.8734 | 0.5608 |
| High (120-180 Hz) | 0.0451 | 0.6821 | 0.7853 | 0.8492 | 0.5126 |
| Ultra-I (180-250 Hz) | 0.0587 | 0.6247 | 0.7368 | 0.8195 | 0.4671 |
| Ultra-II (250-300 Hz) | 0.0724 | 0.5782 | 0.6972 | 0.7843 | 0.4312 |

##### 4.2 Detailed Ablation Study Results

| Model Configuration | Overall MSE | Overall $R^2$ | Avg. Gamma Correlation | Spectral Correlat |
|---|---|---|---|---|
| Full Model | 0.0241 | 0.7590 | 0.8121 | 0.8930 |
| w/o Resonator Banks | 0.0296 | 0.7043 | 0.7530 | 0.8320 |
| w/o Multi-Head Attention | 0.0305 | 0.6958 | 0.7402 | 0.8170 |
| w/o Phase Consistency Loss | 0.0278 | 0.7225 | 0.7683 | 0.8410 |
| w/o Specialized Populations | 0.0319 | 0.6817 | 0.7215 | 0.7950 |
| w/o Curriculum Learning | 0.0283 | 0.7173 | 0.7762 | 0.8540 |
| Standard ESN | 0.0389 | 0.6110 | 0.6823 | 0.7140 |

##### 4.3 Influence Analysis for Key Electrodes

| Electrode | Total Outgoing Influence | Total Incoming Influence | Major Transitions Triggered | Primar |
|---|---|---|---|---|
| E23 | 4.87 | 1.24 | 3 | Initi |
| E25 | 4.32 | 1.56 | 2 | Initi |
| E28 | 3.98 | 1.92 | 2 | Initi |
| E15 | 2.13 | 3.76 | 0 | Rece |
| E18 | 2.56 | 3.45 | 0 | Rece |

#### 5 Complete Hyperparameter Specifications

| Parameter Category | Parameter Name | Value |
|---|---|---|
| 7*Data Parameters | NUM_ELECTRODES | 64 |
| | CONTEXT_SIZE | 330 |
| | PREDICT_SIZE | 50 |
| | SAMPLING_RATE | 1000 |
| | MAX_SEQUENCES | 1000 |
| | BATCH_SIZE | 16 |
| | STRIDE | 50 |
| 6*Gamma Frequency Bands | Low Gamma | (30, 50) |
| | Mid-Low Gamma | (50, 80) |
| | Mid-High Gamma | (80, 120) |
| | High Gamma | (120, 180) |
| | Ultra High Gamma I | (180, 250) |
| | Ultra High Gamma II | (250, 300) |
| 7*Learning Parameters | LEARNING_RATE | 2e-4 |
| | WEIGHT_DECAY | 2e-5 |
| | EPOCHS | 50 |
| | PATIENCE | 15 |
| | CLIP_GRAD_NORM | 0.8 |
| | LR_WARMUP_EPOCHS | 5 |
| | GRADIENT_ACCUMULATION_STEPS | 2 |
| 7*Graph Reservoir Parameters | NUM_POPULATIONS | 8 |
| | NEURONS_PER_POPULATION | 140 |
| | SPECTRAL_RADIUS | 0.94 |
| | LEAKY_RATE | 0.75 |
| | INTER_POPULATION_DENSITY | 0.25 |
| | INTRA_POPULATION_DENSITY | 0.45 |
| | NOISE_LEVEL | 0.015 |
| 6*Oscillator Parameters | USE_OSCILLATORS | True |
| | OSCILLATION_STRENGTH | 0.7 |
| | OSCILLATOR_DAMPING | 0.05 |
| | ADAPTATION_RATE | 0.008 |
| | OSCILLATORS_PER_BAND | 8 |
| | COUPLING_STRENGTH | 0.15 |
| 9*Network Architecture Parameters | HIDDEN_SIZE | 192 |
| | READOUT_HIDDEN | 384 |
| | READOUT_LAYERS | 3 |
| | DROPOUT | 0.25 |
| | USE_ATTENTION | True |
| | ATTENTION_HEADS | 4 |
| | USE_SKIP_CONNECTIONS | True |
| | USE_LAYER_NORM | True |
| | USE_BATCH_NORM | True |
| 4*Loss Function Parameters | TIME_LOSS_WEIGHT | 0.25 |
| | FREQ_LOSS_WEIGHT | 0.75 |
| | PHASE_LOSS_WEIGHT | 0.4 |
| | ENVELOPE_LOSS_WEIGHT | 0.2 |
| 3*Curriculum Learning Parameters | USE_CURRICULUM | True |
| | CURRICULUM_START_EPOCH | 0 |
| | CURRICULUM_PHASE_LENGTH | 10 |

### 6 Computational Resources

All experiments were conducted using the following computational resources:

| Resource | Specification |
|---|---|
| Hardware | NVIDIA A100 GPU with 80GB VRAM<br>AMD EPYC 7742 CPU with 64 cores<br>512GB RAM |
| Software | Python 3.9<br>PyTorch 1.13.1<br>CUDA 11.7 |
| Training Time | 10.5 hours for full model<br>30 minutes for inference on test set<br>2 hours for visualization generation |

### 7 Data and Code Availability

The implementation code and trained models are available in the following GitHub repository:

`https://github.com/username/graph-esn-gamma`

The Allen Mouse Visual Coding Neuropixels dataset used in this work is publicly available at:

`https://portal.brain-map.org/explore/circuits/visual-coding-neuropixels`

The specific drifting gratings dataset can be accessed using the Allen SDK with the following code:

```
from allensdk.brain_observatory.ecephys.ecephys_project_cache import EcephysProjectCache

cache = EcephysProjectCache.from_warehouse(manifest=manifest_path)
sessions = cache.get_session_table()
stimulus_epochs = cache.get_stimulus_epochs()
drifting_gratings = stimulus_epochs[stimulus_epochs['stimulus_name'] == 'drifting_gratin
```

### 8 Extended Population Dynamics Analysis

#### 8.1 Population Contribution to Gamma Prediction

For a detailed analysis of how each specialized population contributes to gamma prediction, we examined the hidden state activations during prediction tasks. The activation patterns of different populations show distinct temporal dynamics for each gamma frequency band.

We quantified the contribution of each population to the overall prediction by analyzing the magnitude of weights connecting each population to the output layer:

| Population | Low Gamma | Mid-Low Gamma | Mid-High Gamma | High Gamma | Ultr |
|---|---|---|---|---|---|
| Pop 0 (Low Gamma) | 0.4872 | 0.1235 | 0.0587 | 0.0321 | 0.01 |
| Pop 1 (Mid-Low Gamma) | 0.1453 | 0.4512 | 0.1324 | 0.0654 | 0.03 |
| Pop 2 (Mid-High Gamma) | 0.0632 | 0.1567 | 0.4387 | 0.1287 | 0.05 |
| Pop 3 (High Gamma) | 0.0387 | 0.0654 | 0.1432 | 0.4265 | 0.12 |
| Pop 4 (Ultra-I) | 0.0213 | 0.0321 | 0.0587 | 0.1342 | 0.41 |
| Pop 5 (Ultra-II) | 0.0187 | 0.0256 | 0.0387 | 0.0587 | 0.12 |
| Pop 6 (Integrator) | 0.1234 | 0.0987 | 0.0876 | 0.0765 | 0.06 |
| Pop 7 (Context) | 0.1022 | 0.0468 | 0.0420 | 0.0779 | 0.16 |

#### 8.2 Cross-Population Information Flow

To analyze how information flows between populations, we calculated the cross-correlation between population states with various time lags. The following table shows the maximum correlation values and corresponding time lags:

| Source/Target | Pop 0 | Pop 1 | Pop 2 | Pop 3 | Pop 4 | Pop 5 | Pop 6 | |
|---|---|---|---|---|---|---|---|---|
| Pop 0 (Low Gamma) | 1.00/0 | 0.72/+3 | 0.54/+5 | 0.32/+8 | 0.21/+12 | 0.15/+15 | 0.48/-2 | [illegible] |
| Pop 1 (Mid-Low Gamma) | 0.68/-2 | 1.00/0 | 0.76/+2 | 0.48/+5 | 0.31/+8 | 0.21/+12 | 0.51/-3 | [illegible] |
| Pop 2 (Mid-High Gamma) | 0.49/-4 | 0.71/-2 | 1.00/0 | 0.75/+3 | 0.52/+6 | 0.34/+9 | 0.56/-4 | [illegible] |
| Pop 3 (High Gamma) | 0.28/-7 | 0.45/-4 | 0.72/-2 | 1.00/0 | 0.78/+3 | 0.53/+5 | 0.62/-5 | [illegible] |
| Pop 4 (Ultra-I) | 0.19/-11 | 0.29/-7 | 0.48/-5 | 0.74/-2 | 1.00/0 | 0.79/+2 | 0.58/-5 | [illegible] |
| Pop 5 (Ultra-II) | 0.12/-14 | 0.18/-11 | 0.31/-8 | 0.49/-4 | 0.76/-2 | 1.00/0 | 0.49/-6 | [illegible] |
| Pop 6 (Integrator) | 0.52/+3 | 0.57/+4 | 0.61/+5 | 0.67/+6 | 0.62/+6 | 0.54/+7 | 1.00/0 | [illegible] |
| Pop 7 (Context) | 0.46/+5 | 0.41/+6 | 0.45/+7 | 0.52/+8 | 0.58/+6 | 0.64/+5 | 0.74/-1 | [illegible] |

Positive lags indicate that the source population activity precedes the target population activity, suggesting causal influence. This analysis reveals a hierarchical information flow from lower to higher gamma bands, with the integrator and context populations showing bidirectional coupling with all gamma-specific populations.

#### 8.3 Electrode-Population Influence Analysis

We analyzed the influence of individual electrodes on specific populations using a partial correlation approach, controlling for the activity of all other electrodes:

| Population | Top Influencing Electrodes |
|---|---|
| Pop 0 (Low Gamma) | E23 (0.687), E25 (0.654), E28 (0.632), E32 (0.587), E35 (0.542) |
| Pop 1 (Mid-Low Gamma) | E25 (0.712), E28 (0.687), E32 (0.645), E35 (0.623), E38 (0.598) |
| Pop 2 (Mid-High Gamma) | E28 (0.732), E32 (0.701), E35 (0.684), E38 (0.643), E42 (0.621) |
| Pop 3 (High Gamma) | E15 (0.745), E18 (0.726), E21 (0.698), E23 (0.662), E25 (0.647) |
| Pop 4 (Ultra-I) | E18 (0.765), E21 (0.743), E23 (0.715), E25 (0.684), E28 (0.653) |
| Pop 5 (Ultra-II) | E15 (0.787), E18 (0.765), E21 (0.732), E23 (0.701), E25 (0.673) |
| Pop 6 (Integrator) | E23 (0.812), E25 (0.798), E28 (0.775), E32 (0.754), E35 (0.732) |
| Pop 7 (Context) | E15 (0.843), E18 (0.821), E21 (0.798), E23 (0.776), E25 (0.754) |

This analysis reveals specialized roles for different electrodes, with some electrodes showing strong influence on specific populations. In particular, electrodes E15, E18, and E21 strongly influence the higher gamma bands and context populations, while electrodes E23, E25, and E28 show stronger influence on the lower gamma bands and integrator population.

### 9 Additional Analysis Details

#### 9.1 Population-Specific Phase Space Analysis

We analyzed the phase space trajectories of each population by applying a time-delay embedding to their hidden states. This analysis reveals distinct attractor-like dynamics for different populations, with gamma-specific populations showing limit-cycle behavior characteristic of oscillatory dynamics, while integrator and context populations exhibit more complex, higher-dimensional attractors.

#### 9.2 Cross-Frequency Coupling Measures

We analyzed the cross-frequency coupling between different gamma bands using phase-amplitude coupling measures. The modulation index calculations reveal significant coupling between low and high gamma bands, with the strongest coupling observed between 40-60 Hz phase and 120-180 Hz amplitude, consistent with known physiological mechanisms.

#### 9.3 Enhanced Causal Flow Analysis

Our causal flow analysis uses Granger causality measures adapted for neural time series data. The analysis reveals strong causal influences flowing from specific initiator electrodes to receiver electrodes, with the pattern varying depending on the gamma frequency band under consideration.

### 10 Extended Experimental Results

#### 10.1 Performance Metrics Across Gamma Bands (Original Dataset)

| Gamma Band | MSE | $R^2$ | Time Correlation | Spectral Correlation | Phase Consistency |
|---|---|---|---|---|---|
| Low (30-50 Hz) | 0.0215 | 0.7845 | 0.8857 | 0.9062 | 0.6352 |
| Mid-Low (50-80 Hz) | 0.0312 | 0.7593 | 0.8616 | 0.8845 | 0.5981 |
| Mid-High (80-120 Hz) | 0.0384 | 0.7245 | 0.8337 | 0.8734 | 0.5608 |
| High (120-180 Hz) | 0.0451 | 0.6821 | 0.7853 | 0.8492 | 0.5126 |
| Ultra-I (180-250 Hz) | 0.0587 | 0.6247 | 0.7368 | 0.8195 | 0.4671 |
| Ultra-II (250-300 Hz) | 0.0724 | 0.5782 | 0.6972 | 0.7843 | 0.4312 |

#### 10.2 Detailed Ablation Study Results

| Model Configuration | Overall MSE | Overall $R^2$ | Avg. Gamma Correlation | Spectral Correlat[illegible] |
|---|---|---|---|---|
| Full Model | 0.0241 | 0.7590 | 0.8121 | 0.8930 |
| w/o Resonator Banks | 0.0296 | 0.7043 | 0.7530 | 0.8320 |
| w/o Multi-Head Attention | 0.0305 | 0.6958 | 0.7402 | 0.8170 |
| w/o Phase Consistency Loss | 0.0278 | 0.7225 | 0.7683 | 0.8410 |
| w/o Specialized Populations | 0.0319 | 0.6817 | 0.7215 | 0.7950 |
| w/o Curriculum Learning | 0.0283 | 0.7173 | 0.7762 | 0.8540 |
| Standard ESN | 0.0389 | 0.6110 | 0.6823 | 0.7140 |

#### 10.3 Performance on Additional Datasets (Graph-ESN Model)

The Graph-ESN model was also evaluated on two additional human neural datasets: the LFP-Gabor-MK2 dataset (referred to as "Gabor Dataset") and the ECoG-Landmark-MKN dataset (referred to as "Human Landmark Dataset"). The overall validation/test loss metrics for the Graph-ESN model on these datasets are presented below:

| Dataset | Overall Loss |
|---|---|
| Gabor Dataset (LFP-Gabor-MK2) | 0.028059 |
| Human Landmark Dataset (ECoG-Landmark-MKN) | 0.022899 |

#### 10.4 Model Performance Comparison on Additional Datasets

To further validate the efficacy of the Graph-ESN architecture, its performance was compared against several other established neural network models on the additional datasets. The comparison metrics, which were consistent across both the LFP-Gabor-MK2 and ECoG-Landmark-MKN datasets, demonstrate the superior performance of the proposed Graph-ESN model.

| Model<br>Test Spectrum Corr | Params | Best Val Loss | Val MSE | Val Corr | Val Spectrum Corr | Test MSE |
|---|---|---|---|---|---|---|
| RNN<br>0.2229 | 4,218,464 | 0.197768 | 0.059178 | 0.0048 | 0.2225 | 0.053239 |
| LSTM<br>0.1518 | 10,867,808 | 0.198031 | 0.057415 | 0.0043 | 0.0102 | 0.050908 |
| GRU<br>0.2539 | 8,651,360 | 0.196347 | 0.057319 | 0.0105 | 0.2523 | 0.051770 |
| Transformer<br>0.1825 | 2,671,904 | 0.196028 | 0.059813 | -0.0045 | 0.1823 | 0.053800 |
| PINN<br>0.1940 | 3,777,440 | 0.197526 | 0.057973 | 0.0106 | 0.1934 | 0.052803 |
| TCN<br>0.1371 | 2,670,176 | 0.196939 | 0.066750 | 0.0019 | 0.1415 | 0.061344 |
| WaveletNN<br>0.1653 | 6,052,192 | 0.196780 | 0.058083 | 0.0065 | 0.1649 | 0.052817 |

##### 10.4.1 Gamma Band Correlation Metrics (Test Set) on Additional Datasets

| Model | 30-50Hz | 50-80Hz | 80-120Hz | 120-180Hz | 180-250Hz | 250-300Hz | Average |
|---|---|---|---|---|---|---|---|
| RNN | 0.0316 | -0.0164 | -0.0083 | -0.0192 | -0.0127 | -0.0281 | -0.0089 |
| LSTM | -0.0075 | -0.0021 | 0.0043 | 0.0119 | -0.0070 | 0.0016 | 0.0002 |
| GRU | 0.0304 | -0.0279 | -0.0178 | -0.0003 | -0.0010 | -0.0197 | -0.0061 |
| Transformer | -0.0317 | 0.0209 | 0.0167 | 0.0206 | -0.0006 | 0.0097 | 0.0060 |
| PINN | -0.0110 | 0.0072 | 0.0100 | -0.0013 | -0.0047 | 0.0145 | 0.0024 |
| TCN | -0.0244 | 0.0027 | 0.0064 | -0.0017 | -0.0197 | 0.0053 | -0.0052 |
| WaveletNN | -0.0367 | 0.0188 | 0.0215 | 0.0260 | 0.0113 | 0.0448 | 0.0143 |

#### 10.5 Influence Analysis for Key Electrodes

| Electrode | Total Outgoing Influence | Total Incoming Influence | Major Transitions Triggered | Primar |
|---|---|---|---|---|
| E23 | 4.87 | 1.24 | 3 | Initia |
| E25 | 4.32 | 1.56 | 2 | Initia |
| E28 | 3.98 | 1.92 | 2 | Initia |
| E15 | 2.13 | 3.76 | 0 | Rece |
| E18 | 2.56 | 3.45 | 0 | Rece |

#### 10.6 Frequency-wise Loss (MSE) for Graph-ESN on Additional Datasets

The performance of the Graph-ESN model was analyzed for different gamma frequency bands on the Gabor Dataset and the Human Landmark Dataset. The Mean Squared Error

(MSE) for each band is presented below, demonstrating the model's performance across the frequency spectrum.

Frequency-wise MSE for Graph-ESN on Gabor Dataset

| Gamma Band | MSE |
|---|---|
| (30, 50) Hz | 0.002218 |
| (50, 80) Hz | 0.001686 |
| (80, 120) Hz | 0.000770 |
| (120, 180) Hz | 0.000468 |
| (180, 250) Hz | 0.000304 |
| (250, 300) Hz | 0.000165 |

Frequency-wise MSE for Graph-ESN on Human Landmark Dataset

| Gamma Band | MSE |
|---|---|
| (30, 50) Hz | 0.000250 |
| (50, 80) Hz | 0.000298 |
| (80, 120) Hz | 0.000665 |
| (120, 180) Hz | 0.000960 |
| (180, 250) Hz | 0.001047 |
| (250, 300) Hz | 0.000747 |
